# Subsistence transition preceded population turnover in the eastern Colombian Andes

**DOI:** 10.64898/2026.03.23.713713

**Authors:** Kendra Sirak, Miguel Delgado, Angélica Triana, Sebastián Rivas, Pedro Argüello, Ana María Boada, Javier Rivera-Sandoval, German Peña, Carl Langebaek, Juan Pablo Ospina, Sonia Archila, Saúl Alberto Torres Orjuela, Martha Beatriz Mejía Cano, Freddy Rodríguez Saza, Alison Barton, Kim Callan, Elizabeth Curtis, Trudi Frost, Lora Iliev, Aisling Kearns, Jack Kellogg, Ann Marie Lawson, Lijun Qiu, J. Noah Workman, Matthew Mah, Mariam Nawaz, Gregory Soos, Alexander Cherkinsky, Carla S. Hadden, Keith M. Prufer, Swapan Mallick, Nadin Rohland, Lars Fehren-Schmitz, David Reich

## Abstract

Northwest South America was a pivotal region for human dispersals and cultural exchange during the Holocene. The Altiplano Cundiboyacense, a high-altitude plateau in the Eastern Cordillera of the Northern Andes of Colombia, preserves one of the most continuous archaeological sequences in the Americas, spanning from late Pleistocene hunter-gatherer groups to final late Holocene Muisca chiefdoms. Increasing the regional ancient DNA sample size 11-fold, we report genome-wide data from 209 individuals who lived over a period of more than 7000 years. This includes hunter-gatherers from the early-middle (10,000-7000 BP) and middle (7000-4000 BP) Holocene, initial late Holocene people (4000-2500 BP) who have the first isotopic evidence of C₄-enriched diets (attributed to maize), and populations associated with increasing sedentism and food production in the Herrera (2200-1300 BP) and Muisca (1200-500 BP) Periods. Previous work identified a major population turnover distinguishing earlier groups from Herrera-Muisca Period populations, but the absence of individuals dating 6000-2000 BP in that study left unresolved whether this ancestry shift was gradual or abrupt and whether it accompanied the earliest isotopic evidence of dietary input from maize or coincided with the later emergence of Herrera culture. We show that individuals predating the Herrera Period form a lineage that persisted for over five millennia, with population structure driven by drift in small groups and no detectable external gene flow. Two individuals who lived ∼2800 years ago – one directly dated to 983-835 calBCE – exhibit genetic profiles entirely consistent with hunter-gatherer ancestry yet have isotopic values consistent with the incorporation of maize into their diets, indicating subsistence change without population replacement. The emergence of Herrera culture ∼2200 BP coincided with a sharp genetic break, reflecting the migration of people carrying ancestry diverged by up to ten millennia into the Sabana de Bogotá and displacing previously established peoples. By co-analyzing ancient data with modern Native Americans, we show these later populations derived from a mixture ∼4000 years ago of groups related to Chibchan language speakers of lower Central America and ones related to present-day people at the Amazonian-Andean interface who may have lived along the Chibchan expansion route. In the Herrera and Muisca Periods, genetic substructure distinguishes people from the southern and northern Altiplano, consistent with the cultural differentiation of these regions in the archaeological record.

**IN BRIEF:** Ancient DNA data from the eastern Colombian Andes reveal five millennia of population continuity during which C₄ plants were incorporated into subsistence systems without population replacement, followed later by a major ancestry turnover involving a population with ancestry admixed between that found in Chibchan-related groups and at the Amazonian-Andean interface.

## INTRODUCTION

Northwest South America is the main gateway into the continent from Central America and has been an important setting for demographic and cultural transformations throughout the Holocene. Within this region, some of the best documented archaeological sequences are found in present-day Colombia. Human occupation is documented by the late Pleistocene (∼13,000 years before present, BP) when groups equipped with diverse lithic technologies and broad-spectrum economies exploited the open environments formed during the transition from a warm and wet climate to cooler and drier conditions^1–8^. Over time, they settled diverse landscapes including the coastal Caribbean lowlands, the Orinoco and Amazon River basins, the inter-Andean valleys, and the Altiplano Cundiboyacense.

The Altiplano Cundiboyacense – a high-altitude plateau in the Eastern Cordillera of the northern Andes that includes the Sabana de Bogotá in the south and the highlands of Boyacá in the north – has archaeological evidence of human habitation since the late Pleistocene^4,5,9,10^, although the earliest skeletal remains date to the early-middle Holocene (10,000-7000 BP). This was a period of climatic amelioration that may have fostered demographic expansion, generalized diets and economies, and inter-regional exchange among hunter-gatherer groups^1,6,11–13^. People relied mostly on C_3_ plants with flexible use of animal protein^12,14,15^. Warmer and drier conditions during the middle Holocene (7000-4000 BP, the hypsithermal interval) have been linked to the fragmentation of forested areas and possible demographic contraction and dispersals of resident hunter-gatherers^11,16^. Zooarchaeological^17^ and isotopic^12^ evidence indicates continued exploitation of C_3_ plants and increased emphasis on guinea pig (*Cavia* sp.) and to a lesser extent white-tailed deer (*Odocoileus virginianus*). Archaeobotanical evidence from middle Holocene sites including Checua, Aguazuque, and Tequendama (**Supplementary Information 1**) documents a broad-spectrum subsistence economy centered on high-Andean tubers and legumes, with limited incorporation of maize (*Zea mays*) at Checua evidenced by starch grains recovered from plant-processing tools and human dental calculus^18^. The cooler and wetter conditions of the early initial late Holocene (4000-2500 BP) facilitated shifts in subsistence practices, including the first isotopic evidence of C₄-enriched resources in human diets^1,8,12,19–21^. Some research has proposed population continuity from the late Pleistocene through the arrival of sedentary farmers on the Altiplano Cundiboyacense in the later initial late Holocene (∼2200 BP), citing the persistence of the Abriense lithic industry and attributing craniodental changes to *in situ* adaptation^20,22–27^. Other research suggests that non-local people migrated to the region during the middle Holocene and mediated dietary shifts and craniofacial changes^1,12,16,21,28,29^. Exchange networks connecting the Sabana de Bogotá with neighboring regions including the Middle Magdalena Valley, Atlantic coast, Cauca Valley, and southwestern Colombia could have been conduits for cultural transformation and gene flow^14,16,22,30,31^, while the appearance of C₄-enriched resources in local food webs could reflect population dispersals across these networks^28^.

Around 2500 BP, there is archaeological evidence of emerging complex sedentary societies on the Altiplano Cundiboyacense, and the period spanning ∼2200-1300 BP is known as the Herrera Period^32,33^. It is characterized by longer-term occupations, cultivation of maize along and other crops, and ceramic production. Archaeological, skeletal, and ancient DNA data indicate that the ancestors of Herrera populations migrated to the region, and connect the incoming population to the spread of Chibchan-speakers from a homeland in lower Central America throughout the Isthmo-Colombian sphere stretching between eastern Honduras and northern Colombia^34–36^. Based on typological similarities between the earliest Herrera pottery tradition (*Mosquera Rojo Inciso*) on the Altiplano Cundiboyacense and the *Arrancaplumas* tradition from the Magdalena River Valley, some archaeologists have hypothesized that Herrera ancestors entered South America and moved southward via the Magdalena River Valley, establishing the *Arrancaplumas* tradition in the lowlands before 2500 BP and expanding to the Sabana de Bogotá via the Magdalena River corridor between 2500-2000 BP, after which they maintained connections with communities who remained there^37,38^. However, *Arrancaplumas* also shares affinities with *Mosquera Triturada,* another variant of early Herrera pottery found on the Altiplano Cundiboyacense, inviting further evaluation of migratory models based on ceramic evidence^33,39,40^.

In the final late Holocene (1200-500 BP), cultural and sociopolitical transformations led to what is known archaeologically as the Muisca Period^32,33^. Studies of skeletal and dental morphology^16,25,26,28,29,41,42^ and ancient DNA^5,36^ documents a close biological relationship between Herrera and Muisca Period people, suggesting population continuity despite cultural shifts^29^. The Late Herrera and Early Muisca Periods were characterized by sustained population growth^33^, a shift in settlement patterns to larger and denser household clusters^43^, and changes in social organization^44,45^, while the Late Muisca Period was characterized by further population expansion^33^, the expansion of chiefly sociopolitical organization, growth in trade networks, craft specialization, and the expansion of settlements located close to fertile agricultural fields^33,46^. At the time of Spanish contact, there were a number of large and socially complex Muisca chiefdoms that that participated in shared religious ceremonies, alliances, and conflicts^44^, spoke Chibchan dialects, and practiced avunculocal post-marital residence and matrilineal political succession^44,45^.

Although the Altiplano Cundiboyacense has yielded abundant human skeletal material suitable for paleogenomic investigation, genome-level ancient DNA from the region remains limited. A previous study^36^ demonstrated that people of the Herrera and Muisca Periods were genetically distinct from a group who lived ∼6000 BP; however, a sampling gap between ∼6000-2000 BP prevented resolution of when, how abruptly, and in association with which subsistence or cultural changes this ancestry shift occurred. As a result, three hypotheses remain to be tested: (1) that the ancestry transition occurred in the middle Holocene (∼6000-4000 BP) during a period primarily featuring C_3_ plant economies; (2) that it occurred in the initial late Holocene (∼4000-2500 BP), when the first clear isotopic signal of C₄-enriched resources appears in human diets; or (3) that it was associated with a later migration of people establishing more sedentary communities and expanding food production, corresponding to the appearance of the Herrera culture.

## RESULTS

We generated genome-wide data using skeletal elements from 240 ancient individuals from the Altiplano Cundiboyacense, enriching for ∼1.4M single nucleotide polymorphisms (SNPs) across the genome and the mitochondrial genome (**Supplementary Information 1**). We report results for 209 individuals with data meeting standards for ancient DNA authenticity (**Figure 1**; **Table S1**; Method Details). We also generated 37 new direct ^14^C dates on bone samples and dietary isotope data for 33 individuals (**Supplementary Information 2**; Method Details). Based on archaeological evidence, strata (layer) dates, or direct ^14^C dates on bone samples, we divided 45 individuals associated with Holocene hunter-gatherer or early food cultivation contexts into three temporal categories: early-middle Holocene, dating 7000 BP or older (n=2); middle Holocene, dating 6000-4000 BP (n=31); and initial late Holocene, dating 4000 BP or more recent (n=12) (**Table S1**). These temporal divisions mark intervals characterized by climatic, cultural, or biological shifts and have been used in previous work^5,12,16,29^. A total of 164 individuals were associated with food-producing contexts, including 18 from the initial late Holocene Herrera Period (∼2200-1300 BP) and 146 dating to the final late Holocene (∼1200-500 BP), encompassing both the Early and Late Muisca Periods. We co-analyzed new data with 21 previously published genomes^36^, including 7 hunter-gatherers from Checua dating ∼6000 BP, 9 Herrera Period individuals from Laguna de la Herrera dating ∼2000 BP, 3 individuals from Las Delicias and Soacha dating to the Early and Late Muisca Periods, and 2 individuals who lived ∼530 BP at the site of Purnia located north of the Altiplano Cundiboyacense in the Santander Department.

**Figure 1.**
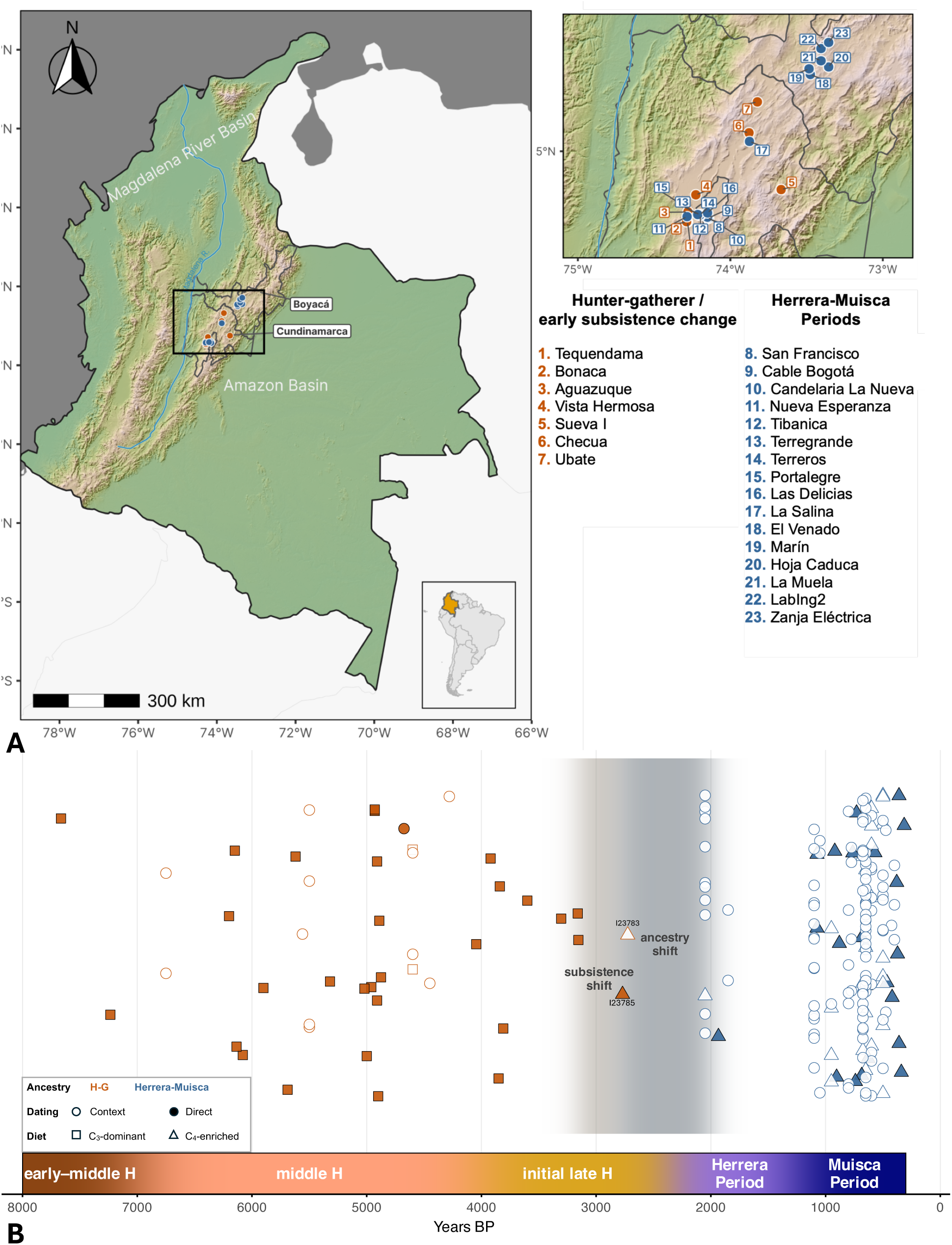
Overview of ancient individuals with new genome-level data in this work. A) Map of sites. Pre-Herrera sites associated with hunter-gatherers or early C_4_ inputs and people with fully hunter-gatherer-associated ancestry are labeled in orange, and sites associated with Herrera-Muisca contexts are labeled in blue. The departments of Cundinamarca and Boyacá, part of present-day Colombia, are outlined and labeled. B) Timeline illustrating the temporal distribution of samples and shifts in subsistence strategy and ancestry that took place between the early-middle and final late Holocene (H). Individuals I23783 and I23785 from initial late Holocene Aguazuque who have a genetic profile entirely consistent with hunter-gatherer ancestry and isotopic values indicative of consuming C₄ resources are labeled. Legend is on the left side of the plot.

For an overview of genetic variation in our data, we carried out Principal Component Analysis (PCA) as in ref. ^47^ (**Figure 2**; **Figures S1** and **S2**; Method Details). Ancient people from the Altiplano Cundiboyacense fall into two clusters corresponding to time period. The “H-G Cluster” includes individuals from contexts spanning over 5000 years from the early-middle Holocene (I28092 from Sueva I is directly dated to 6567-6437 calBCE) through the initial late Holocene (I23785 from Aguazuque is directly dated to 983-835 calBCE). This cluster overlaps some published ancient individuals from South America including from Peru, Chile, and Argentina. The “Herrera-Muisca Cluster” is shifted in the direction of present-day Chibchan-speakers from Costa Rica and ancient people from Panama who have ancestry like that of present-day Chibchan-speaking groups (hereafter referred to as “Chibchan-related ancestry”), although it does not overlap them. All Herrera Period individuals – directly dated as early as 1975 BP^36^ and associated with context dates as early as 2450 BP – fall in this cluster, consistent with a complete shift in ancestry by this time. The clustering of Herrera Period individuals with people from the Muisca Period is suggestive of population continuity from the end of the initial late Holocene through the final late Holocene. Outside of the clusters, two outliers with >15k SNPs are a middle Holocene hunter-gatherer shifted in the direction of Mesoamericans (I23673) and a Late Muisca Period individual (I10932) shifted toward the H-G Cluster and other South Americans.

**Figure 2.**
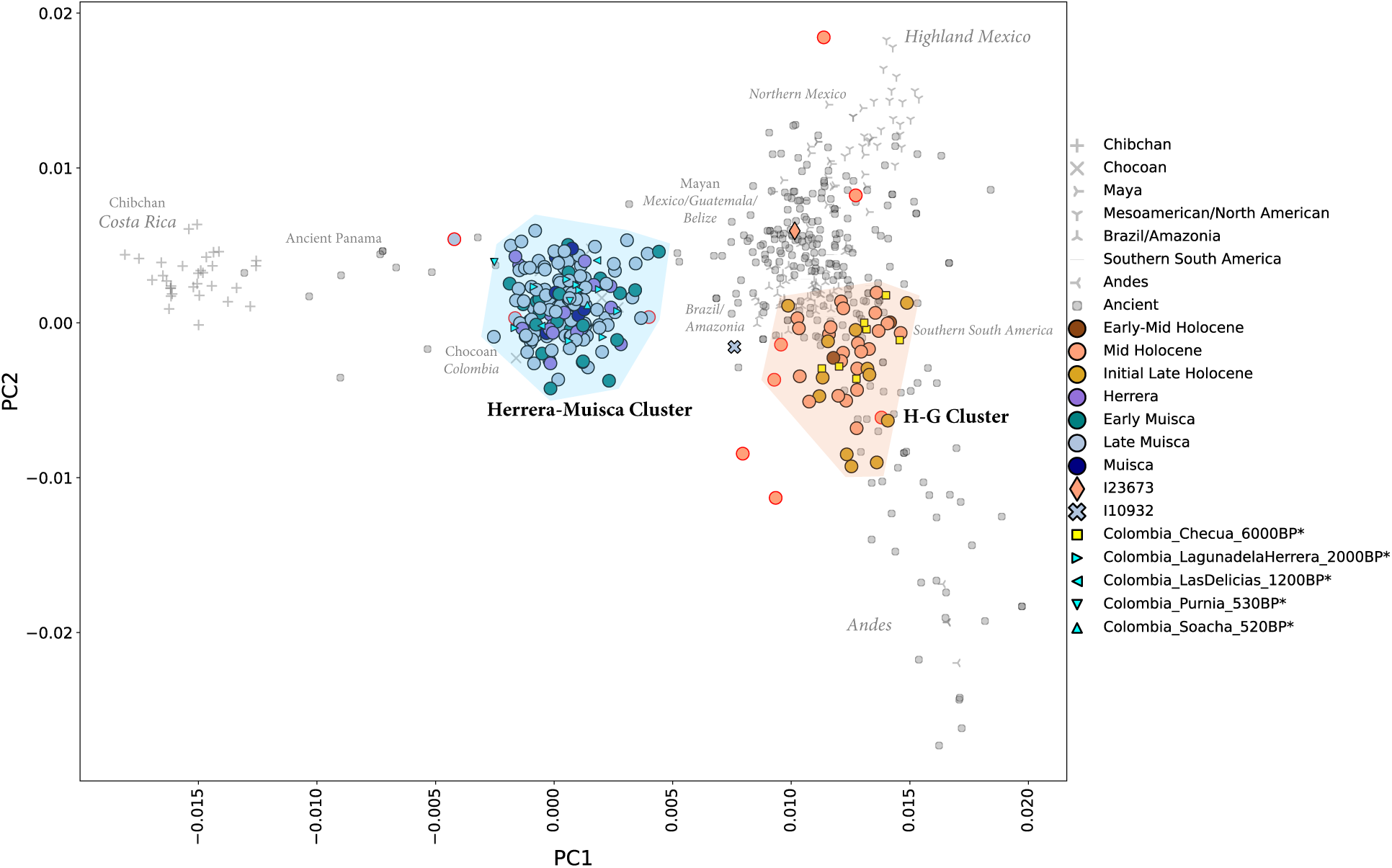
PCA shows two clusters of ancient people from the Altiplano Cundiboyacense. Reference data are in gray (see **Supplementary Figure S2** for a reference data legend). Newly reported data and published ancient Colombian data^36^ represented by large and small colored symbols, respectively. An asterisk next to a sample name in the legend indicates a sample published in ref. ^36^. A red outline denotes low coverage individuals (<15K SNPs covered on the 2M.Illumina dataset). Two PCA outliers (I23673 and I10932) with >15K SNPs are labeled with diamond and X symbols, respectively.

We used pairwise *qpWave* to group newly reported and recently published individuals from the Altiplano Cundiboyacense relative to known genetic diversity in the Americas (**Figure S3**; **Supplementary Information 3**; Method Details). Using a multistep approach, we find relative genetic homogeneity among most individuals who fall in the “H-G Cluster” on PCA within the limits of our resolution. We grouped these people for some analyses as Colombia_SB_HG (‘SB’ to denote the Sabana de Bogotá, and ‘HG’ to denote ancestry associated with pre-Herrera Period populations), excluding three individuals with ambiguous *qpWave* model fits (**Table S2**; **Supplementary Information 3** and **4**) and one lower-coverage first-degree relative of another individual in our dataset. *qpWave* did not support genetic differentiation of PCA outlier I23673 relative to other Colombia_SB_HG individuals, indicating that its PCA position may be driven by technical factors rather than ancestry differences.

Consistent with PCA, *qpWave* confirms genetic differentiation between individuals from pre-Herrera and Herrera-Muisca Period contexts, supporting a major ancestry shift. At the same time, people from Herrera and Muisca contexts appear largely genetically homogeneous aside from a few exceptions (**Supplementary Information 3**). One such case is individual I10932 – a male who we directly date to 1422-1454 calCE – who is both a PCA outlier and exhibits reduced genetic affinity to other Herrera-Muisca Period individuals in *qpWave*, particularly with respect to Chibchan-associated groups (**Supplementary Information 4**; **Tables S3**-**S6**). We tested for cladality at all Herrera-Muisca Period sites with multi-period occupations spanning from the Herrera through Late Muisca Periods (El Venado, La Muela, Nueva Esperanza) or from the Early through Late Muisca Periods (Terreros) and found no evidence of temporal substructure (all groups within each form a clade with p>0.05) (**Supplementary Information 3**). Statistics of the form *f_4_*(Yoruba, *NativeAmerican_Test*; *SiteAPeriodA*; *SiteAPeriodB*) are not significantly different from zero, supporting genetic continuity between the Herrera and Muisca Periods at each site and providing evidence against demic movement driving cultural transformation in this time (**Table S7**).

While there is no evidence of temporal sub-structure between the Herrera, Early Muisca, and Late Muisca Periods, we detect geographic sub-structure among Herrera-Muisca Period people with *qpWave*, whereby people from the southern Altiplano (the present-day department of Cundinamarca) share more genetic affinity with each other than with people from the northern Altiplano (the highlands of present-day Boyacá). We observe additional subtle sub-structure within Boyacá, with people from the sites of Zanja Eléctrica, LabIng2, and Marín forming a group that also includes the individual from La Salina (located in Cundinamarca in the direction of Boyacá relative to other sites, **Figure 1A**) and those from Los Curos (located in southern Santander), while people from La Muela, El Venado, and Hoja Caduca form a second Boyacá cluster. We therefore created three regional groups for subsequent analyses: “Colombia_Cundinamarca”, “Colombia_Boyacá1_LaSalina”, and “Colombia_Boyacá2”. In what follows, we focus primarily on reconstructing the ancestry of the Colombia_SB_HG group and the three regional Herrera-Muisca Period groups, with individuals not incorporated into these broader groups discussed in **Supplementary Information 4**.

### Five millennia of population continuity on the Sabana de Bogotá

Leveraging time-series data from 52 pre-Herrera Period individuals spanning more than five millennia, we asked: 1) Do the genetic relationships among the early groups of the Sabana de Bogotá reflect structure arising from drift or is there evidence for the entrance of non-local ancestry during this time?; and 2) What is the genetic relationship between these people and other ancient Central and South Americans?

We constructed a neighbor-joining (NJ) tree using inverted outgroup-*f_3_* statistics to investigate genetic relationships among Sabana de Bogotá individuals with hunter-gatherer-associated ancestry and between them and other Native American lineages (**Figure 3**; Method Details). These individuals from the Sabana de Bogotá form a distinct genetic clade within the broader Central and South American variation^36^, with sub-clades structured by both temporal and geographic factors. The oldest individual in our dataset (I28092), who we directly date to 6567-6437 calBCE, occupies the most basal position in this clade, suggesting that this early-middle Holocene individual lacks some of the genetic drift shared by later individuals. This highlights I28092 as a key reference point for understanding the earliest Holocene genetic diversity in the region and exploring possible shifts during the middle and initial late Holocene occupations. All remaining pre-Herrera Period people from the Sabana de Bogotá fall into three sub-clades. The first includes individuals from early-middle and middle Holocene Tequendama as well as from middle Holocene Bonacá and Ubaté and I23673 from Aguazuque, suggesting that some early-middle Holocene ancestry persisted into later periods. The second comprises all individuals from middle Holocene Checua, consistent with relatively homogeneous ancestry at that site. The final sub-clade comprises all initial late Holocene individuals from Vista Hermosa and Aguazuque as well as most middle Holocene individuals from Aguazuque, reflecting continuity at that site over time and a unique genetic profile relative to middle Holocene people from other sites. I22318 and I23671, assessed as genetically less similar than others from their same site with *qpWave*, were placed in sub-clades with other individuals from their same site and time period.

**Figure 3.**
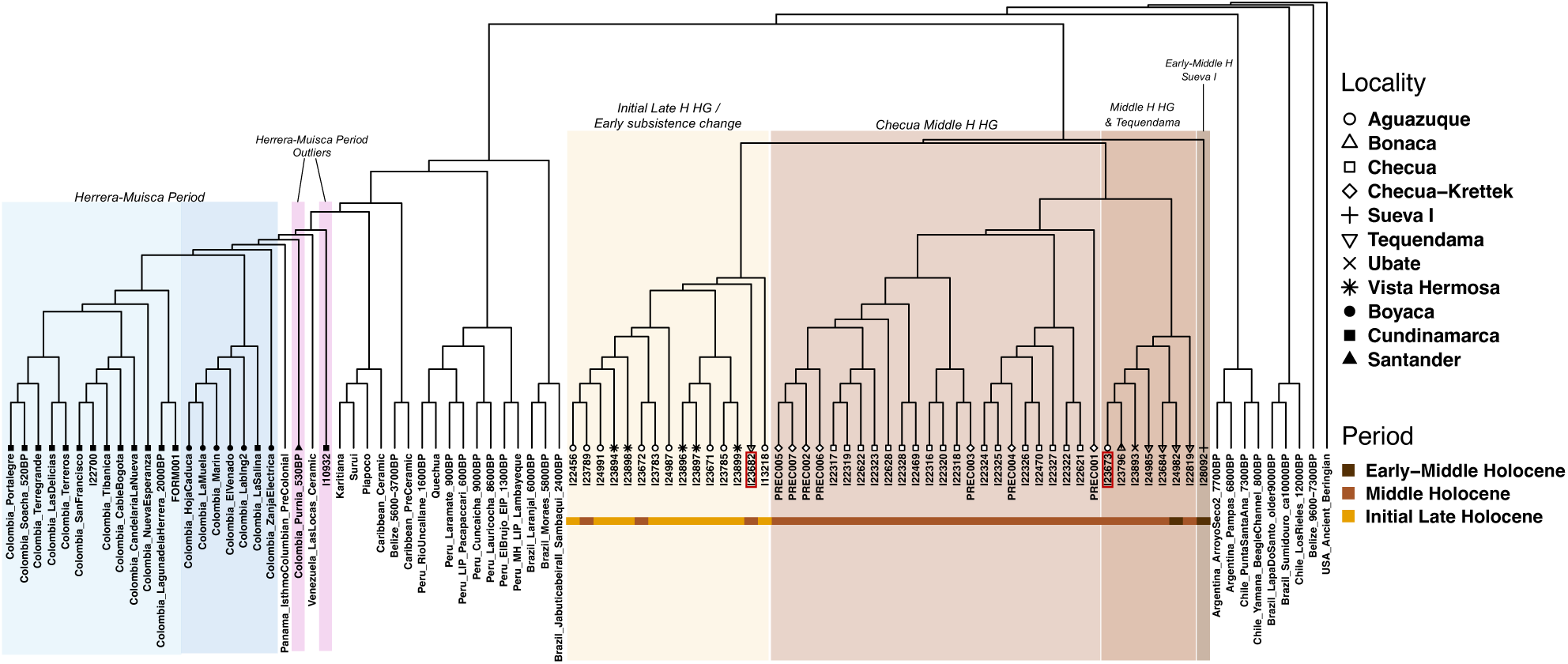
Neighbor-joining (NJ) tree based on inverted outgroup*-f_3_* statistics. We highlight sub-clades of relevance to this work in shaded boxes for easy identification, with individuals from Herrera-Muisca contexts in blue tones (sites from Cundinamarca in light teal and sites from Boyacá in light royal blue) and purple (outliers) and Sabana de Bogotá hunter-gatherers or people from contexts with early C_4_ inputs in yellow and brown tones (initial late Holocene group in yellow, Checua middle Holocene HG group in light brown, middle Holocene HG and Tequendama group in brown, and Sueva I HG in dark brown). Two individuals who are not placed in clades with other individuals from their same site and time periods are identified with red boxes. Top right legend denotes localities and bottom right legend identifies period for the colored strip below individuals from hunter-gatherer and early food producing contexts. ‘Checua-Krettek’ denotes previously published individuals^36^. While Colombia_Purnia_530BP is identified as an outlier in this tree, this group clusters with sites in the department of Boyacá in *qpWave*.

We tested whether we could detect ancestry differences between these sub-clades or whether all are consistent with belonging to the same ancestral population with the sub-structure in the NJ tree fully attributable to genetic drift that accumulates between temporally and spatially separated groups. All tests of the form *f_4_*(Yoruba, *NativeAmerican_Test*; *Colombia_SB_HG_SubClade1*, *Colombia_SB_HG_SubClade2*) returned values indistinguishable from zero, suggesting that every *NativeAmerican _Test* group is equally related to each sub-clade with no signal of asymmetric allele sharing that would indicate gene flow or closer affinity to one sub-clade over another. The only significant statistics (|Z|>3) reaffirm genetic sub-structure among the sub-clades (**Table S8**). Likewise, outgroup-*f₃* values were similar across sub-clades, indicating no meaningful variation in shared genetic drift with other American groups (**Table S9**; **Figure S4**; Method Details).

Our data are consistent with a scenario of genetic continuity on the Sabana de Bogotá from the early-middle Holocene through early initial late Holocene, with no evidence supporting the entry of non-local ancestry during this time. Although we detect sub-structure between sites and time periods, formal tests show this pattern is consistent with genetic drift in a small population rather than external gene flow. Indeed, we estimate effective community sizes of 110-133 (95% confidence interval fitted via the likelihood profile), reflecting the strong drift expected in small, structured groups (Method Details). Despite a very small pool of ancestors, mating between closely related individuals did not take place, with only six pre-Herrera individuals out of 23 tested having any Runs of Homozygosity (ROH) >20cM indicating close-kin unions (**Table S10**; **Figure S5**; Method Details). Consistent with a small community, we identify a three-generation family buried at Checua spanning ∼3765-3630 BP with reproductive partners I22622 and I22323 buried together (the latter, a female, representing the only primary burial) ^48^ while their son (I22317) and his daughter (I22319) were co-buried as part of a separate triple burial (**Supplementary Information 5**).

The ancestry of pre-Herrera people on the Sabana de Bogotá is consistent with deriving wholly from the expansion of Southern Native American (SNA) lineages into South America with no evidence of admixture from other genetic lineages, for example, Anzick-1-related ancestry that spread into Central and South America via subsequent migration events^49^. Likewise, we find no evidence for California Channel Islands-related ancestry (**Table S11**; **Figure S6**). Colombia_SB_HG shares significantly less genetic drift with ancient people from North America and Mesoamerica than with those from Central and South America yet are symmetrically related to ancient groups across Central and South America without specific affinity to any sampled population or region (**Table S9** and **S12**). This pattern is consistent with descent from a deeply diverging ancestral lineage that contributed little to later groups. We confirm the basal positioning and distinctiveness of Colombia_SB_HG with ADMIXTOOLS2, systematically exploring possible admixture graph topologies and evaluating how well each graph fits observed *f*-statistics (**Supplementary Information 6**; Method Details). Colombia_SB_HG represents a distinct lineage that branches from the primary SNA ancestry underlying most South American populations, which dispersed rapidly into and throughout the continent^50^, underscoring both their early divergence from and broader connection to the initial peopling of South America.

### Asynchronous shifts in subsistence and ancestry

As local hunter-gatherer-associated ancestry persisted on the Sabana de Bogotá from the early-middle through initial late Holocene with no evidence for exogenous gene flow during this interval, it is particularly striking that two individuals from Aguazuque – I23785 directly dated to 983-835 calBCE (2770±20 BP, UCIAMS-288524) and I23783 from a context dated to 2725±235 BP (GrN-14479) ^20^, both of whom exhibit genetic profiles entirely consistent with deep local ancestry (**Table S2** and **S13**) – have dietary isotope signatures reflecting a substantive contribution from C_4_-enriched resources (**Figure 4**; **Table S14**; **Supplementary Information 2**). Given the limited availability of wild C_4_ plants in the high-altitude Andean environment^21,51,52^, this isotopic signal is consistent with a meaningful dietary shift, parsimoniously explained by maize playing a more substantial role in the diet at this time^8,12,20,21^ in contrast to its role as a minor, non-staple component of a diversified plant diet at middle Holocene Checua^18^ (**Supplementary Information 2**). Our data therefore decouples the early appearance^18^ and isotopically detectable incorporation of maize into human diets^8,12,20,21^ from the ancestry turnover that took place between pre-Herrera and Herrera-Muisca Periods. This pattern implies that early agricultural practices were adopted or developed within populations showing genetic continuity, rather than introduced through population replacement, and is consistent with the circulation of crops, knowledge, or practices through broader interregional interaction spheres, rather than demic movement^53^.

**Figure 4.**
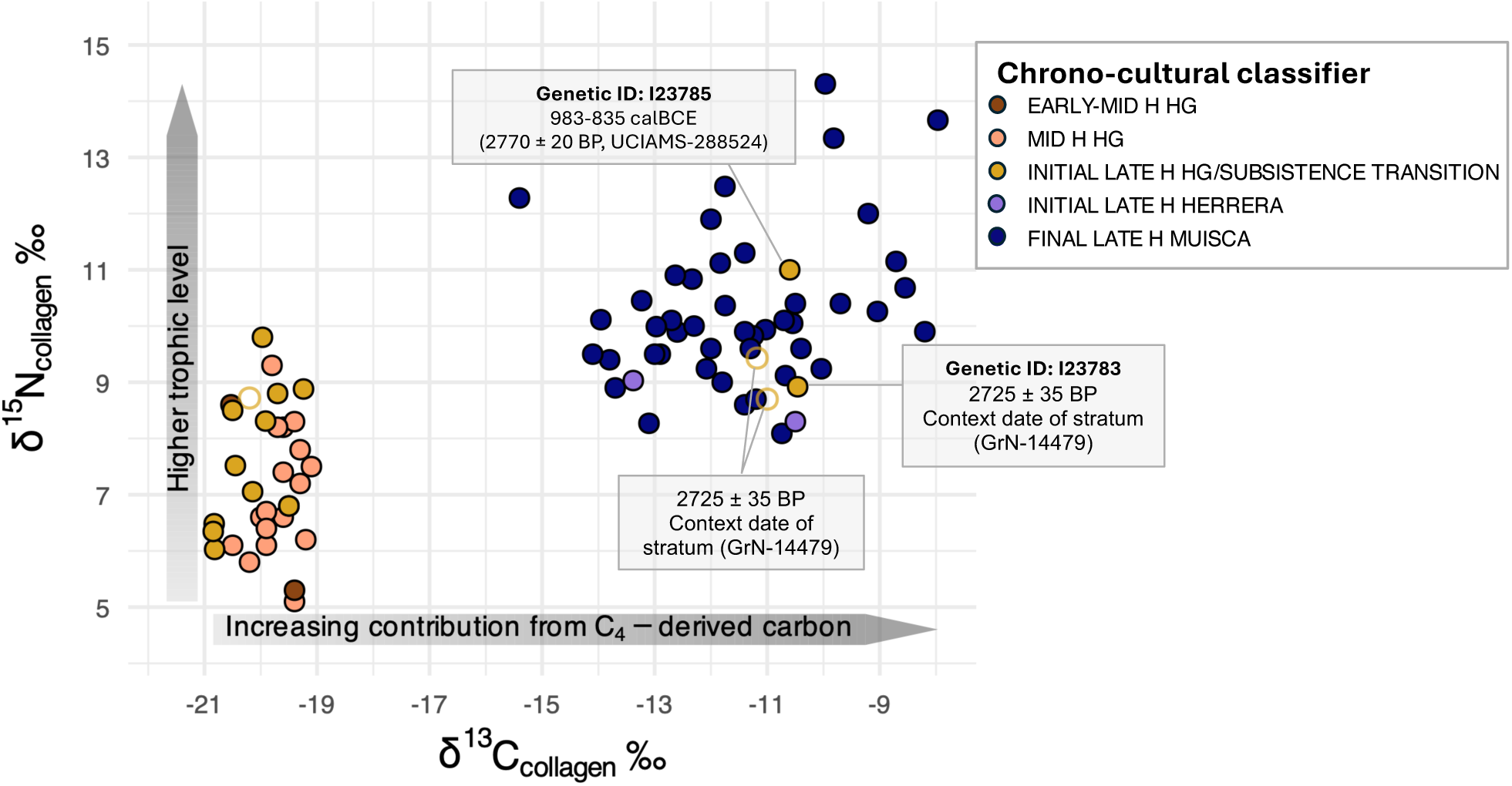
Isotopic measurements on collagen extracted from human bone samples. Scatterplot of δ¹³C_collagen_ versus δ¹⁵N_collagen_ for all individuals with isotopic and ancient DNA data in our dataset (**Table S14**) together with previously published isotopic information (**Supplementary Information 2**). Points are colored by chrono-cultural classifier (H for Holocene): EARLY–MID H HG, MID H HG, INITIAL LATE H HG/SUBSISTENCE TRANSITION, INITIAL LATE H HERRERA, and FINAL LATE H MUISCA. Filled symbols indicate individuals with both isotopic and genomic data, whereas unfilled symbols indicate individuals with isotopic data only (isotopic data from ref. ^20^). The horizontal grey arrow highlights increasing δ¹³C values toward the right, interpreted as increasing contribution from C₄- enriched resources (in this context, maize), and the vertical grey arrow highlights increasing δ¹⁵N values upward, interpreted as higher trophic level and greater reliance on higher-trophic protein sources. Four individuals from initial late Holocene Aguazuque, two with genomic data and one with a direct ^14^C date from bone, who have a diet incorporating C_4_- enriched resources, are labeled.

While hunter-gatherer-associated ancestry persisted for more than 5000 years, by at least 2000 BP, the population living on the Altiplano Cundiboyacense was genetically distinct from the people who lived in the region for many prior millennia. We refer collectively to the ancient individuals postdating 2000 BP as “Herrera-Muisca Period groups,” a term that reflects the absence of temporal sub-structure across these contexts (**Table S7**) but allows us to differentiate them from pre-Herrera people with different ancestry. Individuals from Herrera-Muisca Period contexts form a distinct cluster in PCA and are differentiated from pre-Herrera Period individuals in *qpWave*, indicating a clear ancestry shift and suggesting that the local gene pool was largely, if not entirely, replaced. Consistent with this interpretation, statistics of the form *f_4_*(Yoruba, Colombia_SB_HG; *Colombia_HerreraMuisca, NativeAmerican_Test*) show no significant excess allele sharing between pre-Herrera and Herrera-Muisca Period groups relative to other ancient Native Americans, indicating that the ancestry shift is best explained by a major population replacement rather than admixture (**Table S12**).

The absence of genetic evidence for admixture could reflect genuine reproductive isolation or may result from the absence of genomic data dating 2800-2000 BP – the very period in which interaction and mixing between people with local pre-Herrera hunter-gatherer-associated ancestry and Herrera-Muisca Period-associated ancestry could have occurred. If the latter is true, an admixture signal may have been subsequently diluted or “swamped” by continued gene flow from Herrera-Muisca Period ancestors. Alternatively, if the two groups did remain genetically distinct, their interaction may have been primarily cultural (economic or technological).

Mitochondrial DNA (mtDNA) evidence also supports a demographic shift between pre-Herrera and Herrera-Muisca contexts (**Figure S7**; **Table 15**). While subclades of haplogroup A are frequent in both contexts, the derived A2ac1 lineage which dominates among Herrera-Muisca Period people is absent prior to the Herrera Period. This pattern suggests the introduction or expansion of new maternal lineages. Relative to pre-Herrera contexts, Herrera-Muisca contexts are defined by a pronounced increase in B haplogroups, a sharp decline in C haplogroups, and the appearance of D haplogroups. Haplogroup B is prevalent among northern Andean and Chibchan-speaking groups^54^, while the subgroup B2d that appears most frequently in Herrera-Muisca contexts has a more restricted distribution limited to lower Central America and Chibchan-influenced regions of northern South America^5,49,55–57^. As this lineage is notably absent among pre-Herrera people of the Sabana de Bogotá, its presence in the region may act as a “tracer dye” for the spread of Chibchan-related ancestry onto the Altiplano Cundiboyacense. In contrast, haplogroup D is largely absent in Chibchan-related populations and instead found at higher frequencies in Amazonian and eastern Andean groups, suggesting complexity in the ancestry of Herrera-Muisca Period groups. Consistent with mtDNA, statistics of the form *f_4_*(Yoruba, *Test*; Colombia_SB_HG, *Colombia_HerreraMuisca*) reveal that, relative to pre-Herrera Colombian hunter-gatherers and early food producers, Herrera-Muisca Period people share significantly more alleles with Chibchan-related populations (|Z| up to 17.2) as well as with some Amazonian groups or those with Amazonian-related affinities, primarily those with so-called “Amazonian North” affinity who live on the eastern slopes of the northern Andes in present-day Colombia and Ecuador and speak Andean languages but are traditionally associated with the Amazonian cultural domain^58^ (**Figures S8A** and **S9A**; **Tables S16** and **S17**). In what follows, we focus specifically on reconstructing the ancestry of the people of the Herrera and Muisca Periods who lived in the initial late and final late Holocene.

### The ancestry of Herrera-Muisca Period groups formed ∼4000 years BP

Previous work showed that by 2000 BP, local ancestry on the Altiplano Cundiboyacense was replaced by ancestry connected to Central America associated with the Herrera ceramic complex, who persisted through the Muisca Period^36^. Post-2000 BP people of the Altiplano Cundiboyacense are assessed as being genetically intermediate between ancient Panamanians and other Central and South Americans, but a source for the latter ancestry was previously unidentified^36^.

We find evidence of a marked demographic expansion during the Herrera and Muisca Periods. With *hapROH*, we estimate community sizes ranging from approximately 310–438 individuals during the Herrera Period to 513–601 during the Muisca Period (95% confidence intervals derived from likelihood profiles; Method Details); these numbers do not reflect the size of the population of the whole region, but instead the gene pool of people contributing to any sampled individuals over the last handful of generations, and so are estimates of more localized populations. Our estimates point to continuous population growth on the Altiplano Cundiboyacense, aligning with archaeological indicators of demographic increase, the expansion of food production, technological specialization, and the emergence of larger and more socially stratified communities. The genetic signal of expansion thus complements archaeological evidence for sustained population growth and a transition toward more sedentary and complex societies throughout the Herrera and Muisca Periods^33,44,59^. Three Muisca Period individuals out of 150 – each from a different site (Marín, Portalegre, and El Venado) – are the offspring of close-kin unions, exhibiting 385–406cM of ROH>4cM and 226–255cM of ROH>20cM (**Figure S10**; Method Details). While these values imply consanguineous unions, the absence of a broader shift toward elevated ROH suggests that unions of close kin were rare despite the acceptance of cross-cousin marriage within the Muisca kinship system^44,60,61^. We identify several ‘families’ comprising closely related individuals (up to 3rd-degree relatives), including one spread across the southern Muisca sites of Tibanica, Portalegre, and Candelaria La Nueva (**Supplementary Information 5**).

While Herrera-Muisca Period people form a relatively homogenous genetic group relative to other ancient and modern Americans, we detect sub-structure between Herrera and Muisca Period groups that aligns with archaeologically-attested cultural differences between the southern Muiscas (present-day Cundinamarca) and northern Muiscas (present-day Boyacá), the latter associated with greater cosmopolitanism and more connectivity to Amazonia (**Supplementary Information 3**). This finding is consistent with cultural indicators, like burial practices, which have been found to differ between northern and southern Muisca groups^62^. To explore fine-scale genetic differences, we used *f*₄-statistics to assess asymmetries in allele sharing between the three regional groups of Colombia_Cundinamarca, Colombia_Boyacá1_LaSalina, and Colombia_Boyacá2 with ancient and present-day Native American populations. While all regional groups show broadly symmetric allele sharing with ancient Native Americans, we find distinct geographic patterns in allele-sharing with present-day populations. Colombia_Cundinamarca shares relatively more alleles with “Amazonian North” groups, consistent with proximity to the northwestern Amazon and Andean foothills. In contrast, Colombia_Boyacá1_LaSalina and Colombia_Boyacá2 display stronger Chibchan-related affinities. Additionally, Colombia_Boyacá2 shows a subtle signal of excess Piapoco-related ancestry especially relative to Colombia_Boyacá1_LaSalina, suggesting localized variation in gene flow that may reflect connectivity to the *Llanos Orientales*, plains situated to the east of the eastern Colombian Andes (**Figures S8B** and **S9B**; **Tables S3** and **S4**). Overall, differences among Herrera-Muisca Period groups were close to the limits of *f*-statistic resolution, suggesting they were shaped less by distinct migration pulses than by varying proportions of shared ancestries and geographically structured patterns of interaction across the Altiplano Cundiboyacense.

Herrera-Muisca Period groups are characterized by a significant excess of shared genetic drift with Central American Chibchan-related populations (proxied by ancient Panamanians^49^ and present-day Cabécar) relative to any other ancient or contemporary American group (**Tables S18** and **S19**). However, the two Central American Chibchan-related proxies share significantly more drift with each other relative to Herrera-Muisca groups (|Z|>10.2; **Table S20**) consistent with them descending from a common ancestral population after separating from the lineage leading to Herrera-Muisca Period people, and/or admixture in the Herrera-Muisca ancestors not present in Chibchan-speaking groups. The statistic *f_4_*(Yoruba, *NativeAmerican_Test*; *ChibchanProxy*, *Colombia_HerreraMuisca*) shows that a range of American groups share significantly more genetic drift with Herrera-Muisca Period people than with ancient or present-day Central American Chibchan-speakers (**Tables S21**-**S24**). Populations exhibiting this pattern include representatives of deep South American lineages—for example, from Brazil, the Andes, and the Southern Cone—consistent with drift that accumulated after the divergence from Central American Chibchan-related groups. Additional signals of allele sharing between ancient Colombians and Early/Late Intermediate Period Peruvians as well as Caribbean-related groups could reflect deep shared ancestry and/or later gene flow. Affinities to Mesoamerican groups may point to a distinct ancestry layer separate from the North American–derived component present in Central American Chibchan populations. Present-day groups with “Amazonian North” ancestry^58^ also share more alleles with ancient Colombians relative to Chibchan-related groups. Collectively, these results invalidate a simple clade relationship between Central American Chibchan speakers and Herrera-Muisca Period people.

A possible explanation is that Herrera-Muisca Period people on the Altiplano Cundiboyacense carried additional ancestry in addition to that related to Central American Chibchan speakers. Consistent with this interpretation, *qpAdm* rejects one-source models in which any Herrera-Muisca regional group derives all ancestry from a single source, including Chibchan-related proxies (p<<0.05), (**Supplementary Information 7**). Earlier genomic analyses^36^ did not identify a suitable ancient proxy for a population that admixed with Chibchan-related groups to form the post-2000 BP population of the Altiplano Cundiboyacense, but we use the *qpAdm* framework to fit a model for the ancestry of the Herrera-Muisca Period groups as a mixture of Chibchan-related and northwest Amazonian/Andean-related ancestry, the former proxied in our models by ancient Panamanians and the latter well-proxied by the Kichwa Orellana group living on the eastern slopes of the northern Andes who carry the “Amazonian North” ancestry typical of this region, which split an estimated 4,000 years ago from the “Amazonian Core” ancestry found in the central parts of the Amazon Basin^58^. We detect subtle regional ancestry variation as suggested by *qpWave* and *f_4_-*statistics but find consistent support for primary contributions of both Chibchan-related and Amazonian/Andean-related ancestry to the Herrera-Muisca gene pool. All regional groups can be modeled as two-way mixtures, with Colombia_Cundinamarca fit as having 53.9%±1.4% Chibchan-related and 46.1%±1.4% Amazonian North-related ancestry (p=0.95), Colombia_Boyacá1_Salina fit as having 62.2%±2.0% Chibchan-related and 37.8%±2.0% Amazonian North-related ancestry (p=0.73), and Colombia_Boyacá2 fit as having 59.2%±1.7% Chibchan-related and 40.8% ± 1.7% Amazonian North-related ancestry (p=0.13, with the model fitting only when Piapoco is removed from the reference set, **Supplementary Information 7**). These proportions are consistent with the stronger Amazonian North-related signal in geographically proximate Cundinamarca and stronger Chibchan-related signal in Boyacá observed with *f-*statistics. With the software *DATES*, we estimated the date of admixture between the two ancestries that formed the gene pool of Herrera-Muisca people to have occurred on average 137 ± 15 generations (or 3964 ± 446 years) before those individuals lived (Z=8.9) (**Figure S11**; estimates using varied sources were qualitatively similar, see **Table S25**). This ancient admixture date implies that the genetic foundation of Herrera and Muisca Period populations was established prior to their archaeological prominence in the eastern Colombian highlands.

## DISCUSSION

By analyzing genome-wide data from 209 individuals from the Altiplano Cundiboyacense spanning 7000 years from the early-middle to final late Holocene and integrating them with previously published genomic data from the region^36^, we reconstruct population history at a temporal and spatial resolution not previously attainable. This expanded dataset allows us to delineate fine-scale structure between groups with hunter-gatherer-associated ancestry, demonstrate the persistence of this ancestry across a subsistence shift most parsimoniously related to maize cultivation, constrain the timing and establish the independence of subsistence and ancestry transformations, and clarify the demographic origins of the people who ultimately gave rise to Muisca communities.

We show that the ancestry of pre-Herrera Period people from the Sabana de Bogotá derives wholly from the primary expansion of Southern Native American (SNA) ancestry into South America and forms a distinct lineage that is nevertheless deeply connected to the broader peopling of South America. There was population continuity over five millennia with no evidence of gene flow from genetically distinct populations between the early-middle and early initial final Holocene, despite shifts in subsistence during this time. Sub-structure reflects the accumulation of genetic drift, likely influenced by environmental shifts such as the rising temperatures and decreased humidity of the middle Holocene that resulted in habitat fragmentation and reduced the availability of key resources including both animals (deer and guinea pigs) and plants^4,16,63^. Changes in resource exploitation strategies, mobility, and social networks, potentially increasing isolation, may be one reason why some of the earliest hunter-gatherer lineages are not reflected in later groups. Despite very small community sizes, close-kin mating was avoided.

There is abundant data supporting the early introduction of maize to Andean South America and particularly to the Sabana de Bogotá. In regions of Colombia like the Middle Cauca River Basin (an inter-Andean region located in the western central part of the Colombian Andes), analysis of pollen, starch grains, and phytoliths from tools indicates processing of maize seed likely for direct consumption as early as ∼7000 BP^30,64,65^, while grains of maize have been identified in the intra-Andean basin of Duitama between 3680-2610 BP^66^ and in dental calculus of individuals dating ∼4900 BP from the site of Checua^18^. An important contribution of our study is demonstrating that the incorporation of maize as a meaningful component in diet occurred independently of the major change in ancestry that occurred sometime between 6000 and 2000 BP^36^. Here, we show that two individuals (I23783 and I23785) from the site of Aguazuque, the latter directly dated to 983-835 calBCE, are genetically indistinguishable from the long-standing local hunter-gatherer lineage yet are consuming enough C₄-derived resources to be reflected as an isotopic shift. These results provide genomic support for the incorporation of C₄ plants into subsistence systems by resident populations well before the ancestry turnover associated with the arrival of the ancestors of Herrera-Muisca Period people. The early populations of the Sabana de Bogotá were evidently active participants in subsistence diversification, suggesting well-established economic networks.

Our results also confirm the previous finding of a major genetic turnover occurring on the Altiplano Cundiboyacense^36^ while refining the date of this turnover as having occurred between 2800-2000 BP, with no measurable genetic contribution from autochthonous groups detected among later populations of the Herrera and Muisca Periods. While our data suggest a population replacement took place, future research should address the processes and factors that led to the near-total absence of hunter-gatherer related ancestry persisting into the Herrera and Muisca Periods. While previous work documents a Chibchan-related genetic shift^36^, we find that the ancestors of Herrera-Muisca Period people did not carry exclusively Chibchan-related ancestry but also an additional genetic component absent from the gene pool of Central American Chibchan-speaking groups. Based on the currently available data – which include very limited ancient genomic data from individuals with Amazonian ancestry – this ancestry is best characterized as having Amazonian–Andean affinities and is most closely approximated by the “Amazonian North” genetic profile observed today in the Andean foothills of Ecuador and Colombia, regions that maintain cultural and demographic connections with the Amazon basin^58^. We estimated that the admixture event forming this gene pool took place ∼4000 BP – well before the appearance of Herrera ancestors on the Altiplano Cundiboyacense – implying that the genetic foundations of Herrera and Muisca populations formed outside this region prior to their expansion into the Altiplano.

A hypothesized route of the migration of Chibchan-speaking populations with origins in Central America onto the Altiplano Cundiboyacense is through the Magdalena River Valley; however, this is geographically distant from present-day regions inhabited by genetically Amazonian North groups, leaving open the question of where the admixture that formed the gene pool of Herrera-Muisca Period ancestors may have taken place. Possibly, there was a broader distribution of people with Amazonian North-like ancestry in the past. Alternatively, the absence of a more precise ancient proxy inhibits our ability to give this ancestry an accurate geographic designation. Importantly, the estimated admixture date is coincident with the linguistic evidence for the diversification of Chibchan languages and spread of Chibchan speakers throughout the Isthmo-Colombian region^35,67^.

Consistent with archaeological evidence of sustained population growth and a greater reliance on agriculture, genomic evidence indicates population expansion during the Herrera and Muisca Periods. This is supported by recent paleoenvironmental reconstructions which suggest a significant human impact on the regional landscapes during the Herrera and Muisca Periods^68–70^ and archaeological work that shows significant and sustained population growth from the Herrera to the Late Muisca Period^33,43,45,71^. Herrera-Muisca Period people represent a largely homogeneous yet internally structured genetic cluster, showing minor differentiation between northern and southern subgroups (corresponding to the present-day departments of Boyacá and Cundinamarca, respectively) that parallels archaeological distinctions in material culture, burial practices, and connection networks^72–75^. Subtle allele-sharing patterns reveal greater “Amazonian North” ^58^ affinity in the southern Cundinamarca region and stronger Chibchan-related signals in Boyacá, suggesting culturally and geographically mediated differences in external contact and gene flow throughout the Altiplano Cundiboyacense.

Several centuries ago, the eastern Colombian Andes were home to multiple large Muisca chiefdoms. They practiced multi-crop agriculture, specialized in craft production, and sustained extensive trade networks that linked the northern Andean highlands to the Caribbean and Amazonian lowlands and beyond. The Muisca were renowned for their sophisticated gold metallurgy and ritual use of precious metals, which became central to their economy and cosmology. As Muisca communities flourished, accounts of their wealth circulated widely, ultimately attracting Spanish invaders. In 1537, Spanish forces encountered the Muisca and incorporated their territories into the expanding colonial domain. Although this marked the end of indigenous political autonomy on the Sabana de Bogotá, descendant communities continue to inhabit the region today. Ancient DNA provides a means to illuminate the history of the people who have shaped and reshaped the region’s genetic and cultural landscape.

## ACKNOWLEDGEMENTS

We thank the Instituto Colombiano de Antropología e Historia (ICANH); Museo Nacional de Colombia; Instituto de Ciencias Naturales, Universidad Nacional de Colombia; Universidad de los Andes; Universidad Pedagógica y Tecnológica de Colombia (UPTC); and El Departamento Estudios Sociales, INGETEC, for permissions and support for this project. We thank Nicole Adamski, Kristin Stewardson, and Fatma Zalzala for ancient DNA laboratory work and Brendan Culleton and Laurie Eccles for radiocarbon dates and isotope analysis and for discussions about those data. Sonia Archila, Juan Pablo Ospina, Saúl Alberto Torres Orjuela, and Martha Beatriz Mejía Cano developed the archaeological research program in the Checua River valley, excavating several sites and conducting archaeobotanical and archaeothanatological studies, among other types of analyses. K.S. and M.D. received a pilot grant from the National Geographic Society that supported this work. D.R. acknowledges support from the National Institutes of Health (HG012287); the John Templeton Foundation (grant 61220); from J.-F. Clin; from the Allen Discovery Center, a Paul G. Allen Frontiers Group advised program of the Paul G. Allen Family Foundation; and from the Howard Hughes Medical Institute. The author-accepted version of this article, that is, the version not reflecting proofreading and editing and formatting changes following the article’s acceptance, is subject to the Howard Hughes Medical Institute (HHMI) Open Access to Publications policy, as HHMI lab heads have previously granted a nonexclusive CC BY 4.0 license to the public and a sublicensable license to HHMI in their research articles. Pursuant to those licenses, the author-accepted manuscript can be made freely available under a CC BY 4.0 license immediately upon publication. M.D. dedicates this paper to the memory of Franz Xaver Faust (1953-2025), a beloved friend and teacher.

## RESOURCE AVAILABILITY

### Lead contact

Further information and requests for resources and reagents should be directed to and will be fulfilled by the lead contact, Kendra Sirak.

### Materials availability

Open science principles require making all data used to support the conclusions of a study maximally available, and we support these principles here by making fully publicly available not only the digital copies of molecules (the uploaded sequences) but also the molecular copies (the ancient DNA libraries themselves, which constitute molecular data storage). Those researchers who wish to carry out deeper sequencing of libraries published in this study should make a request to the corresponding authors K.S. or D.R. We commit to granting reasonable requests as long as the libraries remain preserved in our laboratories, with no requirement that we be included as collaborators or co-authors on any resulting publications.

### Data and code availability

Newly reported ancient DNA data of this study are fully publicly available and have been deposited in the European Nucleotide Archive (ENA project accession number PRJEB110479). This paper does not report original code.

## AUTHOR CONTRIBUTIONS

Conceptualization, K.S., M.D., L.F.-S. and D.R.; writing – original draft, K.S.; writing – review & editing, K.S., M.D., and D.R. with the input of all other co-authors; resources – archaeological materials assembly and descriptions, S.R., A.T., P.A., A.M.B., J.R.S., G.P., E.R., C.L., J.P.O., and S.A.; project administration, K.S. and M.D.; methodology – sampling, K.S., M.D., and L.F.-S.; methodology – laboratory work, K.C., E.C., T.F., L.I., J.K., A.M.L., L.Q., J.N.W., and N.R.; methodology – data processing and curation, A.K., M.M., S.M., M.N., G.S., and D.R.; methodology – radiocarbon dates and isotope data, A.C., C.S.H., and K.P.; formal analysis, K.S. and A.B.; visualization, K.S.; supervision, D.R.; funding acquisition, K.S., M.D., and D.R.

## DECLARATION OF INTERESTS

The authors declare no competing interests.

## INCLUSION AND DIVERSITY

We support inclusive, diverse, and equitable conduct of research.

## SUPPLEMENTAL INFORMATION

**Supplementary Information 1**. Archaeological context.

**Supplementary Information 2**. Details of isotope analysis and interpretation of isotope data.

**Supplementary Information 3**. Details of *qpWave* analysis.

**Supplementary Information 4**. Individuals not incorporated into broader analysis groups.

**Supplementary Information 5**. Description of ‘families’ identified with ancient DNA

**Supplementary Information 6**. Description of *qpGraph* analysis.

**Supplementary Information 7**. Description of *qpAdm* analysis.

## STAR★METHODS

### KEY RESOURCES TABLE

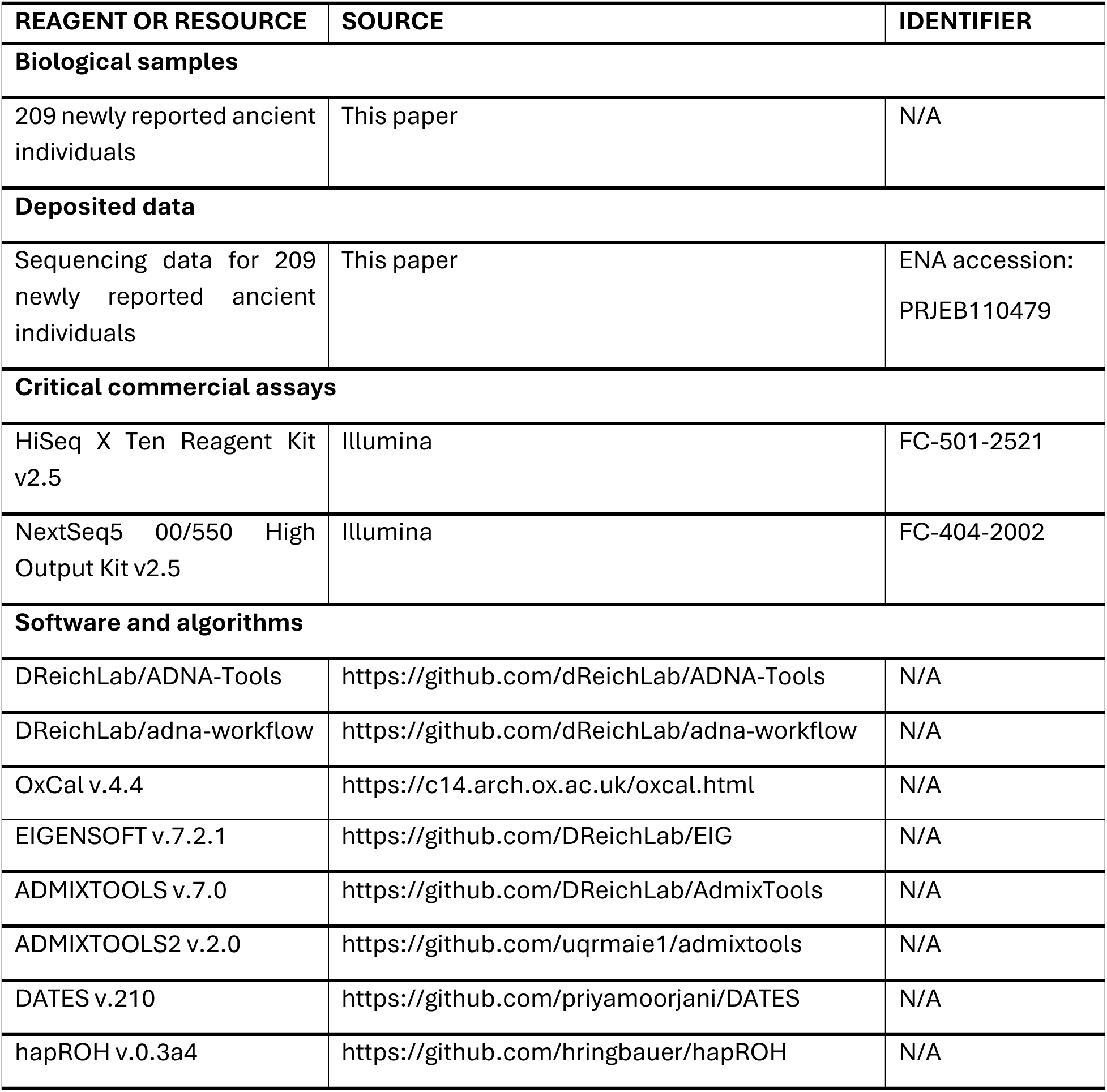
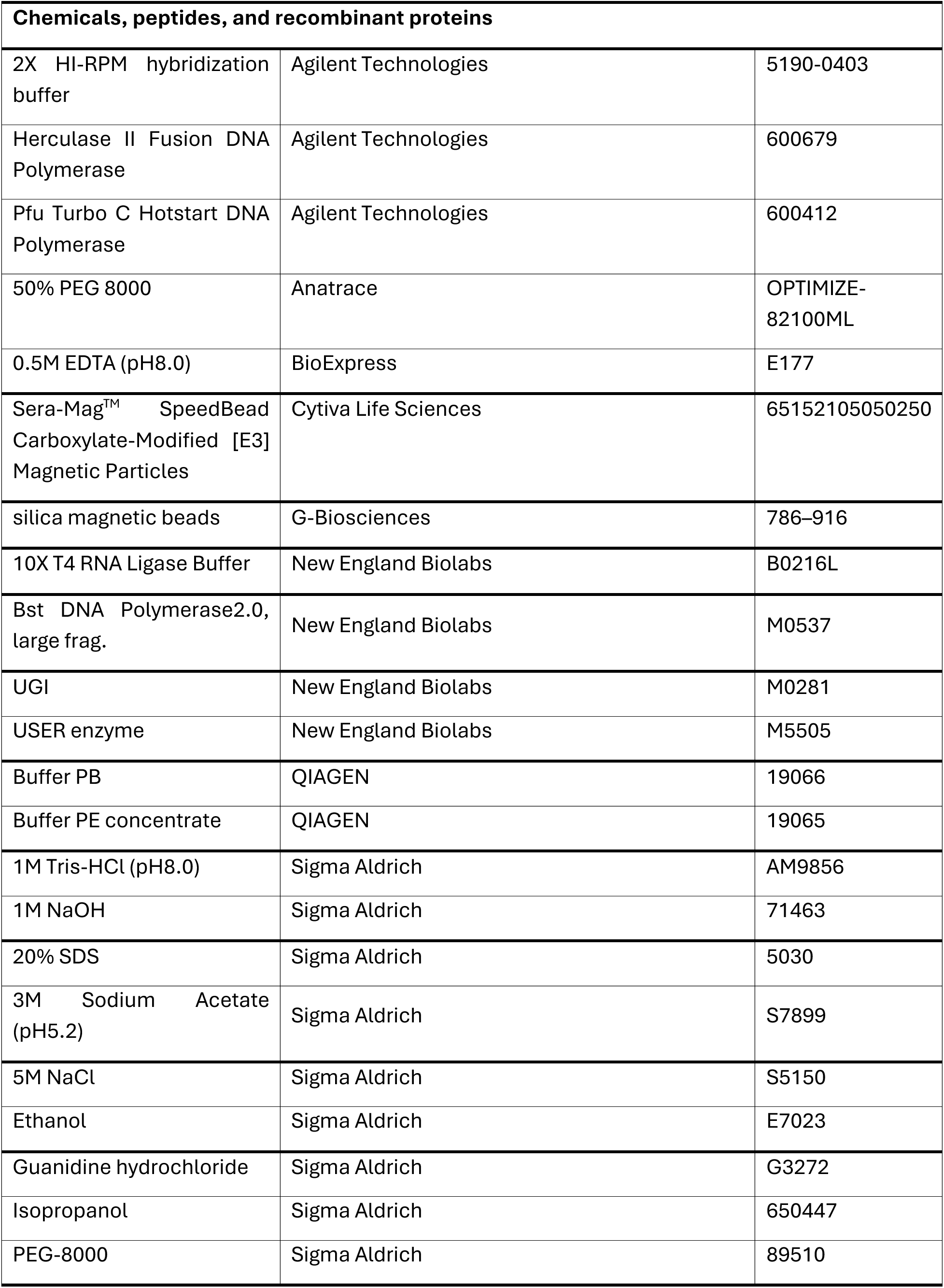
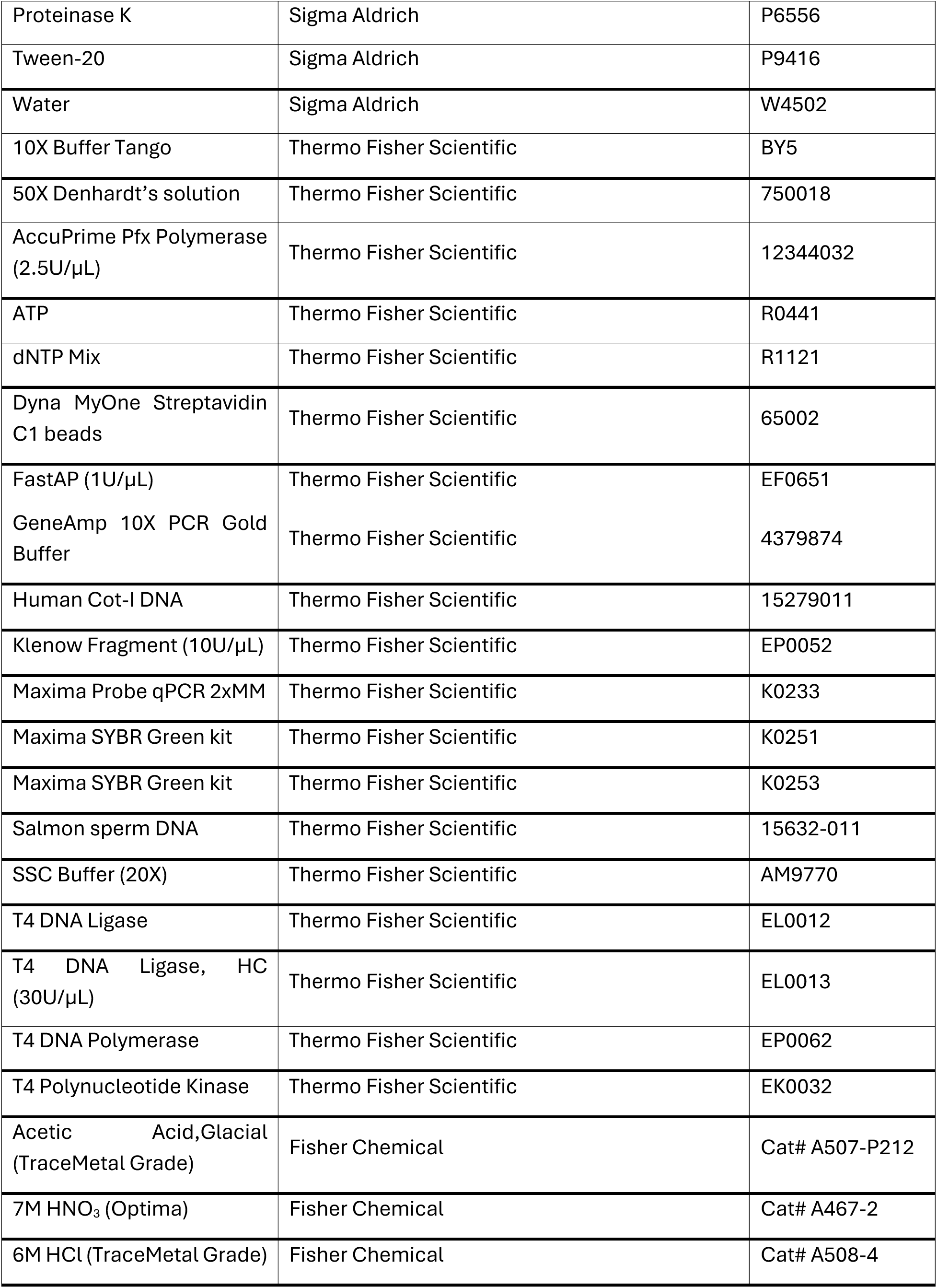
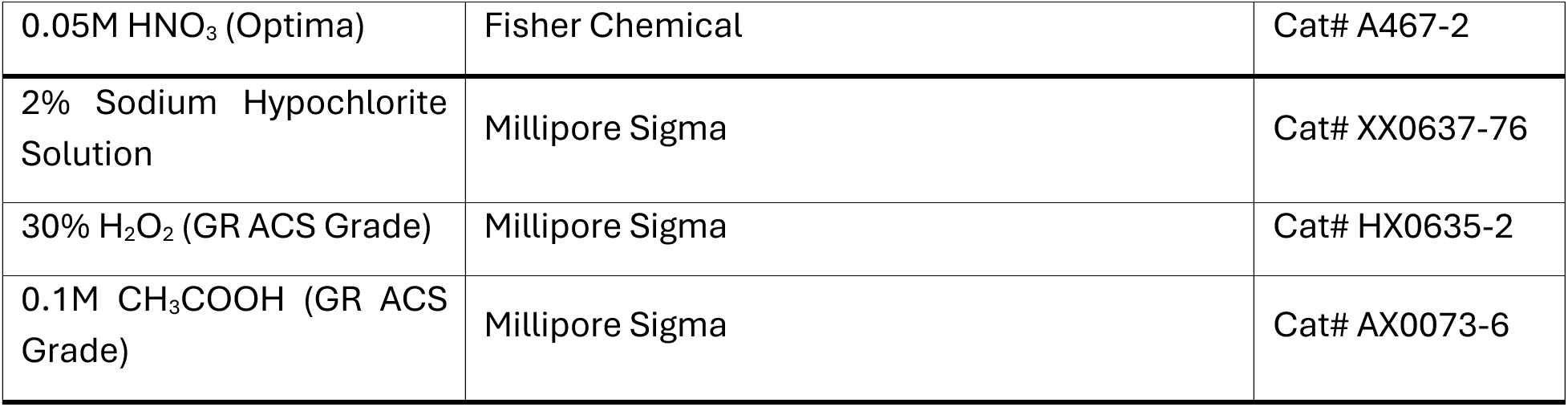

| REAGENT OR RESOURCE | SOURCE | IDENTIFIER |
| --- | --- | --- |
| **Biological samples** |  |  |
| 209 newly reported ancient individuals | This paper | N/A |
| **Deposited data** |  |  |
| Sequencing data for 209 newly reported ancient individuals | This paper | ENA accession: PRJEB110479 |
| **Critical commercial assays** |  |  |
| HiSeq X Ten Reagent Kit v2.5 | Illumina | FC-501-2521 |
| NextSeq5 00/550 High Output Kit v2.5 | Illumina | FC-404-2002 |
| **Software and algorithms** |  |  |
| DReichLab/ADNA-Tools | https://github.com/dReichLab/ADNA-Tools | N/A |
| DReichLab/adna-workflow | https://github.com/dReichLab/adna-workflow | N/A |
| OxCal v.4.4 | https://c14.arch.ox.ac.uk/oxcal.html | N/A |
| EIGENSOFT v.7.2.1 | https://github.com/DReichLab/EIG | N/A |
| ADMIXTOOLS v.7.0 | https://github.com/DReichLab/AdmixTools | N/A |
| ADMIXTOOLS2 v.2.0 | https://github.com/uqrmaie1/admixtools | N/A |
| DATES v.210 | https://github.com/priyamoorejani/DATES | N/A |
| hapROH v.0.3a4 | https://github.com/hringbauer/hapROH | N/A |

| **Chemicals, peptides, and recombinant proteins** |  |  |
| --- | --- | --- |
| 2X HI-RPM hybridization buffer | Agilent Technologies | 5190-0403 |
| Herculase II Fusion DNA Polymerase | Agilent Technologies | 600679 |
| Pfu Turbo C Hotstart DNA Polymerase | Agilent Technologies | 600412 |
| 50% PEG 8000 | Anatrace | OPTIMIZE-82100ML |
| 0.5M EDTA (pH8.0) | BioExpress | E177 |
| Sera-Mag™ SpeedBead Carboxylate-Modified [E3] Magnetic Particles | Cytiva Life Sciences | 65152105050250 |
| silica magnetic beads | G-Biosciences | 786–916 |
| 10X T4 RNA Ligase Buffer | New England Biolabs | B0216L |
| Bst DNA Polymerase2.0, large frag. | New England Biolabs | M0537 |
| UGI | New England Biolabs | M0281 |
| USER enzyme | New England Biolabs | M5505 |
| Buffer PB | QIAGEN | 19066 |
| Buffer PE concentrate | QIAGEN | 19065 |
| 1M Tris-HCl (pH8.0) | Sigma Aldrich | AM9856 |
| 1M NaOH | Sigma Aldrich | 71463 |
| 20% SDS | Sigma Aldrich | 5030 |
| 3M Sodium Acetate (pH5.2) | Sigma Aldrich | S7899 |
| 5M NaCl | Sigma Aldrich | S5150 |
| Ethanol | Sigma Aldrich | E7023 |
| Guanidine hydrochloride | Sigma Aldrich | G3272 |
| Isopropanol | Sigma Aldrich | 650447 |
| PEG-8000 | Sigma Aldrich | 89510 |
| Proteinase K | Sigma Aldrich | P6556 |
| Tween-20 | Sigma Aldrich | P9416 |
| Water | Sigma Aldrich | W4502 |
| 10X Buffer Tango | Thermo Fisher Scientific | BY5 |
| 50X Denhardt's solution | Thermo Fisher Scientific | 750018 |
| AccuPrime Pfx Polymerase (2.5U/μL) | Thermo Fisher Scientific | 12344032 |
| ATP | Thermo Fisher Scientific | R0441 |
| dNTP Mix | Thermo Fisher Scientific | R1121 |
| Dyna MyOne Streptavidin C1 beads | Thermo Fisher Scientific | 65002 |
| FastAP (1U/μL) | Thermo Fisher Scientific | EF0651 |
| GeneAmp 10X PCR Gold Buffer | Thermo Fisher Scientific | 4379874 |
| Human Cot-I DNA | Thermo Fisher Scientific | 15279011 |
| Klenow Fragment (10U/μL) | Thermo Fisher Scientific | EP0052 |
| Maxima Probe qPCR 2xMM | Thermo Fisher Scientific | K0233 |
| Maxima SYBR Green kit | Thermo Fisher Scientific | K0251 |
| Maxima SYBR Green kit | Thermo Fisher Scientific | K0253 |
| Salmon sperm DNA | Thermo Fisher Scientific | 15632-011 |
| SSC Buffer (20X) | Thermo Fisher Scientific | AM9770 |
| T4 DNA Ligase | Thermo Fisher Scientific | EL0012 |
| T4 DNA Ligase, HC (30U/μL) | Thermo Fisher Scientific | EL0013 |
| T4 DNA Polymerase | Thermo Fisher Scientific | EP0062 |
| T4 Polynucleotide Kinase | Thermo Fisher Scientific | EK0032 |
| Acetic Acid, Glacial (TraceMetal Grade) | Fisher Chemical | Cat# A507-P212 |
| 7M HNO<sub>3</sub> (Optima) | Fisher Chemical | Cat# A467-2 |
| 6M HCl (TraceMetal Grade) | Fisher Chemical | Cat# A508-4 |
| 0.05M HNO<sub>3</sub> (Optima) | Fisher Chemical | Cat# A467-2 |
| 2% Sodium Hypochlorite Solution | Millipore Sigma | Cat# XX0637-76 |
| 30% H<sub>2</sub>O<sub>2</sub> (GR ACS Grade) | Millipore Sigma | Cat# HX0635-2 |
| 0.1M CH<sub>3</sub>COOH (GR ACS Grade) | Millipore Sigma | Cat# AX0073-6 |

### EXPERIMENTAL MODEL AND STUDY PARTICIPANT DETAILS

#### Ancient Individuals

An extensive description of the archaeological context of the ancient individuals analyzed in this study is provided in **Supplementary Information 1**.

### METHOD DETAILS

#### Sampling of ancient individuals

All skeletal material screened for ancient DNA was sampled, exported, and analyzed with full permission obtained from Instituto Colombiano de Antropología e Historia (ICANH; this is the agency under the Ministry of Culture in Colombia which oversees the protection and management of archaeological heritage). This project proceeded under archaeological intervention license number 8270, issued to M.D. on August 20, 2019. Samples were taken from skeletal materials curated at different institutions: Museo Nacional de Colombia; Instituto de Ciencias Naturales, Universidad Nacional de Colombia; Universidad de los Andes; Universidad Pedagógica y Tecnológica de Colombia (UPTC); and El Departamento Estudios Sociales, INGETEC. All ancient DNA research was conducted in accordance with the ethical guidelines outlined by the international research community^76^. Descriptions of the archaeological sites and contexts from which the skeletal material in this study derives is in **Supplementary Information 1**. The support we received from local academic and community members contributed to this work.

#### Ancient DNA data generation, bioinformatics, and quality control

When sampling human skeletal material for this work, we prioritized skeletal elements recognized to be especially DNA-rich whenever possible^77–80^. In cases where skulls were complete, we generated powder from the cochlea by approaching it from the cranial base^81^. In other cases, we generated powder from bone and tooth samples^79^ in dedicated ancient DNA clean room facilities at Harvard Medical School (Boston), the University of Vienna (Vienna), or the University of California Santa Cruz (UCSC Paleogenomics) (see **Table S26** for location where powder was prepared). DNA was extracted from powder using a method optimized for retaining small DNA fragments^82^. From the DNA extracts, we prepared truncated dual-barcoded double-stranded libraries^83^ or full-length dual-indexed single-stranded libraries^84^. For all libraries (ds.half and ss.USER), we used *Escherichia coli* UDG (in the enzyme mix “USER” from NEB), which removes Uracils (except when a deaminated Cytosine wasmethylated) and cuts DNA strands at abasic sites, but inefficiently cuts terminal dephosphorylated Uracils and thereby leaving C-to-T ‘damage’ characteristic of aDNA. For all double-stranded libraries we replaced MinElute columns for reaction clean-ups with magnetic silica beads and Qiagen buffer PB and used SPRI beads instead of MinElute columns for PCR cleanup at the end of library preparation^85^. We added two seven-base-pair (bp) indexing barcodes to the truncated adapters of each double-stranded library (for shotgun sequencing to the library or after enrichment) and thereby completed the adapter sites for sequencing (it is not necessary to add these barcodes to single-stranded libraries, which are already indexed and adapter sites are completed from library preparation). We produced DNA libraries from 240 unique individuals, sometimes multiple libraries per sample.

We enriched the libraries using one of the following strategies: 1) for sequences overlapping mtDNA^86^ and approximately 1.2 million nuclear targets (“1240k”) simultaneously^87–89^; or 2) using the Twist Ancient DNA assay^90^ (Twist Bioscience) for approximately 1.4 million SNPs. The Twist reagent includes all SNPs in the “1240k” capture set and has been shown to generate data that is fully co-analyzable with shotgun sequencing data^90^ and shows no evidence of allelic bias^91^. To generate paired-end reads, enriched libraries were sequenced on the Illumina NextSeq500 instrument for 2×76 cycles or on the Illumina HiSeq X10 or NovaSeq instrument for 2×101 cycles. Indices were read with 2×7 cycles (double-stranded libraries) or 2×8 cycles (single-stranded libraries).

The resulting read pairs were separated using library-specific barcode pairs (for double-stranded libraries) or index pairs (for single-stranded libraries) and merged prior to alignment. We retained reads that showed no more than one mismatch between the forward and reverse base if base quality was at least 20, or three mismatches if base quality was less than 20. A custom toolkit (available at https://github.com/DReichLab/ADNA-Tools) was used for merging and trimming adapters and barcodes. Merged sequences were mapped to the human reference genome version (hg19) and the Reconstructed Sapiens Reference Sequence (RSRS) ^92^ using samse command of the Burrows-Wheeler aligner^93^ (v.0.7.15). Duplicate sequences were removed following alignment using the Picard MarkDuplicates tool (v.2.17.10; http://broadinstitute.github.io/picard/). For variant calling, we used an adaptive pseudohaploid approach. We estimated error rates empirically, assuming that monomorphic sites in 1000 Genomes data^94^ are truly monomorphic, and then stratified these error rate estimates by library type, SNP bases (variant or reference), read position, strand, and mapping and base quality, with the base positions more than 10 bases from the 5’ or 3’ end being considered central and merged. These error rates are determined from the sample BAM, allowing for an unbiased adaptive pulldown procedure. Because at a (C,T) SNP the estimated error *E*(C,T) for C->T may be very different from E(T,C) for T->C, we use a symmetric function *S* and calculate (for example) S=max[*E*(C,T),*E*(T,C)]. We threshold *S* with a parametric value of 0.02 and based with *S* below this value go into a pileup of reliable bases out of which a random base is selected. The error achieved is smaller than the threshold which is an upper bound on the error of each potential base that contributes to the pileup.

From the resulting data, we excluded any individuals covered at <15,000 SNPs on the 1240k capture set and excluded or flagged individuals as potentially contaminated if the C->T substitution rate was less than 3% in UDG-treated libraries or less than 10% in non-UDG-treated libraries, the lower bound for ANGSD^95^ or hapConX^96^ was >0.01, the mtDNA consensus match upper bound estimated using contamMix v1.0-12^97^ was <0.90, or the sex ratio was intermediate between 0.05-0.3 (**Table S1**). Twenty-nine individuals were excluded based on coverage or evidence of contamination and two individuals (I23670 and I23795) were excluded because direct radiocarbon dates on the sampled bones were inconsistent with the chronological context of the sites to which they had been attributed. One individual from first-degree relative pairs was excluded from population genetics analysis when the data were grouped; in all cases, we retained the higher-coverage individual. Library information for all individuals studied as part of this work is in **Table S26**.

#### Dataset assembly

We merged newly-generated data from 209 individuals passing quality control checks into a co-analysis dataset of previously-published data from ancient and present-day individuals^36,47,49,50,98–118^ whose genomes had been shotgun sequenced or enriched for sequences covering a canonical set of ∼2M SNPs^90^, which includes targeted SNPs as well as off-target sites (that is, sites not originally targeted by the enrichment protocol but that are commonly captured because of close physical proximity). We refer to this as the ‘2M’ dataset. We also merged the newly-generated data with Affymetrix Human Origins (HO) SNP genotyping data^58,117,119,120^, which allows us to leverage data from a more genetically diverse range of present-day Native American populations. We refer to this as the ‘2M.HO’ dataset. For PCA, we merged these data with a dataset of modern Native American populations genotyped on an Illumina SNP array using unmasked and unadmixed individuals only^121^. We refer to this as the ‘2M.Illumina’ dataset.

#### Radiocarbon (^14^C) dates and isotope data

We report 37 new direct radiocarbon (^14^C) dates on bone fragments generated using accelerator mass spectrometry (AMS) (**Table S1**). Thirty-three dates correspond to individuals with genetic data passing quality control. Most dates were generated at the Pennsylvania State University (PSU) Radiocarbon Laboratory (PSUAMS; n=25, with sample preparation and chemistry for four samples performed at the University of New Mexico (UNM) Center for Stable Isotopes (CSI)), while others were generated at the University of Georgia (UGA) Center for Applied Isotope Studies (CAIS) (UGAMS; n=2) and the University of California, Irvine’s (UCI) W. M. Keck Carbon Cycle Accelerator Mass Spectrometer Facility (UCIAMS; n=12). Dates for I28092 were generated at both PSU and UCI and we report the report the statistically combined date obtained in OxCal^122^ (v.4.4) using the Combine() function (figures in **Supplementary Information 1**). This is possible because agreement indices (Acomb = 121.5%; χ² test, df = 1, T = 0.209) indicate statistical consistency between the two measurements. We calibrated ^14^C ages in OxCal^122^ (v4.4) using the IntCal20 Northern Hemisphere curve^123^ and report 95.4% CI calibrated radiocarbon ages. For 23 additional individuals, genomic data are newly reported here, but ^14^C dates for these individuals were previously reported^18,22,124–126^. In the absence of direct dates for individuals with genomic data, we inferred dates based on context dates from archaeological strata (layers) or based on archaeological context information.

For 33 individuals with genome-wide data passing quality control and direct ^14^C dates newly generated for this work, we also report dietary isotope information (**Table S14**). Dietary isotope data were generated for 19 samples following sample preparation and chemistry at PSU with measurement at Yale University’s Analytical and Stable Isotope Center (YASIC; n=13) or Penn State’s LIME facility (n=6), for four samples at CSI, for 10 samples at UCI, and for one sample at CAIS. Dietary isotope data for I28092 were generated at both PSU/YASIC and UCI, producing nearly identical δ¹³C and δ¹⁵N values. Details of isotope analysis can be found in **Supplementary Information 2**.

### QUANTIFICATION AND STATISTICAL ANALYSIS

#### Population structure overview with PCA

To visualize population structure, we carried out Principal Component Analysis (PCA) using smartpca v.18711 with the options ‘lsqproject: YES’ to project ancient individuals and ‘newshrink: YES’ to remap the points for the individuals used to generate the PCA axes to the positions where they would be expected to fall if they had been projected, thereby allowing the projected and non-projected individuals to be co-visualized. We first used the 2M dataset in a “worldwide PCA” with eigenvectors computed using present-day Yoruba, French, and Han Chinese individuals to check for non-Native American ancestry in the ancient Colombian individuals newly reported in this work. Based on the PCA, we excluded I24983 from all analyses (**Figure S1**). For comparability with previous work, we used the 2M.Illumina dataset and projected ancient individuals onto eigenvectors computed using 12 individuals from 4 present-day American populations (Maleku, Zapotec, Teribe, Aymara) as in ref. ^47^. We identify seven PCA outliers, although only two (I23673 and I10932) are covered at more than 15k SNPs on the 2M.Illumina dataset (the remaining individuals have lower SNP coverage, and their placement on PCA is likely to reflect limited data rather than genuine ancestry differences).

#### Analyses of genetic ancestry

We computed outgroup *f*_3_-statistics as *f*_3_(A, B; Yoruba) using *qp3pop* (v.701) in ADMIXTOOLS^120^ with the option ‘inbreed: YES’ to investigate the genetic distance between pairs of individuals. We excluded 5 newly-reported pre-Herrera Period people with fewer than 50,000 SNPs covered on the 2M dataset (I24988, I23794, I22329, I22623, and I23787) and grouped Herrera and Muisca Period individuals by site with the exception of three individuals who we did not merge into larger analysis groups (I10932, I22700, and FORM001) (**Supplementary Information 4**). We converted the original *f*_3_ values to distances by taking their inverse (1/*f*_3_) and built a Neighbor-Joining (NJ) tree based on these distances in *R* (v4.4.2) with the package *ape*. We rooted the tree with USA_Ancient_Beringian.SG. To optimize visualization, we applied Grafen’s branch length transformation to standardize tip heights, followed by a power transform of node heights (exponent = 0.5) to expand the scale near the leaves while preserving the overall topology. This approach improves resolution of shallow divergences and makes fine structure among closely related samples more visible.

We computed *f_4_*-statistics using qpDstat (v.1154) from ADMIXTOOLS^120^ with ‘f4mode: YES’ and ‘inbreed: YES.’ In all cases where an outgroup was required, we used Yoruba. Standard errors were from a jackknife procedure with units consisting of 5cM blocks.

We assessed genetic homogeneity with pairwise *qpWave* (v.1600) to test cladality between pairs of individuals or groups in our analysis dataset. We used ‘inbreed: NO’, ‘allsnps: YES’ and set Han.DG as ‘basepop’ for all qpfstats calculations. We used the 2M.HO dataset in order to ensure that our reference set was of a consistent data type. We provide a step-by-step description of *qpWave* analyses in **Supplementary Information 3**.

We called mtDNA haplogroups using HaploGrep2^127^ (**Table S15**) and re-examined in greater detail the mtDNA haplogroup calls of I22495 and I22497 with HaploGrep3^128^. We determined Y chromosome haplogroups using both targeted SNPs and off-target sequences that aligned to the Y chromosome based on comparisons to the Y chromosome phylogenetic tree with Yfull version 8.09 (https://www.yfull.com/). We provide two notations for Y chromosome haplogroups in **Table S1**. The first uses a label based on the terminal mutation, while the second describes the associated branch of the Y chromosome tree based on the nomenclature of the International Society of Genetic Genealogy (http://www.isogg.org) version 15.73 (July 2020).

#### Ancestry modelling

We used the *qpAdm* (v.2092) framework^89^ to model ancestry and estimate ancestry proportions. *qpAdm* uses *f_4_*-statistics to detect shared drift between a target individual or group of individuals and possible admixing source proxies relative to a set of differentially related outgroup populations (the ‘reference set’). It evaluates whether a target can be plausibly modelled as descending from a common ancestor of one or more source populations and produces estimates of ancestry proportions from one or more proxy sources. Models associated with p>0.05 are assessed as being consistent with the data. We used ‘inbreed: NO’, ‘allsnps: YES’ and set Han.DG as ‘basepop’ for all qpfstats calculations. Additional details are in **Supplementary Information 7**.

We used the ‘*find_graphs ()*’ function in the ADMIXTOOLS2 software to identify admixture graphs that fit observed *f-*statistics. This function tries to find the best fitting graph topologies for a set of *f*-statistics. We began with 1000 different random graphs each time and searched graph space to find an optimal graph once per random initialization. We used USA_Ancient_Beringian.SG as an outgroup and included a number of populations from our list of ancient Americans tested in *f_4_*-statistics. We filtered graphs to include only those with Z-score < 3.0 and evaluated fit also based on log-likelihood (LL) score, where low LL scores indicate a better fit. We assessed graphs that included 0-3 mixture events, and present graphs with 2 admixture events as this is the category of graphs with the fewest admixture events that still fits the data. Multiple locally optimal graphs which fit the data were found, and all of them share key features. Multiple graphs being found is not surprising given the complexity of graph space and the fact that admixture graphs are simplifications of an underlying true history which is inevitably more complicated; the goal of admixture graph fitting here is simply to show that the proposed model is plausible. The graphs presented exhibit a good statistical fit and are biologically and historically reasonable. Details are in **Supplementary Information 6**.

#### Inferring relationships and community sizes

We tested genetic relatedness between every pair of newly reported individuals following the method described in ref. ^129^. We additionally analyzed shared genomic segments on the X chromosomes of individuals I22495 and I22497 using *hapROH*^130^ who were initially assessed as father-son, a relationship that would be unexpected given evidence of matrilineal kinship in Muisca communities. Details of this analysis are in **Supplementary Information 5**.

For 173 individuals (23 pre-Herrera Period hunter-gatherers/early food producers, 18 Herrera Period individuals, and 132 Muisca Period individuals) with data from at least 300,000 SNPs overlapping a core set of 2M targeted SNPs, we applied *hapROH* to call ROH (**Table S10**) and estimate effective population size (*Ne*). We estimated *2Ne* values and divided that value by 2 to obtain estimates of *Ne*. In the main manuscript we report the 95% confidence interval (CI) fitted via the likelihood profile. For the estimate of the sizes of hunter-gatherer/early food producer communities reported in the main manuscript, we used 21 individuals after excluding 2 with a sum total >50cM of ROH>20cM. We repeated analyses using 14 individuals from middle Holocene Checua who had a sum total >50cM of ROH>20cM and obtained a similar estimate of 101-127 individuals (95% CI). Finally, we repeated analyses using 11 unrelated individuals from middle Holocene Checua who had a sum total >50cM of ROH>20cM and again obtained a similar estimate of 101-130 individuals. For the estimate of community sizes during the Herrera Period, we used all 18 individuals as none had a sum total >50cM of ROH>20cM. For the estimate of community sizes during the Muisca Period, we used 123 individuals after excluding 9 with a sum total >50cM of ROH>20cM.

### Estimation of admixture dates

We ran the software *DATES*^131^ (version 210) with binsize: 001, maxdis: .20, qbin:10, a random seed, and jackknife: YES. This method *measures* the decay of ancestry covariance to infer the time since mixture and estimates jackknife standard errors. Data are in **Table S25**.

## SUPPLEMENTAL FIGURE TITLES and LEGENDS

**Figure S1.**
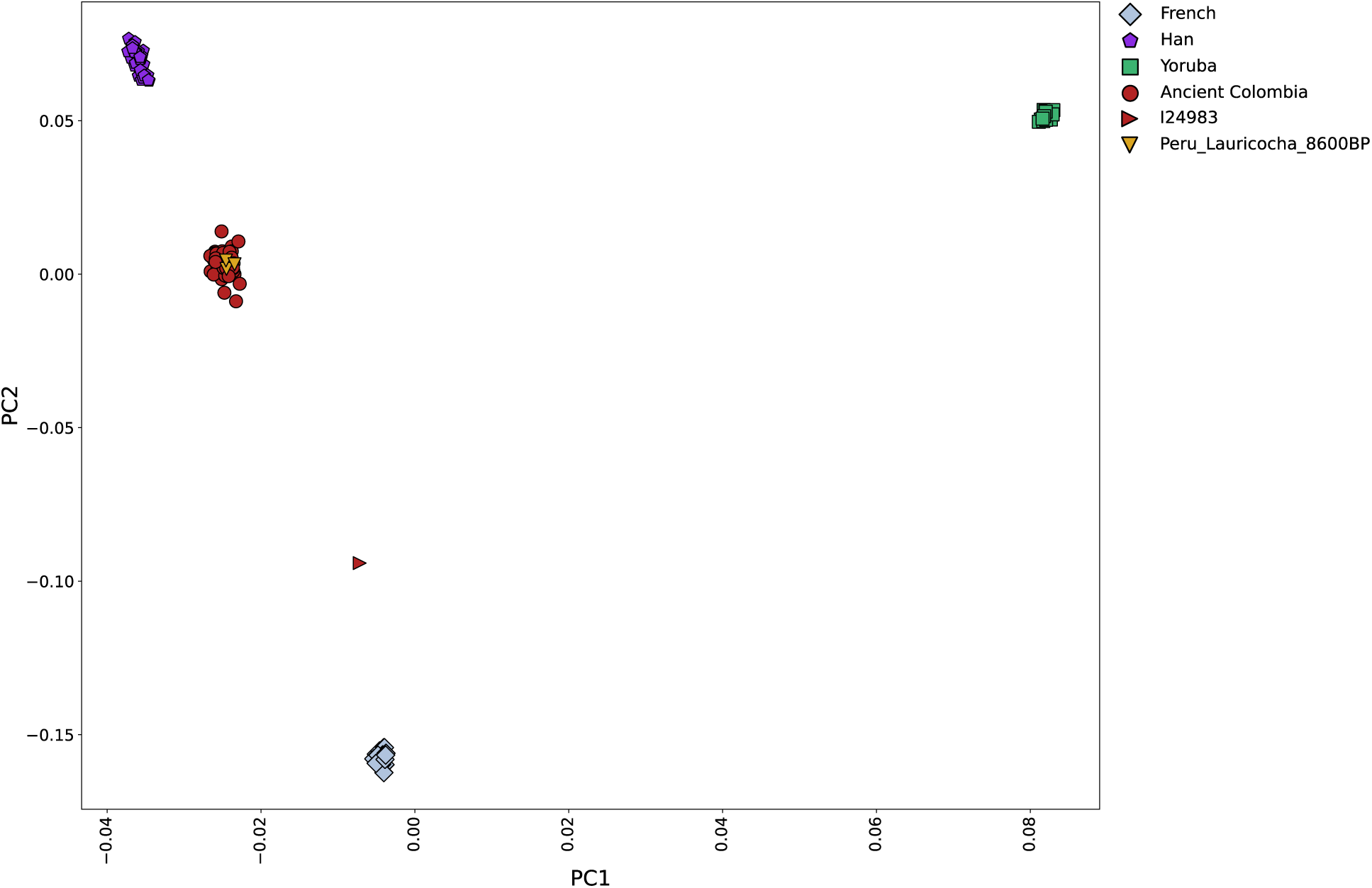
Worldwide PCA with eigenvectors computed using present-day Yoruba, French, and Han Chinese individuals. All ancient individuals were projected. Peru_Lauricocha_8600BP is used to represent unadmixed Native American ancestry. We excluded I24983 from all analyses based on evidence of European-related contamination.

**Figure S2.**
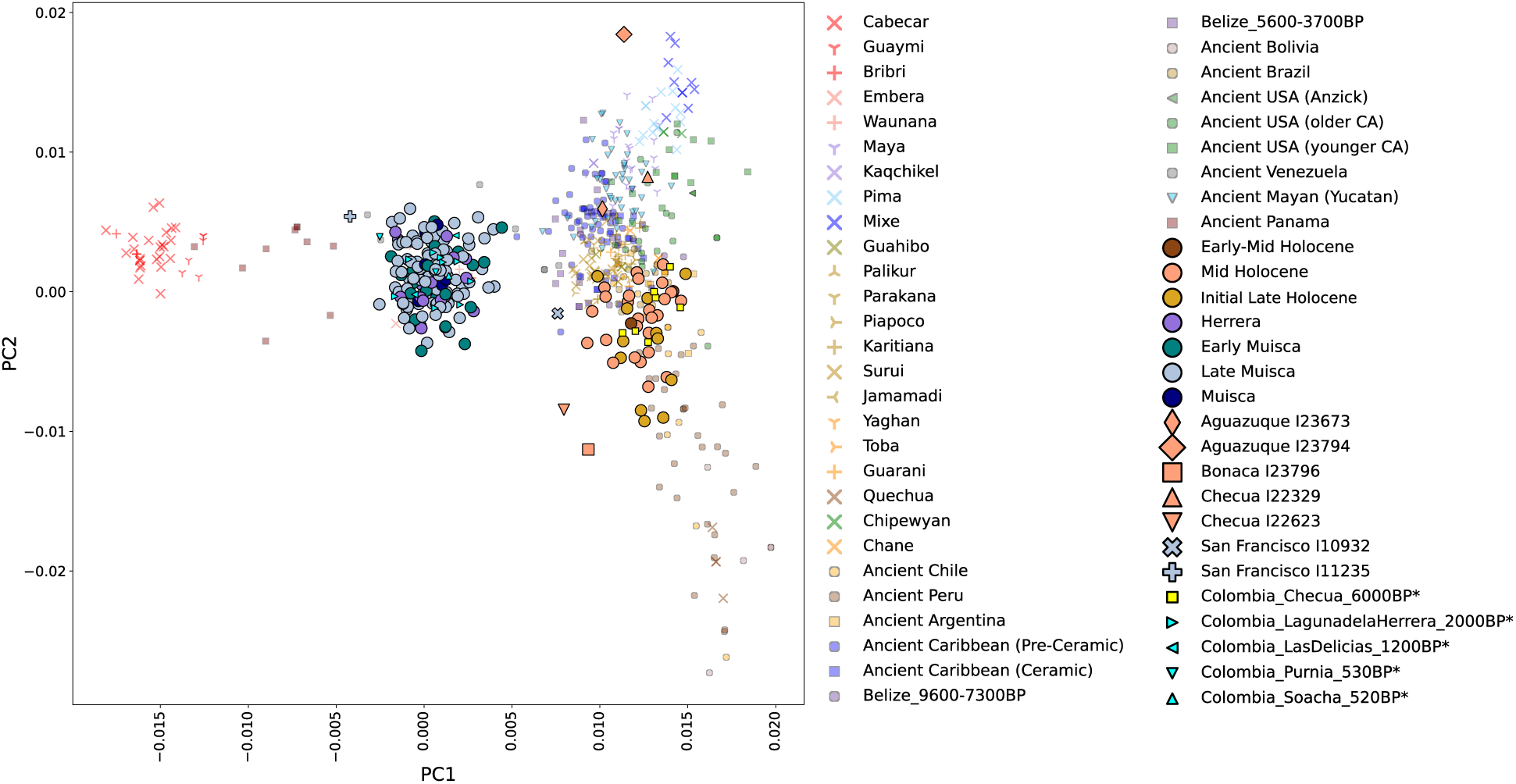
PCA with reference data in color. Newly reported data and published ancient Colombian data from ref. ^36^ represented by large and small colored symbols outlined in black. An asterisk next to a name in the legend indicates its publication in ref. ^36^. A red outline denotes low coverage individuals (<15K SNPs covered on the 2M.Illumina dataset). Two PCA outlier individuals (I23673 and I10932) with >15K SNPs are labeled with diamond and X symbols, respectively.

**Figure S3.**
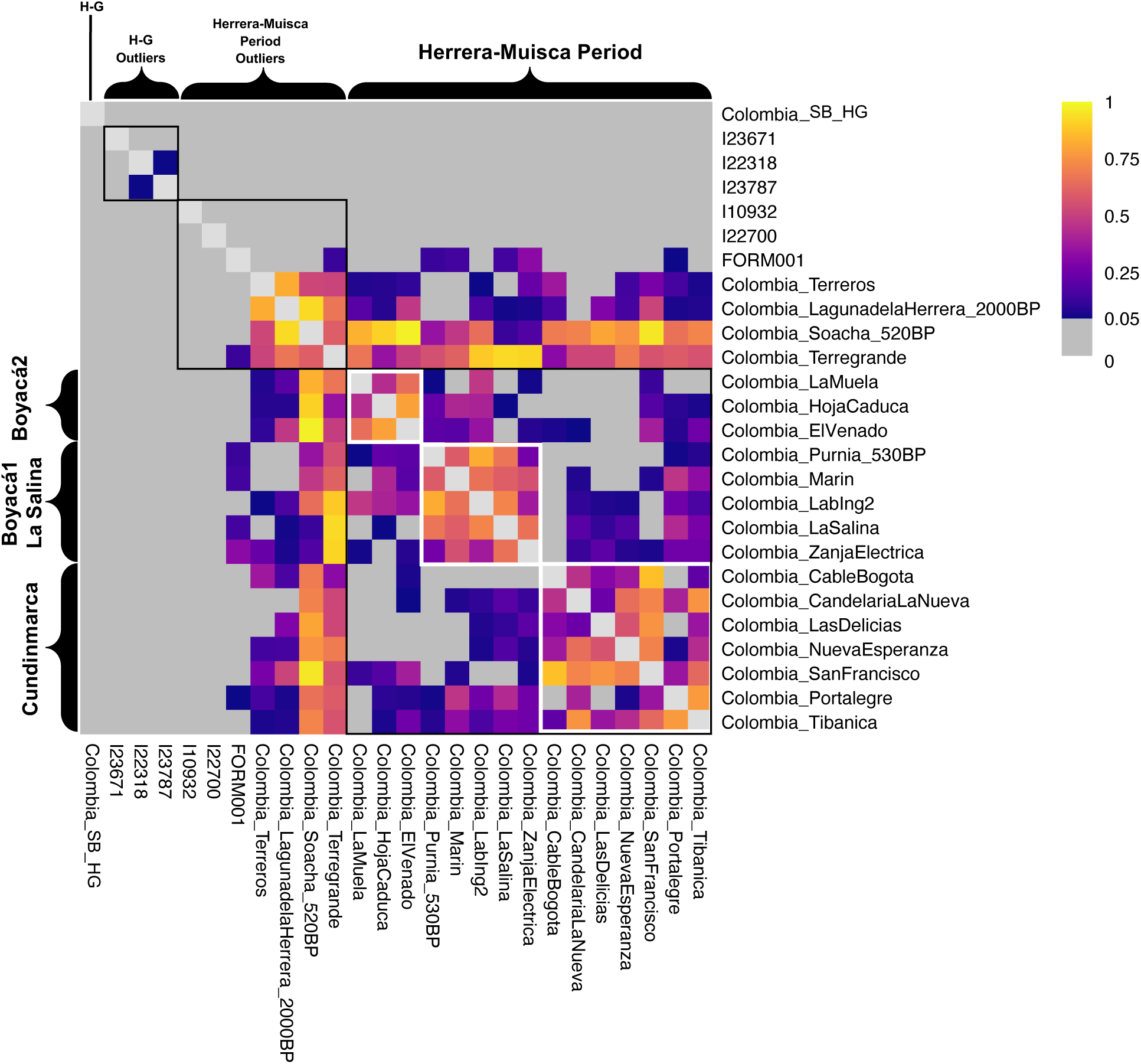
*qpWave* heatmap illustrating genetic differences between pre-Herrera Period people with hunter-gatherer-associated ancestry (H-G) and people from Herrera-Muisca contexts and showing geographic-based genetic substructure within the Herrera-Muisca Period. Heatmap shows pairwise *qpWave* p-values testing whether groups can be modeled as forming a clade relative to a reference set. High p-values (warm-cool colors; p > 0.05) support a clade-like relationship, whereas low p-values (gray; p < 0.05) reject a clade model. *qpWave* analyses are described in **Supplementary Information 3**.

**Figure S4.**
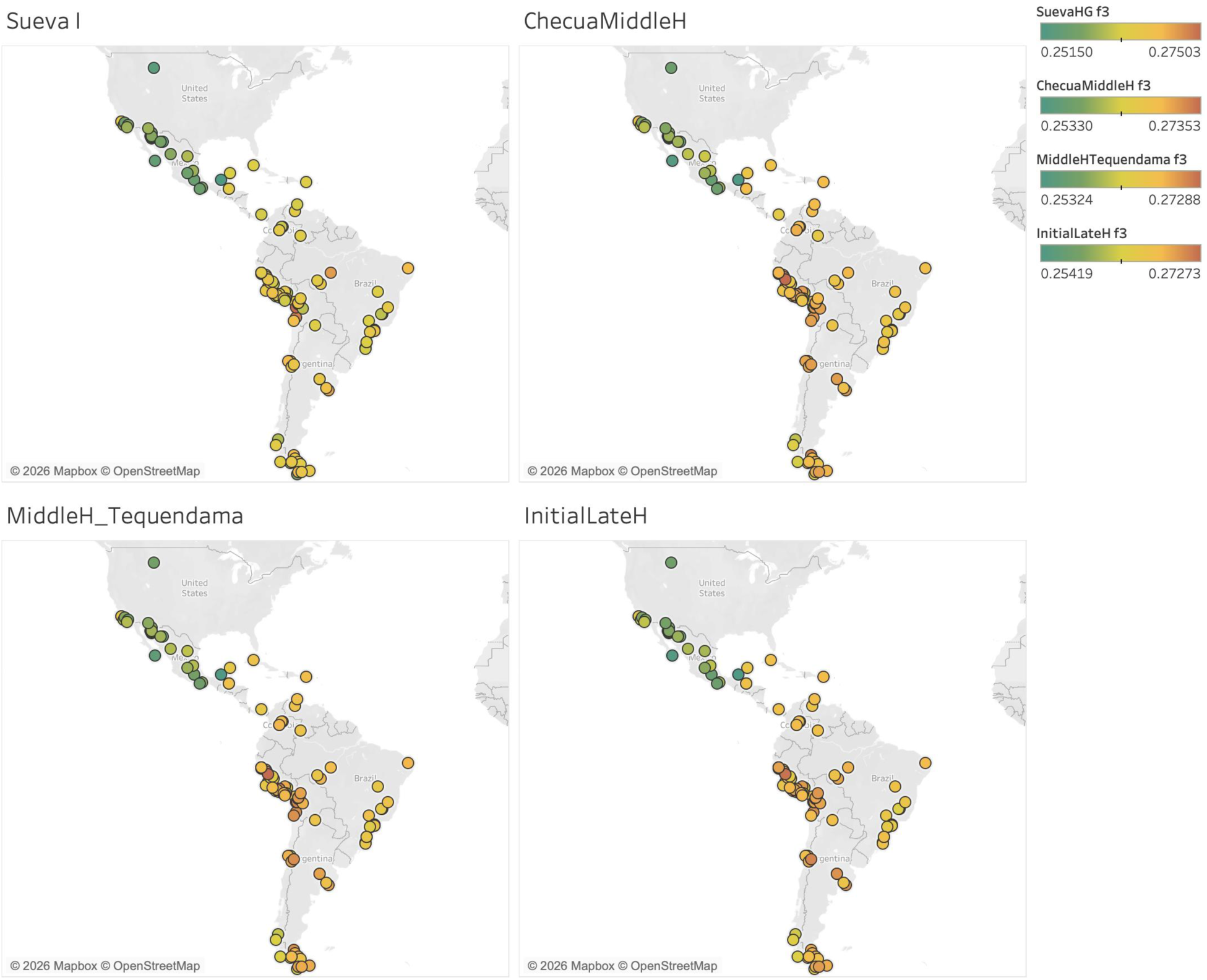
Outgroup-f3 heatmaps showing shared genetic drift between Native American groups and four pre-Herrera Period sub-clades: Sueva I, Checua Middle Holocene, Middle Holocene Tequendama, and Initial Late Holocene (see Figure 3). Each point represents an individual or population plotted at its geographic location (some points jittered slightly for visualization), colored by the magnitude of the *f_3_* value, with warmer colors indicating greater shared drift. Color scales are shown separately for each panel. Basemaps are from OpenStreetMap/Mapbox. Data are in **Table S9**.

**Figure S5.**
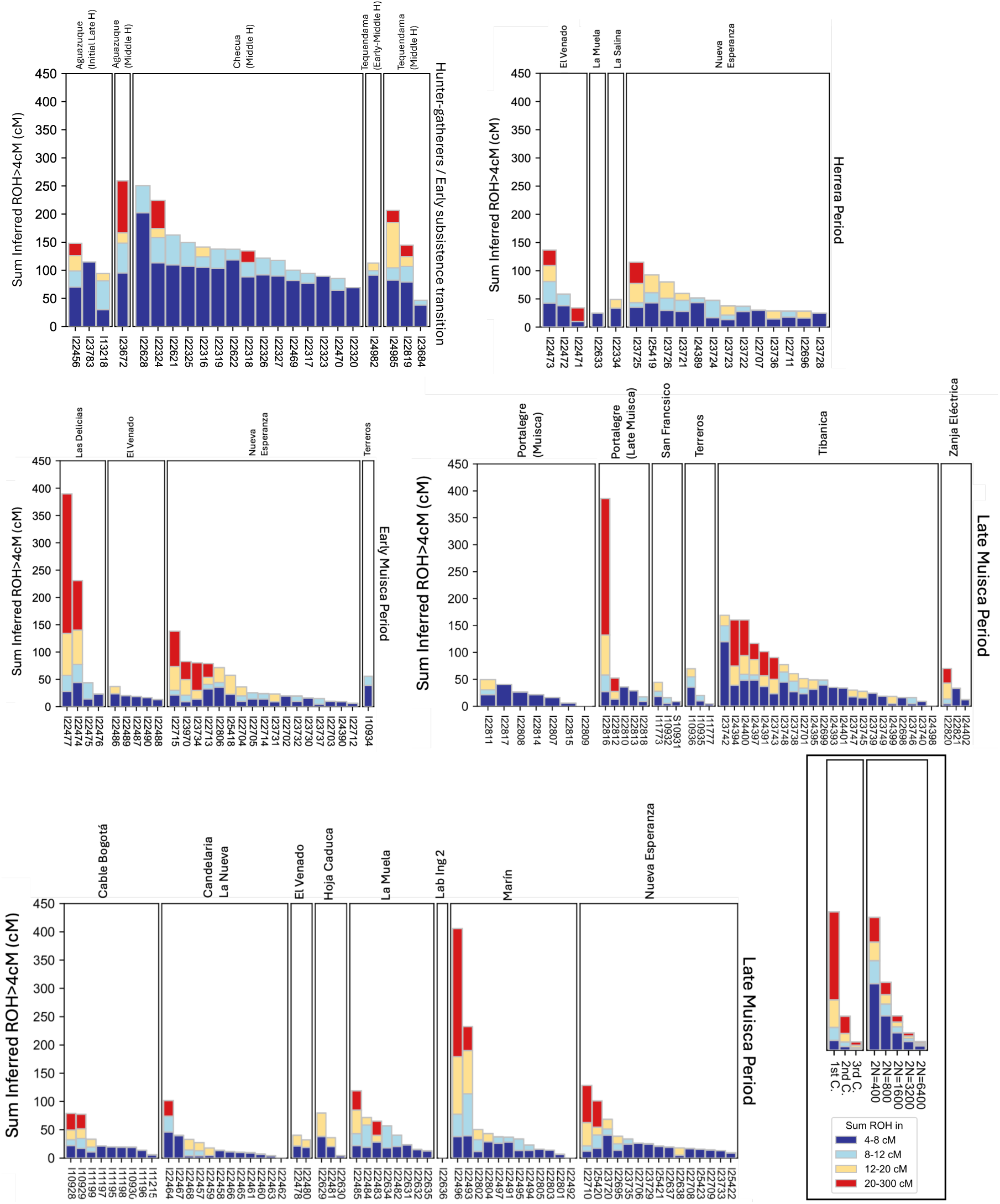
Figure S5. ROH calls for 173 ancient Colombian individuals. We depict inferred ROH for individuals from hunter-gatherer/early subsistence transition contexts (top left), Herrera Period contexts (top right), Early Muisca Period contexts (middle left), and Late Muisca Period contexts (middle right and bottom left). The legend (bottom right) shows analytical expectations calculated using previously-reported formulas^130^ for recent parental relatedness (top right; C represents cousin) and varying population sizes (top left), as well as a color legend for four ROH length categories. Each vertical bar in this figure represents one ancient individual.

**Figure S6.**
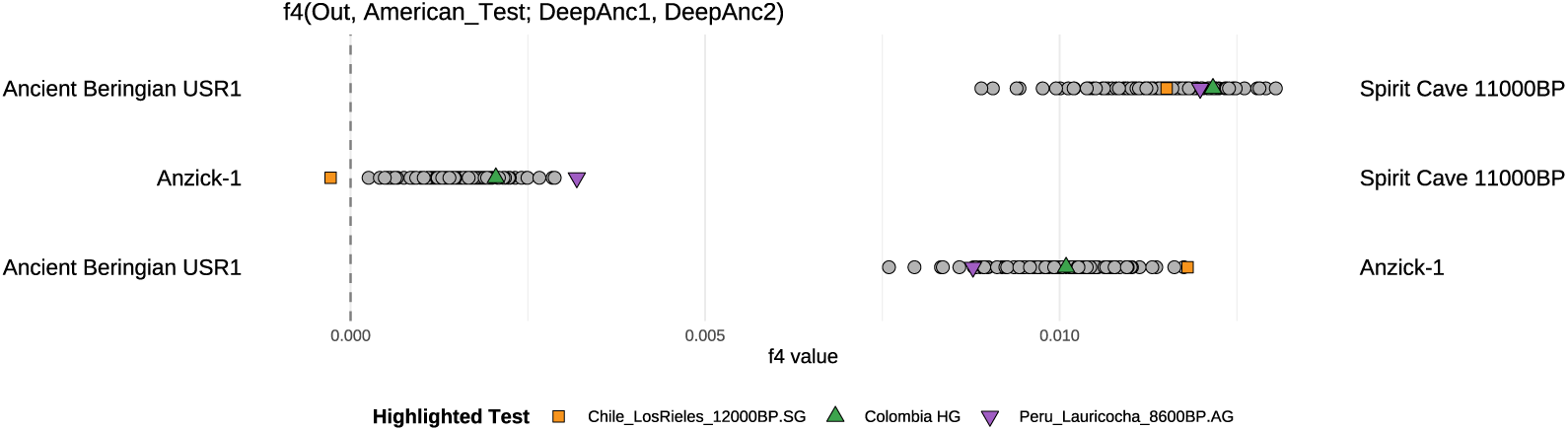
*f_4_*-statistics of the form *f_4_*(Yoruba, *NativeAmerican*_*Test*; *DeepAncient1*, *DeepAncient2*). Colombia_SB_HG (plotted as a green triangle) shows no evidence of ancestry related to Anzick-1 or Spirit Cave 11000BP. The statistic *f_4_*(Yoruba, *NativeAmerican_Test*; USR1, Anzick-1/Spirit Cave 11000 BP), which measures relative allele sharing with Anzick-1 or Spirit Cave 11000BP, indicates that Colombia_SB_HG exhibits affinity to both at a level comparable to other ancient Central and South Americans. In contrast, Chile_LosRieles_12000BP.SG (orange square) shows particularly strong affinity to Anzick-1. Peru_Lauricocha_8600BP.AG (purple upside-down triangle) is a population modeled in previous work as having no Anzick-1-related ancestry and is included here for comparison. Vertical dashed line marks zero. Data are in **Table S11**.

**Figure S7.**
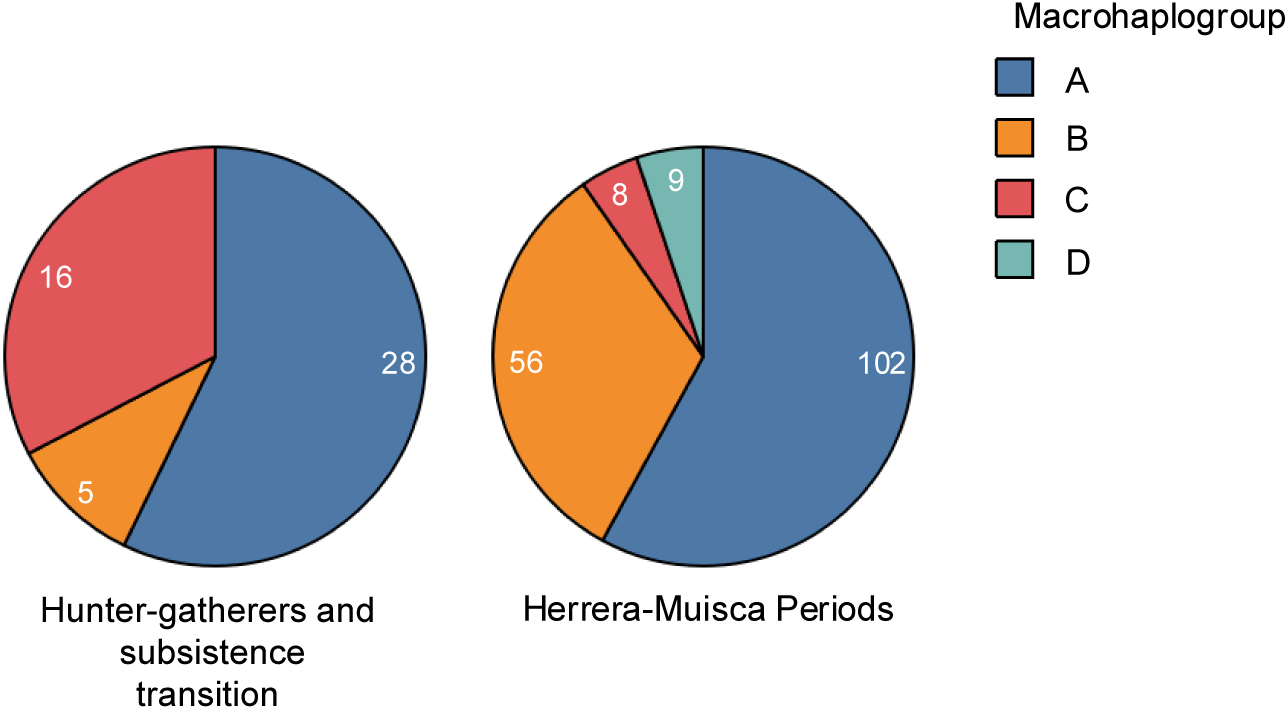
mtDNA macrohaplogroup composition across time on the Altiplano Cundiboyacense. Pie charts show the distribution of macrohaplogroups (A–D) among hunter-gatherers and early food producers (left) and individuals from the Herrera-Muisca Periods (right). Numbers within slices indicate the number of individuals assigned to each macrohaplogroup. The dataset includes 224 individuals in total, of which 20 were previously published^36^. Colors are consistent across panels. Data are in **Table S15**.

**Figure S8.**
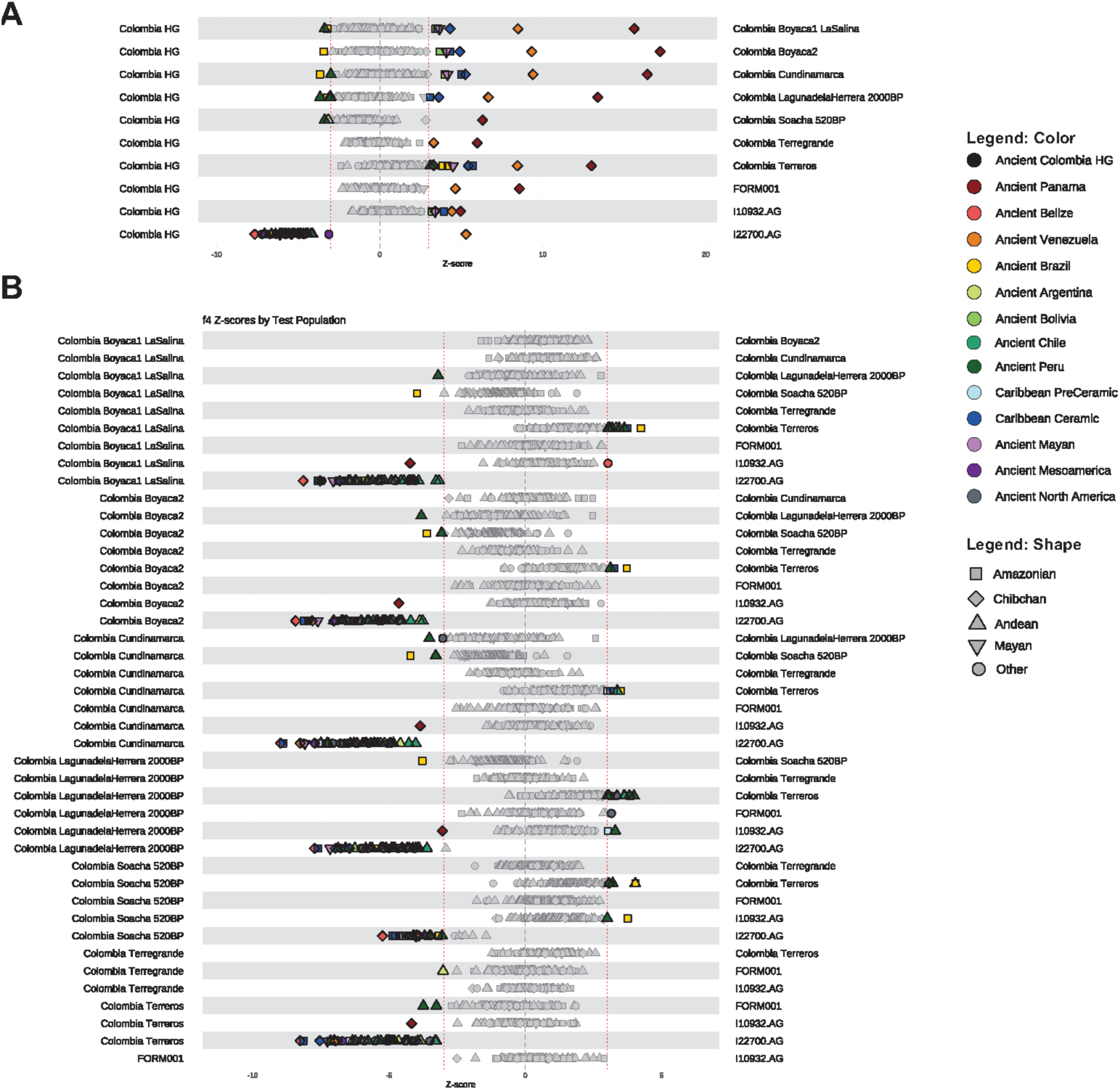
Differences in allele sharing patterns between the people of the Altiplano Cundiboyacense (2M dataset). A) We use the statistic f_4_(Yoruba, *NativeAmerican*_*Test*; Colombia_SB_HG; *Colombia_HerreraMuisca*) to explore differences in allele sharing between hunter-gatherers/early subsistence transition contexts and Herrera-Muisca Period people of the Altiplano Cundiboyacense. Data are in **Table S16**. B) We use the statistic f_4_(Yoruba, *NativeAmerican*_*Test*; *Colombia_HerreraMuisca1*; *Colombia_HerreraMuisca2*) to explore differences in allele sharing between the Herrera-Muisca Period people of the Altiplano Cundiboyacense. Data are in **Table S3.** In both panels, red dotted lines denote |Z|=3. Points associated with |Z|<3 are in gray, while those |Z|>3 are in color, with the legend on the right. The first and second affinity we used to assign the color and shape (respectively) of each *Test* population are included in the table. Only tests based on >50K SNPs are plotted.

**Figure S9.**
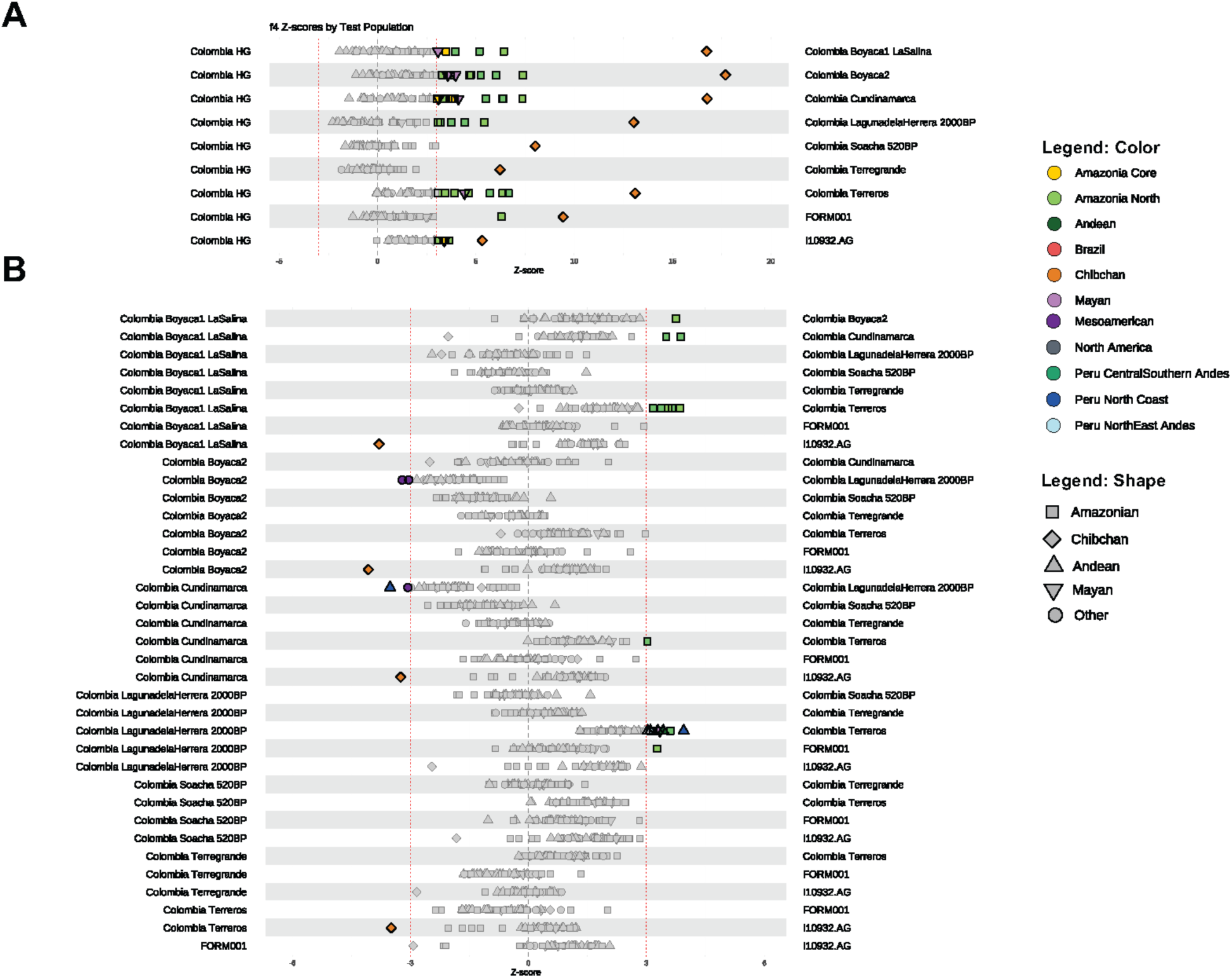
Differences in allele sharing patterns between the people of the Altiplano Cundiboyacense (2M.HO dataset). A) We use the statistic f_4_(Yoruba, *Test_HO*; Colombia_SB_HG; *Colombia_HerreraMuisca*) to explore differences in allele sharing between hunter-gatherers/early subsistence transition contexts and Herrera-Muisca Period people of the Altiplano Cundiboyacense. Data are in **Table S17.** B) We use the statistic f_4_(Yoruba, *Test_HO*; *Colombia_HerreraMuisca1*; *Colombia_HerreraMuisca2*) to explore differences in allele sharing between Herrera-Muisca Period people of the Altiplano Cundiboyacense. Data are in **Table S4.** In both panels, red dotted lines denote |Z|=3. Points associated with |Z|<3 are in gray, while those |Z|>3 are in color, with the legend on the right. The first and second affinity we used to assign the color and shape (respectively) of each *Test* population are included in the table. Only tests based on >50K SNPs are plotted. There were no tests based on >50K SNPs for I22700.

**Figure S10.**
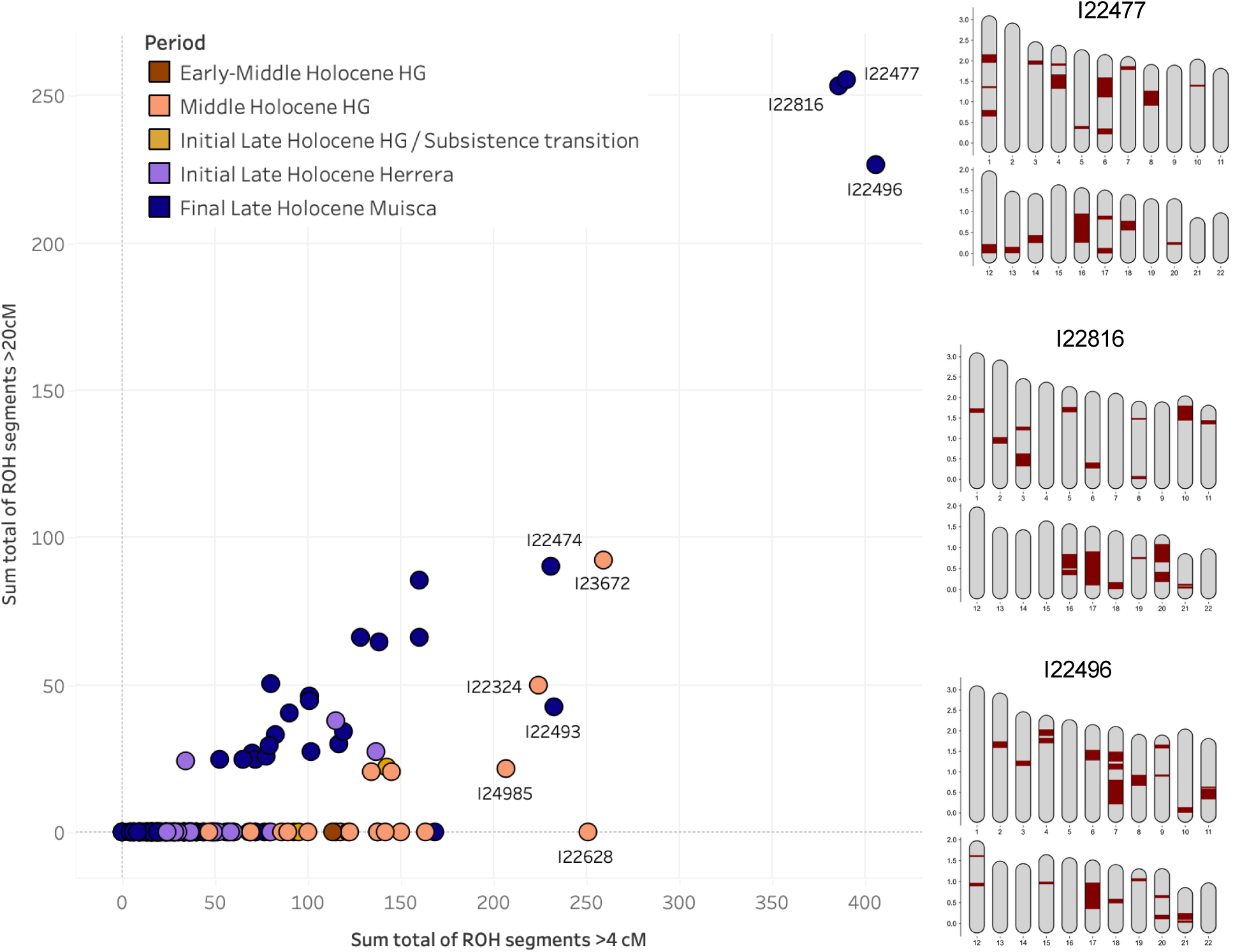
Runs of homozygosity (ROH) across individuals from the Altiplano Cundiboyacense. Left: plot of sum total of sum total of ROH segments >4cM (x-axis) against sum total of ROH segments >20cM (y-axis) for 173 individuals with sufficient coverage (at least 300,000 SNPs overlapping a set of 2M target SNPs). Each point represents one individual, color coded by time period. Nine individuals with >200cM ROH >4cM are labeled. Data are in **Table S10**. Right: karyotypes for three Muisca Period individuals with very high levels of ROH >20cM, indicating consanguinity. Gray outlines represent autosomes and red shaded sections indicate ROH segments.

**Figure S11.**
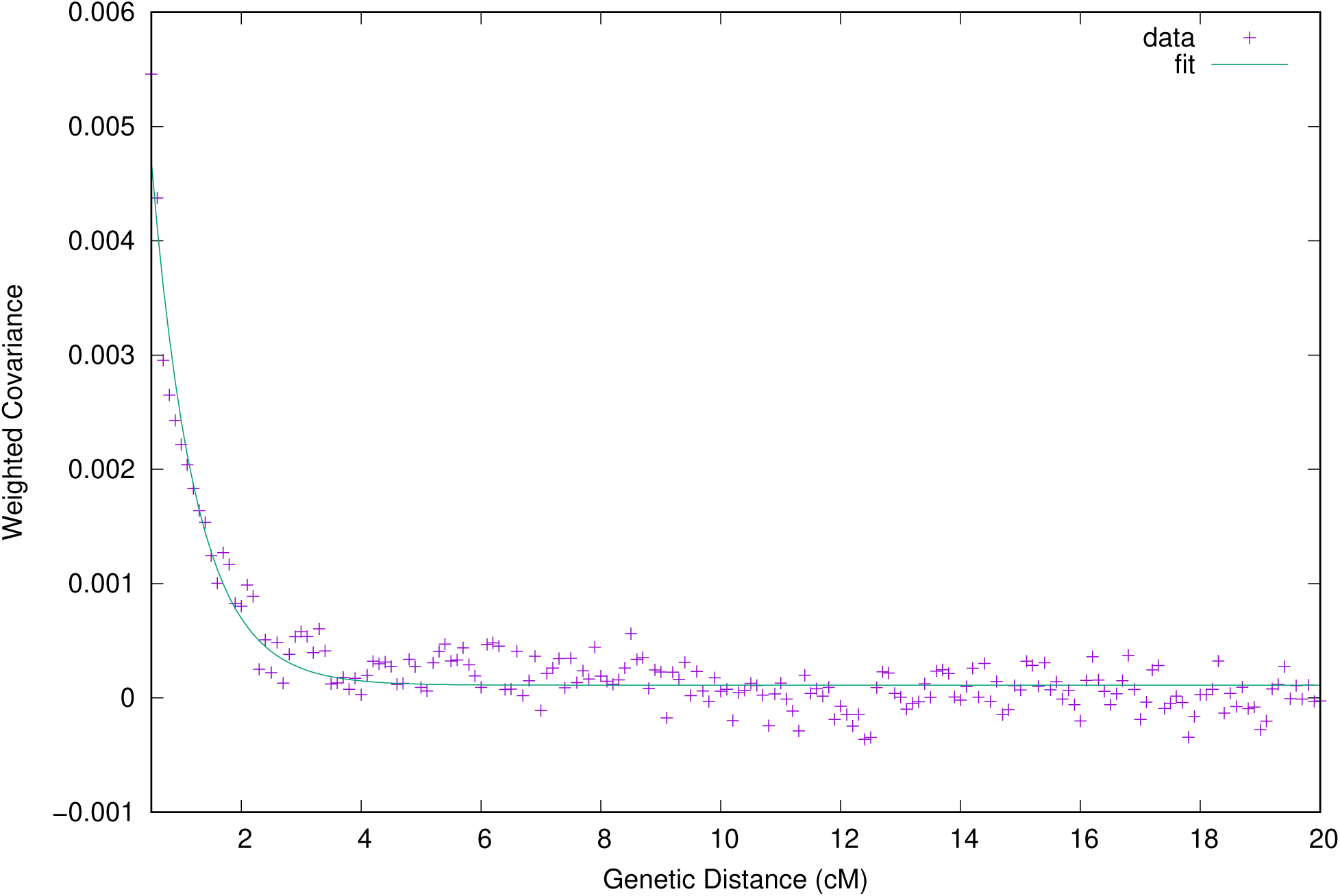
Ancestry covariance decay curve for Herrera-Muisca Period people (target) inferred using a set of pooled ancient and modern Chibchan-related individuals as one reference group and Amazonian-related individuals as a second reference group (composition of groups detailed in Table S25). Weighted ancestry covariance is shown as a function of genetic distance (cM). Purple crosses represent the observed data, and the solid line shows the best-fitting exponential decay model used by DATES to infer admixture timing. The average date of admixture is estimated at 137 *±* 15 generations before the target individuals lived. We observe a strong ancestry covariance decay consistent with admixture, with high statistical support (Z = 8.9).

## Notes

### Competing Interest Statement

The authors have declared no competing interest.

