## Supplemental Information for "Subsistence transition preceded population turnover in the eastern Colombian Andes"

### Supplementary Information

**Supplementary Information 1: Archaeological context**
**Supplementary Information 2: Isotope analysis**
**Supplementary Information 3: Assessing genetic homogeneity with *qpWave***
**Supplementary Information 4: Individuals not incorporated into broader groups**
**Supplementary Information 5: Families**
**Supplementary Information 6: *qpGraph***
**Supplementary Information 7: *qpAdm***
**References**

**Supplementary Information 1. Archaeological Context**

**<u>Pre-Herrera hunter-gatherer and subsistence transition sites</u>**

##### Aguazuque and Tequendama (Cundinamarca, Soacha) – Angélica Triana

***Study Area and Location of Archaeological Sites***

The Sabana de Bogotá is an extensive high-altitude plateau located in the Eastern Cordillera of Colombia. Situated at the geographic center of the country, it maintains an average elevation of 2,600 masl within the municipality of Soacha, Department of Cundinamarca. The archaeological sites of Aguazuque and Tequendama are located within this region, surrounded by the Encantado, Gordo, and Mondoñedo hills[1] (**Supplementary Figure 1.1**).

**Supplementary Figure 1.1.** Location of Aguazuque and Tequendama archaeological sites. Map provided by Angélica Triana.

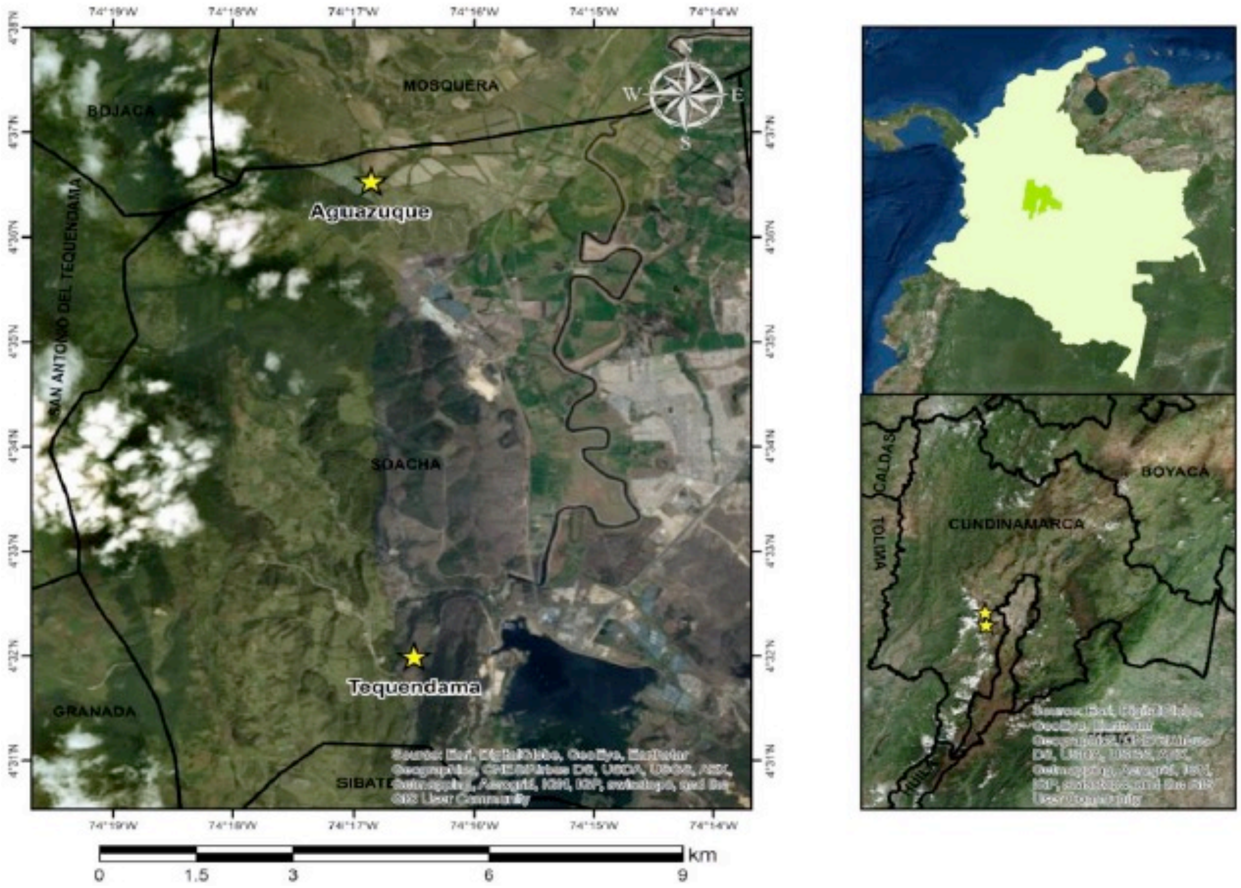


The Sabana de Bogotá comprises two physiographic zones: the high plateau (*altiplano*) and the surrounding mountainous area. It exhibits a bimodal rainfall regime and two thermal tiers: cold (*frío*) and *páramo*. The vegetation in this area responds to altitude and climate, concentrating within two bioclimatic stages: the Andean and High Andean tiers[2-7].

The elevation of these sites ranges between 2,000 and 3,000 masl. Specifically, Tequendama is situated at 2,570 masl and is characterized by montane dry forest, whereas Aguazuque is located at 2,550 masl and is dominated by low montane humid forest[1,8]. Both Tequendama and Aguazuque record a mean temperature between 12°C and 18°C, with an average annual precipitation of 698 mm.

The Tequendama archaeological site is located in a rock shelter 446 meters from the main entrance of the Hacienda Tequendama, oriented at 12º from the Chusacá toll station in the municipality of Soacha, Cundinamarca (**Supplementary Figures 1.2 and 1.3**). Aguazuque is an open-air archaeological site located on private property which, from the 17th century until recently, was known as Hacienda Aguazuque; the site is currently named Hacienda Fute (**Supplementary Figure 1.4**).

**Supplementary Figure 1.2.** Tequendama rock shelter. Photo provided by Angélica Triana.

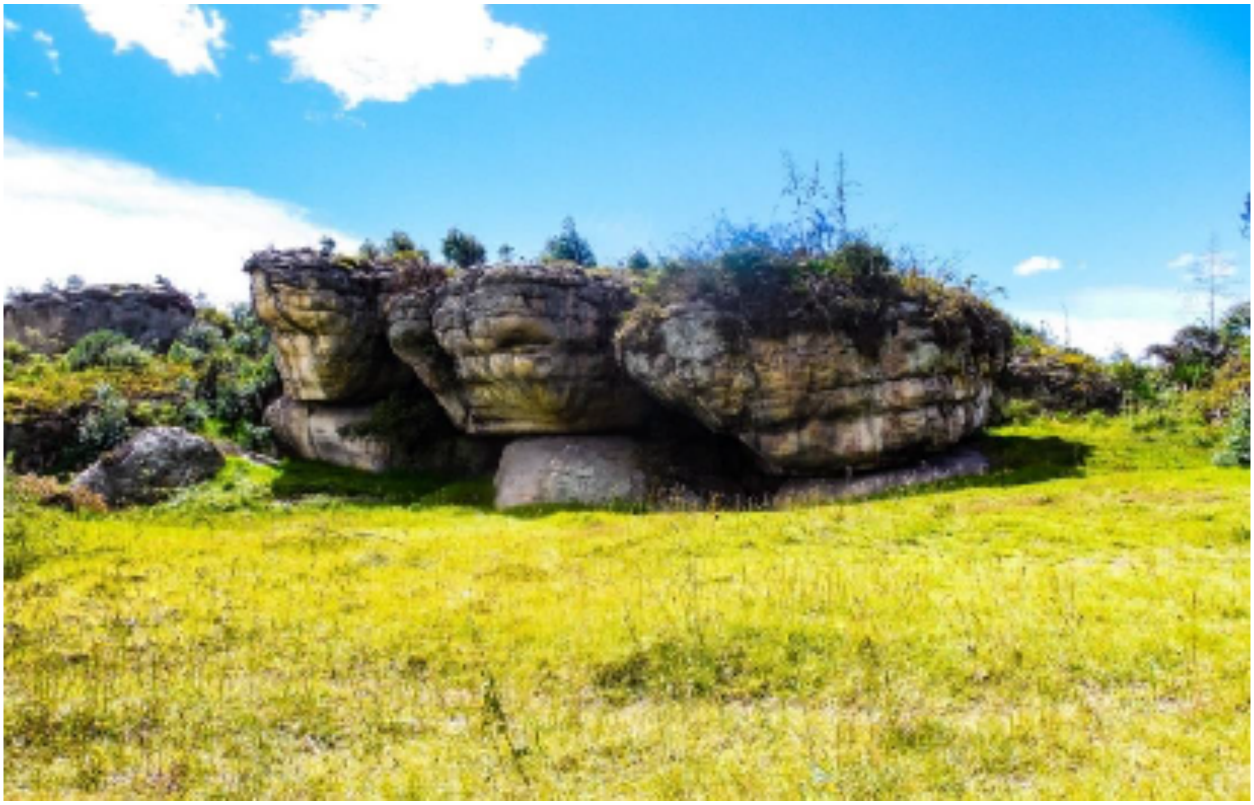

**Supplementary Figure 1.3.** Tequendama rock shelter. Photo provided by Angélica Triana.

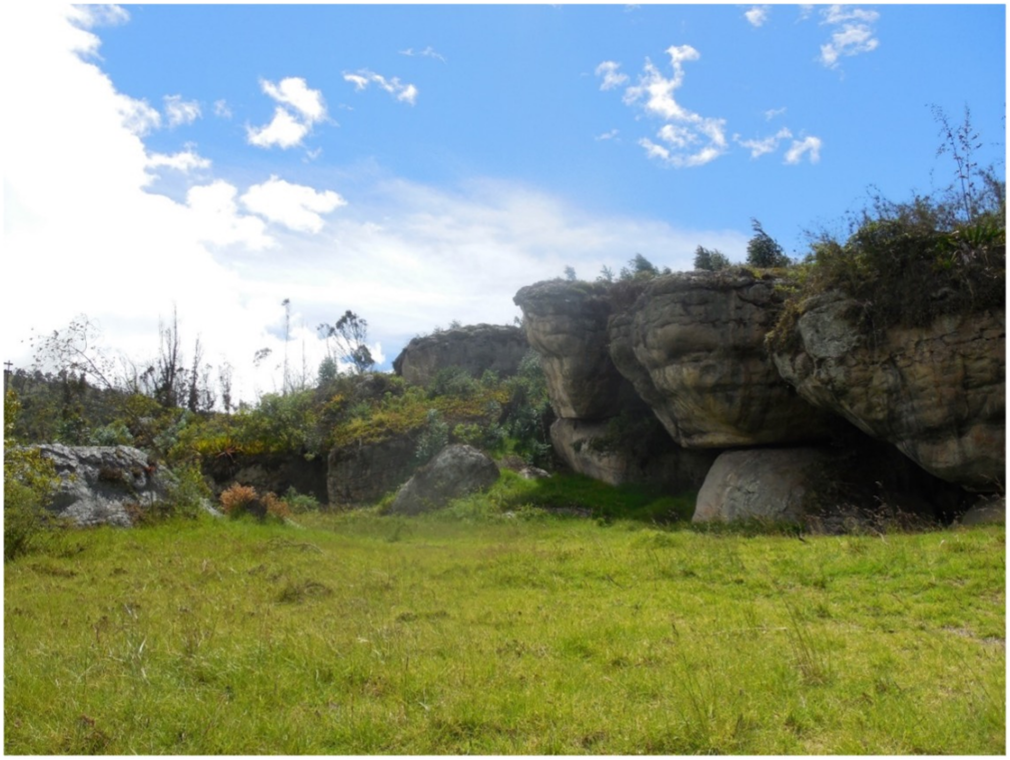

**Supplementary Figure 1.4.** Aguazuque open air site. Photo provided by Angélica Triana.

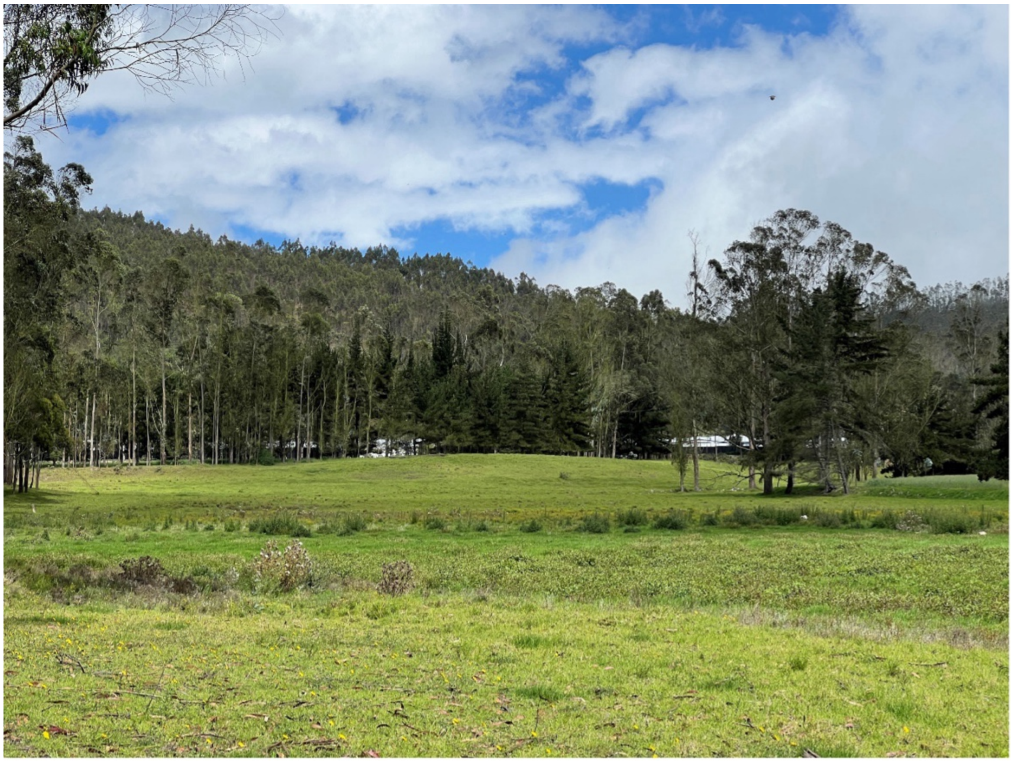

The landscapes of Tequendama and Aguazuque originated through irregular processes due to atypical material deposition, featuring shifts in facies and transport direction. This promoted unique patterns of accumulation and erosion that generated a diverse stratigraphy—specifically, sediment series with unusual proportions characteristic of the Guadalupe Formation (Ksg), which is present at both sites[1].

***Archaeological Context***

The Tequendama and Aguazuque sites represent very different cases; however, both are situated within the framework of pre-agricultural societies. Tequendama is interpreted as belonging to early nomadic hunter-gatherer societies with a high consumption of animal protein. In contrast, Aguazuque is interpreted as a site showing evidence of greater plant dependence and increased sedentism, likely accompanied by higher social complexity[8]. It is noteworthy that both sites were selected due to the evidence of botanical macro-remains and the high frequency of human remains of both sexes. Furthermore, the established occupation dates and the presence of lithic artifacts associated with plant processing—such as hammers, grinding stones (*manos*), and anvils, among others—allow for the understanding and reconstruction of dietary practices developed in each context.

Data regarding the occupational development of Tequendama indicate that the rock shelter site was initially occupied by specialized hunters, followed by hunter-gatherers around 10,000 BP[8] After a long period devoid of data, the first agricultural processes were introduced as an already developed complex, possibly involving human migrations from other areas around 2500 BP[8]. Meanwhile, the archaeological sequence obtained from Aguazuque includes successive occupations. The settlement patterns of these groups differed from those of the hunter-gatherers who occupied the rock shelters of El Abra and Tequendama. Data suggest that by approximately 5000 BP, dwelling sites were no longer located in shelters; populations had adapted to new living conditions, establishing themselves on terraces and low hills outside flood zones, as evidenced by Chía I[9], Vista Hermosa, and Galindo[10].

Based on the above, evidence indicates that while hunting remained a fundamental part of the economy for the Aguazuque population across all occupation periods, gathering was also of great importance as a subsistence activity. Research suggests this based on lithic artifacts such as anvils, hammers, flat mills, and cobbles, indicating an increase in gathering activities. In the words of Correal: "perhaps this increase in gathering activity was the factor that, by broadening the vision of the plant environment—its development and potential utilization—led the groups of the Bogotá Savannah toward the development of horticultural practices around the 4th millennium B.P. This is suggested by the presence of charred plant remains corresponding to plants such as squash (*Cucurbita pepo*) and ibia (*Oxalis tuberosa*)" [1].

The implications of changes in economic activities and settlement have not gone unnoticed. Indeed, a significant portion of archaeological research has focused on understanding dietary patterns; on one hand, through the description of macro-remains[1,8,10,11], and on the other, through isotope studies[12,13]. The latter began with the work of Van der Hammen, Correal, and Van Klinken[12], who analyzed 19 individuals from Tequendama and Aguazuque. They determined a dietary shift from hunting and gathering wild edible plants toward more intensive gathering and, finally, cultivated plants (incipient agriculture) by the end of the human occupation at Aguazuque. Their studies propose that for the period between 3500 and 3000 BP, the diet included maize, and that after 3000 BP, this plant dominated the diet of the inhabitants of these archaeological sites[12].

Another representative study involving stable isotopes in the Sabana de Bogotá reports that at the sites of Tequendama, Aguazuque, Potreroalto, and Checua—dating to the late Pleistocene and the early to mid-Holocene—the human diet was based on the gathering of wild $C_3$ plants associated with tubers, fruits, and nuts from temperate climates[13]. Populations from the mid-to-late Holocene, such as those at Tequendama, show high consumption of meat or certain terrestrial plants, which provides the same increase in $\delta^{15}N_{collagen}$ isotopes[13].

The aforementioned stable isotope studies and the findings of botanical macro-remains at these sites are of great interest for approximating the diet of hunter-gatherer groups in the Sabana de Bogotá. These works have allowed for progress by comparing food consumption patterns of the inhabitants over time. However, no study has yet expanded upon the results of both isotopes and botanical macro-remains, nor has it been established whether dietary differences exist in relation to social organization, specifically gender. This is a significant gap, as it overlooks the fact that, beyond general dietary aspects, hunter-gatherer groups were highly diverse in terms of social organization and gender roles; therefore, it is essential to consider these aspects when studying diet.

Regarding the social organization of Tequendama populations (**Supplementary Figure 1.5**), it has been proposed that they consisted of hunter-gatherer groups throughout nearly all occupation periods. However, evidence appearing around 2500 BP indicates the onset of agricultural processes, judging by the presence of ceramics; nonetheless, it should be noted that the limited evidence prevents a definitive inference of such processes for this period[8]. Regarding Aguazuque, it is stated that during the earliest occupation periods (approx. 8000 BP), societies were organized into hunter-gatherer bands. However, by the 4th millennium BP, these societies transformed into horticulturalists, evidenced by charred plant remains and the discovery of a dwelling floor with a 6-meter diameter containing hearths; this suggests the existence of residences for family units. Additionally, evidence of collective burials and decorated skulls suggests a type of ceremonialism more frequent in horticultural societies than in hunter-gatherer ones[1].

Similarly, human occupation at Aguazuque occurred during the mid-Holocene from approximately 8000 to 2500 BP, with a high presence of human and faunal skeletal remains (**Supplementary Figure 1.6**), lithic artifacts, and plant macro-remains. Aguazuque is perhaps one of the key archaeological sites for understanding domestication processes, resource access, and plant use in Colombia due to its continuous sequence and the abundance of archaeological material[14-16].

Evidence from Aguazuque and Tequendama has allowed for extensive research into human occupation, not only from an archaeological perspective but also through a geoarchaeological approach. This has enabled an understanding of land use and the specific conditions of the pedological, micromorphological, and physicochemical records of the site. The results have yielded interesting data regarding stratigraphic sequences, floor trampling (*apisonamientos*), fire exposure, and various anthropogenic alterations made to the space[15,17,18]. These specific studies had not been previously conducted at these sites and have been performed at few archaeological sites in Colombia. Therefore, the understanding gained from the soil and its formation is a valuable contribution to the study of the hunter-gatherer past and soil formation in general.

**Supplementary Figure 1.5.** Individual I22819 from Tequendama. Photos provided by Angélica Triana.

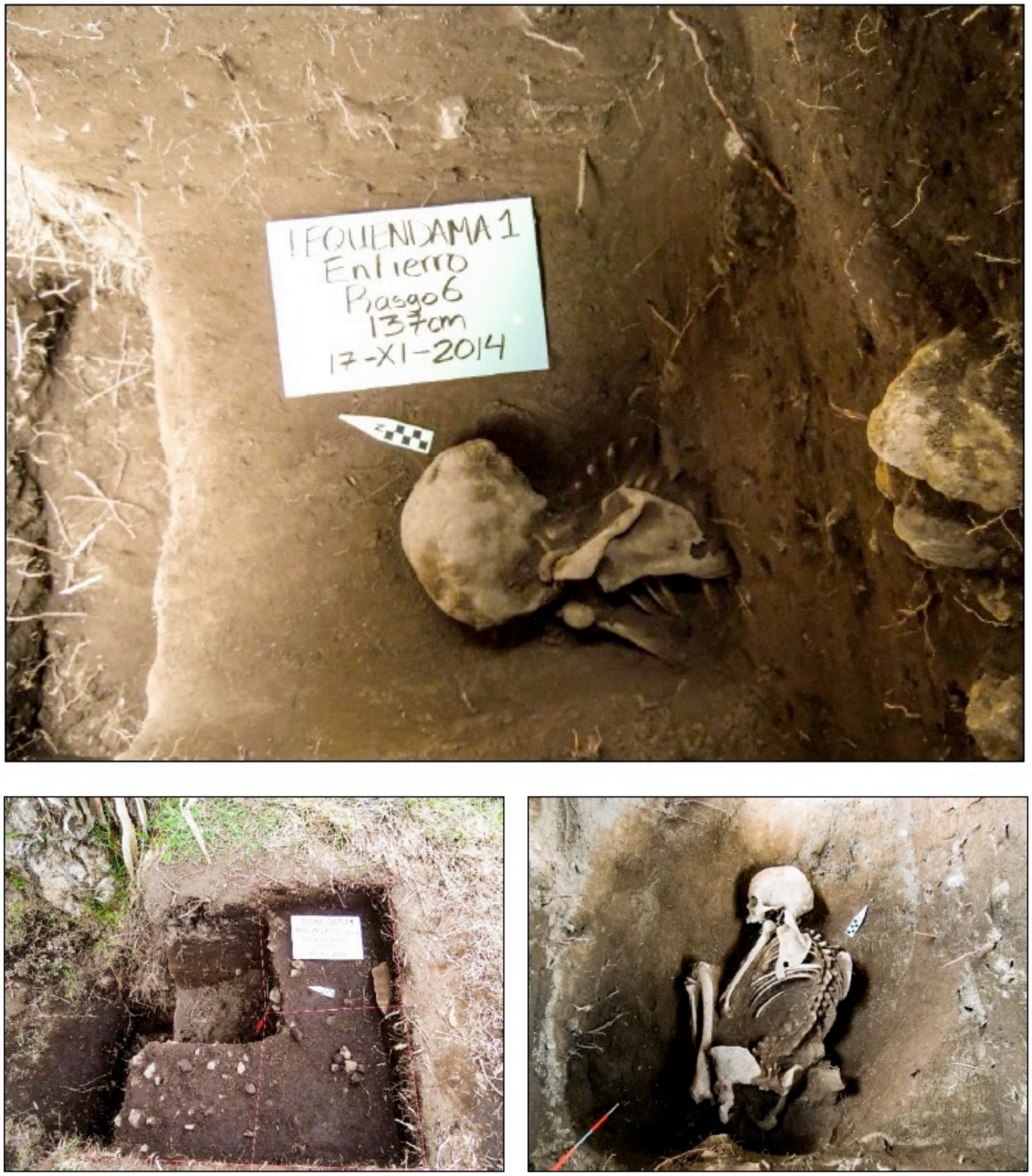

**Supplementary Figure 1.6**. Individual I22456 from Aguazuque. Photos provided by Angélica Triana.

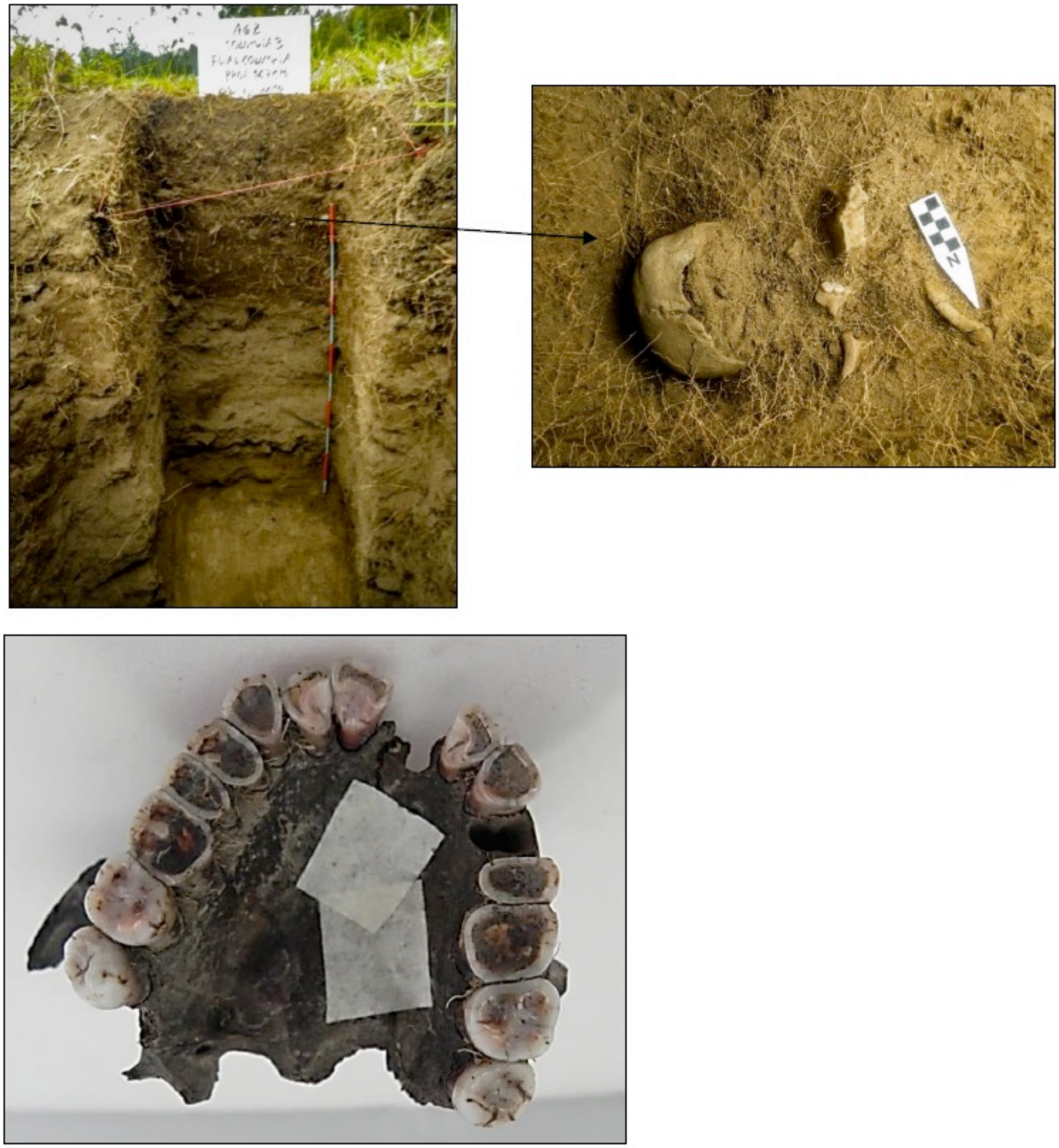

***Área de estudio y ubicación de sitios arqueológicos***

*En la cordillera oriental de Colombia se encuentra una extensa planicie de altura conocida como la sabana de Bogotá, ubicada en el centro geográfico de Colombia, a una altura promedio de 2.600 msnm en el departamento de Cundinamarca, municipio de Soacha. Dentro de la región, encontramos los sitios arqueológicos de Aguazuque y Tequendama, rodeados por el cerro Encantado, el cerro Gordo y el cerro Mondoñedo*[1].

*La sabana de Bogotá comprende dos zonas fisiográficas: el altiplano y la zona montañosa que lo circunda. Presenta concentraciones de lluvia de régimen bimodal y dos pisos térmicos: frío y páramo. La vegetación de esta zona es respuesta a las altitudes y el clima y se concentra en dos pisos bioclimáticos: el piso andino y el alto andino*[2-7].

*La elevación de los sitios está entre 2.000 y 3.000 msnm; sin embargo, Tequendama presenta una altitud de 2.570 msnm y se caracteriza por tener un bosque seco montano, mientras que en Aguazuque la altitud es de 2.550 msnm y predomina un bosque húmedo de baja montaña*[1,8]*. Los sitios arqueológicos de Tequendama y Aguazuque presentan una temperatura media entre 12ºC y 18ºC, el promedio de precipitación media anual es de 698 mm.*

*El sitio arqueológico Tequendama se ubica en un abrigo rocoso a 446 metros de la entrada principal de la Hacienda Tequendama y orientado en dirección 12º desde el peaje de Chusacá en el municipio de Soacha, Cundinamarca. Aguazuque es un sitio arqueológico a cielo abierto se sitúa en un predio privado que, desde el siglo XVII hasta hace algunos años, era llamado Hacienda Aguazuque, actualmente este lugar lleva como nombre Hacienda Fute.*

*Los sitios arqueológicos de Tequendama y Aguazuque presentan paisajes que se originaron de forma irregular, debido a una atípica deposición de materiales con cambios en las facies y en la dirección del transporte, lo que fomentó una acumulación y erosión particular que generaron una estratigrafía diversa, es decir, series de sedimentos con proporciones inusuales características de la formación Guadalupe (Ksg), formación que está presente en ambos sitios*[1].

***Contexto Arqueológico***

*Los sitios Tequendama y Aguazuque representan casos muy diferentes, no obstante ambos se ubican en el marco de sociedades pre-agrícolas. El sitio Tequendama se interpreta como propio de sociedades tempranas de cazadores-recolectores, nómadas, con un alto consumo de proteína de origen animal; Aguazuque se interpreta como un lugar donde existen evidencias de una mayor dependencia de plantas y un mayor sedentarismo, probablemente acompañado de una mayor complejidad social*[8]*. Vale la pena anotar que ambos sitios han sido seleccionados, además de la evidencia de macrorestos vegetales y la alta presencia de restos humanos de ambos sexos; porque las fechas de ocupación establecidas para ambos sitios y el reporte de artefactos líticos asociados a procesamiento de plantas como percutores, manos de moler, yunques, entre otros, permiten comprender y reconstruir las prácticas alimenticias desarrolladas en cada contextos.*

*Los datos acerca del desarrollo de la ocupación de Tequendama*[8] *(sitio explorado por Thomas van der Hammen y Gonzalo Correal, 1977) señalan que el sitio de abrigos rocosos, inicialmente fue ocupado por grupos de cazadores especializados y continuó con cazadores recolectores hacia el 10000 A.P. Después de un largo periodo de ausencia de datos se introducen los primeros procesos de agricultura como complejo ya desarrollado y posiblemente con migraciones humanas desde otras áreas hacia el 2500 A.P aproximadamente*[8].

*Entre tanto, la secuencia arqueológica obtenida del sitio Aguazuque incluye sucesivas ocupaciones, los patrones de asentamiento de estos grupos fueron diferentes a la de los cazadores recolectores que ocuparon los abrigos rocosos del Abra y Tequendama. Los datos permiten inferir que hacia el 5.000 A.P, aproximadamente, los sitios de vivienda ya no lo constituyen los abrigos, las personas se habían adaptado a nuevas condiciones de vida estableciéndose en terrazas y colinas bajas fuera de áreas de inundación como lo muestra la evidencia de Chía I[9], Vistahermosa y Galindo[10].*

*Con base en lo anterior, la evidencia indica que si bien, durante todos los periodos de ocupación la cacería hizo parte fundamental de la economía de la población de Aguazuque; la recolección también tuvo gran importancia como actividad de subsistencia, estas investigaciones lo sugieren, con base en la evidencia de artefactos líticoscomo yunques, percutores, molinos planos y cantos rodados, indicando un incremento de la recolección. En palabras de Correal es: "quizás este incremento de la actividad recolectora, el factor que al ampliar la visión del entorno vegetal, su desarrollo y posible aprovechamiento, condujo a los grupos de la Sabana de Bogotá al desarrollo de prácticas hortícolas, hacia el IV milenio A.P. hecho sugerido por la presencia de restos vegetales calcinados correspondientes a plantas como la calabaza (Cucurbita pepo) y la Ibia (Oxalis tuberosa)"[1].*

*Las implicaciones en cambio de actividades económicas y asentamiento no han pasado desapercibidas. En efecto, parte importante del esfuerzo de los arqueólogos se ha concentrado en entender pautas de alimentación; por un lado a partir de la descripción de macrorestos[1,8,10,11]; por otro a partir del estudio de isótopos[12,13]. Estos últimos inician con el trabajo de Van der Hammen, Correal y Van Klinken[12], autores que analizaron 19 individuos pertenecientes a los sitios de Tequendama y Aguazuque y determinaron un cambio en la dietapasando de la cacería y recolección de plantas comestibles hacia una recolección más intensiva y finalmente plantas cultivadas (agricultura incipiente), al final de la ocupación humana en Aguazuque. Sus estudios plantean que para el períodoentre 3.500 y 3.000 A.P la dieta incluyó maíz y que después del 3.000 A.P dicha planta dominó en la dieta de las personas de estos sitios arqueológicos[12].*

*Otro de los trabajos más representativos que se han realizado en isótopos estables en la sabana de Bogotá reporta que en los sitios de Tequendama, Aguazuque, Potreroalto y Checua, ubicados a finales del Pleistoceno e inicios del Holoceno temprano y medio, la dieta de las personas estaba basada en recolección de plantas silvestres* $C_3$ *asociadas a tubérculos, frutos y nueces de clima templado[6]. También, menciona que las poblaciones que hacen parte del Holoceno medio y tardío, como Tequendama, tienen un alto consumo de carne o de ciertas plantas terrestres, lo cual les proporcionan el mismo aumento de* $\delta^{15}N_{col}$ *en los isótopos[6].*

*Los estudios de isótopos estables mencionados anteriormente y los hallazgos de macrorestos vegetales en estos sitios arqueológicos, son de gran interés para hacer una aproximación acerca del tipo de dieta que poseían los grupos de cazadores recolectores de la sabana de Bogotá. Los trabajos realizados han permitido avanzar al comparar patrones de consumo de alimentos por parte de los habitantes de la Sabana de Bogotá a través del tiempo. Sin embargo, no se ha realizado un estudio que amplíe la información de los resultados de los isótopos y los macrorestos vegetales y tampoco se ha establecido si hay diferencias en la alimentación en relación con aspectos de organización social, específicamente de género. Este es un vacío importante porque desconoce que, además de diferencias en aspectos generales de la dieta, los grupos de cazadores recolectores fueron muy diversas en términos de organización social y roles de género, por lo cual es importante tener en cuenta esos aspectos al estudiar el tema de la dieta.*

*En cuanto a la organización social de las poblaciones de Tequendama se ha planteado que se trató de grupos de cazadores recolectores en casi todos los periodos de ocupación; pero la evidencia que se presenta hacia el 2.500 A.P, aproximadamente, indica el inicio de procesos de agricultura a juzgar por la presencia de cerámica; cabe resaltar que la poca evidencia no permite inferir este tipo de procesos para este periodo de ocupación[8]. En lo que respecta Aguazuque, se afirma que los primeros periodos de ocupación de este sitio 8.000 A.P aproximadamente, las sociedades estaban organizadas en bandas de cazadores recolectores; sin embargo, hacia el cuarto milenio antes del presente las sociedades se transformaron en horticultoras a juzgar por la presencia de restos vegetales calcinados y la evidencia de una planta de vivienda con un diámetro de 6m con fogones en su interior; esto sugiere la existencia de residencias para unidades familiares. Adicionalmente, las evidencias de entierros colectivos y cráneos decorados sugieren un tipo de ceremonialismo que es más frecuente en sociedades horticultoras que cazadoras recolectoras[1].*

*Así mismo, hacia el Holoceno medio se presenta la ocupación humana del sitio arqueológico Aguazuque desde el 8000 al 2500 AP aproximadamente, con alta presencia de restos óseos humanos y de fauna, artefactos líticos y macrorestos de plantas. Aguazuque es quizá uno de los sitios arqueológicos claves para comprender los procesos de domesticación, acceso a recursos y uso de plantas en Colombia debido a la secuencia continua que presenta y la alta presencia de material arqueológico[14-16].*

*Las evidencias de Aguazuque y Tequendama han permitido profundizar en investigaciones amplias acerca de la ocupación humana no sólo desde una perspectiva arqueológica, sino también desde una aproximación geoarqueológica lo cual permitió comprender el uso del suelo y las condiciones puntuales del registro pedológico, micromorfológico y físico químico de este sitio. Los resultados han arrojado datos interesantes que han permitido aproximarse a comprender las secuencias estratigráficas, apisonamientos, exposición al fuego y diversas alteraciones que realizaron los humanos en este espacio[15,17,18]. Estos estudios, no habían sido realizados en estos sitios arqueológicos de forma específica, también han sido realizados en pocos sitios arqueológicos de Colombia. Por lo tanto, la comprensión y los resultados obtenidos a partir del suelo y la formación del mismo, es un aporte muy valioso al pasado de los cazadores recolectores y la comprensión de formación del suelo en general.*

**Bonacá (Cundinamarca, Soacha)**

Bonacá is a middle Holocene open-air site (2,600 masl). Previously reported dates for the site are 6,474–6,276 (5,560±40 $^{14}$C BP Beta-347992) and (5,540±30 $^{14}$C BP Beta-347993)[19].

**Checua (Cundinamarca, Nemocón) – Sonia Archila and Juan Pablo Ospina**

The archaeological site of Checua is located on an isolated hill in Nemocón, Colombia, at coordinates 05º07`24" N and 73º52`50" W, in a transitional zone between the flooding alluvial plane of Checua river and gentle slope hills to the Northeastern edge of the valley in la Sabana de Bogotá, in the eastern range of the Colombian Andes. Since the early 1990's, this site has provided archaeological evidence of early hunter and gatherers societies, who occupied the hill from *ca.* 9000 to 5000 calBP[20,21]. Recent archaeological studies have established that the hill was inhabited in three different moments, described as chronological units 1, 2, and 3[22]. Initially, from 9470 to 8969 calBP (2σ) to 9032-8313 calBP (2σ) (i.e. chronological unit 1), apparently, these early groups visited the place sporadically, and although they left recognizable traces of domestic activity at the site, they did not bury their dead on the hill. Later, during chronological unit 2, from 7580 to 7475 calBP (2σ) to 6398-6280 calBP (2σ) (i.e.), hunter and gatherers groups settled on the hill more intensely, which was inferred by bigger number of archaeological materials on the layers associated to this period, as well as clearer transformations of the occupied zones. It is also remarkable that only from that moment on, the hunter and gatherers began to bury their dead on the hill of Checua. However, during this period, they only performed primary burials by means of sophisticated structures made from wooden sticks[23]. By then, they also established areas for the manufacturing of bone and stone artifacts, hearth zones, refuse deposits and constructed compacted floors.

The third chronological unit of Checua was situated from 5892 to 5659 calBP (2σ) to 5190-5052 calBP (2σ), and archaeological evidence revealed that the hill experienced the densest occupation of all during this period. It is observable the manners in which these groups transformed radically the occupied areas by constructing massive stone floors, domestic structures delimited by posts, compacted living floors, hearth zones as well as by the presence of ritual human and animal burials[22]. It was observed that within this occupation, the concentration of animal bones and lithic objects increased significantly comparing to the earlier moments of occupation, which suggested that hunter and gatherers used the hill to carry forward the process of social life by transforming the landscape as they consolidated physical nodes associated to subsistence, settlement and ritual action. Parallelly, archaeobotany and zooarchaeology studies were conducted, and revealed the use and probable cultivation of plants and domestications of animals[22,24].Chronological unit 3 is also characterized by a strong ritual behavior expressed by the presence of 50 human ritual burials, as well as 2 animal inhumations. Recent works demonstrated that Checua hunter and gatherers constructed varied types of funerary structures, as well as primary and secondary funerary deposits, indicating that the hill was a residential and ceremonial place, intended for the process of dwelling for the living but also for the dead.

Accordingly, by means of diverse lines of evidence, it was proposed that at the beginning of the second chronological unit, at around 7580-7475 calBP, an emerging process of sedentism initiated among the hunter and gatherers that settled on the hill of Checua. It was argued that this process of sedentarization might have increased by the third chronological unit (near 5892-5659 calBP), when the hill apparently had turned into a permanent residence for the hunter and gatherers groups, until it was definitely abandoned at around 5190-5052 calBP.

**Sueva I (Cundinamarca, Sueva) – Angélica Triana**

The Sueva archaeological site is located in the municipality of Junín, Department of Cundinamarca, on the eastern flank of the Eastern Cordillera within the Sueva River basin. This is a transition zone between the Sabana de Bogotá plateau and the foothills descending toward the Eastern Plains (*Llanos Orientales*).

The Sueva rock shelter area is located at an approximate altitude of 2,360 masl within a high Andean forest environment. It possesses a distinct ecology compared to interior Sabana de Bogotá sites (such as El Abra, Nemocón, or Tequendama), suggesting a different adaptation by hunter-gatherer groups.

The artifacts found at this site are predominantly of the "Abriense" tradition. Tools produced through simple percussion on chert flakes were discovered, including scrapers, side-scrapers (*raederas*), and cores. These elements were primarily used for processing hides and meat. Faunal skeletal remains were also identified, including white-tailed deer (*Odocoileus virginianus*) and the intensive consumption of smaller species, such as the guinea pig (*Cavia porcellus*). The faunal remains suggest a subsistence based on deer hunting and, to a lesser extent, animals such as the *curí*, *borugo* (paca), and armadillo. Snails of the genus *Drymaeus* were also identified.

From the recovered plant material (pollen, spores, plant fragments), it was observed that during the deposition of this stratum, the surroundings were covered by alder forests alternating with meadows. The presence of Cyperaceae is linked to humid conditions. The site was thus located at the edge of the Andean forest. The high frequency of lithic reduction waste indicates that tools were manufactured within the shelter. The presence of hammerstones points to gathering activities; this unit features an occupation floor with hearths surrounded by debitage and faunal waste.

*El sitio arqueológico de Sueva se localiza en el municipio de Junín, departamento de Cundinamarca en la vertiente oriental de la Cordillera Oriental y sobre la cuenca del río Sueva. Se trata de una zona de transición entre el altiplano de la Sabana de Bogotá y las estribaciones que bajan hacia los Llanos Orientales.*

*La zona de los abrigos rocosos de Sueva se encuentra a una altitud aproximada de 2.360 msnm en un entorno de bosque altoandino, con una ecología distinta a la de los sitios del interior de la Sabana (como El Abra, Nemocón o Tequendama), lo que sugiere una adaptación diferente de los grupos cazadores-recolectores.*

*En los artefactos hallados en este sitio predominan los artefactos de la Tradición Abriense. Se encontraron herramientas elaboradas mediante percusión simple sobre lascas de chert, tales como raspadores, raederas y núcleos. Estos elementos se utilizaban principalmente para el procesamiento de pieles y carne. También se identificaron restos óseos de fauna de animales como venado de cola blanca (Odocoileus virginianus) y el consumo intensivo de especies menores, cuy (Cavia porcellus). Los restos de fauna sugieren subsistencia basada en la caza de venado y en menor medida de animales como el curí, el borugo y el armadillo. También se identificaron caracoles del género Drymaeus.*

*Del material vegetal recuperado (polen, esporas, fragmentos vegetales) se pudo apreciar que durante el depósito de este estrato los alrededores del sitio estaban cubiertos por bosque de alisos, alternado con praderas. La presencia de Cyperacea se relaciona con condiciones de humedad. El sitio estaba entonces en el límite del bosque andino. La alta frecuencia de desechos de talla indica que las herramientas fueron hechas en el abrigo. La presencia de golpeadores señala que se llevaron a cabo actividades de recolección, esta unidad presenta un piso de ocupación con fogones rodeados por desechos de talla y de fauna.*

***Dating of human remains from Sueva I***

Two direct radiocarbon ($^{14}C$) dates were generated from tooth samples of individual I28092 at two independent laboratories (PSU and UCI; Method Details). The resulting dates of 6592-6439 calBCE (7665±40 BP; PSUAMS-18377) and 6570-6428 calBCE (7640±25 BP; UCIAMS-288521) are statistically consistent and yield overlapping calibrated ranges (**Supplementary Figure 1.7**). Because these dates are consistent within error, they were combined in OxCal[25] (v4.4) using the Combine() function, which calculates an inverse-variance weighted mean. The resulting combined radiocarbon age is 7647±21 BP, which was subsequently calibrated to 6567-6437 calBCE using the IntCal20[26] calibration curve (95.4% CI) (**Supplementary Figure 1.8**).

**Supplementary Figure 1.7.** OxCal calibration and combination of $^{14}C$ dates for I28092. Probability distributions for the two direct $^{14}C$ dates (PSUAMS-18377 and UCIAMS-288521) are shown alongside their combined posterior distribution (I28092_14C_Combined), calculated using the Combine() function in OxCal.

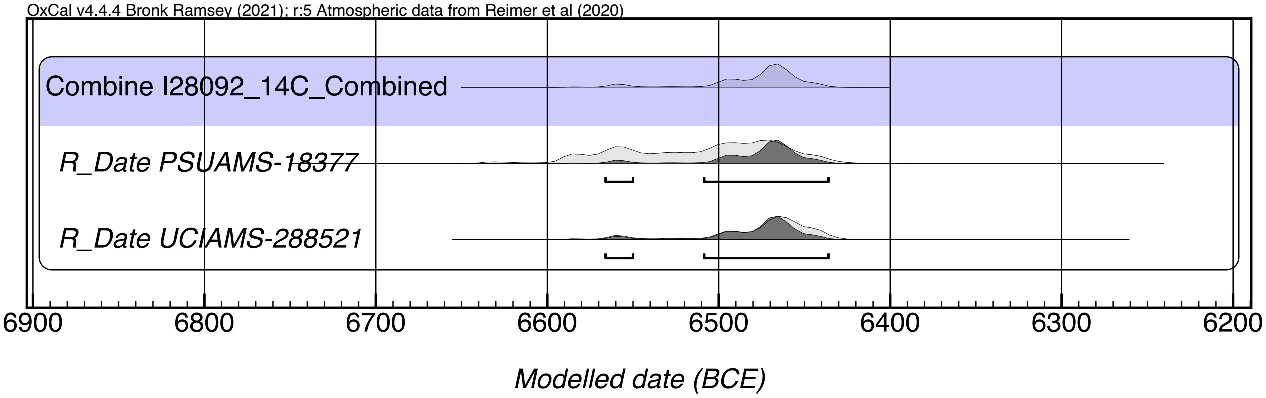

**Supplementary Figure 1.8.** Calibrated probability distribution for the combined radiocarbon date of I28092. The figure shows the posterior calibrated distribution produced by combining two statistically consistent radiocarbon dates. The combined date yields a 95.4% calibrated age range of 6567-6437 calBCE (95.4% CI). Agreement indices (Acomb = 121.5%; $\chi^2$ test, df = 1, T = 0.209) indicate statistical consistency between the measurements.

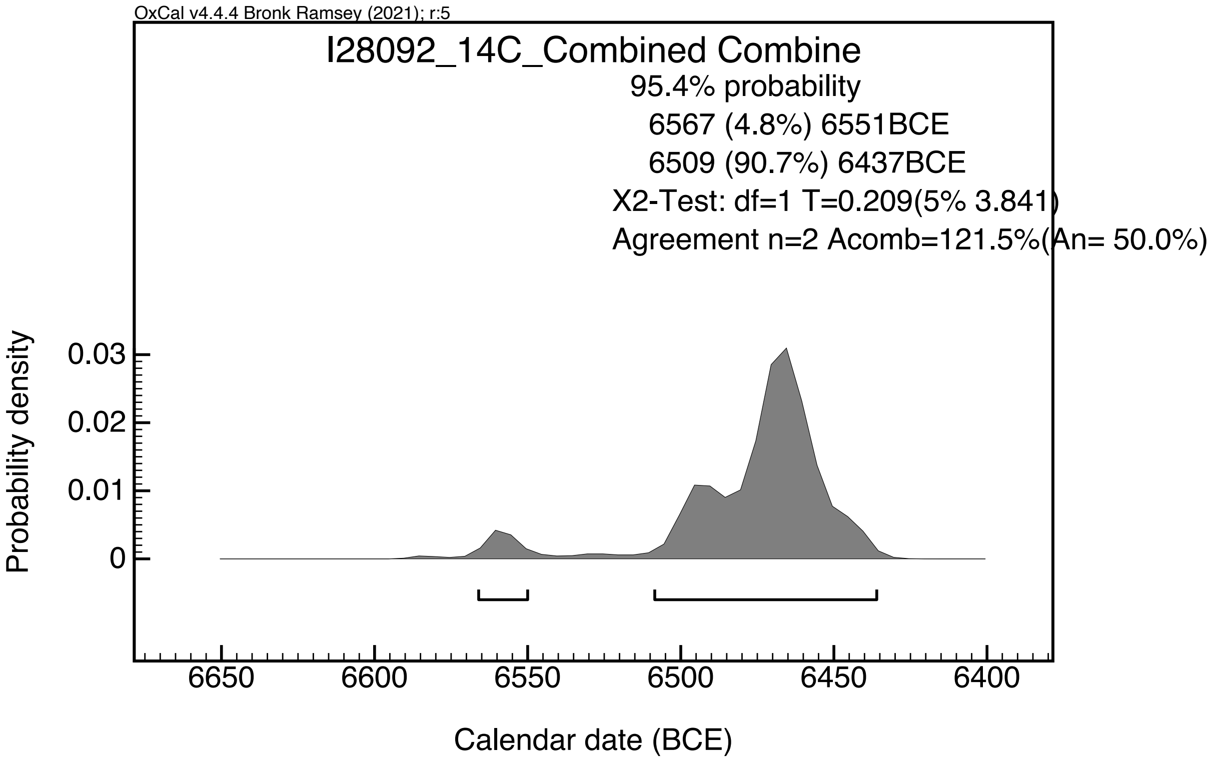


**Ubaté (Cundinamarca, Ubaté) – Sonia Archila and Juan Pablo Ospina**

The Ubaté archaeological site is located on a non-floodable hill in the Sabana de Bogotá, at coordinates 05°19'22.0" N and 73°49'20.6" W. This site was excavated in 2015 as part of a research project by Universidad de los Andes, which revealed evidence for several centuries of occupation by hunter-gatherer groups during the Middle Holocene at the site. Five radiocarbon dates were obtained from human skeletal remains, yielding the following dates: 5545±30, 5655±30, 5700±30 BP, 5400±30 BP, and 5620±30 BP. Five human individuals were excavated at the site, as well as abundant animal bones and lithic objects, which suggested that site was used for both domestic and ritual activity. Archaeobotanical studies conducted at this site have also unveiled important aspects related to the use and consumption of plant resources, demonstrating that the human groups who occupied this hill managed several types of plants, particularly high-altitude Andean tubers. These analyses were performed on sediments extracted from lithic artifacts, as well as micro-remains extracted from human dental calculus.

**Vista Hermosa (Cundinamarca, Mosquera)**

Vista Hermosa is located on the eponymous farm in Mosquera, Cundinamarca, Colombia. G. Correal excavated this site in the early 1980s and published his findings in 1987[27]. It corresponds to a late Holocene hunter-gatherer site where two cultural strata (1 and 2) were found. Stratum 1 contained more cultural evidence than Stratum 2. Stratum 1 yielded an abundance of materials, including faunal skeletal remains, lithic artifacts, and at least seven human burials. The site has close relationships with other hunter-gatherer sites, such as Aguazuque and Chía. Two $^{14}$C dates were obtained from charcoal derived from the burials 1 and 5 respectively: 3,135 ± 35 (GrN-12928) and 3,410 ± 35 (GrN-12929) years $^{14}$C BP. Subsequent studies confirmed the close relationship between the individuals of Vista Hermosa and other mid-Holocene hunter-gatherers from the Sabana de Bogotá region[28-30].

<u>**Herrera-Muisca Period sites**</u>

**Cable Bogotá (Cundinamarca, Bogotá)**

The site is located in the Department of Cundinamarca. We generated a series of $^{14}$C dates on excavated bone material which indicated occupation during the Late Muisca period. The earliest date is 1277-1384 calCE, with multiple tightly clustered dates between 1461-1642 calCE demonstrating continued activity at the site from the 13$^{th}$ through the 15$^{th}$/16$^{th}$ centuries, and possibly into the early 17$^{th}$ century (**Table S1**). This is consistent with a Late Muisca settlement that remained occupied during the period of initial Spanish contact and early colonial transformation.

**Candelaria La Nueva (Cundinamarca, Soacha)**

The Candelaria La Nueva settlement (CLN) (4° 34´8.64” N, 74° 8´58.55” W) is located in an alluvial terrace along the transitional zone between the Eastern mountain range and the valley of the Tunjuelito River. Five circular areas of residence were identified which range from 5.2 to 9.5 meters[31]. Nearly 53 burials were found which were of rectangular and circular shape with abundant biological and cultural remains that include animal remains (deer, armadillo and guinea pig), charcoal, spindle whorls, ground stones, pieces of cooper, pottery, seeds and human skeletal remains[31,32]. The human bodies were buried in supine position following an east and south orientation[33]. A total of 26 individuals of different age and sex have been investigated[32]. Two radiocarbon dates obtained directly from human bone 775 ± 110 B.P (GX-18840-G human bone) and 710 ± 110 B.P (GX-18839-G human bone) which were corrected for isotopic fraction (corrected dates: 999 ± 110 B.P and 934 ± 110 B.P using a mean value $\delta^{13}$C of -11.3‰) place the site during the Early Muisca period[34].

**Cercado de los Santuarios (Boyacá, Tunja) – Pedro Argüello**

In this work, we refer to site names Hoja Caduca, La Muela, Lab Ing2, Zanja Eléctrica, which correspond to archaeological excavation sectors within the broader area of the Cercado Grande de los Santuarios.

The Cercado Grande de los Santuarios (CGS) is an archaeological site located in the city of Tunja, Boyacá (5°33' 94”N, 73°21' 19”W) at 2820 masl, center of Colombia. This name was given by the Spanish in 1539 when the Hispanic city of Tunja was founded[35]. The term "cercado" was the name Europeans used to designate the places where the Muisca chiefs and their relatives lived, and from where they administered and exercised power[36]. The European description indicates that the chief's house and his relatives were surrounded by a fence, hence the term "cercado." At the time of the Spanish arrival in this territory, there were several cercados, probably some in use and others abandoned. The presence of at least eleven cercados have been documented in the Tunja area, and the noun "grande" (large) would imply that this particular one stood out for its size or importance. Sanctuary is the term Europeans used to designate the places, probably bohíos (houses), where rituals were performed and offerings were deposited[37]. In short, in the eyes of Europeans, the Cercado Grande de los Santuarios was, in the mid-16th century, an important residential site for the Muisca elite, where large-scale rituals were performed.

Archaeological investigations carried out since 1937 have demonstrated that this site was used for habitation, burials, and probably rituals, during the pre-Hispanic times. The archaeological importance of the CGS was evidenced by the discovery of monoliths, some of which appear to form a small circular structure[38,39] (**Supplementary Figure 1.9**). Monoliths, and large stone structures, are rare in this region of the country, so the presence of about twenty of them indicates the site's importance. However, the period in which they were crafted, transported to the site, and used is unknown.

**Supplementary Figure 1.9.** Circle of seven monoliths, probably corresponding to the structure of a small house from the Herrera period. Photo by Pedro Argüello.

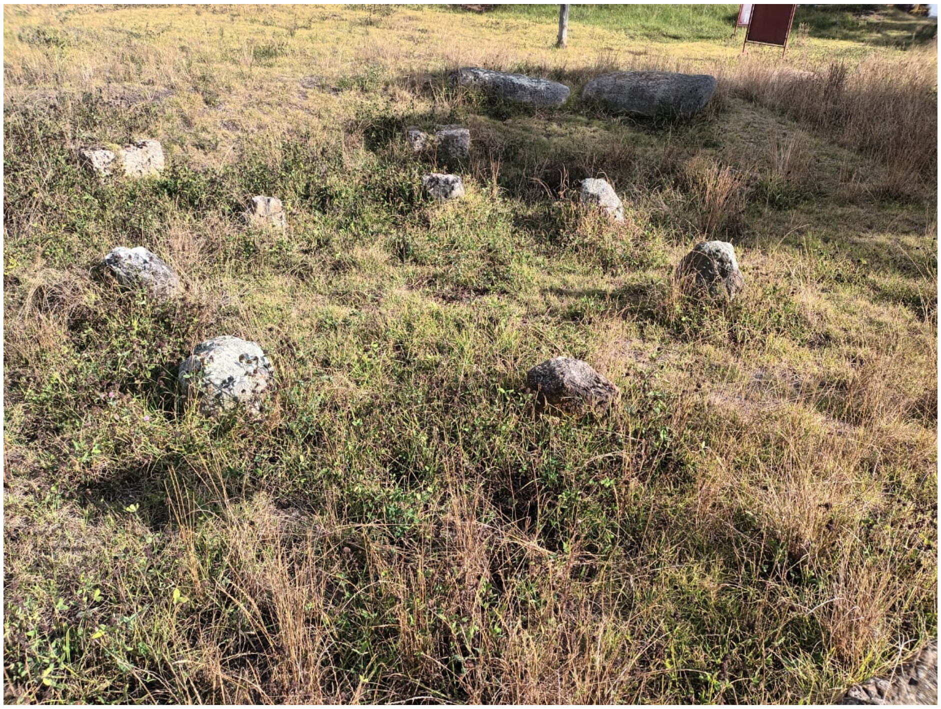

The CGS was continuously inhabited for at least 2,000 years. The earliest settlements date to the Herrera period (2150–1250 BP). These were agricultural communities that used pottery and had permanent settlements. During this period, the CGS was the largest settlement in the entire Tunja region[40], although it contained only a few dwellings, perhaps no more than ten[41,42], and a few burials[43]. The small circular structure may date from the Herrera period, since its excavation yielded pottery sherds typical of this period[38,39], and it is in this area of the cercado that the largest number of domestic structures belonging to this period were located[41,42].

During the succeeding archaeological occupation (Early Muisca period, 1250-950 BP), the settlement grew in size, continued to be the largest in the region, and contained some burials[40]. For the last pre-Hispanic period (Late Muisca, 950-400 BP), although the settlement continued to be of significant size, it was no longer the largest in regional terms and it is even possible that the number of dwellings were reduced[40,42].

The small number of dwellings at the CGS contrasts with the large number of burials. To date, over 400 tombs have been excavated in only a portion of the site. Although few absolute dates exist, changes in burial patterns allow for an approximation of burial typology in relation to the occupation period[44]. It is assumed that the earliest tombs of the Herrera period involved burying the individual in an extended position[43], although tombs found at sites near the CGS contain individuals in fetal or seated positions[45]. Due to the lack of absolute dates, it is not possible to differentiate between tombs corresponding to the Early Muisca period and those of the Late Muisca period. During these periods, individuals were buried in a fetal or seated position, and many of the tombs were sealed with a stone slab (images in ref. [35]). Due to some tombs containing vessels, or fragments of them, it has been possible to assign about twenty tombs to the Late Muisca period.

Although the CGS is assumed to have been an elite site, archaeological data has not yielded convincing evidence of differences that could correlate with sociopolitical differentiation. Comparative analyses of domestic contexts have not found substantial differences between the dwellings in different sectors of the Cercado[41,42,46]. The same is true for funerary contexts. Although some tombs contain grave goods, rare or foreign artifacts power[47,48], its quantity hardly matches the European descriptions for the tombs of the powerful Muisca leaders. Similarly, differences in quality of life and disease incidence do not appear to be correlated with sociopolitical inequality[49]. It is possible that a excavated palisade surrounding a domestic unit corresponds to the cercados described by Europeans as enclosures[50], but, once again, its scale and the labor required to build it is far from the magnificence of the enclosures described by the Spanish.

For this study, we selected samples from 19 individuals and 17 yielded genomic data that are reported here. The tombs come from different sectors of the CGS, including Hoja Caduca, La Muela, Lab Ing2, and Zanja Electrica. One individual (N 37, 42) dates to the Herrera period. (1680 ± 60 BP, Beta 77495) [43], and it is possible that another individual (A 61, 63b) also corresponds to this period. The remaining individuals belong to the Early Muisca and Late Muisca periods. As mentioned, given the lack of variation in burials form and body position between these periods and the absence of absolute dates, it is not currently possible to distinguish the tombs from these two periods. Even so, of these tombs some contain grave goods (vessels or large fragments of broken vessels) that can be dated to the Late Muisca period (i.e. N 56, 61, N54, 62)[35] (**Supplementary Figure 1.10**).

**Supplementary Figure 1.10.** Ceramic vessel found in burial N 56,61. Muisca Tardío period. Photo from the Museo Arqueológico de Tunja, provided by Pedro Argüello.

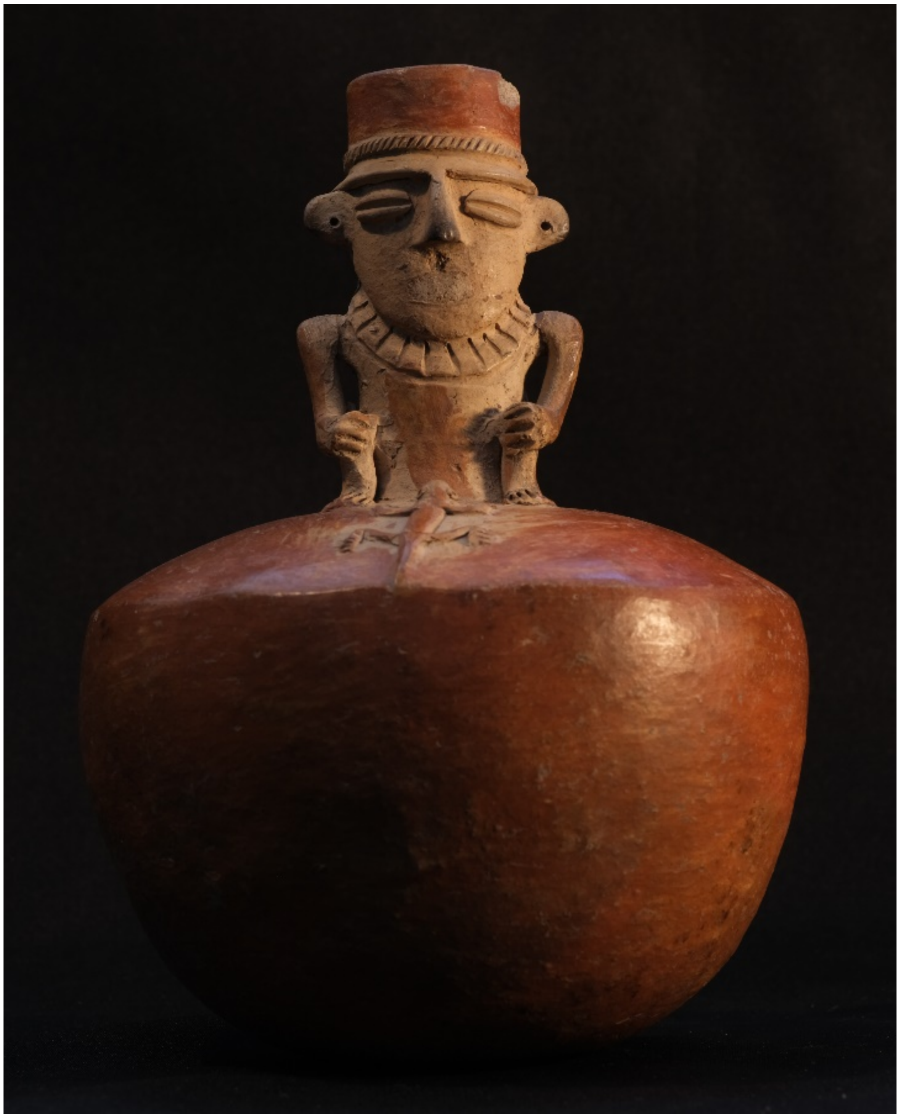

**El Venado (Boyacá, Valle de Samacá) - Ana María Boada**

The archaeological site of El Venado is located at the geographic coordinates 5.501788 N, -73.475234 E, in the Samacá valley, within the municipality of Samacá, about 5 km from Marín (**Supplementary Figure 1.11**).

**Supplementary Figure 1.11. Location of Marín and El Venado archaeological sites.** Map provided by Ana María Boada.

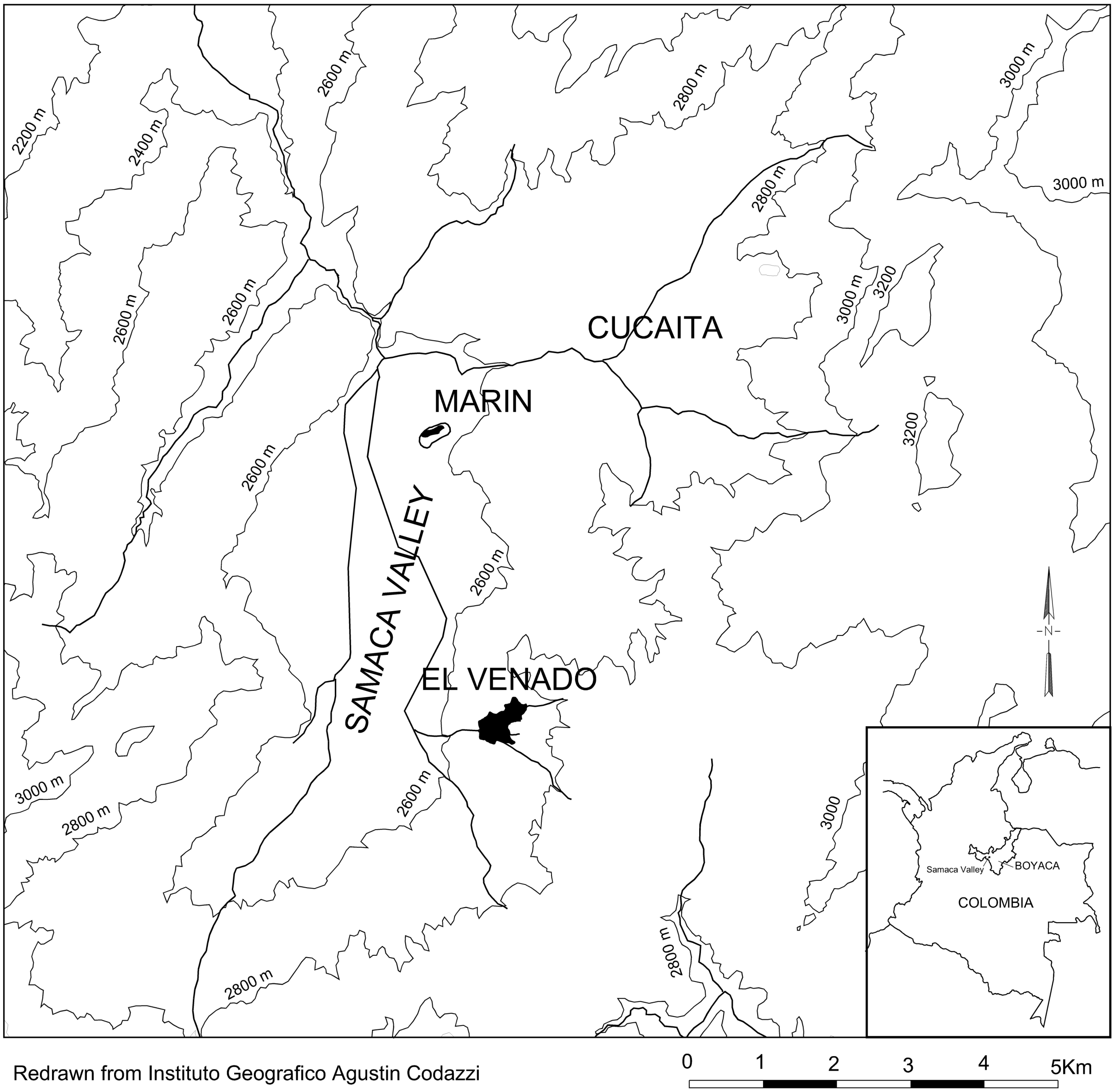


The site sits on a gentle slope, partly eroded and partly covered with fertile soils (**Supplementary Figure 1.12**). This settlement was one of the largest communities in the valley and had the longest continuous occupation, beginning in the Late Herrera period and continuing into modern times[45]. At some point, it was contemporary with the Marín site. Previous research aimed to analyze the development of sociopolitical inequality across the cultural sequence and to identify sources of political power. Two contrasting bases of power were studied at El Venado: one based on prestige obtained through gift-giving, manipulation of symbols and prestige goods, feasting, and ceremonial exchanges; the other based on control over basic resources and wealth accumulation. The study assessed the roles of both bases of sociopolitical hierarchy and how they interacted throughout the cultural sequence of El Venado. The full report on the El Venado site is available in ref. [45].

**Supplementary Figure 1.12**. **View of the El Venado archaeological site.** Photo provided by Ana María Boada.

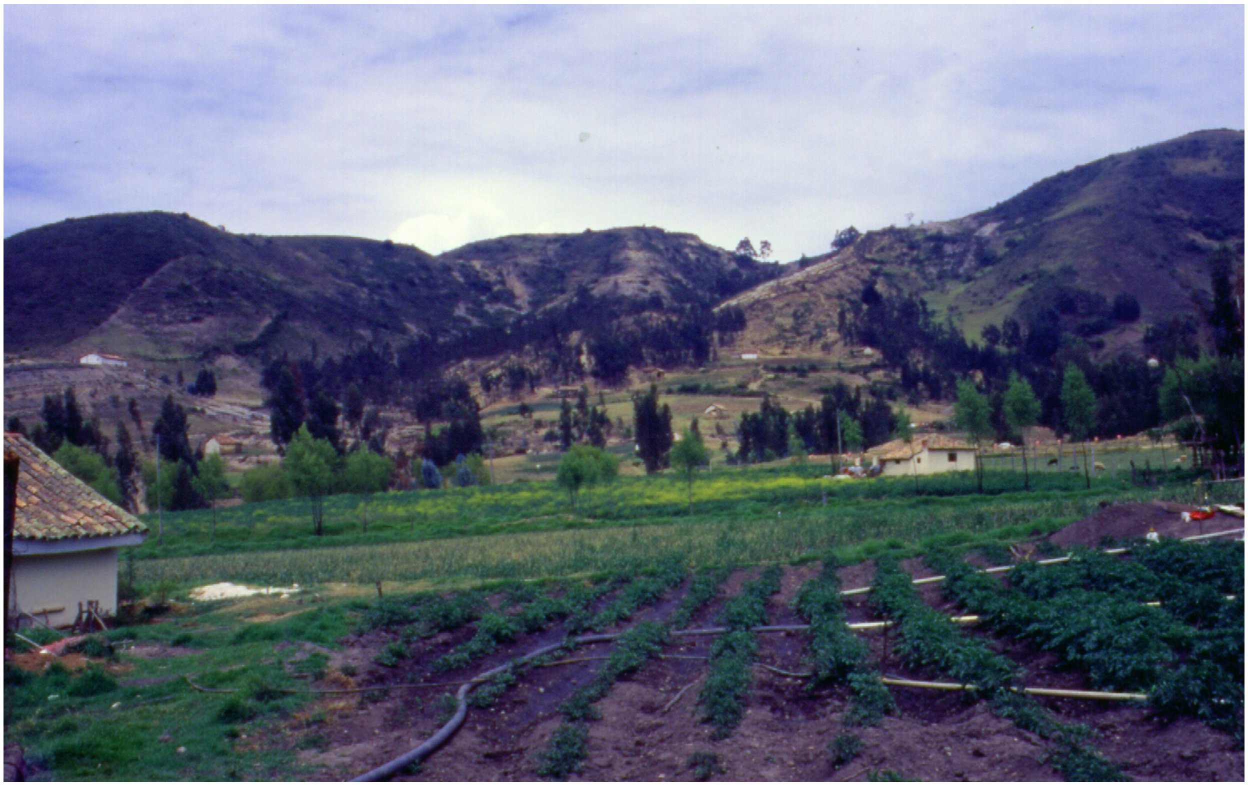

The excavations took place in 1995, and a systematic surface collection was conducted across the area. The surface collection retrieved cultural material from various periods of occupation in different neighborhoods or wards, separated by vacant spaces. Once the site layout was mapped, 65 test pits were dug across the neighborhoods. Some of these were expanded, and the total excavated area reached 190.5 $m^2$[45]. The site's chronology was derived from the proportions of each ceramic type at each level in each test pit. Ceramic material from the fill of each tomb, along with the grave goods, was also analyzed to determine the graves' chronology[45].

Multiple lines of evidence were used to identify sources of power and prestige. Ceramic decoration was interpreted as an indicator of wealth because it adds steps to the pottery production process[51], and acquiring or producing such fine ware is more costly. According to ref. [52], wealthy households tend to display finely made and decorated ware for serving during social gatherings. Therefore, high proportions of decorated ware were interpreted as representing wealthy households. Vessel shapes were used to identify areas where specific activities occurred more frequently to gain prestige. For instance, rituals require formal ware; based on ethnographic research, spoons, miniature vessels, incurved bowls, and cups were likely used for the consumption of hallucinogenic substances and other special purposes. Large quantities of serving ware, such as bowls and pitchers, were interpreted as indicators of locations of drinking parties and ceremonies, while a high proportion of cooking pots was interpreted as evidence of surplus food production. Imported pottery required greater effort to acquire, so it is viewed as a special commodity that requires access to exchange networks. Some imported prestige goods were used in ceremonial exchanges, such as pitchers with anthropomorphic representations, while others may have served as material links to ancestor lineages, such as the Herrera ware. For the analysis of the ceramic material, proportions for each category, residential unit, and period were used to calculate the mean proportion for the settlement as a whole. Then, each residential unit's category proportion was compared against the settlement proportion and the corresponding proportions of the other residential units[45]. Each proportion was then accompanied by three levels of confidence (80%, 95%, and 99%) to evaluate statistical significance, as shown in bullet graphs.

The analysis of faunal remains was conducted to identify differences in abundance, species diversity, meat utility, and meat weight among residential units in each period. This line of evidence served as a marker of control over basic resources. Areas with unusually high proportions of these categories were interpreted as indicating control over access to or consumption of meat. The analysis included the minimum number of individuals of each species or genus (MNI), the total number of identified specimens (NISP), meat weight in pounds based on known values for some species using MNI[53,54], and meat utility (MUI) based on ref. [55], with Caribou meat utility values divided into five ranks assigned to deer parts. The ranks ranged from very poor parts with very little meat, such as the skull, phalanges, or metacarpals, to very good parts with the meatiest portions, like the femur. More details on how the analyses were conducted can be found in ref. [45].

Spindle whorls were stone or ceramic weights suspended from a spindle that, when rotating, spun cotton or sisal fiber into yarn. Cotton was woven into textiles used for clothing, rituals, offerings, ceremonial exchanges, and tribute. Because of these diverse uses, textiles were exchanged for almost anything and served as a form of currency; they were therefore considered wealth. Whorls found at El Venado vary in shape within a similar weight range, and some authors say these differences were intended to produce yarn of different quality; tall, narrow whorls produce thin, tightly twisted yarn, while broader whorls produce thicker, less tightly twisted yarn[56,57] . A combination of different whorl types suggests the use of tool kits. Areas with large concentrations of spindle whorls and weaving tools were interpreted as specialized production sites.

For the analysis of funerary practices, we follow the assumption put forward by ref. [58] that a person's status in life within the status system is reflected in the treatment that person receives at death. Thus, the differences we observe in the funerary treatment of an archaeological population reflect the variety of statuses people held. At one end, the most basic burials involve little energy investment in grave construction, minimal body treatment, and no grave goods, while at the other end, more energy is invested in grave construction, body treatment, and the number of grave goods. All three pre-Hispanic periods provided small samples, in which sex and age were the main dimensions analyzed in relation to wealth, measured by the number, quality, and origin of grave goods, and prestige, measured by energy expenditure, such as grave depth, grave size, and furniture and body treatment.

The last line of evidence, dental health, was used to reconstruct dietary habits that typically reflect social status and wealth. Better dental health is expected among individuals with access to a more balanced, protein-rich diet. Three dental pathologies, caries, antemortem tooth loss, and periapical abscesses, were analyzed in relation to social status[45]. Analyses were conducted on the total number of teeth (for caries) and alveoli (for tooth loss and abscesses) per residential unit and ward. This method, known as tooth count, treats each tooth as a case and provides the number of teeth affected by each disease; these numbers were converted into proportions per residential unit. The comparison of mean proportions between residential units and the mean proportion of each category for the settlement was made using bullet graphs with various attached levels of confidence.

***Late Herrera period (800-1000 CE)***

The systematic surface collection during the Late Herrera period revealed that the settlement of El Venado had two distinct occupied areas, La Esmeralda and El Recuerdo. These were discrete groups of residential units, or wards, with unoccupied areas between them, all considered part of the settlement as a whole. Each ward functioned as a social unit, with members likely linked by kinship. Each ward appears to have operated through smaller economic units called clusters, composed of two to three residential units spatially organized around a small plaza (**Supplementary Figure 1.13**).

**Supplementary Figure 1.13**. **Distribution of wards, residential units, and tombs at El Venado during the Late Herrera period.** Map provided by Ana María Boada.

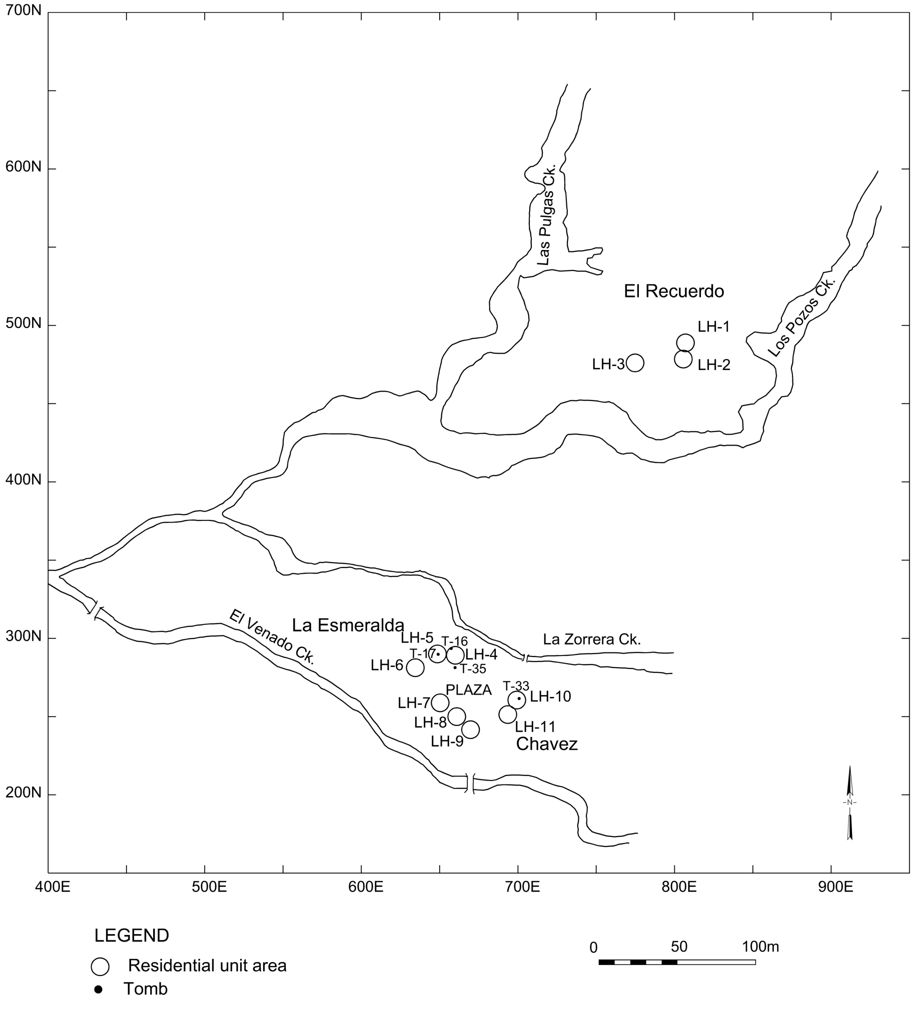

Most residential units performed redundant domestic tasks, but some specialized in a few complementary activities. These specialized activities were likely intended to address economic needs and social obligations. The surplus of food appears to have been intended for communal feasts and special ceremonies.

The archaeological assemblage of the El Recuerdo ward shows low overall proportions of decorated pottery, and no ceremonial artifacts were present. Residential unit (LH-1) contained more imported salt containers, a higher proportion of deer bones (Minimal Number of Individuals [MNI] and Number of Identified Species [NISP]), and a wider range of deer meat cuts, from very poor to very good. In contrast, residential unit LH-2 had a higher proportion of cooking pots and the poorest cuts of deer meat. LH-3 produced a higher proportion of bowls and a lower proportion of decorated pottery. No faunal remains were found in this residential unit. In general, this cluster shows increased effort on specific tasks: one household dedicated to hunting deer, another to acquiring salt likely for curing meat, and a third to cooking food and brewing corn beer. Evidence of other specialized activities, such as textile production, was also found in this ward. However, no graves were found in the El Recuerdo ward.

At La Esmeralda and Chávez wards, most domestic activities were similar across residential units, although specialized activities varied within the cluster. Residential unit LH-4 had a diverse assemblage but a lower proportion of decorated ware and a high proportion of Late Herrera pottery; LH-5 had higher proportions of cooking pots, imported pottery, and Late Herrera pottery, associated with exchange and domestic activities. This unit also contained all types of meat cuts, including the best cuts, which contrasts with the dental analysis showing that residents of LH-5 had poor dental health, with a high proportion of caries and antemortem tooth loss[45]. Greater access to meat would be expected to improve dental health, not the opposite. In contrast, LH-6 produced more pitchers, incurved bowls, and cups, an assemblage linked to ceremonial activities; it also has a high proportion of decorated pottery associated with wealth[45]. The cluster at the southern edge of the plaza includes residential units LH-7 and LH-8, which featured more decorated pottery, a marker of wealth, and more cooking pots. Meanwhile, LH-9 had more salt vessels, suggesting involvement in exchange and cooking activities. The cluster east of the plaza, with residential units LH-10 and LH-11, produced lower proportions of decorated pottery and a higher proportion of bowls, imported pottery, ancestral Herrera ceramics, and salt vessels. This cluster appears to have less wealth and was mainly involved in serving food, exchange, and, in general, more domestic activities. All the spindle whorls (n=3) from this period were found in LH-5, LH-6, and LH-8 at La Esmeralda, suggesting a concentration of yarn production and textile weaving in the ward (**Supplementary Figure 1.14**).

**Supplementary Figure 1.14. Sample of spindle whorls collected by the owner on his property in the La Esmeralda ward.** Map provided by Ana María Boada.

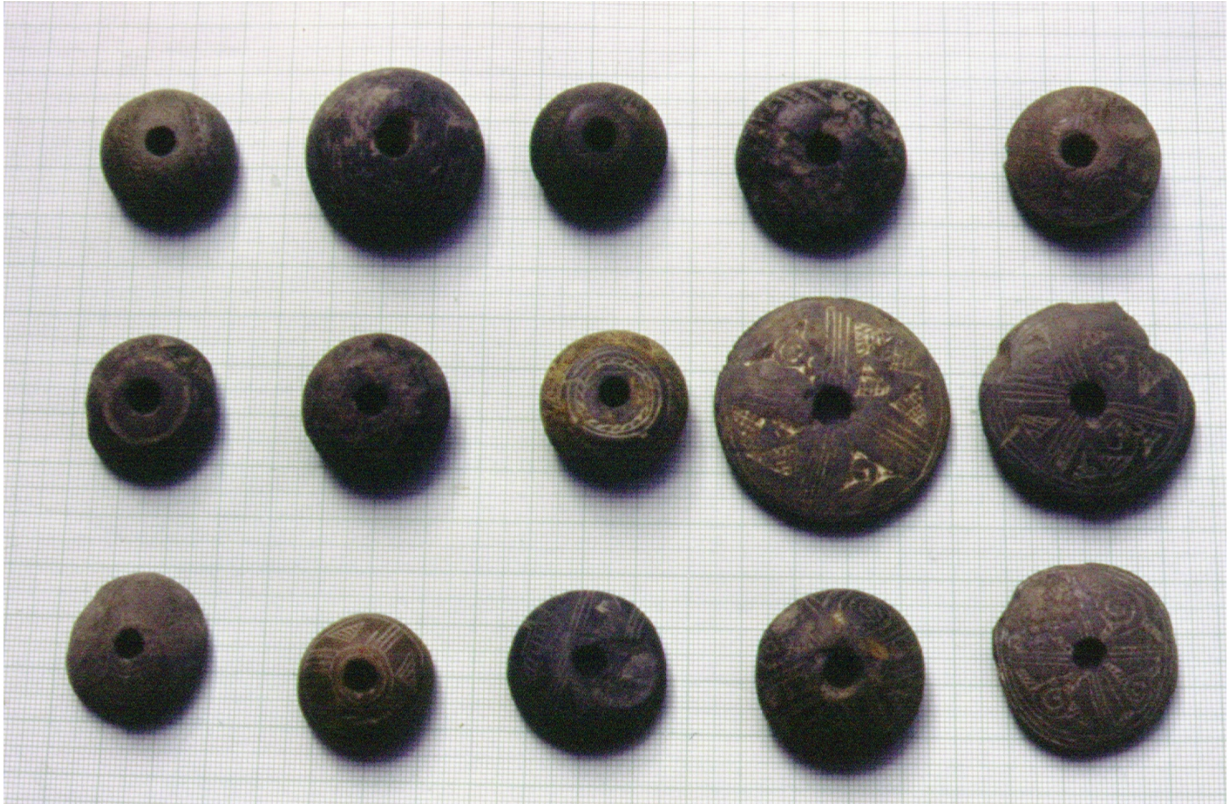

Mortuary practices for this period are based on 14 graves found in the La Esmeralda ward, while El Recuerdo remains unrepresented, although the owner discovered graves during agricultural work (**Supplementary Figure 1.13**). The sample of skeletons consisted of 10 adults and four infants. Of the six adults whose sex was determined, two were female and four were male. All bodies were buried in a fetal upright position in circular shafts, while others were placed in a lateral fetal position in oval shaft graves. Graves ranged in depth from 0.30 to 1.55 m. Some bodies received special treatment, with plaster applied over the entire body or partially covering the hips and feet. This body treatment was very similar to that described for the Marín site. The plaster was a mixture of plant ash and clay, sometimes sprinkled with red ocher. Although there is no evidence of placing a textile over the plaster to wrap the body, textiles were likely used to maintain the bundle, as described for the body treatment in Marín. Other grave furniture, such as stone slabs, well-crafted rectangular flat metates, and irregular rocks, were placed at the entrance of the grave. Most burials lacked grave goods, and only one had nine items. The most common item was pottery vessels (**Supplementary Figure 1.15**), some of which contained charcoal. Other grave goods included imported green stone, shell and gold beads, marine snail shells, agouti teeth, armadillo carapace, and metates. Some individuals' graves contained only imported grave goods, while others had locally made goods.

**Supplementary Figure 1.15. Ceramic vessel found at El Venado in Tomb-13.** Picture provided by Ana María Boada.

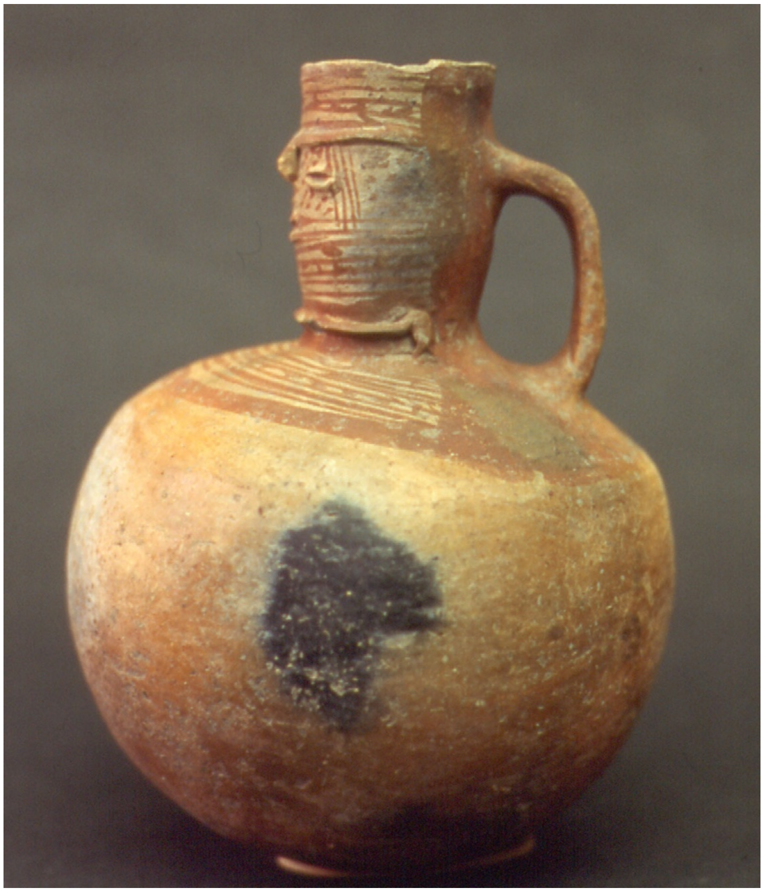

Differences in grave depth suggest differential energy investment: one adult male was buried 1.55 m deep, while others were buried 0.30 m deep, indicating a range of energy investment among adults. Children's graves also varied in depth, reinforcing social differentiation among children associated with ascribed status. Some adults' and children's bodies were wrapped in an ash mixture, but this treatment could not be correlated with other categories of energy investment. Differences in wealth were reflected in more grave goods and imported goods among adults, with females having fewer grave goods than males. However, this difference was not statistically significant ($t = -0.609$, $df = 4$, $p = 0.575$). A few males had the largest number of grave goods in the sample[45].

In sum, the evidence suggests that the La Esmeralda ward was the wealthiest and sponsored ceremonial exchanges, drinking parties, and rituals. It also contained the only evidence of textile production, indicating that this ward concentrated specialists who operated spinning kits for cotton yarn and textile production. Textile production was a specialized activity that likely served as a source of wealth. Although the sample is small, it provides evidence of social differentiation, with males having more wealth and prestige than women and some children having higher status by birthright. Dental analysis results are important because they show that the wealthiest household, with access to the best cuts of meat, may not have consumed them but instead gave them away during feasts and ceremonies.

***Context of the individuals sampled for genetic analysis from Late Herrera period El Venado***

Of a sample of 14 Late Herrera period burials, four individuals were selected for genetic analysis with three yielding genetic data with passing quality control metrics (we excluded Individual 33 (I22625) due to low coverage, **Table S1**). Descriptions of the burials follow.

**Individual 16 (I22471)** was excavated in the La Esmeralda ward, PP-072, residential unit LH-4. Genetic sex analysis identified the skeleton as male, and morphological analysis suggests he was in the 40–45 age cohort. The cranium showed no artificial modification. The tomb is a circular shaft 1.55 m deep, with 11 flat stones covering the surface at mid-depth. The body was in a sitting position, with no plaster cover. Grave goods included four ceramic vessels, comprising a pot, a bowl, a vessel fragment with charcoal, and a foreign cup. A rectangular, polished metate was located in the middle of the shaft fill.

**Individual 17 (I22473)** was found in the La Esmeralda ward, PP-056, household LH-5. Genetic analysis identified the skeleton as female, and morphological analysis suggests she was 45 to 49 years old. The cranium showed no artificial tabular deformation. The body was buried in a flexed sitting position in an oval pit measuring 0.93 by 0.64 meters and 1.10 meters deep. Grave goods included an armadillo carapace (*Dasypus* sp.) in the pit fill, six teeth of Agouti sp. near the body's knee, one spindle whorl in the thorax, and the bottom of a vessel at her feet.

**Individual 35 (I22472)**was found in PP-072 at La Esmeralda ward, residential unit LH-4. Genetic analysis determined that the skeleton was male, and morphological analysis indicates he was in the 30-34 years old age cohort. The cranium shows no artificial tabular deformation. The body was placed in an oval pit in a left-side fetal position. The head was oriented at 225° (clockwise from north). The skeletal remains were in poor condition. No grave goods were found in the grave.

***Early Muisca period (1000-1350 CE)***

During the Early Muisca period, the site expanded as the original wards, La Esmeralda and El Recuerdo, grew, and two new wards, Abejas to the north and San Antonio to the west, were added, with vacant zones separating them (**Supplementary Figure 1.16**).

**Supplementary Figure 1.16. Distribution of wards, residential units, and tombs for the Early Muisca period at El Venado.** Map provided by Ana María Boada.

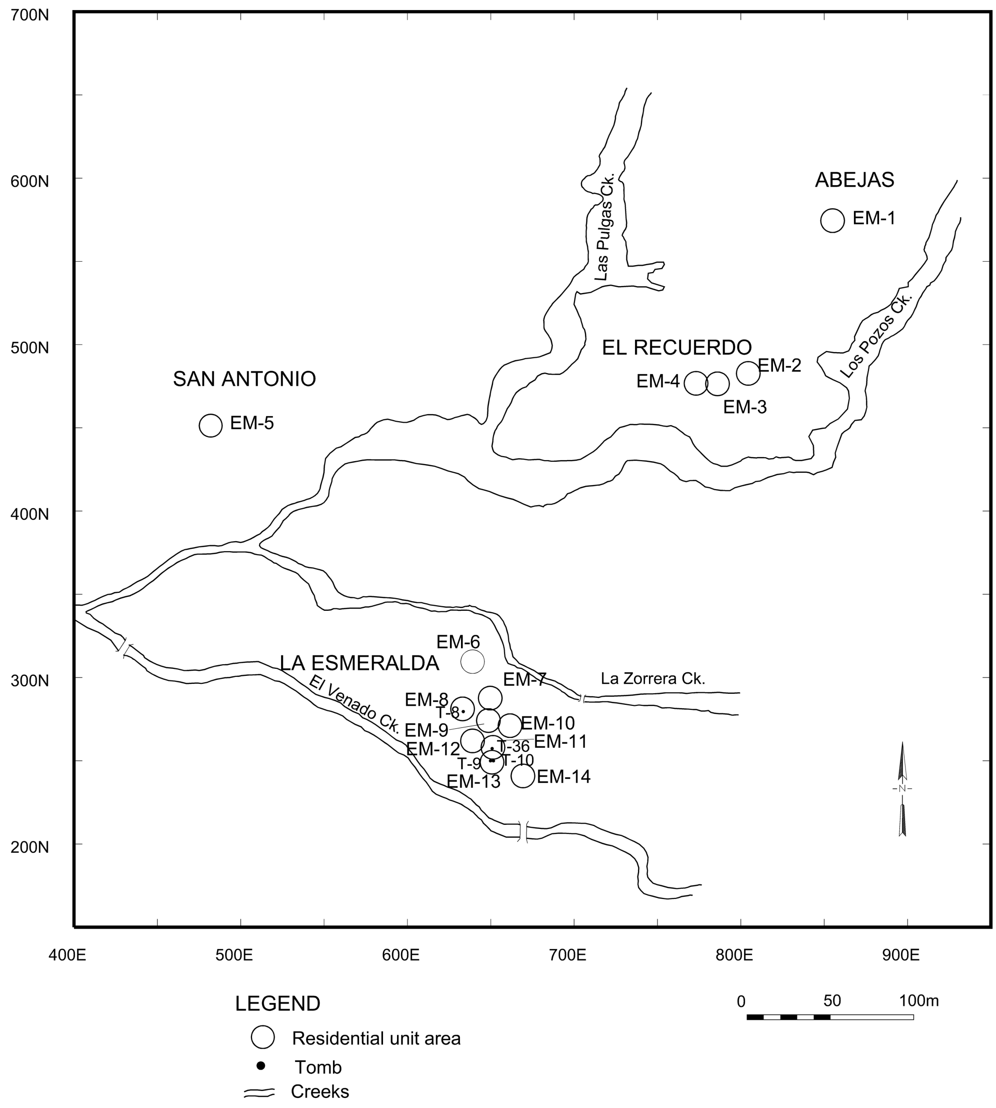


The wards no longer maintained the Late Herrera period's cluster-of-households pattern. Instead, the distribution of residential units appeared more random, with no clear central patio[45]. However, the wards continued to function as distinct social units. The Abejas ward began in this period with a single residential unit, EM-1, which had a higher proportion of cooking pots. Other pottery vessel shapes were at or below the overall settlement proportion. Overall, decorated pottery was scarce compared with other wards, and faunal analysis showed that cuts of deer meat in Abejas ranged from very poor to very good.

The Abejas ward began with one residential unit, EM-1, which featured a ceramic assemblage dominated by cooking pots and with low proportions of decorated pottery and imported goods. It also contained ceremonial objects, such as spoons, which are unusual outside the La Esmeralda ward. It also had a wide range of deer meat cuts, from very poor to very good. Regarding dental health, despite the presence of deer meat, this residential unit showed a high prevalence of caries.

This house did not have spindle whorls. EM-1 had the deepest grave, indicating a high level of energy investment in one burial, while the rest were not deep.

El Recuerdo ward is represented by residential units EM-2, EM-3, and EM-4, which showed low proportions of decorated ceramics but high proportions of cooking pots and imported ceramic salt vessels. These units produced 14.3% of spindle whorls, indicating cotton yarn and textile production. Faunal remains indicate that the residential units in this ward had poor-quality deer meat. No burials were found during the excavations in this ward.

La Esmeralda ward includes nine residential units: EM-6, which had a higher proportion of spoons and decorated pottery but no faunal remains. EM-7 yielded higher proportions of bowls, incurved bowls, and spoons; the only faunal remains were poor cuts of deer meat. EM-8 had high proportions of decorated pottery and deer cuts of meat ranging from very poor to very good; it also had high proportions of dental caries and premortem tooth loss. EM-9 exhibited high proportions of decorated pottery and bowls, but no faunal remains; EM-10 produced no high proportions of any particular ceramic shape and no faunal remains. EM-11 yielded a high proportion of decorated pottery and high proportions of very poor cuts of deer meat, with very low proportions of very good cuts of meat. EM-12 and EM-13 both yielded high proportions of pitchers and decorated ceramics, cuts of deer meat that ranged from very poor to fair, and a greater variety of animal species; yet they showed a high proportion of dental caries, premortem tooth loss, and periapical abscesses. From a total sample of five individuals excavated in El Venado, this residential unit yielded the burial (T-10) of a male with 6 grave goods, the richest of all graves for this period. Women from this ward exhibited more variation in labor investment in grave depth. The last three residential units had higher ratios of faunal bones to sherds than the overall settlement ratio, indicating greater concentrations of faunal remains. The spindle whorl analysis of seven pieces showed that 85.7% came from La Esmeralda ward, with four from EM-12 and two others in EM-8 and EM-9. Although the sample size of whorls is small, the concentration of these tools in La Esmeralda suggests that most cotton yarn production happened there. The different sizes and weights of the whorls also indicate they were used to make various qualities of yarn. Cotton yarn was used to weave textiles that were exchanged for a wide range of products. It is likely that the elite of La Esmeralda produced and/or controlled the textile manufacturing as a source of wealth.

The San Antonio ward is represented by residential unit EM-5, which contains a high proportion of decorated ceramics. Faunal remains were rare, and no spindle whorls or burials were found in this ward.

In sum, the ceramic assemblage from residential units provides evidence of social differentiation, with more decorated pottery at La Esmeralda and San Antonio suggesting these were the richest. Serving bowls and pitchers were more common in certain residential units in the La Esmeralda ward, where food and drink were served, suggesting that feasts and ceremonies took place there. Burials from La Esmeralda suggest that men had the richest graves, and, in general, there is more variation in social status than in the previous period.

Analysis of dental health in the Early Muisca period shows an increase in caries compared with the previous period, suggesting greater consumption of cariogenic carbohydrates. The sample is too small, and the pattern of residential units with better cuts of deer meat yet poor dental health will need to be tested with larger samples[45]. El Recuerdo and Abejas wards had residential units with higher proportions of cooking pots and salt vessels. This pattern suggests more intensive cooking, including salting meat, in these locations rather than in the elite residential units. However, lower proportions of decorated ware suggest less wealth in these wards. Given the distribution of cooking pots, serving bowls, and pitchers, it can be inferred that El Recuerdo provided food for the feasts and celebrations held at La Esmeralda. Abejas shows a slightly different pattern, suggesting that feasting may have also taken place in this ward, although on a smaller scale.

Funerary analysis for this period was based on a sample of 5 individuals, four from La Esmeralda and one from the Abejas ward. All were buried in single graves in a supine fetal position within circular or oval shafts. Individuals in this sample were adults who did not undergo elaborate body treatment. Women showed greater variation in energy investment, with grave depths ranging from .40 to 1.60 m, while the only man was buried in a .40 m deep pit. The differences in grave depth among women and the presence of slabs in one woman's grave suggest that some women had more labor invested in their graves than others, including the only man. These differences indicate social differentiation, with some individuals receiving more effort. This pattern is the opposite of what we saw in the previous period, where males received more labor investment, but this difference may be due to the small sample size. The total amount of grave goods was very small, with most having none and only a few having no more than one. The only man had more grave goods than most women. The absence of children in the sample prevents assessment of social differences among them.

Differences among residential units are associated with activities aimed at gaining status and maintaining power, such as specialized textile production to generate wealth and surplus food and beverage production that likely contributed to feasts and ceremonies. The La Esmeralda ward, with a higher proportion of jars used for serving chicha, spoons, cups, incurved bowls, and miniature vessels, was associated with control over esoteric knowledge, rituals, and ceremonies held there. Such control was essential to maintaining prestige. Higher proportions of decorated pottery and grave goods, along with a more varied range of imported products, indicate that the La Esmeralda ward was the wealthier ward of the settlement. Additionally, cotton yarn and textile production increased, adding to this ward's wealth during this period. Cooking tasks became more concentrated in a few houses of lower-status wards, such as Abejas and El Recuerdo. This increased production of food and beverages suggests surplus was produced to be taken to La Esmeralda for distribution at feasts and ceremonies. Faunal analysis also supports the idea that the La Esmeralda ward had prerogatives over access to deer meat and other species that could have been provided through tribute.

Although some ceramic types are characteristic of the Herrera period, especially the late phase, and differ from typical Muisca material, we generally observed continuity in the ceramic assemblage. This continuity is evident in types such as Fine Sandy ware, Coarse Sandy ware, Orange Burnished ware, and Gray Tempered ware. Shapes and decorative designs (painted and incised) evolved throughout the site's sequence, indicating cultural continuity. Lithic technology and assemblages, which include both polished and expedient production, persist from the beginning to the end of the sequence. Additionally, spindle whorl shapes and designs, along with mortuary practices, have maintained continuity since the Late Herrera period, indicating in situ evolution[45].

***Context of the individuals sampled for genetic analysis from Early Muisca period El Venado***

Of the five individuals from this period, four were selected for genetic analysis, including three from La Esmeralda ward and one from the Abejas ward (**Supplementary Figure 1.16**). The following description provides specific information about each individual in the Early Muisca sample:

**Individual 8B (I22476)** was found in PP-005 at La Esmeralda ward, residential unit EM-8. Burial 8 contained two skeletons: 8A was likely the original interment, removed to accommodate 8B, and then reburied. Genetic analysis indicates that 8B was female, and morphological analysis suggests she was 30-34 years old. The cranium showed no artificial modification. Individual 8B was buried in an oval pit 0.70 m deep. The body was placed in a left-side fetal position, with the head oriented to 120° (clockwise from north). A thin layer of red ochre was applied to the entire body. Overall, the skeleton was well preserved. No grave goods were included.

**Individual 10 (I22475)** was found in PP-006 at La Esmeralda ward, residential unit EM-13. Genetic analysis indicates the skeleton was male, and morphological analysis suggests he was 40-44 years old. The body was buried in an oval pit 0.40 m deep, in a fetal dorsal position with the head oriented at 125° (clockwise from north). No body treatment was evident, except that, to maintain this position, he must have been wrapped. The body was poorly preserved. A necklace with 6 beads—two bird-shaped stone beads, one discoidal seashell, one bell-shaped stone bead, one seashell canine-shaped bead, and one anthropomorphic bead—was included in the grave.

**Individual 30 (I22474)** was found in PP-015, at the Abejas ward, residential unit EM-1. Genetic analysis indicated that the skeleton was female, and morphological analysis placed it in the 25-29 age cohort. The body was buried in a circular pit 1.60 m deep, a depth far greater than the others. The body was placed in a fetal dorsal position, with the head oriented to 165° (clockwise from north), and no apparent body treatment was observed. Preservation of the skeleton was poor.

**Individual 36 (I22477)** was found in PP-051, La Esmeralda ward, residential unit EM-11. Genetic analysis determined that she was female, and morphological analysis suggests she was in the 25-29 age cohort. The body was buried in a circular pit 0.40 m deep, laid in a right-side fetal position, with the head oriented at 80° (clockwise from north). The body is relatively well preserved, with no observable body treatment. The cranium showed no artificial modification. A deer's femur near the head was included as a grave good.

***Late Muisca period (1350-1600 CE)***

Spanish ethnohistorical accounts describe the local chiefdoms of the Samacá Valley and neighboring valleys as small, with chiefs inheriting their positions but limited in their ability to accumulate wealth to support their political interests. Ethnohistoric documents state that the small chiefdoms of Monquirá, Saquencipá, and Sáchica originally inhabited the Samacá Valley. Just before the Spanish conquest, the chief of Tunja, commanding a large and complex chiefdom, along with his allies, set out to conquer the Samacá Valley, displacing the original chiefdoms to nearby valleys such as Sutamarchán, Sáchica, and Leiva, and installing some of his allies, including Cucaita, Sora, and Samacá, in the newly conquered territory[59].

During the Late Muisca period, the El Venado wards maintained their distinct distribution and identity; however, the distance between residential units increased compared to the previous period. The Abejas ward experienced extraordinary growth, reaching a size comparable to La Esmeralda, and the San Antonio ward also shows some growth. El Recuerdo, however, remained roughly the same size, and a new ward, El Rubí, appeared south of La Esmeralda (**Supplementary Figure 1.17**).

**Supplementary Figure 1.17. Distribution of wards, residential units, and tombs for the Late Muisca period at El Venado.** Map provided by Ana María Boada.

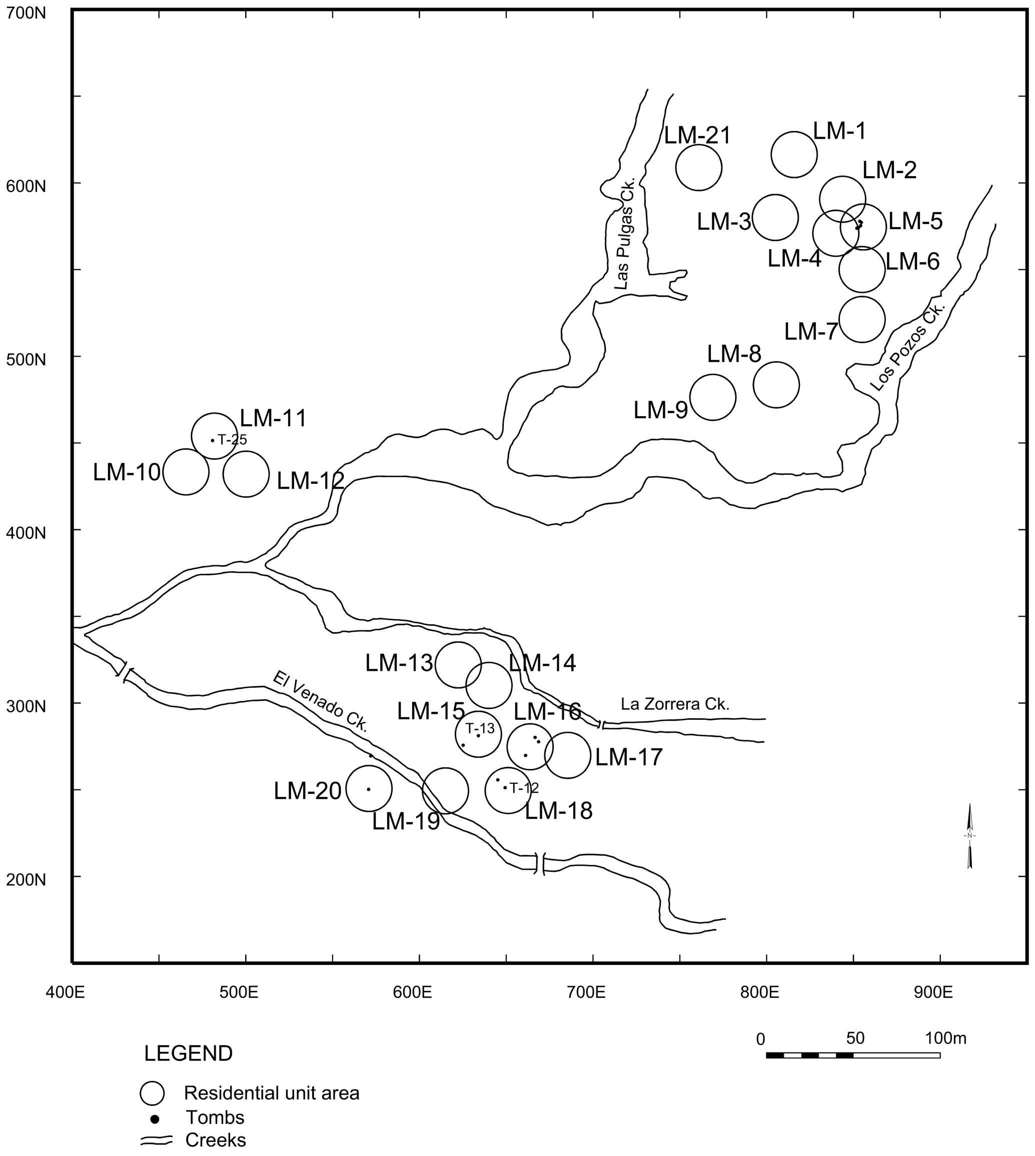

Redundancy in domestic activities, as evidenced by the ceramic and faunal assemblages, was present in most residential units. Some areas identified as residential units lacked the assemblages that characterize domestic areas and may have been part of a residential unit or simply discarded material in a public area. However, these areas were treated as residential units to better understand what they represented. Differences in the ceramic assemblages among residential units during the Late Muisca period suggest modest social differentiation.

The Abejas ward was represented by eight residential units, with much lower proportions of decorated ware than the settlement's mean. LM-1 showed a high proportion of pitchers, suggesting that feasting may also have occurred in this ward. LM-4 produced a large proportion of bowls for serving food and beverages. A small number of spindle whorls (12.5%) were found at Abejas, along with a textile weaving tool. LM-5 shows that almost all ceramic categories were present, including those associated with ceremonial activities. This residential unit produced the deepest burial, indicating more energy investment than the rest, and another tomb with more grave goods. The human remains sample from this residential unit shows the lowest proportion of caries in the settlement. LM-5 also had more variety of faunal genera in the ward and very poor cuts of deer meat. LM-6, LM-7, and LM-21 had vessel shapes with proportions similar to those of the settlement as a whole. No faunal remains were found in the other residential units.

El Recuerdo ward, with residential units LM-8 and LM-9, shows average proportions across all vessel shapes, and ceremonial vessels are absent. However, the two residential units produced higher proportions of imported salt vessels. This ward also had low proportions of decorated pottery. No faunal remains were found in this ward. However, this ward accounted for 12.5% of the spindle whorls.

In the San Antonio ward, residential unit LM-10 has a high proportion of very large, incurved bowls, or ollas-cuenco, and no other vessel shape appears in high proportion. LM-10 and LM-11 had a high proportion of decorated ceramics, though not as high as in La Esmeralda. This residential unit had only very poor cuts of deer meat, whereas the rest of the ward had none. Although no spinning or weaving tools for textile production were found, the ward did produce a bone needle.

The El Rubí ward had a greater variety of vessel shapes, but none of the shapes had particularly high proportions. It also had low proportions of decorated ware, and no evidence of textile production was found in this ward.

In La Esmeralda ward, the higher proportion of decorated pottery in residential units LM-15, LM-18, and LM-19 suggests these units were the wealthiest. However, other residential units in this ward showed a range of decorated pottery proportions, from low to high, indicating a graded distribution of wealth. Vessel analysis revealed differences in domestic and specialized tasks among residential units. At La Esmeralda ward, LM-15 showed a high proportion of pitchers associated with drinking parties and all vessel shapes used in ceremonies, suggesting involvement in rituals and ceremonial activities. This residential unit also had a broader range of proveniences for imported pottery, likely reflecting a wider network of exchange, although the number of imports is small. Notwithstanding LM-18’s high status, it had a high proportion of cooking pots, suggesting it was involved in more intensive cooking tasks, perhaps to provide food for feasts and ceremonies. LM-15 and LM-18 yielded the widest variety of animal genera in the settlement and the most abundant faunal remains. LM-18 had the best cuts of deer meat, but the dental analysis of the human remains showed a high proportion of caries. These results contradict expectations about the relationship between protein consumption and dental health. Of the eight spindle whorls found in the excavations and systematic surface collections, most (75%) were recovered at La Esmeralda, concentrated in the ward's center. The spindle whorls varied in shape and weight, likely producing a range of cotton yarn qualities. Additionally, bone needles were found in LM-18 and LM-15. Textile production appears to have occurred at these locations at a more specialized level. An unsystematic collection of spindle whorls with no chronology, collected by the owners of La Esmeralda, who gathered 94% (n=85) of the whorls in the settlement, supports this pattern.

The Abejas, El Recuerdo, and El Rubí wards had low proportions of decorated ware, suggesting they were the least wealthy in the settlement. In contrast, La Esmeralda ward shows greater wealth, a pattern that extends back to earlier periods.

Among a sample of 15 burials, 10 were adults (4 females and 2 males), and 5 were infants[45]. Infants were buried in shallower graves than adults, but adult graves showed wide variability in depth. The grave of one adult was 0.32 m deep, while that of another was about 1.5 m deep. Children and adults had similar numbers of stone slabs in the tomb and comparable distributions and quantities of grave goods. Within each age group, one individual had more grave goods than the others. A larger number of grave goods associated with an infant is interpreted here as a sign of ascribed status. Females had deeper tombs, more grave goods, and more slabs than males, although these differences were not statistically significant. Females also showed greater variability in the number of grave goods, ranging from none to nine. Notably, graves with greater energy investment (deeper shafts) and greater wealth in the number of grave goods were located in the Abejas ward, not in La Esmeralda. Female burials had more grave goods than male burials, reversing the pattern of previous periods, which may reflect females gaining social importance through increased participation in the economy and the production of household wealth. However, male burials showed greater depth variability than female burials, suggesting higher prestige.

***Context of the individuals sampled for genetic analysis from Late Muisca period El Venado***

Of the 15 individuals from this period identified at the El Venado settlement, two were selected for genetic analysis. Both skeletons are from the La Esmeralda ward (**Supplementary Figure 1.15**).

**Individual 12 (I22480)** was found in PP-006, La Esmeralda ward, residential unit LM-18. Genetic analysis identified the skeleton as female. Morphological analysis suggests she was 35 to 39 years old. The body was buried in a grave niche with a circular shaft 0.68 m deep, in a right-side fetal position, with the head oriented 160° (clockwise from north). No evidence of body treatment was observed, and no grave goods were present.

**Individual 13 (I22478)** was found in PP-005, La Esmeralda ward, residential unit LM-15. Genetic analysis determined that the skeleton was female, and morphological examination suggests she was in the 30-35 age cohort. The body was buried in an oval pit 0.80 m deep, in a left-side fetal position, with the head oriented at 230° (clockwise from north). The body was covered with a layer of brown, soft soil resembling a partial wrapping, similar to that used in Marin, but it was not composed of ashes. The skeleton is well-preserved, and there is no artificial cranial modification. A pitcher with an anthropomorphic design on the neck (**Supplementary Figure 1.15**) was found at her feet, along with two lithic flakes with no signs of use near the head, and a necklace on the thorax featuring five discoidal black stone beads with carved designs, 12 black tubular glass beads, a black glass barrel-shaped bead, six rectangular glass beads, and four green glass discoidal beads, all of European origin. Additionally, 46 shell beads, each 5 mm in diameter and 1 mm thick, were found.

***Conclusions***

The archaeological assemblage shows variation in wealth and activities across the wards. Some wards, such as La Esmeralda, appear wealthier than others throughout the sequence. Residential units within each ward also reveal differences in wealth and status, suggesting that even in the wealthiest ward, some units had less wealth, indicating a range of wealth and prestige among the elite. The same pattern is found in residential units of the lower-status wards. There are no sharp distinctions between poor and rich residential units; instead, there is a gradual differentiation of wealth within and among wards.

The evidence shows that elites across different periods of occupation did not rely on a single source of power—economic, political, social, or religious—to establish social hierarchy. Instead, they used a mix of strategies to gain and maintain prestige and wealth that evolved over time. During the Late Herrera period, inherited succession appears to have been a primary source of power. Ascribed status is evident in the archaeological record through infants buried with more grave goods than many adults. Additionally, early Herrera ceramics, likely brought by the early settlers of El Venado, are concentrated in the La Esmeralda ward. This pottery may have originated in Tunja, the regional center, where such ceramics were common during the earliest occupation[60]. The Herrera pottery in La Esmeralda is utilitarian rather than highly decorative, suggesting its value lies more in its ancestral significance than in its appearance; it was likely used to legitimize the lineage of the founding families. The La Esmeralda ward sponsored and performed rituals and ceremonies in the settlement of El Venado throughout the cultural sequence. Thus, power relied heavily on ideological strategies based on genealogy and the performance of rituals, ceremonial exchanges, and drinking parties.

Inherited status, along with the privileges of high-ranking families, is also reflected in their wealth, as evidenced by higher proportions of decorated pottery and more finely crafted imports from a broader range of sources, indicating a larger exchange network. Best cuts of deer meat, greater faunal diversity, and more abundant identified faunal bone genera suggest privileged access to meat among the inhabitants of the La Esmeralda ward, although not necessarily for their consumption. Specialized textile production at La Esmeralda further suggests that the elites used textiles as a source of wealth.

Over time, the importance of ancestral pottery to the founding lineage waned, and elites sought new ways to uphold their status and power during the Early Muisca period. Inherited status continued to limit who could reach the top of the social hierarchy and become a chief. Once in office, chiefs employed various strategies to maintain power and prestige. The La Esmeralda ward remained the center for special ceramics used in hosting celebrations, rituals, and ceremonial exchanges. This ward also retained prerogatives over access to basic resources, such as better cuts of deer meat. More importantly, it remained the wealthiest, as evidenced by its high proportion of decorated ware, the specialized production of cotton yarn and textiles, and the importation of more goods, all of which contributed to the accumulation of wealth to support chiefly enterprises and maintain their high status and power.

During the Late Muisca period, La Esmeralda was not the only ward to host celebrations; the Abejas ward also held them, challenging La Esmeralda's prerogative. The initial ideological strategies for acquiring and maintaining prestige and power seem to have lost their significance. Although sponsoring drinking parties in Abejas may have been a challenge for La Esmeralda's elite, they continued to host ceremonies and rituals featuring games that gave young men opportunities to display their aptitudes, gain prestige, and rise in the social hierarchy. Inherited rights remained a powerful ideological strategy for maintaining power during this period. The elite from La Esmeralda ward also retained prerogatives over economic resources, such as the best cuts of deer meat and rights to hunting grounds, as described in ethnohistorical accounts. They continued to display their wealth through decorated ware, which became more restricted, and to maintain specialized production of cotton yarn and textiles, a critical source of wealth. Ethnohistoric accounts also describe the payment of tribute to the chief. Archaeologically, the distribution of faunal remains could be evidence of tribute payments in hunted species. Other goods, produce, arms, exchanged goods, and raw materials were also used as tribute. People also paid tribute in labor by cultivating the chief's lands every year and building his houses and temples. At this point in the cultural sequence, tribute and wealth became the primary sources of economic support for maintaining the chief's power.

Traditional interpretations attributed changes in ceramic shapes and decoration to new people colonizing the area[61]. However, despite these changes, ceramic vessels, spindle whorls, and lithic tools show continuity in shape, manufacture, and decoration. This continuity also extends to the settlement pattern, grave shapes, and mortuary traditions (including body treatment, positioning and orientation, and grave goods), as well as to bodies interred within the settlement's living areas throughout the three periods of occupation. Taken together, this evidence indicates cultural continuity throughout the three periods. The observed cultural continuity is now supported by the genetic analysis, which also shows biological continuity between the Herrera and Muisca period populations.

**La Salina (Cundinamarca, Nemocón)**

The site of La Salina is located in the municipality of Nemocón, within the department of Cundinamarca. Analysis of materials recovered as part of archaeological excavations at the site were included in the publication of the work at the Zipaquirá saltworks[62]. A radiocarbon date yielded a date of 2350±90 calBP (400 ± 90 BCE) [63], originally reported in ref. [62]. Subsequent research documented evidence of diverse human activities dating back to the Herrera period that may be correlated with the social organization of salt production in the area surrounding the salt mine[64].

Detailed information about La Salina as a salt production site in the Checua River valley can be found in ref. [65]. We attempted to generate ancient DNA data from three individuals who were buried in a flexed position and successfully obtained data from one individual.

**Las Delicias (Cundinamarca, Bogotá)**

The Las Delicias site (LD) (4° 35´ 54.22” N, 74° 8´58.32” W) is located southwest the city of Bogotá near the Tunjuelito River and represents both a residential and burial site. Two radiocarbon dates of 1180 ± 70 (Beta-39874 wood charcoal) and 1010 ± 60 B.P (Beta-39873 wood charcoal) (Enciso, 1993) suggest occupations during the early Muisca period (some radiocarbon dates obtained from human bone using beta-counting techniques were corrected for isotopic fractionation using measured $\delta^{13}C$ values and calibrated with CALIB[66] 7.0 v. 7 based on the computations of ref. [67]. The ratio $^{14}C/^{12}C$ and a mean human value of $\delta^{13}C$ derived from each sample were used to perform the correction). Nevertheless, the author on the basis of the finding of the late Chocontá vidriada pottery type suggests a continuous occupation for several centuries[68]. Eighteen individuals of both sexes and different ages were recovered apparently related to the earliest occupation[69]. Abundant animal and plant remains were recovered that represent deer, guinea pig, mollusks, maize, beans, cotton and potato[70,71].

**Marín (Boyacá,Valle de Samacá) - Ana María Boada**

The archaeological site of Marín is located in the municipality of Cucaita, Boyacá Department, with geographic coordinates 5.538671 N, -73.483820 E, at an altitude of 2600 m above sea level, in the eastern branch of the Colombian Andes (**Supplementary Figure 1.11**). The settlement occupied the smooth, eroded northern slope of an isolated hill. The climate is cool, with average temperatures between 12 and 18°C, though some nights can drop below freezing, especially in January and occasionally in June. The climate supports a low dry forest, mainly on the mountains surrounding the valley. Most of the valley is currently used for the cultivation of grains, onions, and tubers.

Marín was part of a settlement system that included several other contemporary settlements from this period. An unsystematic survey of the Cucaita and Sora valleys revealed two or three contemporaneous settlements of similar size[72-74]. Ethnohistorical information indicates that the valley was conquered by the chief of Tunja a few years before the Spanish conquest. The Tunja chief displaced most of the independent small chiefdoms living in the Samacá Valley to the nearby Leiva and Sáchica valleys, leaving the land for the newcomers to settle in the northern part of the Samacá Valley[59,75].

The settlement of Marín, also known as El Santuario, was a community with a ceramic assemblage marking the transition from the late Early Muisca to the early Late Muisca period (AD 1180-1480 Cal), although radiocarbon dates extend the range at both ends. A wood charcoal sample from a cup in Tomb 20 yielded a conventional radiocarbon age of 700±80 BP (Beta-22667, uncalibrated). Calibration with OxCal[25] v.4.4.4 using the IntCal20[26] Northern Hemisphere curve (produced two ranges: 1177 (1.6%) to 1193 calAD and 1202 (93.9%) to 1414 calAD. A second wood charcoal sample from Tomb 10, taken from the body-ash wrap (60 cm deep), yielded a radiocarbon age of 600±100

BP (Beta-22668, uncalibrated). Calibration with OxCal[25] v.4.4.4 using the IntCal20[26] Northern Hemisphere curveproduced a range of 1220–1481 calAD with 95.4% probability.

The site was first excavated in 1986, with several additional excavations through 1992[72,75-77]. Multiple artificial terraces were built on the hillside by cutting into the surface to create flat areas, but they were not reinforced with structural features to maintain them (**Supplementary Figure 1.18**).

**Supplementary Figure 1.18**. Terrace 3 on the northern side of the Marín hill. Photo provided by Ana María Boada.

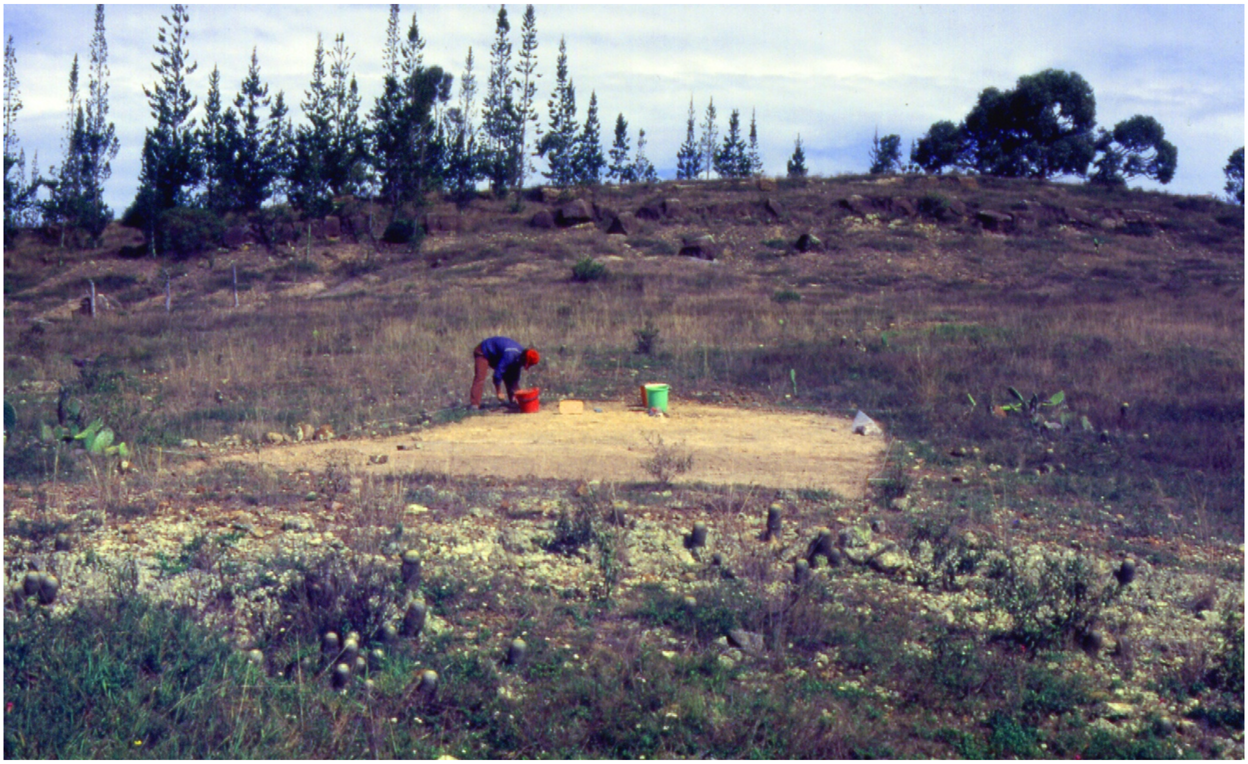

Excavations revealed the remains of structures, including post molds for round huts and a partial post alignment. We excavated about 650 square meters and recovered information on houses, pits, hearths, post alignments, and most of 70 burials[76,78]. Although systematic sampling was not performed, I believe we covered several areas, which could be considered a stratified sample. The high degree of erosion on the hill allowed excavations to proceed quickly. Unfortunately, in most areas, the cultural floor was nonexistent, and a great deal of information had been lost[19].

Marín shows very modest economic differences among households in fine pottery, exotic objects, goldwork, funerary treatments of the dead, and grave goods. Most houses were built on artificially flattened terraces of varying sizes. Terrace 3, located in the center of the settlement, is the largest flattened area. On this terrace, a circular house about 7.2 meters in diameter was found, built with posts 0.25 m in diameter, placed every meter, and a large central post about 0.45 m in diameter (**Supplementary Figure 1.19**). The walls were likely made of small bamboo, possibly covered with mud, to protect against the cool wind. Nearby, there was an east-west oriented alignment of 16 posts that might have been part of a palisade surrounding the house or a structure serving another purpose. This structure was the largest circular house in the settlement and seems to have been the community's elite residence. Interestingly, most women, men, and children in this large house displayed artificial oblique tabular cranial modification (**Supplementary Figure 1.20**), distinguishing this group from the rest of the population, which did not exhibit cranial modification[78,79]. The other houses excavated in the settlement were also built on artificial terraces; they were small, circular, thatched structures. Subsurface features such as earth ovens, pits, and graves were found both inside and outside the houses[77,80].

**Supplementary Figure 1.19**. **Distribution of features and burials in Terrace 3 at Marín Site**. The burials analyzed for ancient DNA were excavated from this context and are described below. Drawing provided by Ana María Boada.

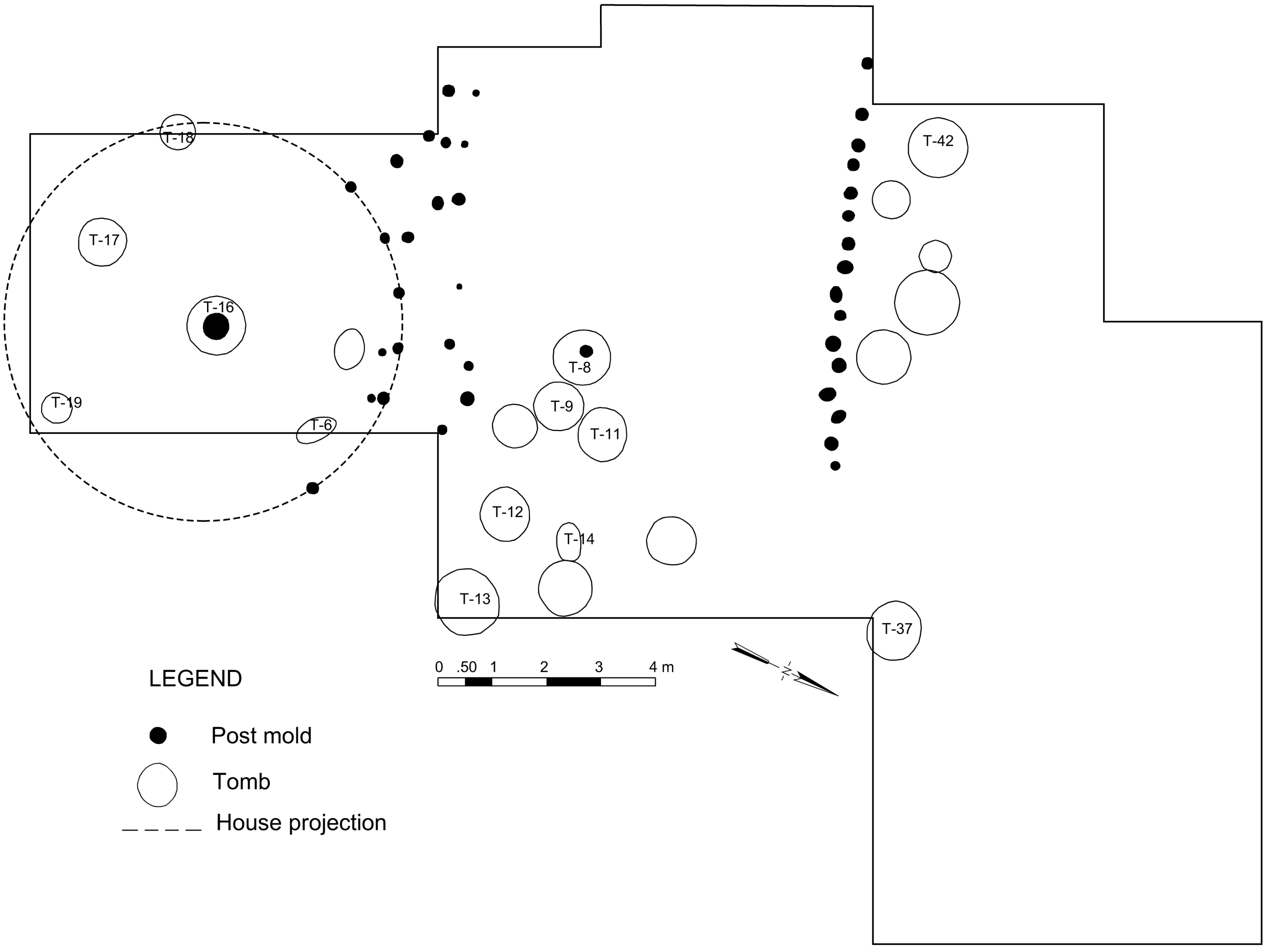

**Supplementary Figure 1.20. Skull with artificial oblique tabular cranial modification from Marín.** Photo provided by Ana María Boada.

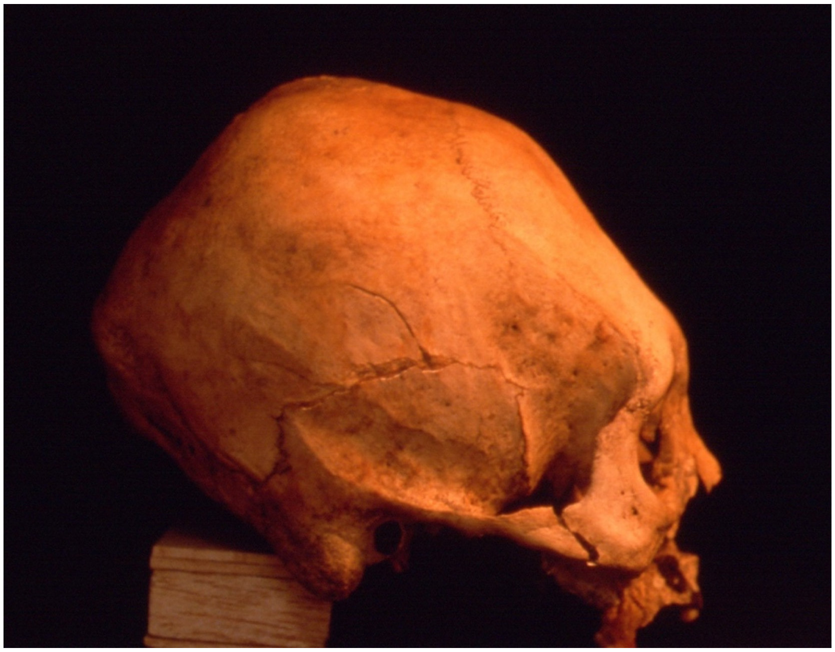

Graves took various forms, including oval and circular shafts, as well as shafts with a small niche containing human bodies in a horizontal fetal or sitting position. Some corpses were wrapped in textiles and covered with a plaster made from a mixture of ashes from burnt plants and clay. The plaster was then covered with textiles and secured with cords. Some bodies had only partial ash mix wrapping around the feet and hips, while others were wrapped completely (**Supplementary Figure 1.21**).

**Supplementary Figure 1.21. Illustration of Tomb 17, a shaft with a lateral niche containing a wrapped body at the Marín site.** Photo provided by Ana María Boada.

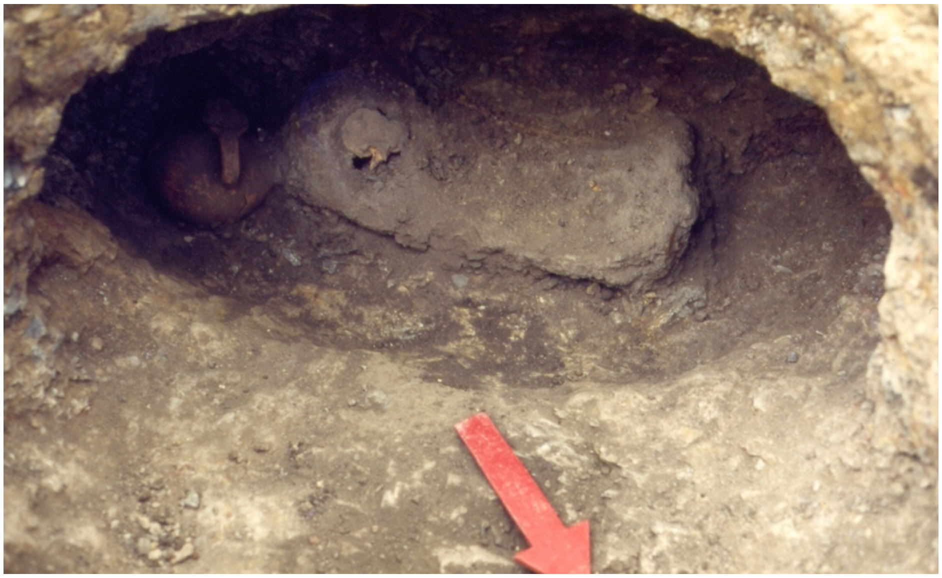

Social stratification is evident in some children being buried with more grave goods and in deeper tombs than most adults, indicating ascribed social differences at birth. Similarly, artificial cranial modification in children points to social differences assigned at birth. Additionally, a very modest level of economic differentiation is indicated by a few more objects and exotic artifacts in some graves (**Supplementary Figure 1.22**).

**Supplementary Figure 1.22. Photo of grave goods from Tomb 27 at the Marín site.** Photo provided by Ana María Boada.

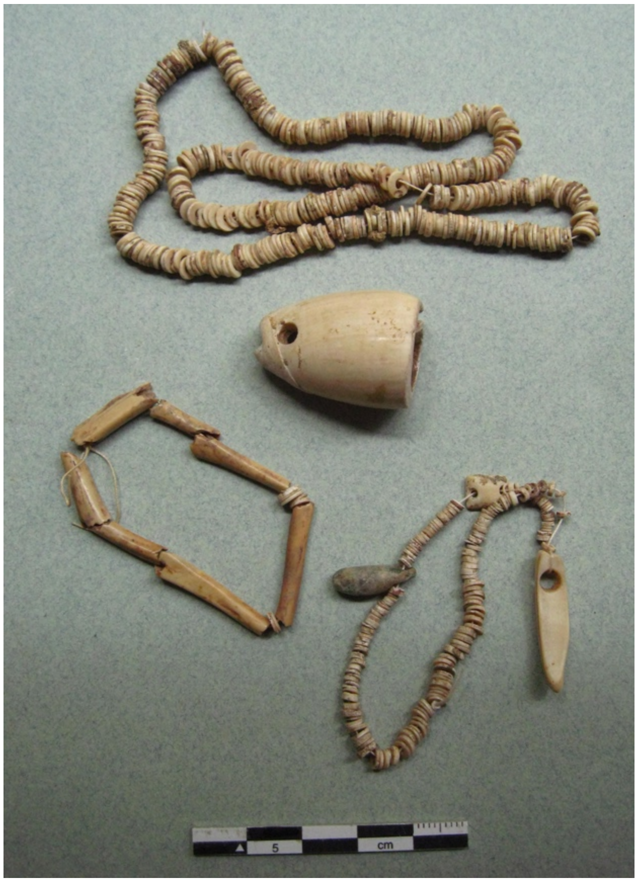


Given the group's higher status, its central location on Terrace 3 within the settlement, and its occupancy of the largest house, several questions arose regarding relationships among household members. A sample of skeletons from this household was collected to analyze their DNA with an aim to determine their kinship ties.

##### The individuals sampled from Marín

Of the 70 individuals detected at the site, 12 were selected for genetic analysis and all yielded ancient DNA data that passed quality control checks. All individuals are from the same area and are likely from the same household that occupied the most prominent house structure on the site, with posts aligned on the north side (**Supplementary Figure 1.19**). Most of them show cranial modification. Sex determination of the human remains was made using standard criteria for cranial, pelvic, and long bones morphology[81-83]. For sex determinations, each observed feature was graded on a scale of -2 to 2, according to the format developed by ref. [84]. In their system, positive values indicate masculinity, while negative values indicate femininity; individuals with a total score between -0.5 and 0.5 were considered of indeterminate sex.

Several criteria were also combined for age determination, including pubic symphysis morphology[81], auricular surface morphology[85], dental eruption[86,87], degree of dental attrition[88-90], and ectocranial suture closure[91]. Each criterion was used to determine an average age, which was then placed in an age cohort. These guidelines were followed for the samples of Marín and El Venado. Descriptions of the burials follow.

**Individual 6 (I22803)** was found in Terrace 3, within the large house. The individual's head was oriented at 120°. The person was a male (based on genetic sex), five years old, and was lying in a shallow, oval-shaped grave in a flexed position, with the entire body wrapped in a mixture of clay and ashes. No grave goods were present. The cranium showed no distinguishable artificial modification. Genetic analysis indicates that he is unique, as he has no relatives in the settlement.

**Individual 8 (I22492)** was found in Terrace 3, between the large house and the palisade. The individual is a female (based on genetic analysis) and falls within the 40-45 age group. The skeleton is well-preserved, and the cranium exhibits artificial oblique tabular modification. A post mold was found in the grave but did not contact the skeleton. The body was found in a sitting position, entirely wrapped in a mixture of ash and clay that shows textile imprints. The grave contained no goods. This female was the daughter of Individual 42 (I22491).

**Individual 9 (I22494)** was found in Terrace 3, between the large house and the palisade, in a grave niche with a shaft. This person was a male (based on genetic analysis), aged 25-29. The skull shows artificial tabular modification. He was buried seated, fully wrapped in a plaster mixture of clay and plant ashes and covered with a textile. A small animal bone was found at the feet of the skeleton as a grave good.

**Individual 11 (I22495)** was located in Terrace 3, between the large house and the row of posts. This person was buried in a circular shaft grave. He was male (based on genetic analysis), in the age range of 30-34 years old. The cranium shows signs of artificial tabular modification. The body was buried seated, wrapped in a mixture of ash and clay, and then covered with cotton textiles. Two animal bones were buried with the body.

**Individual 12 (I22805)** was found in Terrace 3, between the large house and the post line. Genetic analysis indicates the individual is female, and approximately 2 years old. The cranium shows artificial tabular modification. The child was buried in a fetal position, wrapped in a clay-and-ash mixture and covered with textiles secured with cords. She was placed in a circular shaft 1.10 meters deep, seated on a metate fragment. The grave goods included two discoidal shell beads.

**Individual 13 (I22801)** was located in Terrace 3, between the large house and the line of posts. Genetic analysis indicates the individual is female, and 25-29 years old. The cranium shows artificial tabular modification. She was buried in a deep grave, covered with multiple irregular stones, with a small niche containing a ceramic fragment on which the body was seated. The grave contained several grave goods: a half-bowl filled with charcoal, a shell bead, and a lithic scraper at her feet. The body was wrapped in a mixture of ash and clay, then covered with textiles. Over her head, there was a layer of clay with imprints of a mesh.

**Individual 14 (I22804)** was found in Terrace 3, between the large house and the line of posts. Genetic analysis determined that the individual is female and that she is estimated to be between 35 and 39 years old. The cranium shows no signs of artificial modification. She was buried in a dorsal fetal position in an oval, shallow grave. This body shows no evidence of ash wrapping but may have been wrapped in a textile, as a flat clay piece bearing a fingerprint and a textile imprint was found around her neck. A fragmented bowl was buried alongside her.

**Individual 16 (I22497)** was found in Terrace 3, at the center of the large house. Above this burial was a large post mold, but the post did not disturb the skeleton. The cranium shows artificial oblique tabular modification. This individual was a male (based on genetic analysis) in the 40-44 age cohort. He was buried in a round shaft 1.70 m deep, in a fetal upright position, wrapped in a mixture of ash and clay; he was presumably wrapped in textiles. He was buried without any goods. His burial at the center of the house suggests high status and possibly his role as a lineage founder.

**Individual 17 (I22496)** was found in Terrace 3, within the large house. Genetic analysis indicates he was male and approximately 4 years old based on dental eruption patterns. The cranium shows signs of artificial oblique tabular modification. The body was buried in a shaft with a small niche grave, placed in a left-side fetal position. It was wrapped in a layer of clay, covered by a textile that left an imprint in the clay. The body was then wrapped in a mixture of ash and clay and covered again with a textile. This child was buried with a pitcher, a necklace made of four teeth from *Tayassu pecari*, a mandible from *Didelphis marsupialis*, a bone bead, 14 discoidal shell beads, and a fragment of an ulna.

**Individual 18 (I22493)** was found in Terrace 3, inside the large house. Genetic analysis indicates the body was male and 3 years old based on dental eruption patterns. The cranium shows artificial oblique tabular modification. The body was buried in a seated position within a large cooking pot placed in the grave's niche at a depth of 78 cm. The pot was covered with a thin layer of dark clay, similar to that used for other wrapped bodies. Three animal bones, not yet identified, were found inside the pot as grave goods.

**Individual 19 (I22802)** was found in Terrace 3, within the large house. Genetic analysis indicated that the individual was female. Based on dental eruption patterns she was 2 years old. The cranium shows oblique, tabular artificial modification. The body was buried in a large cooking pot, then placed over a flat rock in a grave with a circular shaft 0.50 m deep. No grave goods were present.

**Individual 42 (I22491)** was found in Terrace 3 on the north side of the post line. Genetic analysis identified the individual as female, and morphological analysis estimated an age of 35-39 years. The cranium shows artificial tabular deformation. The body was buried in a shaft with a niche grave, 1.07 meters deep, in a right-side fetal position. The head was oriented at 40°. The body was partially wrapped in ash and clay, covering the hips and feet, and a small, gray, bell-shaped bead served as a grave good. This female was the mother of female 8. Information about the 11-member family from Terrace 3 at Marín is in **Supplementary Information 5**.

**Nueva Esperanza (Cundinamarca, Bogotá) – Pedro Argüello**

The Nueva Esperanza (NE) site lies on a natural terrace of approximately 20 hectares at 2,570 meters above sea level at coordinates E977235 N977318 (source: Magna Sirgas Colombia - Bogotá). It was excavated as part of a rescue archaeology contract and is located on the right bank of the Bogotá River, near the Tequendama Falls and just 7 km from the municipality of Soacha, an ancient Muisca indigenous settlement that was reconfigured during the Spanish colonial period in the 16th century. Soacha and NE are part of a large and diverse territory called Sabana de Bogotá (SB), which covers an area of approximately 2,500 $km^2$ where biodiversity allowed the development of multiple human groups with particular historical trajectories, but which shared some cultural traits that were strengthened over time.

For more than 2,000 years, the Nueva Esperanza site was inhabited by indigenous Herrera and Muisca families, who slowly turned the place into one of the most important settlements in the region. It was here that various forms of political, economic, and ideological expression took place, changing over time and recorded archaeologically as the Herrera period (400 BC–200 AD) Early Muisca (200 AD – 1000 AD) and Late Muisca (1000 AD – 1600 AD).

Archaeological evidence indicates that the first inhabitants of NE formed families that in many cases extended their lineages until the Late Muisca period, building circular and rectangular dwellings, which, in addition to being used for living and carrying out domestic and economic activities, were also spaces for burying and worshipping their dead. Agriculture and textile production were apparently the two economic pillars of this population, which managed to consolidate trade routes with neighboring populations within the Muisca area and in more distant territories, covering distances of up to approximately 950 kilometers in what is now the Sierra Nevada de Santa Marta, occupied by the Taironas.

The inhabitants of Nueva Esperanza expressed cultural phenomena such as social inequality through their funeral practices. In tombs associated with the Herrera period, material indicators associated with power and political prestige have been found, which were not displayed by all inhabitants and allow us to infer differences in access to resources, which could have been controlled by social groups with some type of kinship, since tombs from this period were characterized by collective burial.

During the Early Muisca period, burial practices were transformed in response to political and ideological changes, whereby prestige could be achieved through individual actions. However, these processes took time and were slowly consolidated over a period of approximately 800 years. Around the year 1000 AD, new cultural changes were recorded, which were reflected in burial patterns, where not only was individuality privileged, but changes were also identified in the transmission of power or prestige, which was possibly obtained through bloodline regardless of personal merit.

*El sitio Nueva Esperanza (NE) yace sobre una terraza natural de aproximadamente 20ha a 2570 msnm en las coordenadas E977235 N977318 (origen Magna Sirgas Colombia - Bogotá), excavado en el marco de la arqueología por contrato de rescate y se encuentra localizado en la margen derecha del Rio Bogotá, en cercanías del Salto del Tequendama y a escasos 7 km del municipio de Soacha, un antiguo poblado indígena muisca reconfigurado durante la colonia española en el siglo XVI. Soacha y NE hacen parte de un amplio y diverso territorio denominado Sabana de Bogotá (SB) el cual cubre una extensión aproximada de 2500km$^2$ en donde la biodiversidad permitió el desarrollo de múltiples grupos humanos con trayectorias históricas particulares, pero que compartieron algunos rasgos culturales que se fortalecieron con el paso del tiempo.*

*Durante más de 2000 años el sitio Nueva Esperanza fue habitado por familias indígenas muiscas, las cuales lentamente convirtieron el lugar en uno de los más importantes poblados de la región y en donde tuvieron lugar diversas formas de expresión política, económica e ideológica las cuales cambiaron a través del tiempo, registradas arqueológicamente como periodo Herrera (400 a.C. – 200 d.C.) Muisca Temprano (200 d.C. – 1000 d.C.) y Muisca Tardío (1000 d.C. – 1600 d.C.).*

*Las evidencias arqueológicas indican que los primeros habitantes de NE conformaron familias que en muchos casos extendieron sus linajes hasta el periodo Muisca Tardío, construyendo estructuras habitacionales de planta circular y rectangular, las cuales además de ser usadas para vivir y desarrollar actividades domésticas y económicas, también fueron espacios para enterrar y rendirle culto a sus muertos. La agricultura y producción de textiles fueron al parecer los dos pilares económicos de esta población, logrando consolidar rutas de intercambio comercial con poblaciones vecinas dentro del área muisca, y en territorios más lejanos, cubriendo distancias de hasta 950 kilómetros aproximadamente en la actual Sierra Nevada de Santa Marta, ocupada por los Taironas.*

*Los habitantes de Nueva Esperanza, expresaron a partir de las prácticas funerarias, fenómenos culturales como la desigualdad social. En las tumbas asociadas al periodo Herrera se han registrado indicadores materiales asociados al poder y el prestigio político, los cuales no eran ostentados por todos los pobladores y nos permiten inferir diferencias al acceso de recursos, los cuales podrían ser controlados por grupos sociales con algún tipo de parentesco ya que las tumbas de este periodo se caracterizaron por la inhumación colectiva.*

*Durante el periodo Muisca Temprano, las prácticas de enterramiento se transformaron respondiendo a cambios políticos e ideológicos en donde el prestigio pudo ser alcanzado por acciones individuales. Sin embargo, estos procesos tomaron tiempo y se consolidaron lentamente en un lapso de tiempo de 800 años aproximadamente. Aproximadamente en el año 1000 d.C., se registraron nuevos cambios culturales, los cuales se reflejaron en el patrón de funerario, en donde no solo se privilegió la individualidad, también se identificaron cambios en transmisión del poder o prestigio posiblemente se obtenía por línea consanguínea sin importar los méritos personales.*
*Hasta el momento se han excavado arqueológicamente cerca de 10 hectáreas y se han identificado más de 10.000 rasgos, cerca de 3.000 cuerpos en estado esquelético y decenas de miles de objetos elaborados en cerámica, piedra, hueso y metales que dan cuenta de las trayectorias históricas de los pobladores de Nueva Esperanza.*

**Portalegre (Cundinamarca, Bogotá)**
The Portalegre settlement (PTG) (4° 35´12.55" N, 74° 12´ 50.17" W) is located in the Soacha county south Bogotá city near the homonym river. The excavated area of 1200 $m^2$ indicates that the site corresponds to a large settlement with its corresponding cemetery[92]. At least seven residential platforms of circular shape (*Bohíos*) were identified with a size that ranges from 5.2 m to 6.2 m. The *Bohíos* were located very close each other and some of them were superimposed which has been interpreted as a continuous use of the site[93]. At least 130 burials were found with a high number of human skeletons and very diverse grave goods that includes different kinds of pottery, spindle whorls, shell necklace beads, animal bones tools and *Tumbaga* pieces[92,93]. The human skeletons were found in supine position mostly located in a south and east direction[33]. Only six adult individuals have been investigated for stable isotopes[13]. The chronology of the site is given for two radiocarbon dates 915±115 BP (GX-18842 human bone) and 720±110 BP (GX-18841 human bone), which were corrected for isotopic fractionation (corrected dates: 1115±115 BP and 920±110 BP using a mean value $\delta^{13}C$ -12.8‰) indicating that those individuals belong to the Early Muisca period.

**San Francisco (Cundinamarca, Bogotá)**
The San Francisco site dates to the Late Muisca Period; four direct 14C dates on bone material suggest occupation of the site between 1230-1377 calCE and 1457-1620 calCE (**Table S1**). We sampled eight individuals for ancient DNA and successfully generated data from seven, all attributed to Late Muisca contexts. Genetic outlier individual I10932 comes from San Francisco (**Supplementary Information 4**).

**Terregrande (Cundinamarca, Bogotá)**
The site of Terregrande provides important insight into the late pre-Hispanic occupation of the southern Sabana de Bogotá. Direct 14C dates on bone material newly reported in this work place the occupation of the site in the Muisca Period as early as 895-1023 calCE and spanning through 1224-1275 calCE (**Table S1**). Four individuals were sampled for ancient DNA analysis. All were evaluated as having cranial deformation. Only one individual yielded authentic ancient DNA data that passed quality control checks (**Table S1**).

**Terreros (Cundinamarca, Bogotá)**
The site of Terreros provides important insight into the late pre-Hispanic occupation of the southern Sabana de Bogotá. Direct $^{14}C$ dates on bone material newly reported in this work place the occupation of the site in the Muisca Period as early as 895-1023 calCE and spanning at least through 1322-1410 calCE (**Table S1**).

**Tibanica (Cundinamarca, Bogotá)**
The *Alameda de Tibanica* settlement (TIB) located in the Soacha County south of Bogotá was excavated since 2005 and yielded nearly 700 burials with abundant cultural and biological remains (faunal and botanical remains, pottery, lithics, and human skeletal remains) [94]. There are nearly 50 associated radiocarbon dates from the site. The site corresponds to a circular settlement with habitation complexes, dump areas and a cemetery integrated by four burial groups[95]. Different

cultural evidence (pottery, burial form, etc.) as well as seventeen AMS radiocarbon dates obtained from human bone allows place the site occupation between 640±20 BP and 990±20 BP[95-97].Data about Tibanica can be found in refs. [96,98,99].

##### *No ancestry differences among groups at Tibanica*

A key archaeological question is whether genetic ancestry differences among individuals from Tibanica are detectable at the resolution of allele-sharing statistics. In our dataset, we have working genome-wide data from 24 individuals from Tibanica from elite and non-elite contexts who are divided archaeologically into 5 groups (Tibanica 0-4). All date to Late Muisca times (~1200-1600 CE).

**Supplementary Table 1.1. Division of 24 people from Tibanica into 5 archaeological groups with elite and non-elite designations.**

| Group – Burial Context | Number of individuals in our study |
|---|---|
| Group 0 – Elite | 1 |
| Group 0 – Non-Elite | 2 |
| Group 1 – Elite | 3 |
| Group 1 – Non-Elite | 2 |
| Group 2 – Elite | 3 |
| Group 2 – Non-Elite | 3 |
| Group 3 – Elite | 2 |
| Group 3 – Non-Elite | 2 (genetic outlier I22700 is also in this group) |
| Group 4 – Elite | 2 |
| Group 4 – Non-Elite | 3 |

With PCA, we see no difference in ancestry between elite/non-elite people from Tibanica (**Figure 2**). With $f_4$ statistics of the form $f_4$*(Yoruba, NativeAmerican_Test; TibanicaA, TibanicaB)* for all permutations of the Tibanica groups (elite/non-elite contexts from groups 0-4), we find no significant evidence of systematic differences in genetic ancestries between elite and non-elite people buried at Tibanica at the limits of our resolution (**Table S27**). An exception is for outlier individual I22700 who shares less drift with essentially all Native American reference populations and is discussed further in **Supplementary Information 3** and **4**. We merged people from elite contexts from all cemeteries and non-elite contexts from all cemeteries for more power and ran the statistic $f_4$*(Yoruba, NativeAmerican_Test; Tibanica_Elite, Tibanica_NonElite)*. We still identified no systematic ancestry differences between these groups (**Table S27**). Likewise, systematic genetic differences between groups at Tibanica are identified with *qpWave* (**Table S22**).

**Supplementary Information 2: Isotope analysis**

For 33 individuals with genome-wide data passing quality control and direct $^{14}C$ dates, we also generated dietary isotope information (**Table S14**). Here we detail the methods to generate these data and discuss dietary changes on the Altiplano Cundiboyacense as documented by dietary isotopes.

#### Methods

***PSU (sample preparation and chemistry)/YASIC or LIME (measurement); n=19***

Dietary isotope data were generated for 19 samples following sample preparation and chemistry at the Pennsylvania State University Radiocarbon Laboratory and measurement at the YASIC facility at Yale University (n=13) and the LIME facility at Penn State (n=6).

After physical cleaning, all of the bone samples (200-1000mg each) were rinsed in successive washes of American Chemical Society (ACS)-grade methanol, acetone and dichloromethane for 20 min each at room temperature, followed by multiple washes in >18.2 MOhm/cm water, to remove possible consolidants, glues, etc. Bone collagen was extracted following a modified Longin method, demineralizing the samples in 0.5N HCl for 2-3 days at ~5°C. The resulting pseudomorph was gelatinized in 0.01N HCl for 10h at 60°C, lyophilized and weighed to estimate crude collagen yield and evaluate gelatin quality. Well preserved gelatin (e.g., high yield, white to pale tan color, opalescent sheen) was rehydrated and ultrafiltered in precleaned CentriPrep or Amicon centrifugal filters with nominal 30kDa molecular weight cutoff, centrifuged three times for 20 min, diluted with >18.2 MOhm/cm $H_2O$, and centrifuged three more times for 20 minutes to desalt the solution. Poorly preserved gelatin was processed using amino acid hydrolysis and XAD purification[100]. Gelatin was hydrolyzed in 2ml of 6N HCl for 24 hours at 110°C. Supelco ENVI-Chrom SPE (solid-phase extraction; Sigma-Aldrich) columns with 0.45-μm polyvinylidene difluoride filters were equilibrated with 50 ml of 6N HCl and the washings discarded. Two milliliters of hydrolysate were pipetted onto the SPE column and driven with an additional 10 ml of 6N HCl dropwise with a syringe into a 20-mm culture tube. The hydrolysate was dried into a viscous syrup by passing UHP (ultra-high purity) $N_2$ gas over the sample heated at 50°C for ~12 hours.

Carbon and nitrogen isotope ratios were measured on 0.7mg/1.0mg of the extracted and purified ultrafiltered gelatin/amino acid hydrolysate packed in Sn boats. Carbon and nitrogen concentrations and stable isotope ratios were measured at the Yale Analytical and Stable Isotope Center (YASIC) with a Costech elemental analyzer (ECS 4010) and Thermo DELTA Plus analyzer. Samples submitted to the Penn State LIME lab were measured with a Costech elemental analyzer (ECS 4010) and Thermo DELTA V analyzer. Sample quality was evaluated by % crude gelatin yield, %C, %N, and C/N ratios (atomic).

We combusted ultrafiltered gelatin/amino acid hydrolysate samples (~2.1 mg/4.5mg) for 3h at 900°C/800°C in vacuum-sealed quartz tubes with CuO and Ag wires. Sample $CO_2$ was cryogenically purified and reduced to graphite at 550°C using $H_2$ and a Fe catalyst, drawing off reaction water with $Mg(ClO_4)_2$[101]. We pressed graphite samples into aluminum cathode blank and loaded them onto a target wheel with process-specific backgrounds and secondaries and the OXII oxalic acid primary standard. $^{14}C$ measurements were made on a modified National Electronics Corporation compact accelerator mass spectrometer with a 0.5 MV accelerator (NEC 1.5SDH-1). Sample measurements were normalized to the OXIIs, background corrected with process-specific background measurements and $^{13}C$-corrected with the $13^+/12^+$ ratio measured on the AMS to produce the reported conventional radiocarbon ages.

***CSI; n=4***

Dietary isotope data were generated for 4 samples at the CSI facility at the University of New Mexico. This number includes stable isotope and $^{14}C$ data for I28092 which were generated at this facility as well as at UCI.

Two methods were used for extracting bone collagen for AMS 14C measurements: ultrafiltration[102] and XAD-purification[103]. Bone collagen was first extracted and purified using the modified Longin[104] method followed by ultrafiltration[105] or XAD amino acid purification[106] depending on the preservation of the sample. Bone material was suspended in 0.5 N HCl at 5 °C for 24-48 hours until the bone matrix was demineralized, followed by two rinse cycles of UltraPure 18Ω H2O until neutrality. The resulting collagen pseudomorph was gelatinized for 12 hours at 60 °C in 0.01 N HCl and lyophilized for 48 hours. Samples were resuspended in UltraPure 18Ω H2O and transferred to pre-cleaned Sartorius Vivaspin 20 ultrafilters (to retain >30 kDa molecular weight gelatin) and centrifuged 6 times. Purified collagen samples were lyophilized for 48 hours and evaluated for crude gelatin yields. Samples with low collagen yields were further processed using XAD-purification to isolate amino acids in the ultrafiltered bone collagen[106]. The purified collagen was hydrolyzed in 1.5 mL 6 N HCl for 22 hours at 110 °C. The 2 mL collagen hydrolysate solution was transferred to a dropwise syringe and passed through a pretreated Supelco ENVI- Chrom® SPE (Solid Phase Extraction; SigmaAldrich) column and eluted with 10 mL 6 N HCl in a 20 mm culture tube. The purified hydrolysate was heated to 50 °C and dried under UHP N2 gas for ~12 hours. Sample quality was evaluated by % crude gelatin yield, and %C and %N, C:N ratios to assess collagen preservation and contamination[107]. Approximately 0.7 mg of ultrafilter bone sample and 1.5 mg XAD-purified bone collagen were weighted into tin capsules and analyzed on a Thermo Scientific Delta V mass spectrometer with a dual inlet and Conflo IV interface connected to a Costech 4010 elemental analyzer (EA). Samples with Atomic C:N rations between 3.0 and 3.4 progressed to sample combustion and graphitization. Samples were combusted at 900°C in vacuum sealed quartz tubes along with prebaked CuO2 and Ag wire. Resulting C02 was cryogenically purified under vacuum and reduced using a Bosch reaction onto SigmaAldrich Fe powder at 550°C in an H2 atmosphere.The resulting Fe/graphite powder was pressed into 1mm Al targets and send the PSUAMS facility for 14C measurements.

***UCI[108]; n=10***

Dietary isotope data were generated for 10 samples at the UCI Keck Carbon Cycle AMS Facility. This number includes stable isotope and $^{14}C$ data for I28092 which were generated at this facility as well as at CSI.

Bone collagen for AMS $^{14}C$ dating was extracted using a modified Longin[104] collagen extraction followed by ultrafiltration[105]. Mechanically cleaned, crushed bone (typically 50–150 mg) was decalcified for 24-36 h in HCl (0.5 N HCl in the UCI protocol[108]; UCI additionally reported 1 N HCl for samples included here), then rinsed with Milli-Q water to near-neutral pH. Demineralized collagen pseudomorphs were gelatinized overnight at 60 °C in weak acid (0.01 N HCl; UCI reported gelatinization at 60 °C, pH ~2 for samples reported here), and the gelatin was purified by ultrafiltration using pre-cleaned Centriprep YM-30 devices (30 kDa MWCO; YM-10 used in some cases). Ultrafiltration included repeated centrifugation and dilution steps to concentrate the >30 kDa fraction and reduce salts, after which purified gelatin was transferred to heavy-walled tubes, frozen, and freeze-dried overnight for yield determination. For measurement, collagen-derived carbon was converted to $CO_2$ by combustion in pre-baked quartz tubes with CuO and Ag wire; tubes were evacuated, sealed, and combusted at 900°C for ~3 h. $CO_2$ was cryogenically purified and quantified, then reduced to graphite via the Bosch reaction using $H_2$ over −325 mesh Fe catalyst, with water trapped by $Mg(ClO_4)_2$; Fe catalyst was preconditioned under $H_2$ prior to use. Graphitization was run at 550 °C with $H_2$ added at ~2× the reactor $CO_2$ pressure for a ~3 h reaction cycle while reactor pressures were monitored. Graphite was pressed into aluminum targets and mounted in a 40-position UCI wheel. UCI reported subtraction of preparation backgrounds based on $^{14}C$-free mammoth and whale bone. Reported results follow fraction modern conventions and are corrected for isotopic fractionation using $\delta^{13}C$ measured on prepared graphite by the AMS system.

***UGA; n=1***

Dietary isotope data were generated for 1 sample at the Center for Applied Isotope Studies (CAIS), University of Georgia.

Collagen was extracted using a modified Longin[104] collagen extraction, modified as described here. A sub-sample of bone was removed using a Dremel tool outfitted with a diamond cutting wafer. Surface contamination was removed from the sub-sample using a scalpel and wire-bristle brush; the sub-sample was simultaneously reduced to smaller fragments (approximately 3–5mm in size). These small fragments were demineralized in cold (4°C) 1N HCl for 24 hours, the acid was decanted, and demineralized fragments of bone were rinsed three times with ultrapure water (MilliQ). The bone fragments were then treated with 0.1M NaOH to dissolve and remove humic acids, followed by a series of ultrapure water rinses. Atmospheric CO2 was eliminated through the rinsing of bone fragments with cold 1N HCl. The fragments were then rinsed again in ultrapure water to ~ pH 4 (slightly acidic) and heated at 80ºC for 8 hours. The solution was subsequently filtered through a glass fiber filter, isolating the total acid insoluble fraction ("collagen"), which was then freeze-dried. A ~5mg sub- sample of collagen was combusted at 575°C in an evacuated and sealed Pyrex tube in the presence of CuO, producing CO2. The CO2 sample was cryogenically purified from the other reactionproducts and catalytically converted to graphite following the method of ref.37. Graphite 14C/13C ratios were measured using the 0.5 MeV accelerator mass spectrometer (AMS) housed at CAIS. Sample ratios were compared to the ratio measured from the Oxalic Acid I standard (NBS SRM 4990). All results are presented as percent Modern Carbon (pMC). The quoted uncalibrated dates are given in radiocarbon years before 1950 (years BP), using a 14C half-life of 5568 years. The date has been corrected for isotope fractionation using the δ13C value measured by EA-IRMS.

**Isotope research on the ancient populations of the Altiplano Cundiboyacense**

Stable carbon and nitrogen isotope analyses from the Altiplano Cundiboyacense provide insight into changes in subsistence strategies across the Holocene. In humans, $\delta^{13}C$ values measured on bone collagen ($\delta^{13}C_{collagen}$) primarily reflect the values of protein an individual consumed, more specifically the diet of animals consumed with smaller contributions from plant carbohydrates[109].Omnivores like humans derive most of their protein from animal sources, which are routed directly from source to consumer tissues[110].A lesser amount of protein is derived from plants. Enriched $\delta^{13}C_{collagen}$ thus primarily reflects the consumption of resources like marine foods or $C_4$ plants with high carbon-13 ($^{13}C$) relative to carbon-12 ($^{12}C$), or in the case of nursing infants, the diets of their mothers[111].Conversely, $\delta^{13}C$ values measured from hydroxyapatite (tooth or bone apatite) reflect primarily the isotopic values of dietary carbohydrates, with much smaller contributions from proteins and lipids; $\delta^{13}C_{apatite}$ is generally interpreted as an approximate measure of the "whole diet" [112]. Elevated $\delta^{13}C_{collagen}$ values can be interpreted as reflecting the presence of $C_4$ plants (like maize) in a food web that includes humans along with the animals they eat and primary producer plants at the base of that food web.

Highland Andean environments, like most neotropical forests and savannas, are dominated by $C_3$ vegetation, with local vegetation values on the Sabana de Bogotá ranging from −32‰ to −22‰[113]. $C_4$ taxa are uncommon: Poaceae (grasses) are largely restricted lower elevations and Cyperaceae (sedges and related monocots) are found primarily in marginal water-stressed environments; both have $\delta^{13}C$ values ranging from −16 to −11‰[114,115]. Archaeofaunal data from deer (*Odocoileus virginianus*) and guinea pigs (*Cavia* sp.) at Tequendama and Aguazuque that date to the terminal Pleistocene and Holocene have $\delta^{13}C$ values consistent with $C_3$ forest environments[113]. This suggests minimal incorporation of $C_4$ biomass into local animal food webs. In this ecological context, elevated human $\delta^{13}C$ values cannot be readily attributed to background vegetation or wild herbivore consumption and instead reflect meaningful dietary shifts. Maize and amaranth (*Amaranthus* sp.), are the only $C_4$ plants cultivated regionally, exhibiting high $\delta^{13}C$ values[116]. Archaeobotanical analysis of micro-remains recovered from lithic tools surfaces and dental calculus have documented the presence of maize starch grains, suggesting that maize was the most likely $C_4$ resource to contribute to the signal observed in this context[22,117-119].

Previous research in the Sabana de Bogotá linked enriched $\delta^{13}C_{collagen}$ values in humans to the incorporation of $C_4$ plants into local human food-webs[113]. Animals consume human cultivated foods both through direct animal management or because wild animals feed on cultivated crops and are then hunted. Garden hunting, where animal prey is opportunistically selected from cultivated fields is well documented globally. Varying degrees of human-animal interactions have been proposed to explain $C_4$ consuming animals found in lowland South America[120], Panama[121], and the Caribbean[122].

Importantly, the timing and magnitude of isotopic change identified by ref. [113] (as well as previous work, including refs. [123-125]) – elevated $\delta^{13}C$ values (-10 to -11‰) at Aguazuque ~2900-2750 calBP –

align closely with patterns observed in our dataset, including enriched $\delta^{13}C$ values in some individuals from Aguazuque during the initial late Holocene. When considered alongside our ancient DNA data, these isotopic data demonstrate that the incorporation of $C_4$ resources, likely maize, occurred during a period of long-term genetic continuity on the Sabana de Bogotá. This decoupling of subsistence change from population replacement provides strong evidence that early agricultural practices were adopted locally through cultural transmission rather than introduced by incoming populations, with major demographic turnover occurring only later.

#### Isotopic data newly reported in this work

##### *Early–middle and middle Holocene*

Previous isotopic work[123-125] showed that early Holocene hunter-gatherers consumed mostly (80-100%) $C_3$ vegetal resources with a lesser emphasis on animal protein. Middle Holocene hunter-gatherers also primarily consumed $C_3$ vegetal resources with a slight increase in C3 animal protein intake[123-125].

In our data, the earliest individuals from the sites of Aguazuque and Sueva I exhibit tightly clustered $\delta^{13}C$ values between –20.5 and –19.4‰. These values are characteristic of consumers whose diets are high in $C_3$ resources. These individuals show no isotopic evidence for the consumption of $C_4$-enriched resources. Corresponding $\delta^{15}N$ values are relatively low, particularly in the earliest individual (e.g., I28092: ~5.0–5.3‰), suggesting protein intake from low trophic-level terrestrial or avian animals.

##### *Initial Late Holocene*

Previous work shows a trend toward mixed $C_3/C_4$ diets during the initial late Holocene, with some suggesting that this reflects a move toward a food production system[12,13,123]. Research has suggested that $C_4$ plants were present on the Sabana de Bogotá by ~3500 calBP[123,124]. Archaeobotanical evidence from across the Sabana de Bogotá for sweet potato (*Ipomoea batata*), calabash tree (*Cresentia cujete*), avocado (*Persea americana*), cherry tree (*Prunos serotina*), and maize (grains and cobs of *Zea mays*) [1,10]suggests that by the initial late Holocene, $C_3$ and $C_4$ domesticates were being consumed on the Sabana de Bogotá. Refs. [123-125] suggest that the period between 3800-2800 BP was the main dietary change during which people shifted to the consumption of significantly more $C_4$ plants and animals (although it is possible that this date may be as early as 4000 BP).

In our data, a pronounced isotopic shift is first observed in the initial late Holocene. I23785, directly dated to 983-835 calBCE (2770±20 BP, UCIAMS-288524), exhibits a $\delta^{13}C_{collagen}$ value of –10.6‰, representing an enrichment of approximately +9‰ relative to earlier individuals. A shift of this magnitude cannot be explained by environmental baselines alone and instead is likely to indicate dietary input from $C_4$ plants, almost certainly maize. Notably, this isotopic signal appears only in two initial late Holocene individuals who lived ~2800 BP (I23785 and I23783, an individual who is dated by context to 2725±35 BP (GrN-14479) and has isotopic data published in van der Hammen et al. 1990) and not in individuals who lived only several hundred years prior at this same site, for example, I22456 and I13218 who are dated to 2128-1781 calBCE (3600±40 BP, UNAM-Col-AAA) and 2340-

2146 calBCE (3810±20 BP, PSUAMS-5501), respectively. Two other individuals also from Aguazuque also dated by context to 2725±35 BP (GrN-14479) lack ancient DNA data but have the same isotopic signal reflecting dietary input from $C_4$-derived carbon. Thus, our data are consistent with the incorporation of $C_4$ plants into the food web on the Sabana de Bogotá between 3800 and 2800 BP.

***Herrera and Muisca Periods***

Previous work indicates increasing reliance on agriculture during the final late Holocene and varied diets comprising $C_4$ and $C_3$ crops, $C_3$-$C_4$ feeding animals, and fresh-water resources[123,124,126].

In our data, individuals dating to the initial and final late Holocene associated with Herrera and Muisca contexts show substantial inter-individual variability in $\delta^{13}C_{collagen}$ values, ranging from approximately –15.4 to –8.0‰. These values indicate mixed $C_3$-$C_4$ diets, implying varying degrees of maize consumption both within and between sites. The absence of a narrow $\delta^{13}C_{collagen}$ range suggests that agricultural intensification did not result in dietary homogenization. Instead, it is more indicative of maize being integrated alongside other plant and animal resources, consistent with a diversified subsistence economy.

**Previously published isotope data used for Figure 4**

**Supplementary Table 2.1. Previously published isotope data for 50 individuals with corresponding genetic IDs for 47.**

| Site | Archaeological ID | Genetic ID | 14C date or age estimated with archaeological stratum date or context | $\delta^{13}C_{coll}$ | $\delta^{15}N_{coll}$ | Reference for isotope data |
|---|---|---|---|---|---|---|
| **Checua** | Individual 1 (Individuo 1) | I22316 | 3795-3649 calBCE (4960±30 BP, Beta-503359) | -19.6 | 6.6 | Arias Conchas 2019[127] |
| **Checua** | Individual 2 (Individuo 2) | I22317 | 3706-3632 calBCE (4875±22 BP) [R_combine: (4850±30 BP, Beta-493119); (4900±30 BP, Beta-509235)] | -19.3 | 7.2 | Arias Conchas 2019[127] |
| **Checua** | Individual 3 (Individuo 3) | I22318 | 3766-3638 calBCE (4910±30 BP, Beta-493120) | -19.6 | 7.4 | Arias Conchas 2019[127] |
| **Checua** | Individual 4 (Individuo 4) | I22319 | 3766-3638 calBCE (4910±30 BP, Beta-493121) | -19.3 | 7.8 | Arias Conchas 2019[127] |
| **Checua** | Individual 6 (Individuo 6) | I22320 | 3771-3645 calBCE (4930±30 BP, Beta-503360) | -19.2 | 6.2 | Arias Conchas 2019[127] |
| **Checua** | Individual 9 (Individuo 9) | I22323 | 3771-3645 calBCE (4930±30 BP, Beta-503362) | -19.5 | 6.8 | Arias Conchas 2019[127] |
| **Checua** | Individual 10 (Individuo 10) | I22324 | 4841-4711 calBCE (5900±30 BP, Beta-503363) | -20.0 | 6.6 | Arias Conchas 2019[127] |
| **Checua** | Individual 12 (Individuo 12) | I22325 | 4609-4450 calBCE (5690±30 BP, Beta-503364) | -19.6 | 8.2 | Arias Conchas 2019[127] |
| **Checua** | Individual 13 \| Individual 22 (Individuo 13 \| Individuo 22) | I22326 | 5209-4996 calBCE (6135±22 BP) [R_combine: (6140±30 BP, Beta-503365), (6130±30 BP, Beta-503366)] | -19.4 | 5.1 | Arias Conchas 2019[127] |
| **Checua** | Individual 15 (Individuo 15) | I22327 | 5209-5004 calBCE (6150±31 BP, Beta-493123) | -19.9 | 6.1 | Arias Conchas 2019[127] |
| **Checua** | Individual 5 (Individuo 5) | I22469 | 3943-3655 calBCE (5000±30 BP, Beta-493122) | -19.8 | 9.3 | Arias Conchas 2019[127] |
| **Checua** | Individual 11 (Individuo 11) | I22470 | 5292-5046 calBCE (6200±30 BP, Beta-493124) | -20.2 | 5.8 | Arias Conchas 2019[127] |
| **Checua** | Individual 16 (Individuo 16) | I22622 | 3765-3636 calBCE (4900±30 BP, Beta-493125) | -19.4 | 8.3 | Arias Conchas 2019[127] |
| **Checua** | Individual 20 (Individuo 20) | I22623 | 3763-3632 calBCE (4890±30 BP, Beta-493128) | -19.1 | 7.5 | Arias Conchas 2019[127] |
| **Checua** | Individual 18 (Individuo 18) | I22628 | 3946-3708 calBCE (5020±30 BP, Beta-493127) | -19.9 | 6.7 | Arias Conchas 2019[127] |
| **Tibanica** | Ind. 945 (Non-Elite; Group 1) | I23744 | 1000-1600 CE | -12.0 | 11.9 | Aristizábal Losada 2015[128] |
| **Tibanica** | Ind. 1188 (Elite;Group 2) | I23746 | 1000-1600 CE | -12.1 | 9.2 | Aristizábal Losada 2015[128] |
| **Tibanica** | Ind. 1189 (Elite;Group 2) | I23747 | 1000-1600 CE | -13.0 | 10.0 | Aristizábal Losada 2015[128] |
| **Tibanica** | Ind. 3237 (Elite; Group 4) | I24400 | 1250-1450 CE | -10.0 | 9.2 | Aristizábal Losada 2015[128] |
| **Las Delicias** | 90-IX-001 (LD-01) | I22486 | 800-1200 CE | -13.7 | 8.9 | Cárdenas 1993[69] |
| **Las Delicias** | 90-IX-002 (LD-02) | I22487 | 800-1200 CE | -12.9 | 9.5 | Cárdenas 1993[69] |
| **Las Delicias** | 90-IX-15 (LD-15) | I22488 | 800-1200 CE | -11.8 | 9.0 | Cárdenas 1993[69] |
| **Las Delicias** | 90-IX-11 (LD-11) | I22489 | 800-1200 CE | -10.4 | 9.6 | Cárdenas 1993[69] |
| **Candelaria La Nueva** | 87-X-0004 | I22460 | 1200-1500 CE | -11.4 | 11.3 | Cárdenas 1996[32] |

| | | | | | | |
|---|---|---|---|---|---|---|
| **Candelaria La Nueva** | 87-X-0041 | I22462 | 1200-1500 CE | -11.4 | 8.6 | Cárdenas 1996[32] |
| **Candelaria La Nueva** | 87-X-0007 | I22466 | 1200-1500 CE | -13.8 | 9.4 | Cárdenas 1996[32] |
| **Aguazuque** | AZ50 (AZ-458-50) | I23672 | 2831-2466 calBCE (Context date based on date of stratum 4.1, 4030±35 BP (GrN-12930)) | -20.0 | 9.8 | Cárdenas 2002[13] |
| **Tequendama** | TEQ1-E12C (TEQ-I E12) | I24982 | 6227-5996 calBCE (7235±60 BP, GrN-7477) | -20.5 | 8.6 | Cárdenas 2002[13] |
| **Ubaté** | UB03 (Individual 3) | I23893 | 4534-4361 calBCE (5620±30, Beta-398660) | -19.7 | 8.2 | Archila and Langebaek 2015[129] |
| **Tibanica** | Ind. 1054 (Elite; Group 1) | I22698 | 1300-1600 CE | -10.7 | 8.1 | Miller 2016[97] |
| **Tibanica** | Ind. 2809 (Non-Elite; Group 0) | I22699 | 1300-1600 CE | -10.7 | 10.1 | Miller 2016[97] |
| **Tibanica** | Ind. 50 (Non-Elite; Group2) | I23738 | 1300-1600 CE | -11.2 | 8.7 | Miller 2016[97] |
| **Tibanica** | Ind. 177 (Non-Elite; Group2) | I23740 | 1000-1600 CE | -10.5 | 10.4 | Miller 2016[97] |
| **Tibanica** | Ind. 924 (Elite; Group 1) | I23742 | 1000-1600 CE | -14.1 | 9.5 | Miller 2016[97] |
| **Tibanica** | Ind. 947 (Non-Elite; Group 1) | I23743 | 1000-1600 CE | -13.0 | 9.5 | Miller 2016[97] |
| **Tibanica** | Ind. 1186 (Elite;Group 2) | I23745 | 1000-1600 CE | -12.7 | 10.1 | Miller 2016[97] |
| **Tibanica** | Ind. 2888 (Elite; Group 3) | I24393 | 1300-1600 CE | -11.4 | 9.9 | Miller 2016[97] |
| **Tibanica** | Ind. 3020-b (Elite; Group 1) | I24394 | 1300-1600 CE | -9.7 | 10.4 | Miller 2016[97] |
| **Tibanica** | Ind. 3119 (Non-Elite; Group 0) | I24395 | 1300-1600 CE | -11.3 | 9.6 | Miller 2016[97] |
| **Tibanica** | Ind. 3162 (Non-Elite; Group 4) | I24397 | 1250-1450 CE | -12.3 | 10.0 | Miller 2016[97] |
| **Tibanica** | Ind. 3164 (Non-Elite; Group 4) | I24398 | 1300-1600 CE | -8.2 | 9.9 | Miller 2016[97] |
| **Tibanica** | Ind. 3189 (Non-Elite; Group 4) | I24399 | 1300-1600 CE | -12.0 | 9.6 | Miller 2016[97] |
| **Nueva Esperanza** | OHI-364 (TCE09-H9-N3-ACU4-R58B-T33-I63) | I23728 | 500 BCE-300 CE | -10.5 | 8.3 | Rivas et al. 2024[126] |
| **Aguazuque** | AGZ-4-9 (AGZ-C4-R9-E2, AGZ; Column 4; Rasgo 9) | I22456 | 2128-1781 calBCE (3600±40 BP, UNAM-Col-AAA) | -19.5 | 6.8 | Triana-Vega et al. 2020[18] |
| **Tequendama** | TEQ/AT (TEQ1-R9-E2, Rasgo 6) | I22819 | 5206-4846 calBCE (6080±40 BP, UNAM-Col-AAA) | -19.9 | 6.4 | Triana-Vega et al. 2020[18] |
| **Aguazuque** | AZ32 (AZ-683 32 E5.2) | I23783 | 912-797 calBCE (Context date based on date of stratum 5.2, 2725±35 BP (GrN-14479)) | -10.5 | 8.9 | van der Hammen et al. 1990[12] |
| **Aguazuque** | AZ51 (AZ-683 51 E5.1) | I23787 | 2831-2466 calBCE (Context date based on date of stratum 4.1, 4030±35 BP (GrN-12930)) | -19.2 | 8.9 | Correal 1990[1] |
| **Aguazuque** | AZ-m.2 85 cm | no DNA | 2457-2204 calBCE (3850 ± 35 BP (GrN-Col.593)) | -20.2 | 8.7 | van der Hammen et al. 1990[12] |
| **Aguazuque** | AZ-683 33 E5.2 | no DNA | 912-797 calBCE (Context date based on date of stratum 5.2, 2725±35 BP (GrN-14479)) | -11.2 | 9.4 | van der Hammen et al. 1990[12] |
| **Aguazuque** | AZ-m.3-40 cm | no DNA | 912-797 calBCE (Context date based on date of stratum 5.2, 2725±35 BP (GrN-14479)) | -11.0 | 8.7 | van der Hammen et al. 1990[12] |

**Supplementary Information 3. Assessing genetic homogeneity with *qpWave***

We assessed genetic homogeneity in our newly reported data and data from ref. [130] to understand if and how we could meaningfully group ancient individuals for increased power in statistical analyses. We used the software *qpWave* applied to the 2M.HO dataset, carrying out pairwise tests to assess if individuals or groups were genetically homogenous relative to a genetically diverse reference set (also referred to as an outgroup set, or "right" populations): Chipewyan, Zapotec, Mixe, Mixtec, Surui, Cabécar, Piapoco, Karitiana, Inga, Wayuu, Apalai, Arara. For all *qpWave* modeling, we set 'allsnps: YES', 'inbreed: NO', and 'basepop: Han.DG'.

**Assessing genetic homogeneity among the pre-Herrera people of the Sabana de Bogotá**

We began by running pairwise *qpWave* on 45 individuals from pre-Herrera hunter-gatherer and subsistence transition contexts newly-reported in this work as well as the seven previously published individuals from Checua who lived ~6000 BP[130] (**Table S2**; **Figure S3**). We generated pairwise *qpWave* p-values and reshaped them into a symmetric matrix with each cell representing the p-value (the "fit") of the model. To highlight significantly poor fits, all p-values < 0.05 were assigned a gray color in the heatmap. We applied hierarchical clustering using complete linkage and Euclidean distance, grouping individuals based on the similarity of their *qpWave* p-value profiles.

With this approach we identified eight individuals who were subtly genetically dissimilar from others relative to the reference set (**Supplementary Figure 3.1**):

- I22323
- PREC007[130]
- I22326
- I23898 (note: low resolution, <50K SNPs covered on 2M.HO dataset)
- I22623 (note: low resolution, <50K SNPs covered on 2M.HO dataset)
- I22318
- I23671 (note: low resolution, <50K SNPs covered on 2M.HO dataset)
- I23787 (note: low resolution, <50K SNPs covered on 2M.HO dataset)

These subtle deviations should be interpreted with caution, as *qpWave* is sensitive to missing data and reduced SNP overlap, and several of the individuals identified here are represented by fewer than 50,000 SNPs, raising the possibility that their distinctiveness reflects technical artifacts rather than genuine ancestry differences.

PCA outlier I23673 was not assessed as genetically dissimilar to other individuals with *qpWave*.

**Supplementary Figure 3.1.** Symmetry matrix of *qpWave* values with each cell representing the fit (p-value) of the model. Gray cells represent p-values < 0.05. Clustering approach described above.

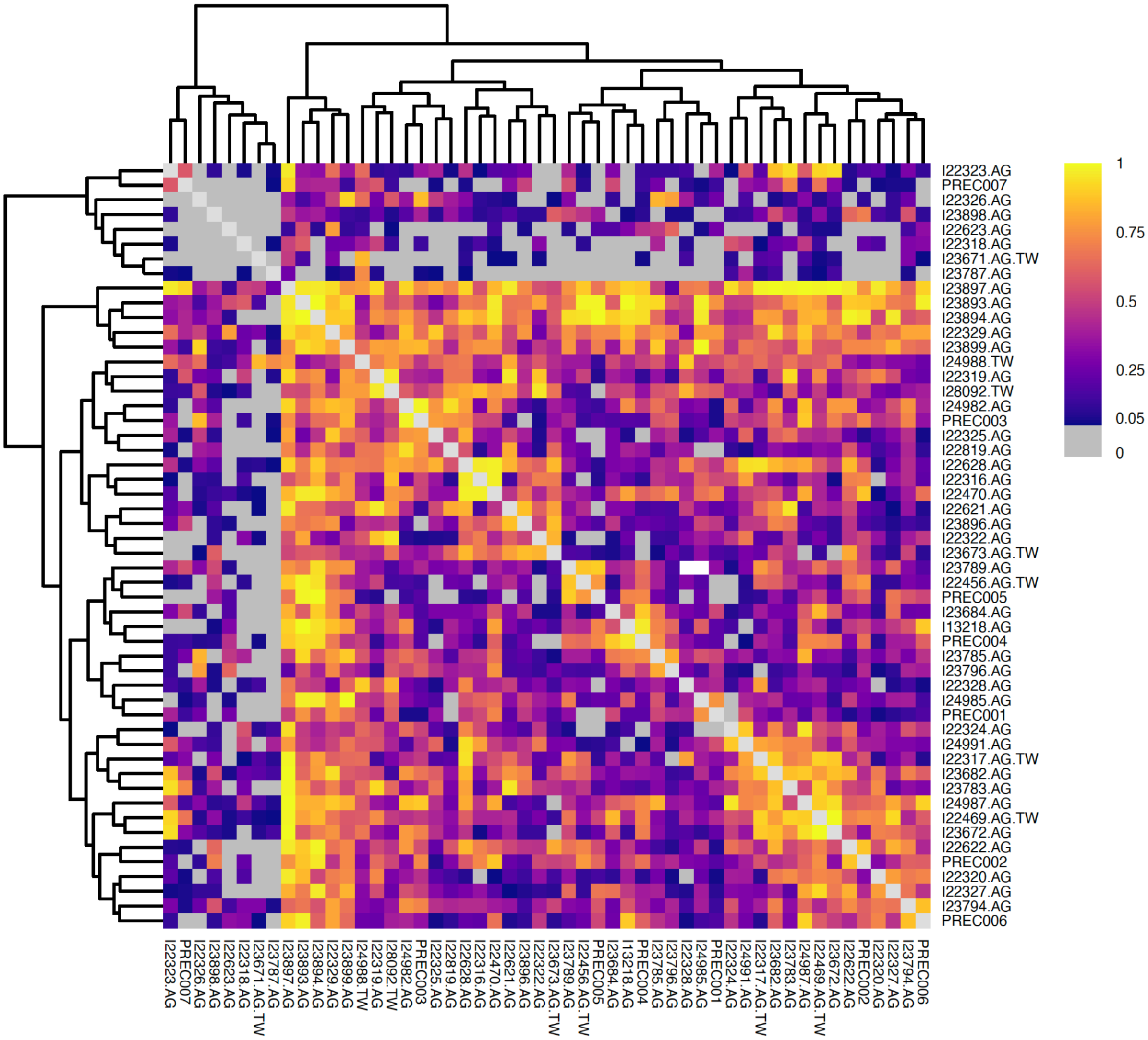


We grouped all individuals by site and time period except the eight individuals listed above and I22317, who is the lower coverage first-degree relative of another individual in our dataset. We re-ran *qpWave* with the following individuals/groups (Table S2; **Supplementary Figure 3.2**):

- Colombia_Aguazuque_MidH_HG
- Colombia_Aguazuque_InitialLateH_HG
- Colombia_Bonacá_MidH_HG (note: low resolution, <50K SNPs covered on 2M.HO dataset)
- Colombia_Checua_MidH_HG
- Colombia_Checua_6000BP[130]
- Colombia_Sueva1_EarlyMidH_HG
- Colombia_Tequendama_EarlyMidH_HG
- Colombia_Tequendama_MidH_HG
- Colombia_Ubaté_MidH_HG

- Colombia_VistaHermosa_InitialLateH_HG
- I22323
- PREC007[130]
- I22326
- I23898 (note: low resolution, <50K SNPs covered on 2M.HO dataset)
- I22623 (note: low resolution, <50K SNPs covered on 2M.HO dataset)
- I22318
- I23671 (note: low resolution, <50K SNPs covered on 2M.HO dataset)
- I23787 (note: low resolution, <50K SNPs covered on 2M.HO dataset)

**Supplementary Figure 3.2.** Symmetry matrix of *qpWave* values with each cell representing the fit (p-value) of the model. Gray cells represent p-values < 0.05. Clustering approach described above.

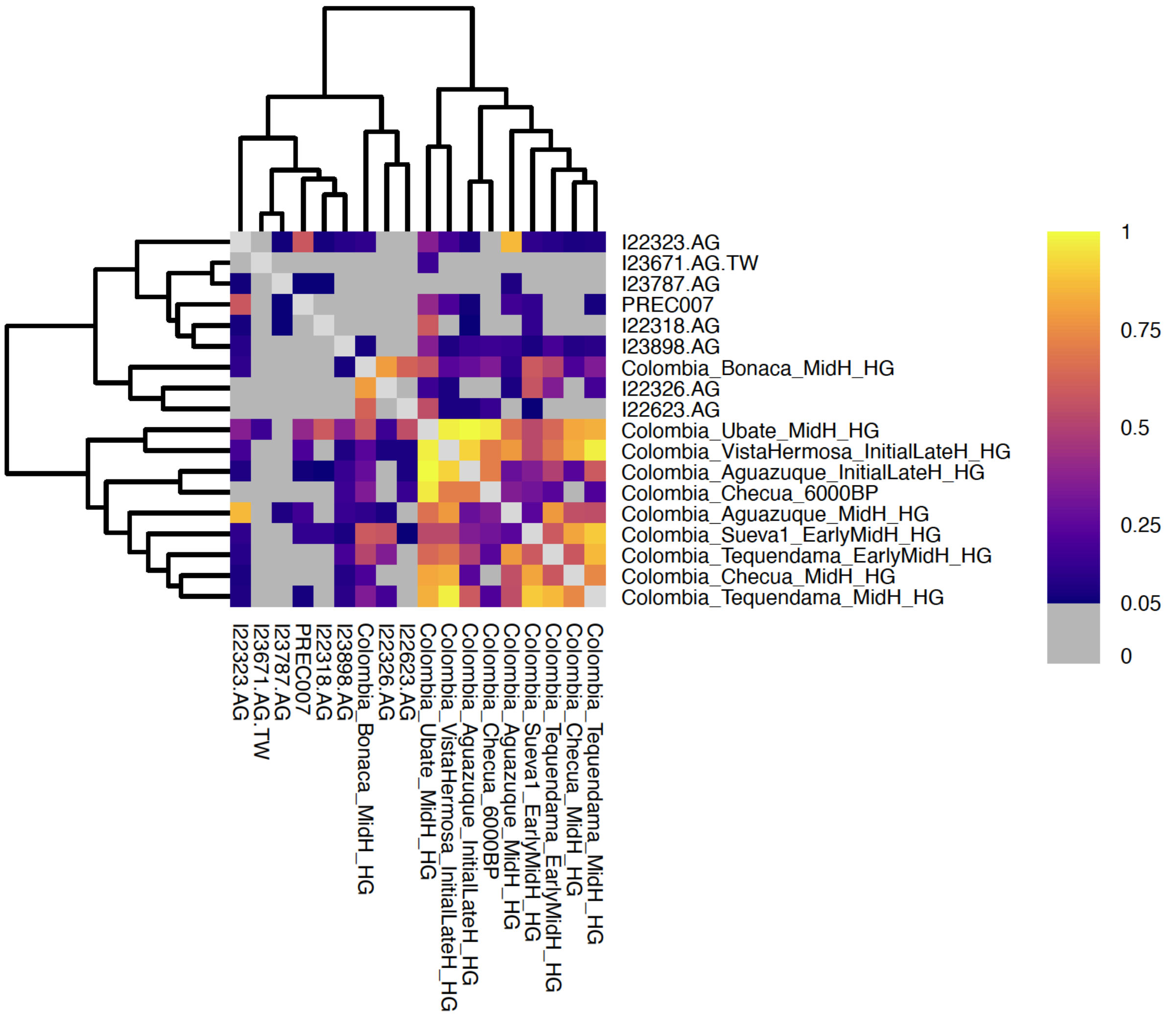

In addition to the eight individuals identified above as being genetically dissimilar to others relative to the reference set, Colombia_Bonacá_MidH_HG clusters separately from the primary group of hunter-gatherers in this round of analysis. However, we again interpret this result with caution as this is a single individual represented by fewer than 50,000 SNPs.

Finally, we merged 42 individuals from Ubaté, Vista Hermosa, Aguazuque, Sueva I, Checua, and Tequendama who form a cluster in **Supplementary Figure 3.2** as Colombia_SB_HG and ran a final round of *qpWave* to see if any of the 9 individuals identified as possibly subtly distinct in our previous analyses formed a clade with the Colombia_SB_HG group at a relaxed threshold of p>0.01 (**Supplementary Table 3.1**). We adopt a less stringent threshold here to limit our outlier classification to only those individuals who are most likely to be genetically distinct. If the p-value of the *qpWave* clade test with Colombia_SB_HG was >0.01, the individual was merged into the Colombia_SB_HG analysis group.

**Supplementary Table 3.1. *qpWave* results.** Individuals in green cells were merged into the Colombia_SB_HG analysis group based on the *qpWave* model p-value (p>0.01). Asterisk (*) indicates less than 50,000 SNPs covered on 2M.HO dataset.

| Pop1 | Pop2 | qpWave model p-value |
|---|---|---|
| Colombia_SB_HG | Colombia_Bonacá_MidH_HG* | 0.24238793 |
| Colombia_SB_HG | I22318 | 0.0073586 |
| Colombia_SB_HG | I22323 | 0.05925565 |
| Colombia_SB_HG | I22326 | 0.02114137 |
| Colombia_SB_HG | I22623* | 0.04170332 |
| Colombia_SB_HG | I23671* | 0.00459542 |
| Colombia_SB_HG | I23787* | 0.00674283 |
| Colombia_SB_HG | I23898* | 0.10928204 |
| Colombia_SB_HG | PREC007 | 0.01585982 |

Three individuals - I22318, I23671, and I23787 - did not form a clade with Colombia_SB_HG at a threshold of p>0.01, although only I22318 had coverage at more than 50,000 SNPs on the 2M.HO dataset. To investigate if I22318 might be a true ancestry outlier, we used the statistic $f_4$(Colombia_SB_HG, I22318; *qpWave_Outgroup1*, *qpWave_Outgroup2*) to explore whether poor model fit was driven by a single highly informative outgroup but could not identify a clear signal of significantly distinctive ancestry for this individual (**Table S28**). In the absence of a clear explanation for the poor model fit, we conservatively did not merge these individuals into the Colombia_SB_HG group for subsequent analyses.

**Assessing genetic homogeneity among Herrera Period and Muisca Period people**

For the 164 newly reported individuals from Herrera-Muisca Period contexts and the 14 individuals from post-2000 BP contexts reported in ref. [130], we first tested for cladality between individuals at each site. Results for all pairwise tests are summarized below with data in **Table S29**.

- **Cable Bogota** (n=11; Late Muisca): all individuals are consistent with forming a single genetic group

- **Candelaria La Nueva** (n=12; Late Muisca): all individuals are consistent with forming a single genetic group

- **El Venado** (n=9; Herrera, Early Muisca, Late Muisca): all individuals are consistent with forming a single genetic group
    - Temporal groups (Herrera-Early Muisca-Late Muisca) also form a clade and are merged as Colombia_ElVenado (**Supplementary Table 3.2**)

**Supplementary Table 3.2. Testing for cladality among temporal groups at El Venado with *qpWave*.**

| Pop1 | Pop2 | qpWave model p-value |
|---|---|---|
| Colombia_ElVenado_Herrera | Colombia_ElVenado_EarlyMuisca | 0.690232067 |
| Colombia_ElVenado_Herrera | Colombia_ElVenado_LateMuisca | 0.242158394 |
| Colombia_ElVenado_EarlyMuisca | Colombia_ElVenado_LateMuisca | 0.297118676 |

- **Hoja Caduca** (n=3; Late Muisca): all individuals are consistent with forming a single genetic group

- **LabIng2** (n=1; Late Muisca): single sample

- **La Muela** (n=10; Herrera, Late Muisca): all individuals are consistent with forming a single genetic group
    - Temporal groups (Herrera-Late Muisca) also form a clade and are merged as Colombia_LaMuela (**Supplementary Table 3.3**)

**Supplementary Table 3.3. Testing for cladality among temporal groups at La Muela with *qpWave*.**

| Pop1 | Pop2 | qpWave model p-value |
|---|---|---|
| Colombia_LaMuela_Herrera | Colombia_LaMuela_LateMuisca | 0.899135195 |

- **La Salina** (n=1; Herrera): single sample

- **Las Delicias** (n=7, with 2 published in ref. [130] and 5 newly reported in this work; Early Muisca): all individuals are consistent with forming a single genetic group

- **Laguna de la Herrera**[130] (n=9, Herrera): all individuals except FORM001 are consistent with forming a single genetic group

- We used the statistic $f_4$(Colombia_LagunadelaHerrera_2000BP, FORM001; *qpWave_Outgroup1*, *qpWave_Outgroup2*) to explore whetherthe poor model fit was driven by a single highly informative outgroup (**Table S5**). Statistics suggest that FORM001 has a stronger affinity to Wayuu than other individuals from Laguna de la Herrera.

- **Marin** (n=12; Late Muisca): all individuals are consistent with forming a single genetic group

- **Nueva Esperanza** (n=47; Herrera, Early Muisca, Late Muisca): all individuals are consistent with forming a single genetic group
  - Temporal groups (Herrera-Early Muisca-Late Muisca) also form a clade and are merged as Colombia_NuevaEsperanza (**Supplementary Table 3.4**)

**Supplementary Table 3.4. Testing for cladality among temporal groups at Nueva Esperanza with *qpWave*.**

| Pop1 | Pop2 | qpWave model p-value |
|---|---|---|
| Colombia_NuevaEsperanza_Herrera | Colombia_NuevaEsperanza_EarlyMuisca | 0.84158348 |
| Colombia_NuevaEsperanza_Herrera | Colombia_NuevaEsperanza_LateMuisca | 0.134946639 |
| Colombia_NuevaEsperanza_EarlyMuisca | Colombia_NuevaEsperanza_LateMuisca | 0.108353991 |

- **Portalegre** (n=12; Late Muisca, Muisca [unknown if Early or Late]): all individuals are consistent with forming a single genetic group
  - Temporal groups (Muisca-Late Muisca) also form a clade and are merged as Colombia_Portalegre (**Supplementary Table 3.5**)

**Supplementary Table 3.5. Testing for cladality among temporal groups at Portalegre with *qpWave*.**

| Pop1 | Pop2 | qpWave model p-value |
|---|---|---|
| Colombia_Portalegre_Muisca | Colombia_Portalegre_LateMuisca | 0.232386295 |

- **Purnia**[130] (n=2, Los Curos region): all individuals are consistent with forming a single genetic group

- **San Francisco** (n=7; Late Muisca): all individuals except I10932 are consistent with forming a single genetic group
  - We used the statistic $f_4$(Colombia_SanFrancisco, I10932; *qpWave_Outgroup1*, *qpWave_Outgroup2*) to explore whether the poor model fit was driven by a single highly informative outgroup (**Table S5**). Statistics suggest that I10932 has a weaker affinity to Inga and Cabécar than other individuals from San Francisco and stronger affinity to people living in lowland Amazonian South America.

- **Soacha**[130] (n=1, Late Muisca): single sample

- **Terregrande** (n=1; Early Muisca): single sample

- **Terreros** (n=6; Early Muisca, Late Muisca): all individuals are consistent with forming a single genetic group
    - Temporal groups (Early Muisca-Late Muisca) also form a clade and are merged as Colombia_Terreros (**Supplementary Table 3.6**)

**Supplementary Table 3.6. Testing for cladality among temporal groups at Terreros with *qpWave*.**

| **Pop1** | **Pop2** | **qpWave model p-value** |
|---|---|---|
| Colombia_Terreros_EarlyMuisca | Colombia_Terreros_LateMuisca | 0.423984704 |

- **Tibanica** (n=24; Early Muisca, Late Muisca + Elite, Non-Elite): all individuals except I22700 are consistent with forming a single genetic group (I22700 does not form a clade with any other individual at p>0.05)
    - Temporal groups (Early Muisca-Late Muisca) or groups based on status (Elite-Non-Elite) or cemetery (0-4) form a clade and are merged as Colombia_Tibanica (data are in **Table S29** due to the number of tests)
    - We used the statistic $f_4$(Colombia_Tibanica, I22700; *qpWave_Outgroup1*, *qpWave_Outgroup2*) to explore whether the poor model fit was driven by a single highly informative outgroup (**Table S5**). Statistics suggest that I22700 has less Cabécar-like ancestry and raises the possibility that there is contamination in this sample (especially given an intermediate sex ratio of 0.303, **Table S1**).

- **Zanja Eléctrica** (n=3; Late Muisca): all individuals are consistent with forming a single genetic group

We then grouped all individuals by site except five lower coverage 1st-degree relatives of other individuals in our dataset (I10929, I22495, I22492, I23276, FORM004) and the 3 distinct individuals identified with *qpWave* (FORM001, I10932, and I22700) and re-ran *qpWave* with 22 individuals/groups (**Table S30**; **Supplementary Figure 3.3**).

**Supplementary Figure 3.3.** Symmetry matrix of *qpWave* values with each cell representing the fit (p-value) of the model. Gray cells represent p-values < 0.05. Clustering approach described above.

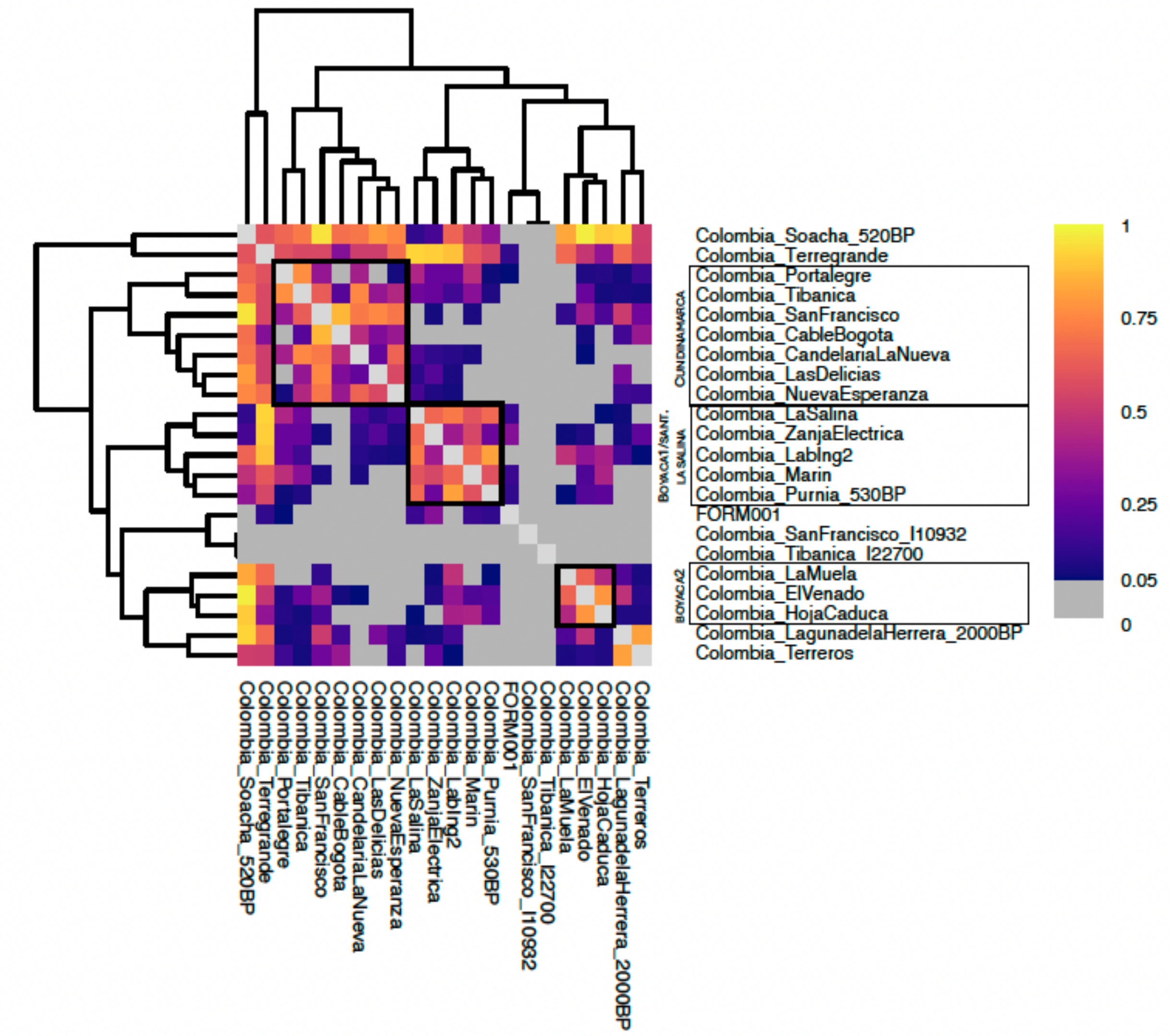


Here, we see geographic-based substructure separating sites in Cundinamarca from sites in Boyacá and Santander (an exception is La Salina; while the site is located in Cundinamarca, the single individual is clustered with individuals from Boyacá). There are two individuals – Colombia_Terregrande and Colombia_Soacha_520BP – who share affinity with individuals from both departments. There are two groups – Colombia_LagunadelaHerrera_2000BP and Colombia_Terreros – who share affinity with each other but less so with other groups in the same department. Finally, individuals I10932 and I22700 do not form a clade at $p>0.05$ with any other Herrera or Muisca Period individual or group, while FORM001 forms a clade with a small number of Herrera or Muisca Period individuals or groups, but notably does not form a clade with other individuals from Laguna de la Herrera ($p=0.003$). These groups and individuals are discussed in greater detail in **Supplementary Information 4**.

Finally, we ran *qpWave* using a final set of 26 individuals or groups and present the resulting heat map as **Figure S3**.

**Supplementary Information 4. Individuals not incorporated into broader groups**

**Hunter-gatherer and early food producer contexts**

We identified one individual from a Sabana de Bogotá hunter-gatherer context – I22318 from middle Holocene Checua – who had coverage at >50,000 SNPs on the 2M.HO dataset and was assessed as being somewhat genetically dissimilar to other people with Sabana de Bogotá hunter-gatherer-associated ancestry (Colombia_SB_HG) relative to the reference set used for *qpWave*. A key question was if we could determine the extent and nature of ancestry differences; however, in every case the statistic $f_4$(Yoruba, Colombia_SB_HG; I22318, *NativeAmerican_Test*) showed significant excess allele sharing between Colombia_SB_HG and I22318 relative to any *NativeAmerican_Test* population (**Table S12**), consistent with these individuals deriving from the same ancestral population. Non-significant statistics of the form $f_4$(Yoruba, *NativeAmerican_Test*; Colombia_SB_HG, I22318) revealed that no population splits a clade between Colombia_SB_HG and I22318 (**Table S31**). This individual was also placed in a sub-clade containing other individuals from the same site and time period in an NJ tree constructed using inverted outgroup-$f_3$ statistics (**Figure 3**). As discussed in **Supplementary Information 3**, we also used the statistic $f_4$(Colombia_SB_HG, I22318; *qpWave_Outgroup1*, *qpWave_Outgroup2*) to explore whether the lack of cladality between I22318 and Colombia_SB_HG in *qpWave* was driven by a single informative outgroup but identified no clear signal (**Table S6**). We conclude that any true ancestry differences between I22318 and other people with Sabana de Bogotá hunter-gatherer-associated ancestry fall below the limits of detection with allele frequency-based methods, precluding our ability to confidently distinguish between true ancestry differences and technical or data-quality effects.

**Herrera-Muisca contexts**

Individuals not incorporated into the three Herrera-Muisca Period regional groups (Colombia_Cundinamarca, Colombia_Boyacá1_LaSalina, and Colombia_Boyacá2) include those from Laguna de la Herrera and Terreros who show somewhat less genetic similarity to all Herrera-Muisca Period people; individuals from Terregrande and Soacha who show similar affinity to Herrera-Muisca Period people from Cundinamarca and Boyacá departments; and three individuals (I10932, I22700, and FORM001) who are dissimilar to other Herrera-Muisca Period people in *qpWave*. These exclusions reflect a conservative grouping strategy designed to avoid over-interpreting subtle differences that may arise from limited coverage, individual-level drift, or technical artifacts, rather than unambiguous evidence for ancestry differences.

**Laguna de la Herrera[130]**

The site of Laguna de la Herrera is located in the present-day Cundinamarca department of Colombia, yet *qpWave* suggests that the people who lived at this site share somewhat less affinity to people from other sites within this department. We used the statistic $f_4$(Yoruba, *NativeAmerican_Test*; Colombia_Cundinamarca, Colombia_LagunadelaHerrera_2000BP) in an attempt to understand any ancestry differences (**Tables S3** and **S4**). Only a few statistics reached statistical significance – the strongest (Z=-3.5) showing greater allele sharing between ancient

Peru_LIP_Pacapaccari_600BP or present-day Narihuala and Colombia_Cundinamarca relative to Colombia_LagunadelaHerrera_2000BP – and no ancient or modern genotyped population sharing significantly more alleles with Colombia_LagunadelaHerrera_2000BP. Considering results falling below a Z=|3| significance threshold, it appears that Colombia_Cundinamarca shares more drift with a range of ancient and modern Native American reference populations than Laguna de la Herrera. Because this pattern is not restricted to any single geographic region, it does not clearly identify a specific source of ancestry difference and could instead reflect increased drift in the Laguna de la Herrera individuals or technical factors affecting allele-sharing analyses. An alternative explanation is that the individuals from Laguna de la Herrera have some ancestry that is not well-represented among genotyped *NativeAmerican_Test* populations.

**Terreros**

The site of Terreros is also located in the Cundinamarca department, but the people we analyze here appear to share somewhat less affinity to other people from sites within this department in *qpWave*. We again used the statistic $f_4$(Yoruba, *NativeAmerican_Test*; Colombia_Cundinamarca, Colombia_Terreros) to attempt to understand any ancestry differences (**Tables S3** and **S4**). Relative to the Colombia_Cundinamarca regional group, we find a signal of asymmetric allele sharing between Colombia_Terreros and some ancient Brazilian individuals and groups, including Brazil_Moraes_5800BP, and sambaqui-associated Brazil_Limao_Sambaqui_500BP, Brazil_JabuticabeiraII_Sambaqui_2200BP as well as Caribbean_Ceramic (a population documented to have ancestry like that found in present-day Arawakan speakers from northeastern South America[131]) and an ancient individual from Chile (Chile_PuntaSantaAna_7300BP). This pattern suggests subtle ancestry heterogeneity within Cundinamarca, though the signal is relatively weak and does not point to a single external geographic source as driving the difference.

**Terregrande**

The site of Terregrande is in the present-day department Cundinamarca, but the individual we analyze from this site shares similar affinity with people from both the Cundinamarca and Boyacá regional groups in *qpWave*. Consistent with this pattern, $f_4$(Yoruba, *NativeAmerican_Test*; *Colombia_HerreraMuisca_RegionalGroup*, Colombia_Terregrande) indicates no significant ancestry differentiation between this individual and either regional group (**Tables S3** and **S4**). This result may reflect shared ancestry or limited analytical power from a single individual.

**Soacha_520BP[130]**

Soacha is in the present-day department Cundinamarca, but the individual from this site shares similar affinity with people from both the Cundinamarca and Boyacá regional groups in *qpWave*. Only a few statistics reached statistical significance – the strongest (Z=-4.2) showing greater allele sharing between Brazil_PalmierasXingu_Sambaqui_500BP as well as relatively recent ancient Peruvians and Colombia_Cundinamarca relative to Colombia_Soacha_520BP – and no ancient or modern genotyped population sharing significantly more alleles with Colombia_Soacha_520BP (**Tables S3** and **S4**). Considering results falling below a Z=|3| significance threshold, it appears that Colombia_Cundinamarca shares more drift with a range of ancient and modern Native American reference populations than Soacha_520BP. Because this pattern is not restricted to any single geographic region, it does not clearly identify a specific source of ancestry difference and could instead reflect subtle local drift in the Soacha_520BP individual or technical factors. An alternative explanation is that the Soacha_520BP individual has some ancestry that is not well-represented among genotyped *NativeAmerican_Test* populations.

**FORM001**

FORM001 is a male individual from the site of Laguna de la Herrera who is assessed as somewhat genetically dissimilar to other people from Laguna de la Herrera in *qpWave* (p-value of 0.002 for a clade test). Statistics of the form $f_4$(Yoruba, *NativeAmerican_Test*; Colombia_LagunadelaHerrera_2000BP, FORM001) provide limited insight into the differences between FORM001 and the other individuals from Laguna de la Herrera (**Table S3**), with the strongest statistic when present-day Wayuu is in the position of *NativeAmerican_Test* (Z=3.3, **Table S4**). Tests of the form $f_4$(Colombia_LagunadelaHerrera_2000BP, FORM001; *qpWave_Outgroup1*, *qpWave_Outgroup2*) similarly suggest that FORM001 shares more alleles with a population proxied by present-day Wayuu than other individuals from Laguna de la Herrera (**Table S5**). Wayuu are a present-day population who live in the Guajira Peninsula of northern Colombia and Venezuela, genetically associated with northern Amazonian lowland-related and Caribbean-related South Americans; increased affinity to Wayuu in this context is most consistent with subtle within-site ancestry heterogeneity rather than strong evidence for direct Wayuu-related gene flow.

**I10932**

I10932 – a male individual from San Francisco (in the department of Cundinamarca) who we directly date to 1422-1454 calCE (465±20 BP, PSUAMS-5492) – is a PCA outlier (**Figure 2**), shifted toward the H-G cluster and South Americans and away from Chibchan-related groups relative to other Herrera-Muisca Period people. This individual also appears as genetically distinct in *qpWave*, forming a clade at a p>0.05 threshold with no other individuals or groups from the Altiplano Cundiboyacense (**Figure S3**). Additionally, an NJ tree based on inverted outgroup-$f_3$ statistics (**Figure 3**) places I10932 basal to all Herrera-Muisca Period groups. $f_4$-statistics exceeding our significance threshold involve Chibchan-associated references (ancient Panama_IsthmoColumbian and present-day Cabécar), in the direction of greater allele sharing with Colombia_Cundinamarca than with I10932 (**Tables S3** and **S4**) or with the other individuals from San Francisco relative to I10932 (**Table S5**). Across other ancient and present-day Native American references, $f_4$ values are not statistically significant, suggesting that differentiation is primarily driven by reduced affinity to Chibchan-related populations. We tested whether I10932 might have retained ancestry from people with Sabana de Bogotá hunter-gatherer-associated ancestry; however, the statistic $f_4$(Yoruba, Colombia_SB_HG; I10932, *Colombia_HerreraMuiscaPeriodGroup*) is consistent with zero, suggesting no excess hunter-gatherer ancestry in I10932 relative to other Herrera-Muisca Period people (**Table S12**). We interpret I10932 as being modestly differentiated from other Herrera-Muisca Period people from the site of San Francisco as well as other sites in Cundinamarca.

**I22700**

I22700, a non-elite individual attributed to Group 3 from the site of Tibanica in Cundinamarca, is of relatively low coverage (~90,000 SNPs covered on the 2M dataset and ~32,000 SNPs covered on the 2M.HO dataset) and does not appear as a PCA outlier but appears genetically dissimilar from all other Herrera-Muisca Period people in *qpWave*. With tests of the form $f_4$(Yoruba, *NativeAmerican_Test*; Colombia_Cundinamarca, I22700) we obtain consistently negative statistics indicating that I22700 shares less drift with essentially all Native American reference populations relative to Colombia_Cundinamarca (**Tables S3** and **S4**). This likely reflects contamination which is also supported by an intermediate sex ratio of 0.303 (there are too few SNPs for contamination analysis using ANGSD or hapConX) (**Table S1**).

**Supplementary Information 5 – Families**

In the text below, "d" represents "degree" (e.g., 2d = 2$^{nd}$ degree). Genetic relatedness was inferred using the method described in ref. [132].

__Hunter-gatherer/subsistence transition contexts__

We identify three groups of closely-related individuals ("families") from pre-Herrera Period hunter-gatherer/subsistence transition contexts. All are from the site of Checua.

- **Checua Family A** (5 members): I22317-I22622 are son-father; I22319, I22622 have a 2d relationship; I22317-I22319 are father-daughter; I22319, I22328 have a 2d relationship; I22319, I22323 have a 2d relationship; I22317-I22323 are son-mother. Pedigree shown as **Supplementary Figure 5.1**.

**Supplementary Figure 5.1**. Pedigree for Checua Family A.

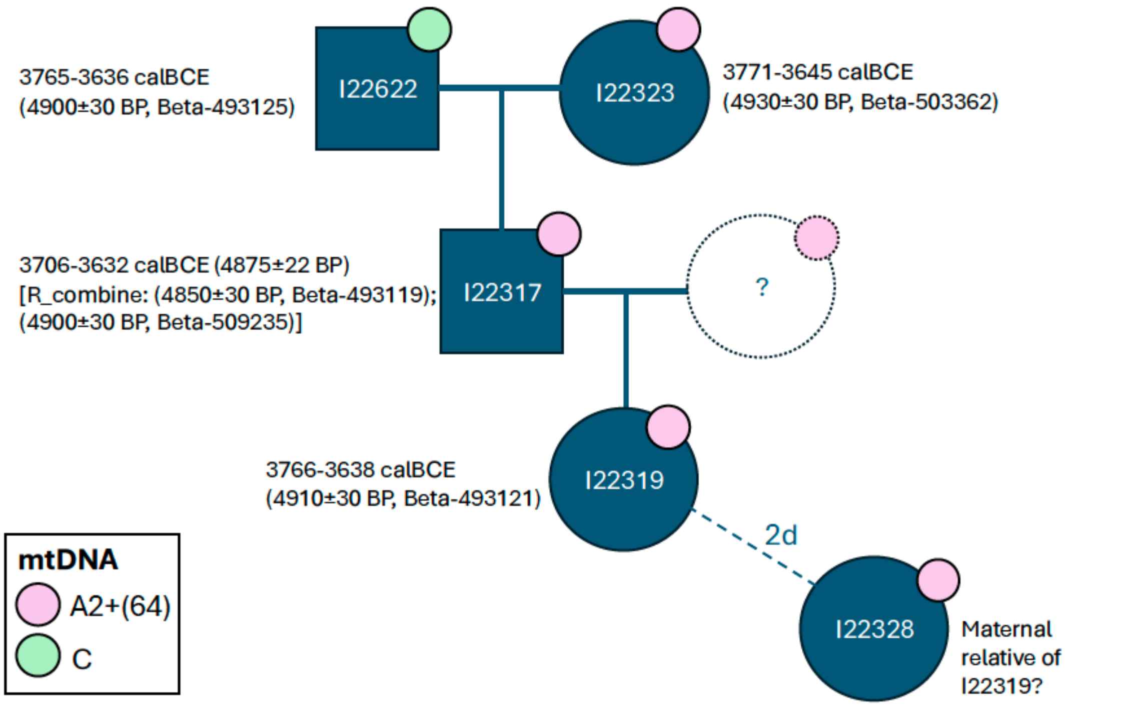


- **Checua Family B** (3 members): I22320, I22329 have a 2d relationship; I22321 (excluded from population genetics analysis following quality control checks for authenticity), I22329 have a 2d relationship

- **Checua Family C** (2 members): I22322, I22621 have a 3d relationship

**Herrera-Muisca Period contexts**

We identify 15 groups of closely-related individuals ("families") from Herrera-Muisca Period contexts.

- **Cable Bogotá Family A** (2 members): I10928-I10929 are sisters
- **Candelaria La Nueva Family A** (2 members): I22457, I22460 have a 2d/3d relationship
- **La Muela Family A** (3 members): I22483, I22632 have a 2d relationship; I22484, I22483 have a 2d/3d relationship
- **Nueva Esperanza Family A** (18 members): I23729-I23720, I22710-I25423, I22714-I23720, I23720-I23730 have a 2d relationship; I23729-I22695, I23729-I22710, I23729-I22714, I23729-I23730, I22695-I23730, I22710-I22638, I22710-I23720, I22710-I23733, I22710-I23735, I22710-I22714, I22710-I22715, I22710-I23730, I22710-I23970, I22710-I24390, I22714-I23730, I22714-I25423, I23720-I23733, I23720-I25423, I23720-I24390, I23720-I25418, I23730-I23733, I23730-I25418, I22638-I22715, I23733-I22712, I23733-I25423, I23735-I25423, I25423-I22715, I25423-I24390, I22715-I22637, I24390-I22704, I24390-I22709 have a 2d/3d relationship. Note that this family spans the Early and Late Muisca Periods.
- **Nueva Esperanza Family B** (4 members): I23726-I23722 are siblings; I23722-I23723 have a 2d relationship; I23726-I23723, I23726-I24389, I23722-I24389 have a 2d/3d relationship
- **Nueva Esperanza Family C** (2 members): I23727-I23737 have a 2d/3d relationship
- **Portalegre Family A** (3 members): I22813-I22816 have a 2d relationship;I22813-I22818, I22816-I22818 have a 2d/3d relationship
- **Portalegre Family B** (2 members): I22814-I22811 have a 2d/3d relationship
- **San Francisco Family A** (2 members): I11772-I10931 have a 2d/3d relationship
- **Terreros Family A** (2 members): I11227-I11228 (excluded from population genetics analysis following QC) have a 1d relationship
- **Terreros Family B** (2 members): I11233 (excluded from population genetics analysis following QC)-I10934 have a 2d/3d relationship
- **Tibanica Family A** (3 members): I22698-I22699, I22699-I22700 have a 2d/3d relationship

- **Zanja Eléctrica Family A** (2 members): I22820-I22821 have a 2d/3d relationship

- **Cross-site Family A** (7 members): I22812 (Portalegre, M) - I22810 (Portalegre, F) have a 2d relationship; I22465 (Candelaria La Nueva, F) - I22701 (Tibanica, M), I22701 (Tibanica, M) - I24397 (Tibanica, M), I22701 (Tibanica, M) - I24400 (Tibanica, F), I22701 (Tibanica, M) - I22812 (Portalegre, M), I22701 (Tibanica, M) - I23739 (Tibanica, F), I24397 (Tibanica, M) - I24400 (Tibanica, F) have a 2d/3d relationship. Note that the cross-site relationships seem to be mediated by I22701 (Male) who is directly dated 1265-1287 calCE (740±15 BP, PSUAMS-12579); this individual has a relative at both Candelaria La Nueva and Portalegre.

- **Marín Family A** (11 members): I22495-I22497 are likely brothers (alternative = father-son); I22492-I22491 are daughter-mother; I22493-I22495, I22493-I22496, I22493-I22497, I22493-I22804, I22494-I22802, I22495-I22496, I22495-I22804, I22496-I22497, I22496-I22804, I22497-I22804 have a 2d relationship; I22493-I22494, I22493-I22802, I22493-I22805, I22494-I22497, I22494-I22804, I22495-I22805, I22496-I22492, I22496-I22805, I22497-I22801, I22497-I22802, I22497-I22805, I22804-I22491, I22804-I22492, I22804-I22805 have a 2d/3d relationship

**Supplementary Figure 5.2. Marín family A pedigree version A.** In this configuration, I22495 and I22497 are father-son (respectively).

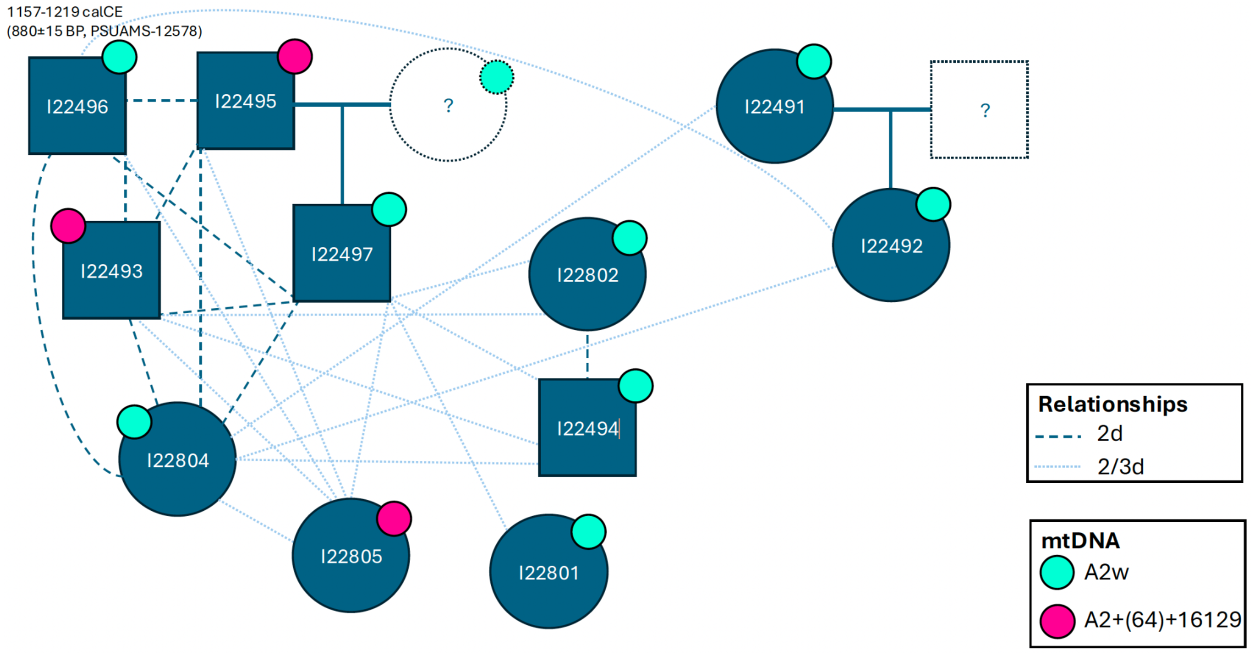

**Supplementary Figure 5.3. Marín family A pedigree version B.** In this configuration, I22495 and I22497 are full siblings.

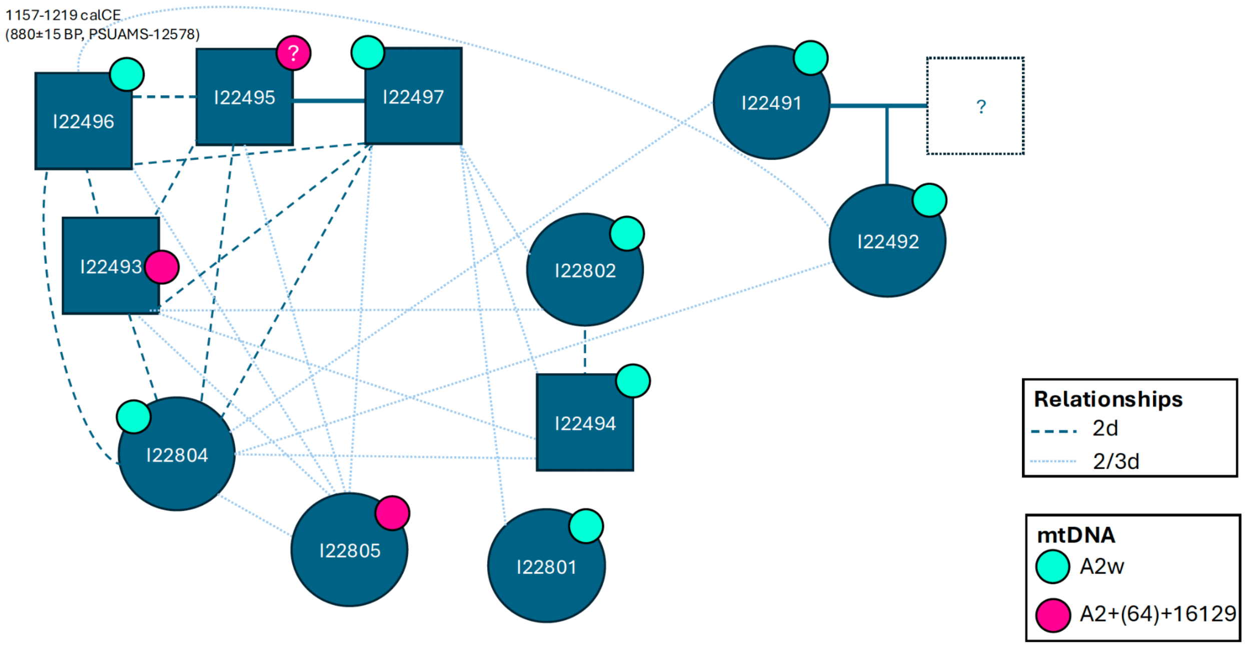


Within the 11-member family at Marín, we identified a 1st-degree relative pair, I22495 and I22497, using a previously described method[132]. Both individuals were adult males buried among the Terrace 3 elite group who exhibited cranial deformation, a symbol of elite status in Muisca society. While initially assessed as father-son (I22495 and I22497, respectively) (**Supplementary Figure 5.2**), the finding of a father and son buried in close proximity is unexpected given evidence for a system of matrilineal kinship and avunculocal residence in Muisca society[45,75], particularly in elite lineages. For this reason, we examined the genetic details of their proposed relationship in greater detail, specifically asking if these individuals could be brothers instead (**Supplementary Figure 5.3**).

We first created a table of the relationships between seven other individuals at Marín and I22495 and I22497. We summarize these relationships in **Supplementary Table 5.1**. The 2nd-degree relationships that each individual shares with I22496, I22493, and I22804 support an interpretation of I22495 and I22497 belonging to the same nuclear family. This would be consistent with an interpretation that they are full siblings (brothers) although we cannot exclude the possibility that they are father-son based on these more distant relationships alone.

**Supplementary Table 5.1. Table of relationships of 7 individuals from Marín with I22495 and I22497.** Cells are colored teal when the other Marín individual shares the same degree of relationship with both I22495 and I22497.

| Other Marín individual | Relationship with I22495 | Relationship with I22497 |
|---|---|---|
| I22496 | 2d | 2d |

| | | |
|---|---|---|
| I22493 | 2d | 2d |
| I22804 | 2d | 2d |
| I22802 | - | 2d/3d |
| I22494 | - | 2d/3d |
| I22801 | - | 2d/3d |
| I22805 | 2d/3d | 2d/3d |

Next, we looked at mitochondrial DNA (mtDNA) haplogroups, which are passed from mother to offspring (meaning that we would expect brothers to share the same mutations and the same mtDNA haplogroup). The mitochondrial genomes of I22495 and I22497 were covered at an average of 75.7x and 90.1x, respectively. Using *haplogrep3*[133], we identified the top 3 haplogroup calls for each individual and assigned a quality score to each call (**Supplementary Table 5.2**).

**Supplementary Table 5.2.** Haplogroup calls for I22495 and I22497.

| ID | haplogroup | rank | Quality score |
|---|---|---|---|
| I22495 | A2+16129! | 1 | 0.9253 |
| I22495 | A2+(64T) | 2 | 0.9188 |
| I22495 | A2+153! | 3 | 0.9188 |
| I22497 | A2w | 1 | 0.9575 |
| I22497 | A2w1 | 2 | 0.9457 |
| I22497 | A2+(64T) | 3 | 0.9396 |

Both individuals share a nearly identical pattern of mutations, consistent with belonging to the A2 lineage. A notable difference is the back mutation at 16129 (16129G!) in I22495, while the mutation is present as 16129G in I22497. However, we note that I22495 has a number of missing sites (denoted with a number followed by 'N') in regions used by PhyloTree[134] to define haplogroup A2w. Therefore, we cannot reject the possibility that I22497 belongs to A2w. Conservatively, we cannot rule out the possibility that these individuals share a haplogroup.

We then analyzed shared genomic segments on the X chromosome with *hapROH*[135] and found that I22495 and I22497 share a single segment that is 6.7 cM long. Both individuals inherit a single X chromosome exclusively from their mother, and they can either inherit the same maternal X (in which case they would share the full X chromosome) or the opposite maternal X (in which case no IBD X would be expected). Father-son pairs deterministically share no IBD X. The observed 6.7 cM of shared IBD on the X chromosome is small compared to the expected ~180 cM for a full shared X chromosome, and is thereby inconsistent with a full sibling pair that inherited the same maternal X.

However, the existence of such a segment would not be expected under a strict father–son model or model of full siblings who inherited opposite maternal X chromosomes.

It is possible that the shared segment arises from background genetic relatedness such as could have occurred generations in the past. Regardless, this result does not definitively distinguish between a scenario of full siblings or a father-son dyad for I22495 and I22497.

While we cannot robustly support a father-son or full sibling relationship for I22495 and I22497 using our SNP data alone, the genetic evidence considered alongside archaeological evidence for a general absence of father-adult son pairs in burial contexts resulting from a system of matrilineality and avunculocal residence in Muisca society lends greater support for the interpretation of these two individuals as being full siblings whose parents also shared a common ancestor at some point in the past.

**Supplementary Information 6. *qpGraph***

We first refit the informative portion of the admixture graph published in ref. [130] which shows individuals from the site of Checua dating ~6000 BP (we use Colombia_SB_HG in this work) as a distinct lineage branching from the main lineage ancestral to most South Americans that spread rapidly into and throughout South America (**Supplementary Figure 6.1**).

**Supplementary Figure 6.1. Admixture graph co-modeling major American genetic lineages.** The log-likelihood score (a weighted average of f-statistic residuals) for this graph was 17.0, and the maximum Z-score for the difference between the observed and predicted f-statistics was 2.6.

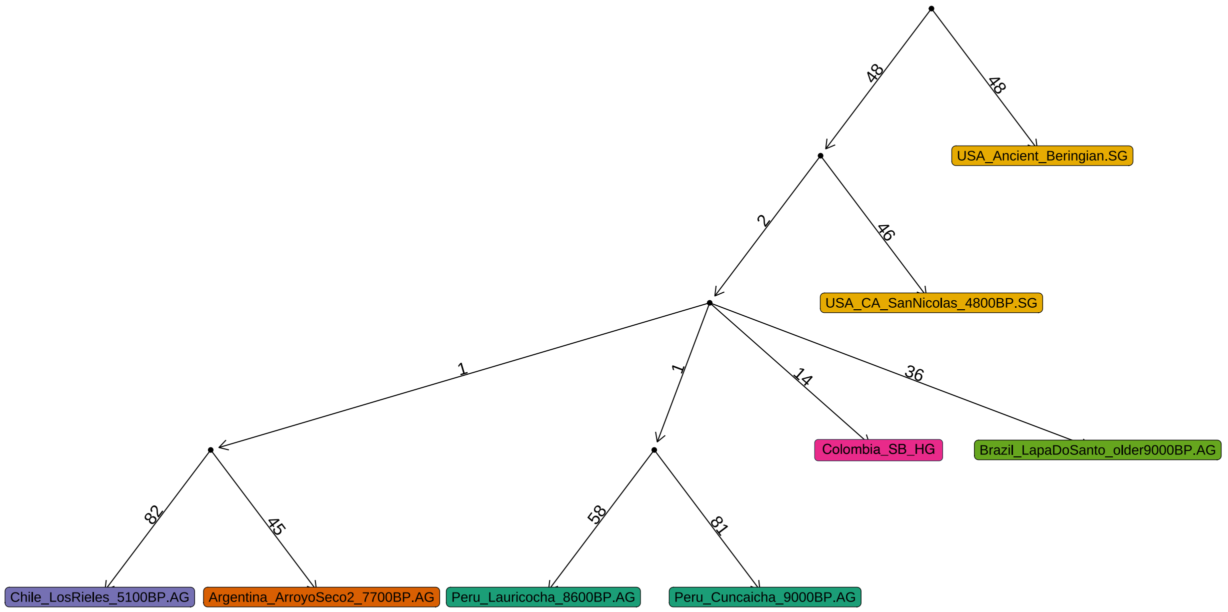


We then added Colombia_Cundinamarca to the graph as a group to represent the Herrera-Muisca Period people of the Sabana de Bogotá (we also removed USA_CA_SanNicolas_4800BP.SG based on *f*-statistics analysis showing no Channel Islands-related ancestry in our samples). Co-modeled with other American lineages, Colombia_Cundinamarca fit as a separate lineage branching from the main radiation event (**Supplementary Figure 6.2**).

**Supplementary Figure 6.2. Admixture graph co-modeling major American genetic lineages, including Colombia_SB_HG and Colombia_Cundinamarca.** The log-likelihood score for this graph was 22.4 and the maximum Z-score for the difference between the observed and predicted f-statistics was 2.6.

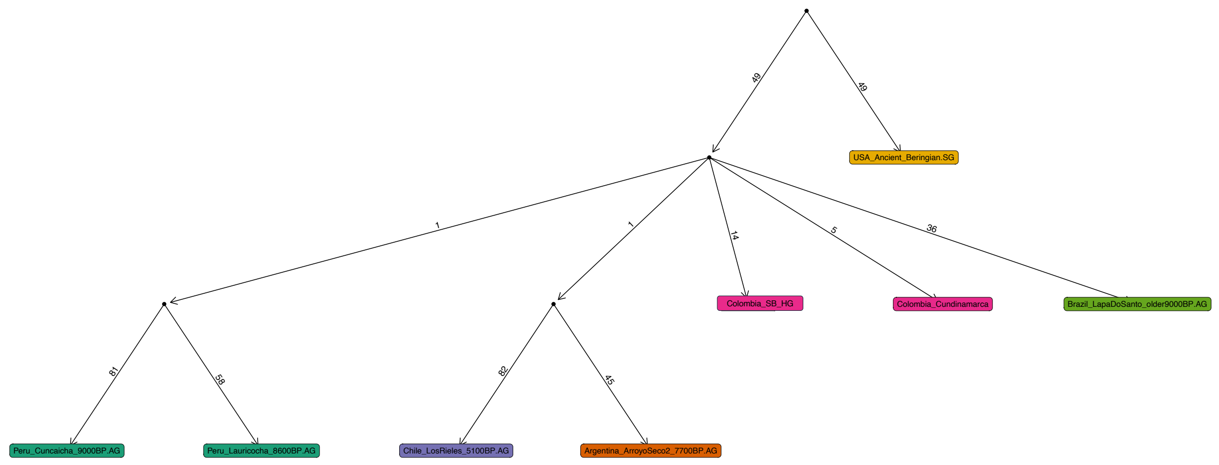


Based on $f_4$-statistics, we wondered if Colombia_Cundinamarca would fit in an admixture graph as a mix of Chibchan-related ancient Panamanians and another lineage. We added ancient Panamanians to the graph and found that a model with zero or one admixture events did not fit the data (Z>4.6 and 3.8, respectively). Graphs that included two admixture events were a much better fit for the data.

The fitting graph that we show here fits Colombia_Cundinamarca as having a minority proportion of ancestry from a node that gives rise to ancient Panamanians, with the remainder coming from a lineage branching off of the main radiation that is not represented by any other population in the graph (**Supplementary Figure 6.3**). An alternative well-fitting model had Colombia_Cundinamarca receiving a smaller proportion of ancestry from a source ancestral to ancient Panamanians (**Supplementary Figure 6.4**).

**Supplementary Figure 6.3. Admixture graph co-modeling major American genetic lineages, including Colombia_SB_HG and Colombia_Cundinamarca.** The log-likelihood score for this graph was 27.9 and the maximum Z-score for the difference between the observed and predicted f-statistics was 2.7.

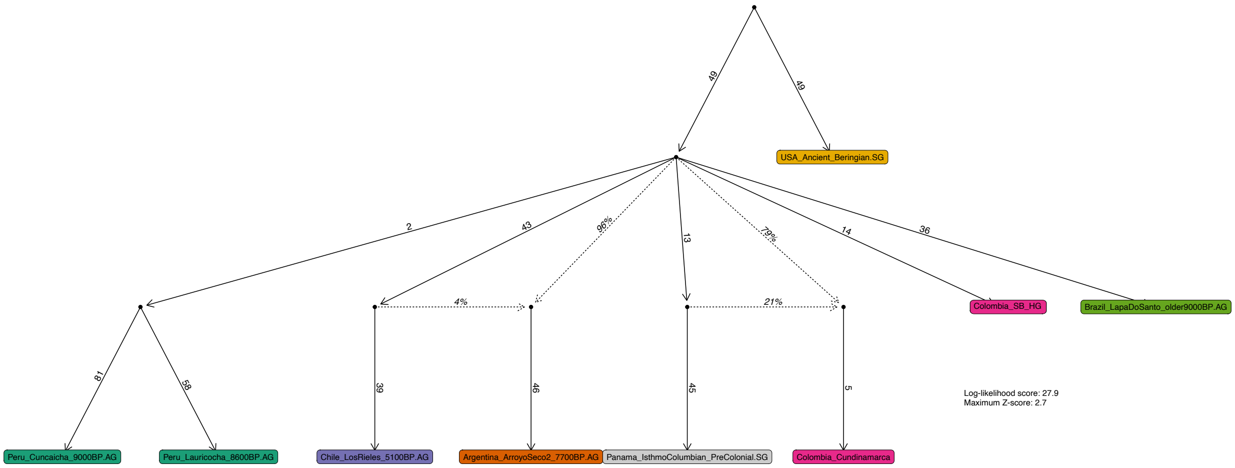


**Supplementary Figure 6.4. Admixture graph co-modeling major American genetic lineages, including Colombia_SB_HG and Colombia_Cundinamarca.** The log-likelihood score for this graph was 27.3 and the maximum Z-score for the difference between the observed and predicted f-statistics was 2.7.

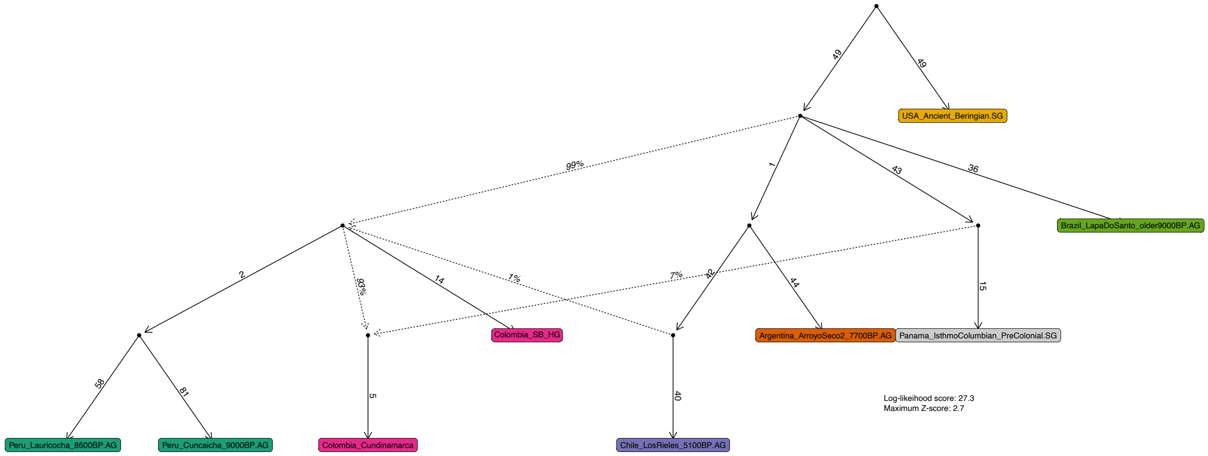


Additional alternative fitting models have gene flow in the opposite direction, with the node leading to ancient Panama receiving majority ancestry from the node leading to Colombia_Cundinamarca (**Supplementary Figure 6.5**).

**Supplementary Figure 6.5. Admixture graph co-modeling major American genetic lineages, including Colombia_Cundinamarca.** The log-likelihood score for this graph was 28.5 and the maximum Z-score for the difference between the observed and predicted f-statistics was 2.5.

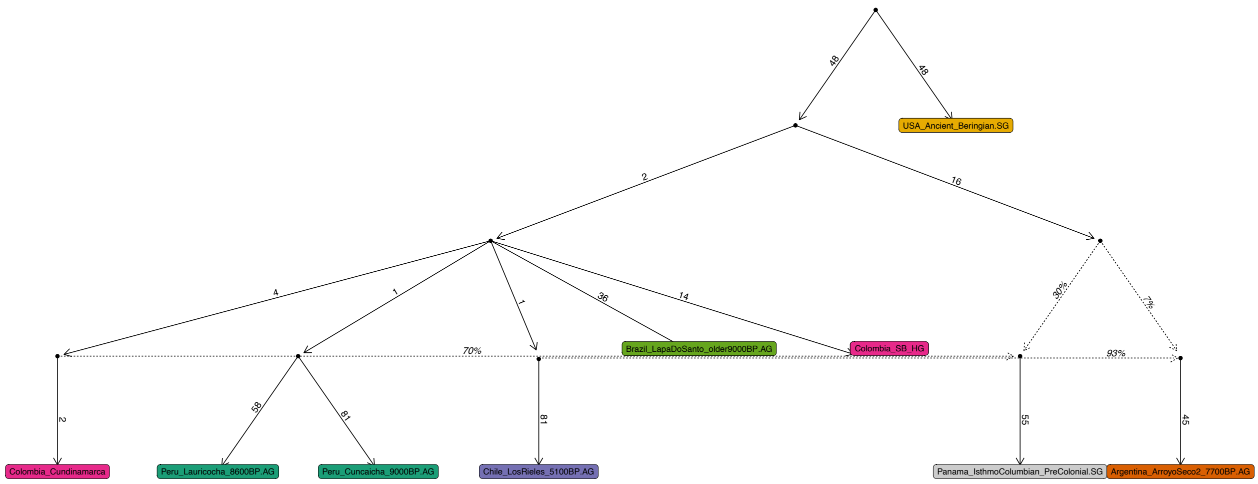

48
48
USA_Ancient_Beringian.SG
2
16
4
1
1
36
14
30%
7%
70%
93%
Brazil_LapaDoSanto_older9000BP.AG
Colombia_SB_HG
2
58
81
81
55
45
Colombia_Cundinamarca
Peru_Lauricocha_8600BP.AG
Peru_Cuncaicha_9000BP.AG
Chile_LosRieles_5100BP.AG
Panama_IsthmoColumbian_PreColonial.SG
Argentina_ArroyoSeco2_7700BP.AG

#### Supplementary Information 7 - *qpAdm*

We carried out ancestry modeling using *qpAdm*. Our primary focus was modeling the ancestry of the three regional groups of the Altiplano Cundiboyacense (Colombia_Cundinamarca, Colombia_Boyacá1_LaSalina, and Colombia_Boyacá2), but we also attempted to model the ancestry of six other individuals/groups (Colombia_Soacha_520BP, Colombia_Terregrande, Colombia_LagunadelaHerrera_2000BP, Colombia_Terreros, I10932, FORM001) who were not incorporated into the regional groups (**Supplementary Information 4**), although smaller sample sizes and lower coverage reduce our ability to reject models (we exclude I22700 from modeling due to evidence of contamination). We used the 2M.HO dataset as it allows for a "right" reference set of a consistent data type that is meaningful for differentiating ancestries in the Americas. Unless stated otherwise, we used the same base reference set as for *qpWave*: Chipewyan, Zapotec, Mixe, Mixtec, Surui, Cabécar, Piapoco, Karitiana, Inga, Wayyuu, Apalai, and Arara. We used the options 'basepop: Han.DG', 'allsnps: YES', and 'inbreed: NO'.

##### One-source models

We started by examining the fit of models with one ancestry source using the base reference set (**Supplementary Figure 7.1**). If there was population continuity between pre-Herrera times and the onset of the Herrera-Muisca Periods, pre-Herrera people (Colombia_SB_HG) would fit as a single source for the ancestry of the Herrera-Muisca Period people. Alternatively, if a Chibchan-related population, proxied here by ancient Panamanian individuals (Panama_IsthmoColumbian_PreColonial) or present-day Cabécar, migrated to the Altiplano Cundiboyacense and gave rise to Herrera and Muisca Period groups without subsequent gene flow, one of these proxies would be sufficient as a single ancestry source in our models.

We found that no ancient or present-day American population fits as a single ancestry source for the three Altiplano Cundiboyacense regional groups relative to the reference set ($p<0.05$ for each model).

**Supplementary Figure 7.1.** Matrix of results for one-source ancestry models for Herrera-Muisca Period targets with the base reference set. Shaded cells represent "fitting" models with p-values >0.05 where ancestry proportions are between 0-1 for both sources, standard errors are between 0-1, and ancestry proportions ± 1 standard error do not go outside of the 0-1 range. In other words, these are models that are consistent with the data.

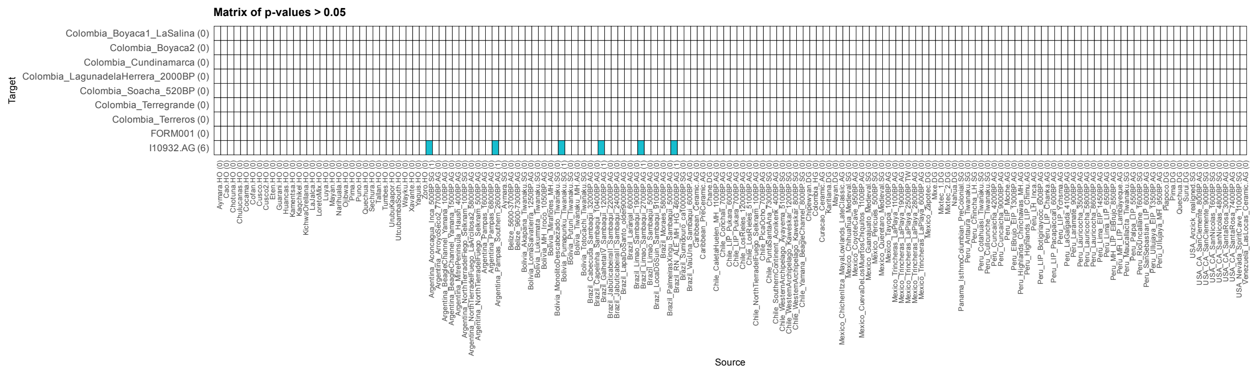


Five of the six groups/individuals that were not incorporated into the three regional groups also had no fitting one-source models; however, there were some fitting one-source models for I10932 (covered at ~380K SNPs on the 2M.HO array), who formed a clade with genetically diverse ancient individuals from Argentina, Bolivia, and Brazil. All fitting sources comprise single individuals of varying coverage, and therefore we caution that our ability to reject models is relatively poor. An interesting result is the one-source model fitting I10932 as a clade with Argentina_Aconcagua_Inca_500BP ($p=0.08$), an individual found in the southern Andes but most closely related to populations from the North Peru coast where previous work has detected a signal of Amazonian-related ancestry relative to the neighboring highlands groups[136].

We removed Cabécar from the reference set in order to 1) test it as a source, and 2) retest ancient Panamanians as a single source without a genetically similar population in the reference set. Neither Cabécar nor ancient Panamanians fit as a single source for any target ($p<<0.05$ for all models). This suggests that multiple sources are needed to explain the ancestry of the agriculturalists of the Altiplano Cundiboyacense.

##### Two-source models

One-source models were insufficient for explaining the ancestry of the regional groups, so we proceeded to test two-source models. We first set Colombia_SB_HG as Source 1 and rotated through all other genotyped ancient and modern American populations as Source 2 to test the hypothesis that an incoming group mixed with the local hunter-gatherer/early food producing population of the Sabana de Bogotá to give rise to the later Herrera and Muisca Period people of the Altiplano Cundiboyacense (**Supplementary Figure 7.2)**. We found no fitting models for three regional groups. These results are consistent with other analyses suggesting negligible hunter-gatherer ancestry persisted into the Herrera and Muisca Periods. This is consistent with the incoming population largely or fully replacing the autochthonous population.

**Supplementary Figure 7.2.** Matrix of results for two-source ancestry models for Herrera-Muisca Period targets with Colombia_SB_HG set as Source 1. Shaded cells represent models with p-values >0.05 where ancestry proportions are between 0-1 for both sources, standard errors are between 0-1, and ancestry proportions ± 1 standard error do not go outside of the 0-1 range. In other words, these are models that are consistent with the data.

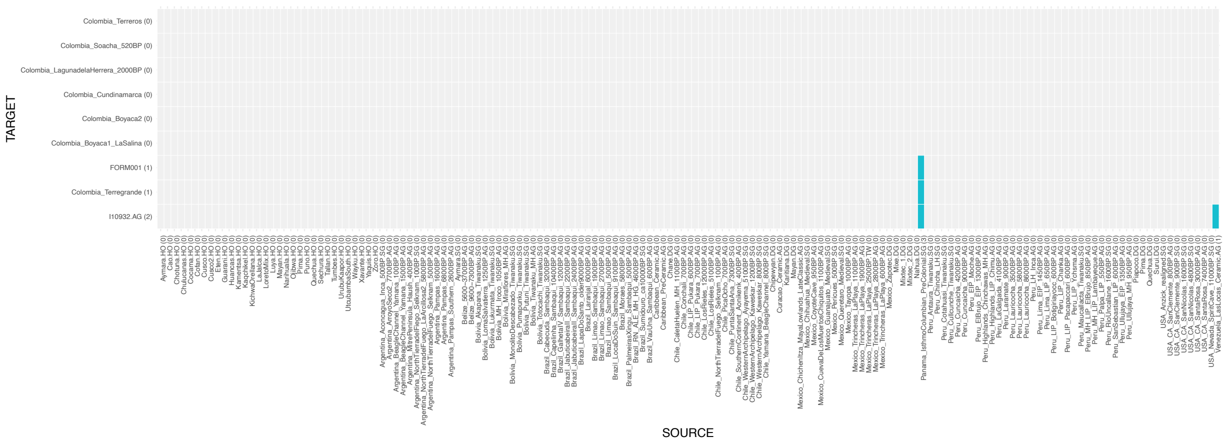


We cannot robustly reject every two-source model for the single individuals FORM001, I10932, and Colombia_Terregrande. A model for FORM001 (covered at ~400K SNPs on the 2M.HO dataset) as 35.9%±5.2% Colombia_SB_HG and 64.1%±5.2% Panama_IsthmoColumbian_PreColonial fits the data in *qpAdm* (p=0.17), however, this is inconsistent with $f_4$-statistics of the form $f_4$(Yoruba, Colombia_SB_HG; FORM001, *Colombia_OtherHerreraMuisca*), where all tests are equivalent to zero, suggesting that FORM001 does not share significantly more alleles with Colombia_SB_HG than other Herrera-Muisca groups do (we would expect these tests to be significantly negative if FORM001 had ancestry from Colombia_SB_HG that was not found in the other Herrera-Muisca Period groups) (**Supplementary Table 7.1**).

**Supplementary Table 7.1.** $f_4$-statistics of the form $f_4$(Yoruba, Colombia_SB_HG; *Test*, *Colombia_OtherHerreraMuisca*).

| A | B | C | D | f4 | std err | Z | SNPs |
|---|---|---|---|---|---|---|---|
| Yoruba | Colombia_SB_HG | FORM001 | Colombia_Soacha_520BP | -0.000213 | -0.000427 | -0.5 | 586709 |
| Yoruba | Colombia_SB_HG | FORM001 | Colombia_LagunadelaHerrera_2000BP | -0.000106 | -0.000336 | -0.316 | 662896 |
| Yoruba | Colombia_SB_HG | FORM001 | Colombia_Terregrande | 0.000102 | -0.000578 | 0.177 | 132157 |
| Yoruba | Colombia_SB_HG | FORM001 | Colombia_Cundinamarca | 0.000074 | -0.000313 | 0.237 | 674011 |
| Yoruba | Colombia_SB_HG | FORM001 | Colombia_Boyacá1_LaSalina | 0.000098 | -0.000323 | 0.302 | 672251 |
| Yoruba | Colombia_SB_HG | FORM001 | Colombia_Boyacá2 | 0.000217 | -0.000321 | 0.676 | 672913 |
| Yoruba | Colombia_SB_HG | FORM001 | Colombia_Terreros | 0.000378 | -0.000341 | 1.106 | 638138 |
| Yoruba | Colombia_SB_HG | FORM001 | I10932.AG | 0.00014 | 0.00046 | 0.305 | 547359 |

The same is true for Colombia_Terregrande individual (covered at ~86K SNPs on the 2M.HO array), which can be fit as having 44.8%±6.2% Colombia_SB_HG-related ancestry and 55.2%±6.2% Panama_IsthmoColumbian_PreColonial-related ancestry (p=0.09), again inconsistent with $f_4$-statistics (**Supplementary Table 7.2**).

**Supplementary Table 7.2.** $f_4$-statistics of the form $f_4$(Yoruba, Colombia_SB_HG; *Test*, *Other_Colombia_HerreraMuisca*).

| A | B | C | D | f4 | std err | Z | SNPs |
|---|---|---|---|---|---|---|---|
| Yoruba | Colombia_SB_HG | Colombia_Terregrande | Colombia_Terreros | 0.00074 | 0.000441 | 1.679 | 157482 |
| Yoruba | Colombia_SB_HG | Colombia_Terregrande | I10932.AG | -0.000091 | 0.000581 | -0.157 | 140109 |
| Yoruba | Colombia_SB_HG | Colombia_Terregrande | FORM001 | -0.000102 | 0.000578 | -0.177 | 132157 |
| Yoruba | Colombia_SB_HG | Colombia_Terregrande | Colombia_Soacha_520BP | -0.000133 | -0.000544 | -0.244 | 146151 |
| Yoruba | Colombia_SB_HG | Colombia_Terregrande | Colombia_LagunadelaHerrera_2000BP | 0.000345 | -0.000428 | 0.806 | 161506 |
| Yoruba | Colombia_SB_HG | Colombia_Terregrande | Colombia_Cundinamarca | 0.000476 | -0.000377 | 1.264 | 163764 |
| Yoruba | Colombia_SB_HG | Colombia_Terregrande | Colombia_Boyacá1_LaSalina | 0.000592 | -0.000397 | 1.492 | 163441 |
| Yoruba | Colombia_SB_HG | Colombia_Terregrande | Colombia_Boyacá2 | 0.00067 | -0.00039 | 1.717 | 163584 |

Likewise, I10932 (covered at ~380K SNPs on the 2M.HO array) can be fit as having either 71.1%±4.9% Colombia_SB_HG-related ancestry and 28.9%±4.9% Panama_IsthmoColumbian_PreColonial-related ancestry (p=0.05) or 28.4%±12.6% Colombia_SB_HG-related ancestry and 71.6%±12.6% Venezuela_LasLocas_Ceramic-related ancestry (p=0.15), but also is not supported in sharing significantly more alleles with Colombia_SB_HG than other Herrera and Muisca Period people in $f_4$-statistics (**Supplementary Table 7.3**).

**Supplementary Table 7.3.** $f_4$-statistics of the form $f_4$(Yoruba, Colombia_SB_HG; *Test*, *Other_Colombia_HerreraMuisca*).

| A | B | C | D | f4 | std err | Z | SNPs |
|---|---|---|---|---|---|---|---|
| Yoruba | Colombia_SB_HG | I10932 | Colombia_Soacha_520BP | -0.000336 | -0.00043 | -0.782 | 619719 |
| Yoruba | Colombia_SB_HG | I10932 | Colombia_LagunadelaHerrera_2000BP | -0.000166 | -0.000335 | -0.496 | 700909 |
| Yoruba | Colombia_SB_HG | I10932 | FORM001 | -0.00014 | -0.00046 | -0.305 | 547359 |
| Yoruba | Colombia_SB_HG | I10932 | Colombia_Cundinamarca | 0.000031 | -0.000317 | 0.097 | 712547 |
| Yoruba | Colombia_SB_HG | I10932 | Colombia_Terregrande | 0.000091 | -0.000581 | 0.157 | 140109 |
| Yoruba | Colombia_SB_HG | I10932 | Colombia_Boyacá1_LaSalina | 0.000098 | -0.00033 | 0.299 | 711103 |
| Yoruba | Colombia_SB_HG | I10932 | Colombia_Boyacá2 | 0.000172 | -0.000323 | 0.531 | 711701 |
| Yoruba | Colombia_SB_HG | I10932 | Colombia_Terreros | 0.000244 | -0.000342 | 0.712 | 680611 |

An interpretation is that FORM001, Colombia_Terregrande, and I10932 have some ancestry that is not well-characterized by the available reference data and cannot be well discerned using this reference set in *qpAdm*. This is further supported by the multiple fitting one-source models for I10932 (discussed above), which suggests we have poor resolution to robustly model the ancestry of this individual. One way to test this interpretation is to set Panama_IsthmoColumbian_PreColonial as Source 1 and rotate all ancient and modern American populations into the position of Source 2 to assess if Colombia_SB_HG is an exclusively fitting second source for FORM001, Colombia_Terregrande, and I10932.

Based on the signal of Chibchan-related ancestry on the Altiplano Cundiboyacense during the Herrera-Muisca Period observed in $f_4$-statistics combined anthropological, archaeological, and linguistic evidence of Chibchan influence in this region, and also to test whether FORM001, Colombia_Terregrande, and I10932 share excess alleles exclusively with Colombia_SB_HG, we next set Panama_IsthmoColumbian_PreColonial as Source 1 (we selected this proxy so that we could include Cabécar in the right reference set for improved discretionary power) and rotated through other ancient and modern American populations as Source 2 (**Supplementary Figure 7.4**).

**Supplementary Figure 7.4.** Matrix of results for two-source ancestry models for Herrera-Muisca Period targets with Panama_IsthmoColumbian_PreColonial set as Source 1. Shaded cells represent models with p-values >0.05 where ancestry proportions are between 0-1 for both sources, standard errors are between 0-1, and ancestry proportions ± 1 standard error do not go outside of the 0-1 range. In other words, these are models that are consistent with the data.

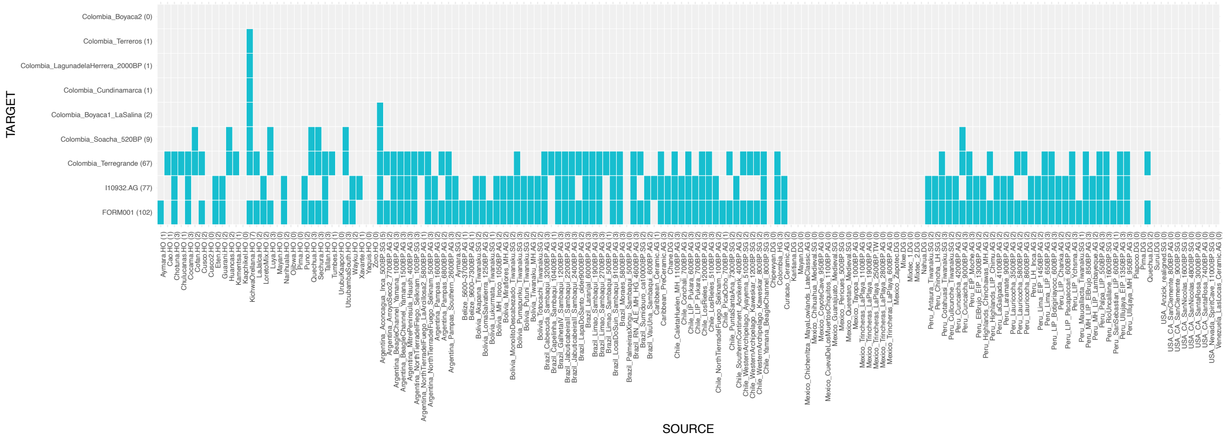


We find that the regional group of Colombia_Cundinamarca as well as the groups from Colombia_Terreros and Colombia_LagunadelaHerrera_2000BP can be fit exclusively as a two-source mixture of ancient Panama-related ancestry and ancestry related to a population with ancestry like that proxied by Kichwa Orellana. This latter ancestry has been characterized as "Amazonian North"-related, found in northern Ecuador and populations from the eastern slope of the northern Andes in Colombia[137]. This model also fit for Colombia_Boyacá1_LaSalina, Colombia_Soacha_520BP, Colombia_Terregrande, and FORM001, although not exclusively, but was a poor fit for Colombia_Boyacá2 (p=0.006) and I10932 (p=0.00001).

Colombia_Cundinamarca can be exclusively fit as having 53.9%±1.4% ancient Panama-related ancestry and 46.1%±1.4% Kichwa Orellana-related ancestry (p=0.95), while Colombia_Terreros and Colombia_LagunadelaHerrera_2000BP were fit as having 54.7%±2.3% and 54.2%±2.6% ancient Panama-related ancestry and the rest Kichwa Orellana-related ancestry (p=0.24 and 0.41, respectively) (**Supplementary Figure 7.5**). All of these sites are located within the Cundinamarca department. This is consistent with the Herrera-Muisca Period population in the Cundinamarca department being a mixture of Chibchan-related ancestry and ancestry related to that found at the Andean-Amazonian interface.

**Supplementary Figure 7.5.** Plot of ancestry models for Herrera-Muisca Period targets with Panama_IsthmoColumbian_PreColonial and Kichwa_Orellana as sources. Color indicates targets for which model p-values are >0.05, while gray indicates targets for which model p-values are <0.05. Error bars represent 1 standard error.

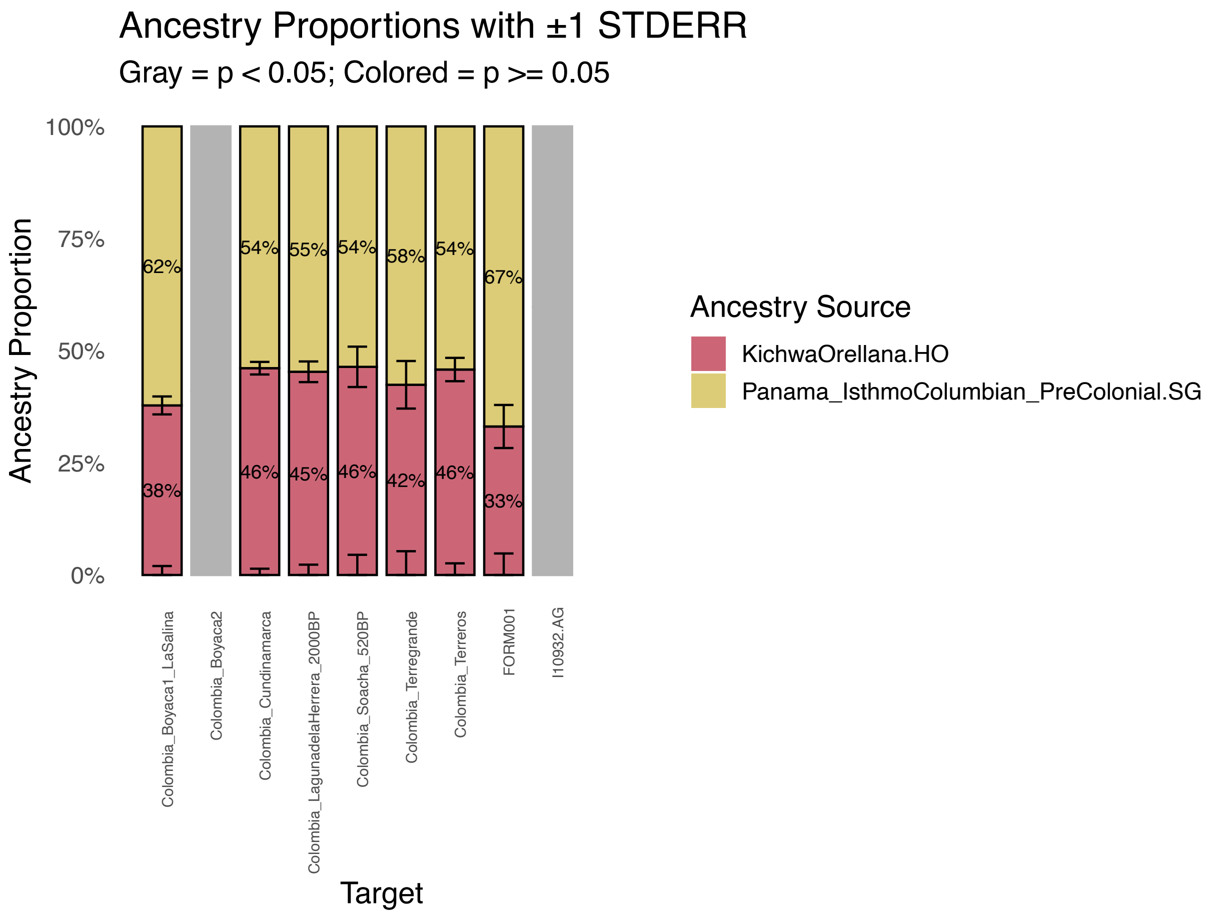


Colombia_Boyacá1_LaSalina was fit as having 62.2%±2.0% ancient Panama-related ancestry and 37.8%±2.0% Kichwa Orellana-related ancestry (p=0.73), while a model with 54.4%±3.2% Panama-related ancestry and 45.6%±3.2% Argentina_Aconcagua_Inca_500BP-related ancestry also fit (p=0.05) (**Supplementary Figure 7.5**). Fitting models for Colombia_Soacha_520BP included a second source with ancestry from the western Andes or circa-Andean region, but we were not able to discern a specific location for this ancestry (populations fitting as Source 2 in our models had either Amazonia North (Kichwa Orellana, Cofan), north coast Peru (Sechura), north highland Peru (Huancas, Utucubamba South, Luya), or central Southern Andes (Quechua) ancestry; Peru_Cuncaicha_9000BP and Argentina_Aconcagua_Inca_500BP also fit as Source 2 at p>0.05). There were many fitting models for Colombia_Terregrande, I10932, and FORM001, suggesting that we have little ability to meaningfully discern ancestry for these individuals with the data currently available.

There were no fitting two-source models for the Colombia_Boyacá2 regional group. To investigate the ancestry of this group in greater detail, we began by removing each of the right reference groups one-by-one to see if model fit improved, using this approach to identify outgroups that revealed additional ancestry complexity beyond a simple two-source framework (**Supplementary Table 7.4**).

**Supplementary Table 7.4.** Table of p-values for two-source ancestry models of Panama_IsthmoColumbian_PreColonial + KichwaOrellana for Colombia_Boyacá2 as target.

| **Right group removed** | **2 source model p-value** |
|---|---|
| Chipewyan | 0.02 |
| Zapotec | 0.03 |
| Mixe | 0.01 |
| Mixtec | 0.003 |
| Surui | 0.004 |
| Cabecár | 0.004 |
| Piapoco | **0.13** |
| Karitiana | 0.004 |
| Inga | 0.004 |
| Wayuu | 0.004 |
| Apalai | 0.01 |
| Arara | 0.006 |

We found that removing Piapoco from the right reference set improved model fit past a threshold of $p>0.05$, consistent with the signal of attraction between Colombia_Boyacá2 and Piapoco especially relative to Colombia_Boyaca1_LaSalina that we observe with $f_4$-statistics (**Table S4**). When Piapoco is removed from the reference set, Colombia_Boyacá2 can be fit as having 59.2%±1.7% ancient Panama-related ancestry and the remaining 40.8% ± 1.7% Kichwa Orellana-related ancestry, the exclusively fitting model for this target (**Supplementary Figures 7.6** and **7.7**). This shift from a non-fitting to fitting model is because *qpAdm* relies on the reference set to distinguish the ancestry sources from one another. Each reference population contributes axes of allele-frequency drift against which the model is evaluated. If a model is a poor fit with the full reference set but model fit improves when a reference population is removed, the improvement can be interpreted as reflecting a loss of discriminatory power or as suggesting that the original (rejected) model does not capture all the ancestry components present in the target.

**Supplementary Figure 7.6.** Matrix of results for two-source ancestry models for Herrera-Muisca Period targets with Panama_IsthmoColumbian_PreColonial set as Source 1 and Piapoco removed from the reference set. Shaded cells represent models with p-values >0.05 where ancestry proportions are between 0-1 for both sources, standard errors are between 0-1, and ancestry proportions ± 1 standard error do not go outside of the 0-1 range. In other words, these models are consistent with the data.

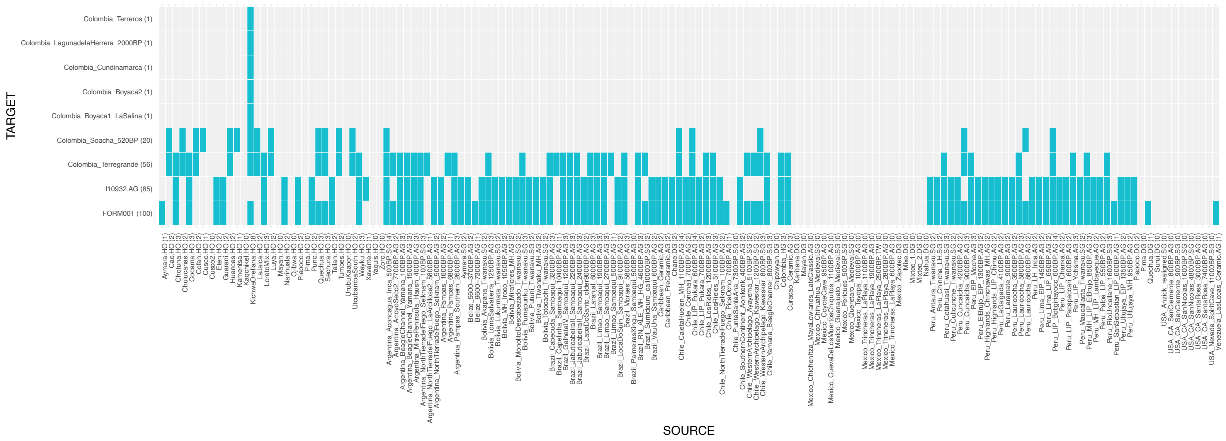


**Supplementary Figure 7.7.** Plot of ancestry models for Herrera-Muisca Period targets with Panama_IsthmoColumbian_PreColonial and Kichwa Orellana as sources and Piapoco removed from the reference set. Color indicates targets for which model p-values are >0.05, while gray indicates targets for which model p-values are <0.05.

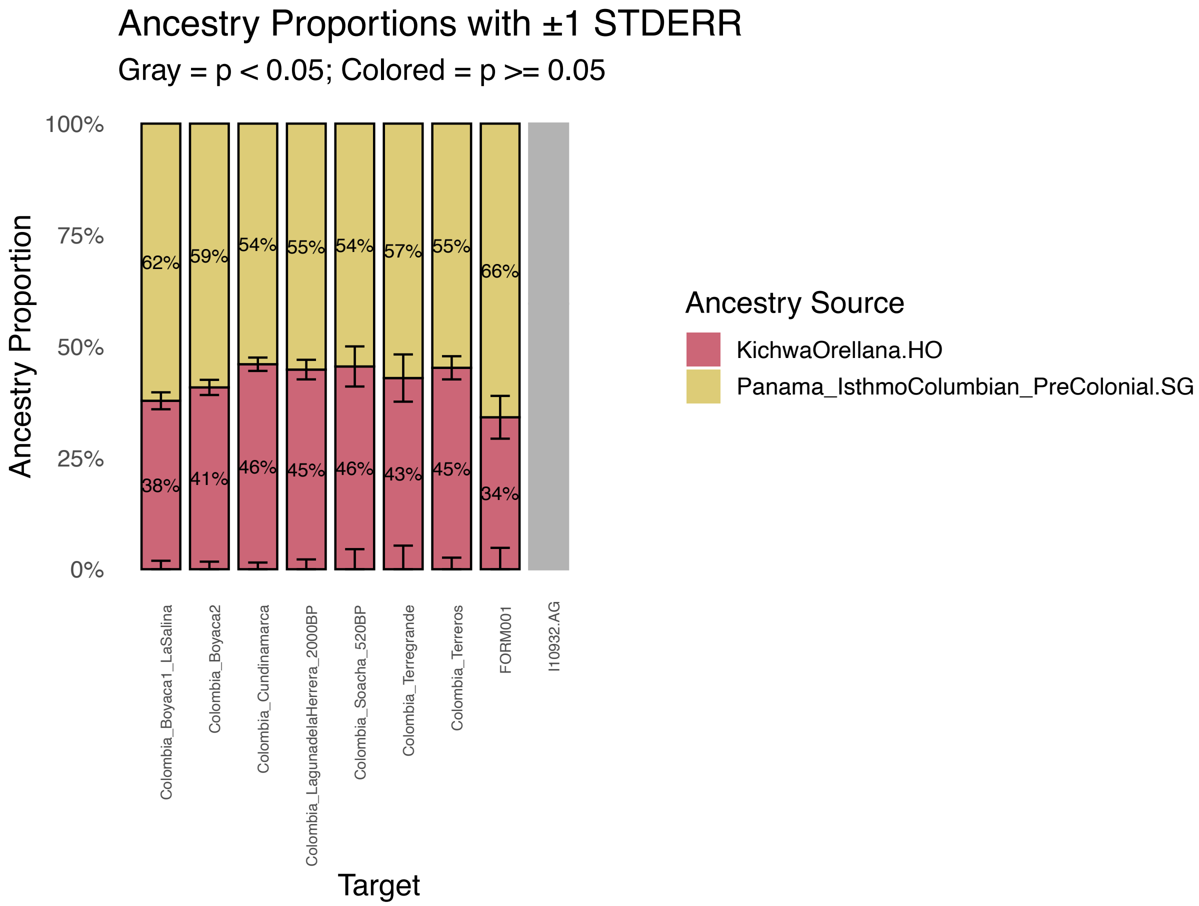


Based on these results, a question was whether there was quantifiable ancestry well-proxied by Piapoco or another source in the gene pool of Colombia_Boyacá2. To test this, we first set

Colombia_Boyaca1_LaSalina or Colombia_Cundinamarca as Source 1 and rotated through all ancient and modern Americans as Source 2 with the full base reference set but found no fitting models at p>0.05. We removed Piapoco from the right reference set and re-ran this analysis but still identified no fitting models at p>0.05.

##### Three-source models

To test the hypothesis that a model fitting the ancestry in Colombia_Boyacá2 requires three-sources, we set Panama_IsthmoColumbian as Source1, Kichwa Orellana as Source 2, and rotated through all ancient and modern Americans as Source 3 with the full base reference set. There were no fitting models at p>0.05. We removed Piapoco from the right reference set and tested the same three-source models. Without this right population, there were 41 fitting models for Colombia_Boyacá2 at p>0.05 (**Supplementary Figure 7.8**); however, these fitting models were non-unique: a wide range of Native American populations fit as Source 3 while yielding very similar estimates for the two major ancestry components (58%–60% Panama_IsthmoColumbian_PreColonial-related and ~40%-42% KichwaOrellana-related). This suggests that acceptance reflects reduced constraint in the absence of Piapoco in the right reference set rather than strong evidence for a particular third ancestry source. We therefore interpret these results as consistent with Colombia_Boyacá2 being well-described primarily by Panama- and KichwaOrellana-related ancestry, while the identity of any minor additional component is not well resolved.

**Supplementary Figure 7.8.** Matrix of results for three-source ancestry models for Colombia_Boyacá2 with ancient Panamanians set as Source 1 and Kichwa Orellana set as Source 2 and Piapoco.HO removed from the reference set. Shaded cells represent models with p-values >0.05 where ancestry proportions are between 0-1 for both sources, standard errors are between 0-1, and ancestry proportions ± 1 standard error do not go outside of the 0-1 range. In other words, these models are consistent with the data.

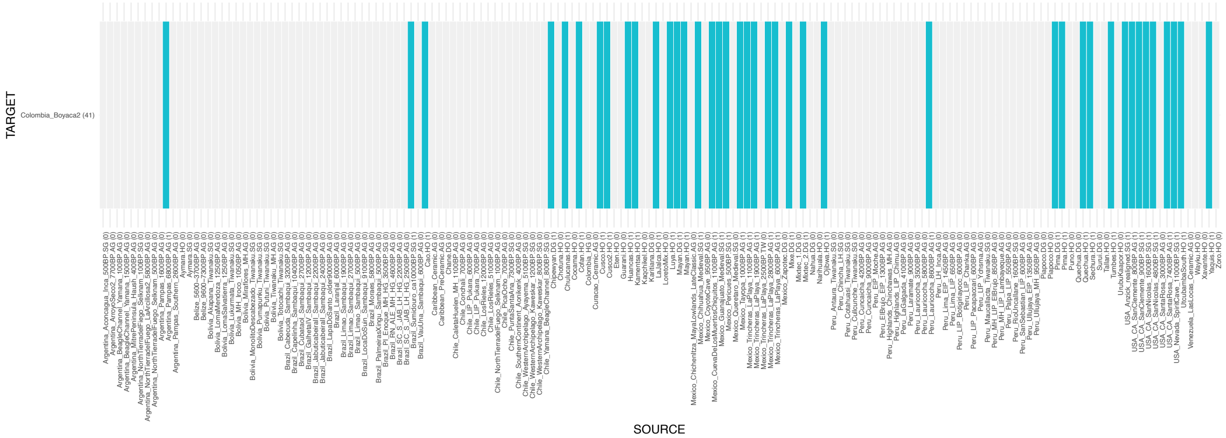

**Confirming model robustness**

To evaluate the robustness of our working models—particularly the use of Kichwa Orellana as a source from the Amazonian/Andean interface that, together with a Chibchan-related group, can be used in a two-source model to explain a major part of the ancestry found in the regional groups of the Altiplano Cundiboyacense in the Herrera-Muisca Periods—we carried out additional tests.

First, we fixed Kichwa Orellana as Source 1 and rotated every ancient and present-day American population into the Source 2 position to assess whether the combination of Kichwa Orellana-related and Panama-related ancestry was uniquely supported by the data. A model with Panama_IsthmoColumbian as Source 2 was an exclusive fit for Colombia_Cundinamarca and Colombia_Boyacá1_LaSalina (no two-source model fit Colombia_Boyacá2 when the full right reference set was used, consistent with our earlier results) (**Supplementary Figure 7.9**).

**Supplementary Figure 7.9.** Matrix of results for two-source ancestry models for Herrera-Muisca Period targets with Kichwa Orellana set as Source 1. Shaded cells represent models with p-values >0.05 where ancestry proportions are between 0-1 for both sources, standard errors are between 0-1, and ancestry proportions ± 1 standard error do not go outside of the 0-1 range. In other words, these are models that are consistent with the data.

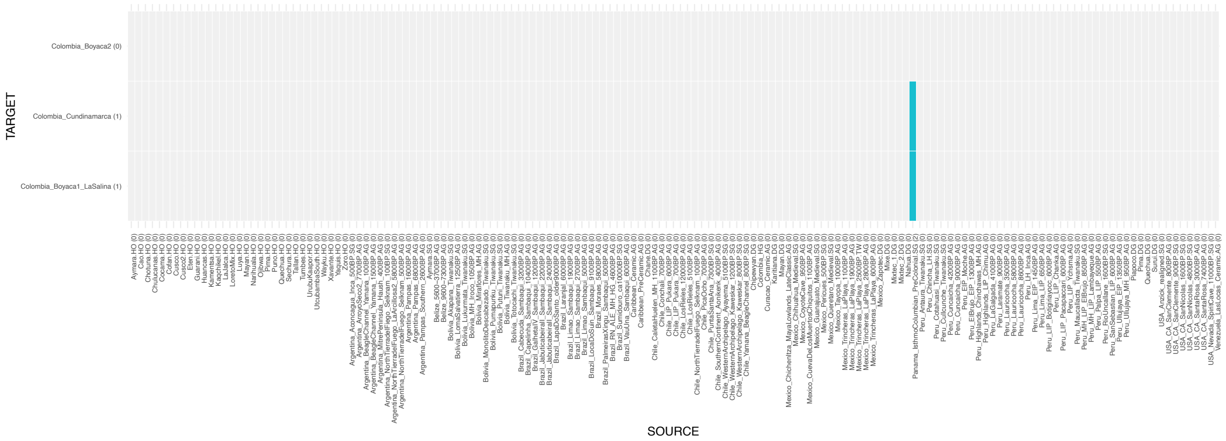


Second, we tested robustness by systematically manipulating the right reference set. We first added Kichwa Orellana to the base right reference set. When Kichwa Orellana is on the right, no model is a good fit for Colombia_Cundinamarca or Colombia_Boyacá1_LaSalina, indicating that the signal for Kichwa Orellana-related ancestry is not an artifact of outgroup choice.

Next, we implemented a rotation strategy in which South American populations from the right reference set were removed one at a time and tested as potential sources in two-source models with ancient Panamanians fixed as Source 1. Each test was performed both with and without Kichwa Orellana in the right reference set. This approach ensures that no suitable source population is inadvertently excluded from being tested as a possible source population and allows us to more comprehensively assess whether alternative ancestry combinations could account for the data.

Removing Surui, Karitiana, Apalai, Arara, or Wayuu from the right had no effect on model fits: a model with ancient Panamanians and Kichwa Orellana still fit exclusively at p<0.05 for Colombia_Cundinamarca and Colombia_Boyacá1_LaSalina. None of these populations fit in a model as Source 2, either with or without Kichwa Orellana in the right reference set. As discussed in more detail above, removing Piapoco from the reference set has no effect on the fit of models for Colombia_Cundinamarca and Colombia_Boyacá1_LaSalina (ancient Panamanians + Kichwa Orellana was still an exclusively fitting model), but its exclusion from the right shifts the fit of a two-source model for Colombia_Boyacá2 from non-fitting to fitting.

If we remove Cabécar from the right reference set, a model with Cofan as Source 2 also fits only for Colombia_Boyacá1_LaSalina (p=0.07) (**Supplementary Figure 7.10**).

**Supplementary Figure 7.10.** Matrix of results for two-source ancestry models for Herrera-Muisca Period targets with Panama_IsthmoColumbian_PreColonial.SG set as Source 1 and Cabécar removed from the right reference set. Shaded cells represent models with p-values >0.05 where ancestry proportions are between 0-1 for both sources, standard errors are between 0-1, and ancestry proportions ± 1 standard error do not go outside of the 0-1 range. In other words, these are models that are consistent with the data.

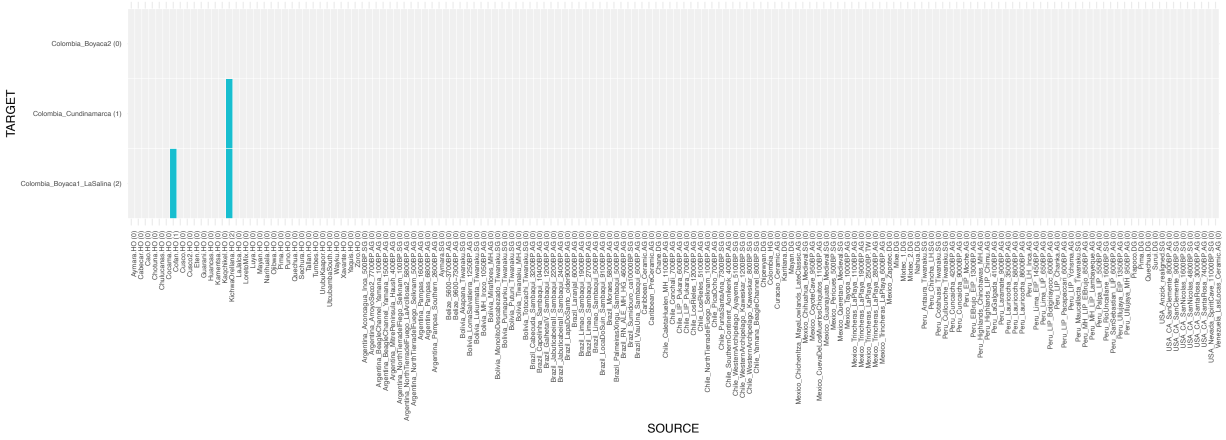


When we add Kichwa Orellana to the right, a two-source model of Panama + Cofán still fits Colombia_Boyacá1_LaSalina (p=0.07). However, when we re-add Cabécar to the right and rerun Panama + Cofán, the fit disappears (p=7.4e-05), showing that the fit was contingent on a weaker right reference set. While under reduced Chibchan discrimination, the Amazonian/Andean-related ancestry in Colombia_Boyacá1_LaSalina can be proxied by other northern Andean lineages, including a Chibchan outgroup restores the specificity of the Panama + Kichwa Orellana model and eliminates the Cofán alternative.

Removing Inga from the right reference reduced the Andean/Amazonian contrast that distinguishes the sources, and multiple otherwise incompatible two-source models passed for Colombia_Cundinamarca and Colombia_Boyacá1_LaSalina, indicating that Inga adds discriminative power necessary for rejecting models (**Supplementary Figure 7.11**).

**Supplementary Figure 7.11.** Matrix of results for two-source ancestry models for Herrera-Muisca Period targets with Panama_IsthmoColumbian_PreColonial.SG set as Source 1 and Inga removed from the right reference set. Shaded cells represent models with p-values >0.05 where ancestry proportions are between 0-1 for both sources, standard errors are between 0-1, and ancestry proportions ± 1 standard error do not go outside of the 0-1 range. In other words, these are models that are consistent with the data.

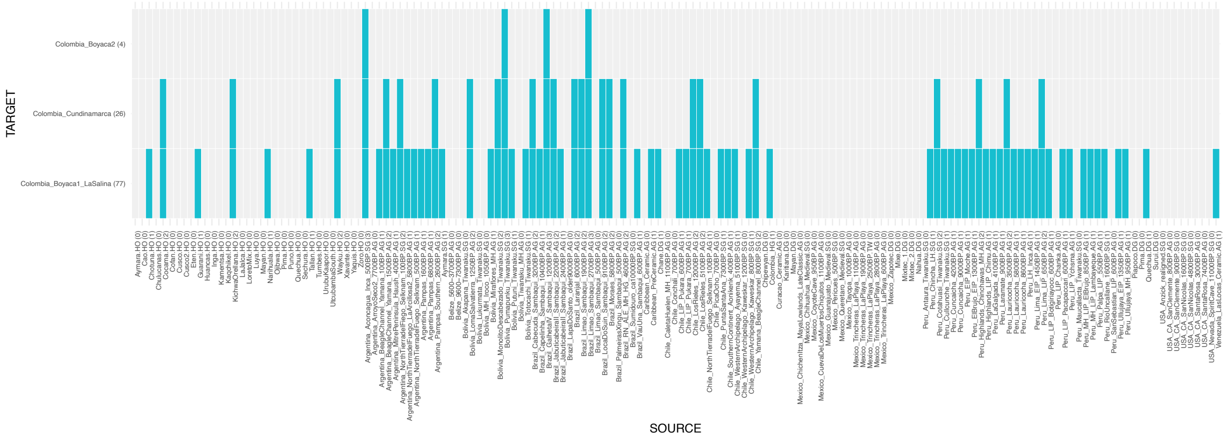


Although Inga provides critical discriminatory power when included in the right reference set it does not serve as a fitting proxy source for ancestry in Colombia_Cundinamarca or Colombia_Boyacá1_LaSalina, possibly due to population-specific drift or postcolonial admixture.

To assess whether including Inga on the right might bias our results, we substituted it with the genetically similar Cofán, another 'Amazonian North' population[137] and re-ran all two-source models with Panama fixed as Source 1 (**Supplementary Figure 7.12**). If the Panama + Kichwa Orellana model were truly robust, this substitution should not alter its status as the only fitting model for Colombia_Cundinamarca and Colombia_Boyacá1_LaSalina. Consistent with this expectation, Panama + Kichwa Orellana remained the exclusive fit for both groups when Cofán replaced Inga in the reference set.

**Supplementary Figure 7.12.** Matrix of results for two-source ancestry models for Herrera-Muisca Period targets with ancient Panama set as Source 1 and Inga replaced by Cofán in the right reference set. Shaded cells represent models with p-values >0.05 where ancestry proportions are between 0-1 for both sources, standard errors are between 0-1, and ancestry proportions ± 1 standard error do not go outside of the 0-1 range. In other words, these are models that are consistent with the data.

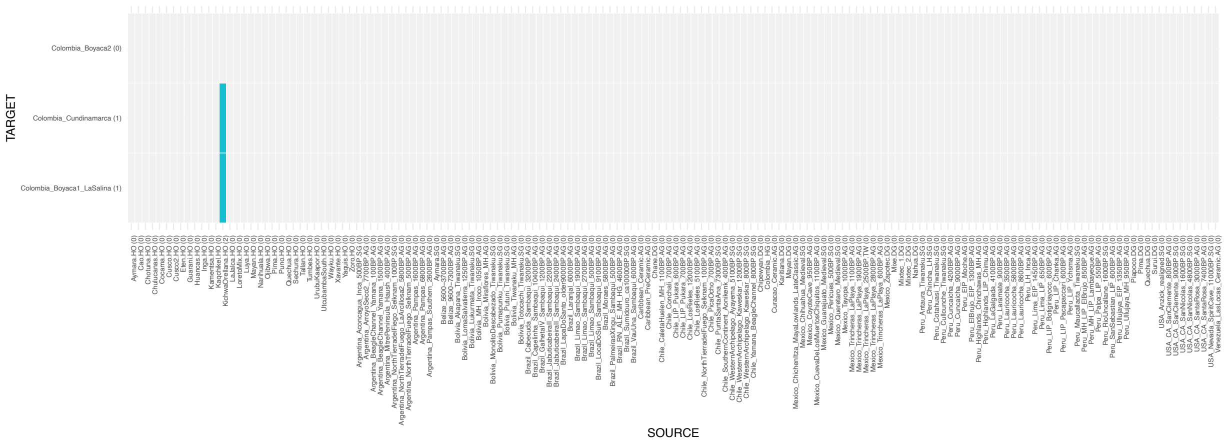


In contrast, if we replace Inga with Kichwa Orellana, no models fit. Inga or Cofán (though geographically proximate and genetically similar) cannot therefore be considered a more appropriate proxy source for the Andean/Amazonian ancestry than Kichwa Orellana. The absence of valid fits under right reference set rotations supports the specificity of the Panama + Kichwa Orellana model and reduces the likelihood that the signal reflects overfitting.

A question is if the signal of Amazonian/Andean-related ancestry proxied by Kichwa Orellana in the populations of the Altiplano Cundiboyacense is related to shared Chibchan-related ancestry. However, our robustness tests suggest that this is unlikely to be the case. Across multiple *qpAdm* rotations, the two-source model of Panama_IsthmoColumbian_PreColonial (Chibchan-related) + Kichwa Orellana (Amazonian/Andean-related) was the only consistently fitting solution. When Kichwa Orellana was added to the right reference set, all models failed. Alternative models with northern Andean or Amazonian groups (e.g., Cofán, Inga) only fit when Chibchan discriminators such as Cabécar were removed, and these fits collapsed once those Chibchan groups were restored. Taken together, these results strongly support the presence of an Amazonian-related ancestry stream, well proxied by present-day Kichwa Orellana, that is distinct from Chibchan-related ancestry in the Herrera-Muisca Period populations. We emphasize, however, that present-day Kichwa Orellana are used here as a proxy: the true source of this ancestry could have been an unsampled population from other parts of northern Amazonia or the Andean foothills.
